# Back-pocket optimization of 2-aminopyrimidine-based macrocycles leads to potent dual EPHA2/GAK kinase inhibitors with antiviral activity

**DOI:** 10.1101/2024.02.18.580805

**Authors:** Joshua Gerninghaus, Rezart Zhubi, Andreas Krämer, Marwah Karim, Do Hoang Nhu Tran, Andreas C. Joerger, Christian Schreiber, Lena M. Berger, Benedict-Tilman Berger, Theresa A.L. Ehret, Lewis Elson, Christopher Lenz, Krishna Saxena, Susanne Müller, Shirit Einav, Stefan Knapp, Thomas Hanke

## Abstract

Macrocyclization of acyclic compounds is a powerful strategy for improving inhibitor potency and selectivity. Here, we developed a 2-aminopyrimidine-based macrocyclic dual EPHA2/GAK kinase inhibitor as a chemical tool to study the role of these two kinases in viral entry and assembly. Starting with a promiscuous macrocyclic inhibitor, **6**, we performed a structure-guided activity relationship and selectivity study using a panel of over 100 kinases. The crystal structure of EPHA2 in complex with the developed macrocycle **23** provided a basis for further optimization by specifically targeting the back pocket, resulting in compound **55** as a potent dual EPHA2/GAK inhibitor. Subsequent front-pocket derivatization resulted in an interesting *in cellulo* selectivity profile, favoring EPHA4 over the other ephrin receptor kinase family members. The dual EPHA2/GAK inhibitor **55** prevented dengue virus infection of Huh7 liver cells, mainly via its EPHA2 activity, and is therefore a promising candidate for further optimization of its activity against dengue virus.

## INTRODUCTION

Macrocyclization of acyclic compounds serves as a powerful tool in medicinal chemistry that recently gained increasing attention.^1^ The structural preorganization of these rigid molecules ideally locks their bioactive conformation, resulting in several beneficial properties, including enhanced potency, selectivity, and pharmacokinetics.^2–5^ The most recent examples in the field of kinase inhibitors include the approval of the macrocyclic ALK/Ros1 inhibitor lorlatinib (Lorbrena^®^) and the JAK2/IRAK1 inhibitor pacritinib (Vonjo^®^) by the U.S. Food and Drug Administration (FDA)^6,7^, highlighting the growing importance of macrocyclic compounds in drug development. Particularly, for a protein family characterized by a highly conserved binding pocket, such as protein kinases, the approach of developing macrocyclic kinase inhibitors offers a promising avenue to rapidly achieve selectivity within this family.^8^

The ephrin receptors (EPH), short for erythropoietin-producing human hepatocellular receptors, are transmembrane tyrosine kinases that represent the largest subfamily of receptor tyrosine kinases (RTK). These transmembrane proteins consist of an N-terminal ligand-binding domain (LBD), followed by a cysteine-rich region and repeats of fibronectin type III on the extracellular side, linked by a transmembrane helix to an intracellular kinase domain, a sterile alpha motif (SAM) domain and the PDZ-binding motif at the C-terminus. The 14 human EPHs can be classified into nine type-A and five type-B receptors, depending on their ligand binding preferences for cell-bound type-A or type-B ephrins. EPH kinases play a central role in a variety of cellular processes such as cell positioning, tissue and organ patterning, cell survival, proliferation, and development.^9–12^ They enable bidirectional cell-to-cell communication through forward signaling in EPH-bearing cells and reverse signaling in the respective ephrin-bearing cells.^13^ Co-expression of different EPHs and ephrins on the surface of the same cell leads to complex clustering and crosstalk with different signaling pathways.^14,15^ Due to the widespread expression of EPHs and their involvement in numerous physiological processes, dysregulation of EPHs has been associated with the development of cardiovascular, oncological, and neurological diseases, as well as viral infections.^16,17^ For example, Lupberger *et al.* reported that EPHA2 acts as a host factor for hepatitis C virus (HCV) entry by controlling the formation of essential CD81-CLDN1 co-receptor complexes.^18^ HCV belongs to the *Flaviviridae* family, a group of single-strand RNA viruses, that comprises other prominent representatives such as the yellow fever virus, zika virus, and dengue virus (DENV). Transmitted by mosquitoes of the genus *Aedes*, DENV is the causative agent of dengue fever. With an estimated 390 million infections per year worldwide, DENV poses a major health threat to people living in endemic regions.^19^ Up to five percent of symptomatic individuals progress within several days postinfection to severe dengue, characterized by an acute shock syndrome, bleeding, organ impairment, and sometimes death.^20^

Cyclin-G associated kinase (GAK) is another well-known kinase suitable as a target for antiviral therapy, relevant not only in the context of HCV but also in addressing DENV infections.^21,22^ This serine/threonine protein kinase is widely expressed in the cell and is primarily localized in the perinuclear region and the Golgi apparatus.^23^ GAK plays a pivotal role as a regulator of clathrin-mediated intracellular trafficking of cellular cargo proteins.^24–26^ GAK facilitates viral entry by phosphorylating the clathrin adaptor protein complex 2 (AP-2), which together with other endocytotic factors promotes endocytosis.^27^ In addition, GAK regulates AP2M1 (µ subunit of AP-2), thereby stimulating its binding to cargo proteins that are essential for viral assembly.^28^

Given the pathophysiological relevance of EPHA2 and GAK to the mechanisms underlying viral entry and assembly, we hypothesized that the dual inhibition of both kinases could be effective against viral infection. We were particularly interested in addressing DENV infection, as current treatment focuses primarily on symptomatic relief (e.g., antipyretic and antiphlogistic agents) and no targeted antiviral therapy is currently available.^29^ To achieve this goal, we chose a macrocyclic inhibitor scaffold which potently but unselectively inhibits EPHA2 and GAK. By introducing variations in moieties interacting with the kinase back pocket and simultaneously monitoring kinase off-targets and other members of the EPH family, we developed a potent dual inhibitor with significantly improved selectivity that can serve as a lead for further translational studies of this dual targeting strategy.

The macrocyclic diaminopyrimidine **6** (**Figure 1C**) was reported by Lücking *et al.* as dual CDK2 and VEGF-R inhibitor.^30^ Due to its low molecular weight of 384.25 g/mol and the presence of a bromine in position 5 of the pyrimidine core as chemical handle, compound **6** was considered as an ideal starting point for our target-oriented SAR studies. The crystal structure of **6** in complex with CDK2 (PDB: 2J9M) served as the basis of our structure-guided design strategy.^30^ Macrocycle **6** binds to the hinge region of CDK2 by interacting with the amide backbone of L83 via two hydrogen bonds, formed by the 2-aminopyrimidine motif (**Figure 1A**). The bromine residue in position 5 of the pyrimidine points towards the gatekeeper residue, F80, which restricts access to the hydrophobic back pocket due to its relatively large size. We hypothesized that derivatization in position 5 of the pyrimidine ring would increase the selectivity of macrocycle **6** by extending the inhibitor into the hydrophobic back pocket located between the gatekeeper and the αC-helix, and thus kinases that contain larger gatekeeper residues, such as a phenylalanine in CDK2, should no longer be targeted by **6**. Since both kinases, EPHA2 and GAK, have smaller gatekeeper residues, this structure-based design was expected to significantly increase selectivity in our representative panel of approximately 100 kinases (**Figure 1D**).

**Figure 1.**
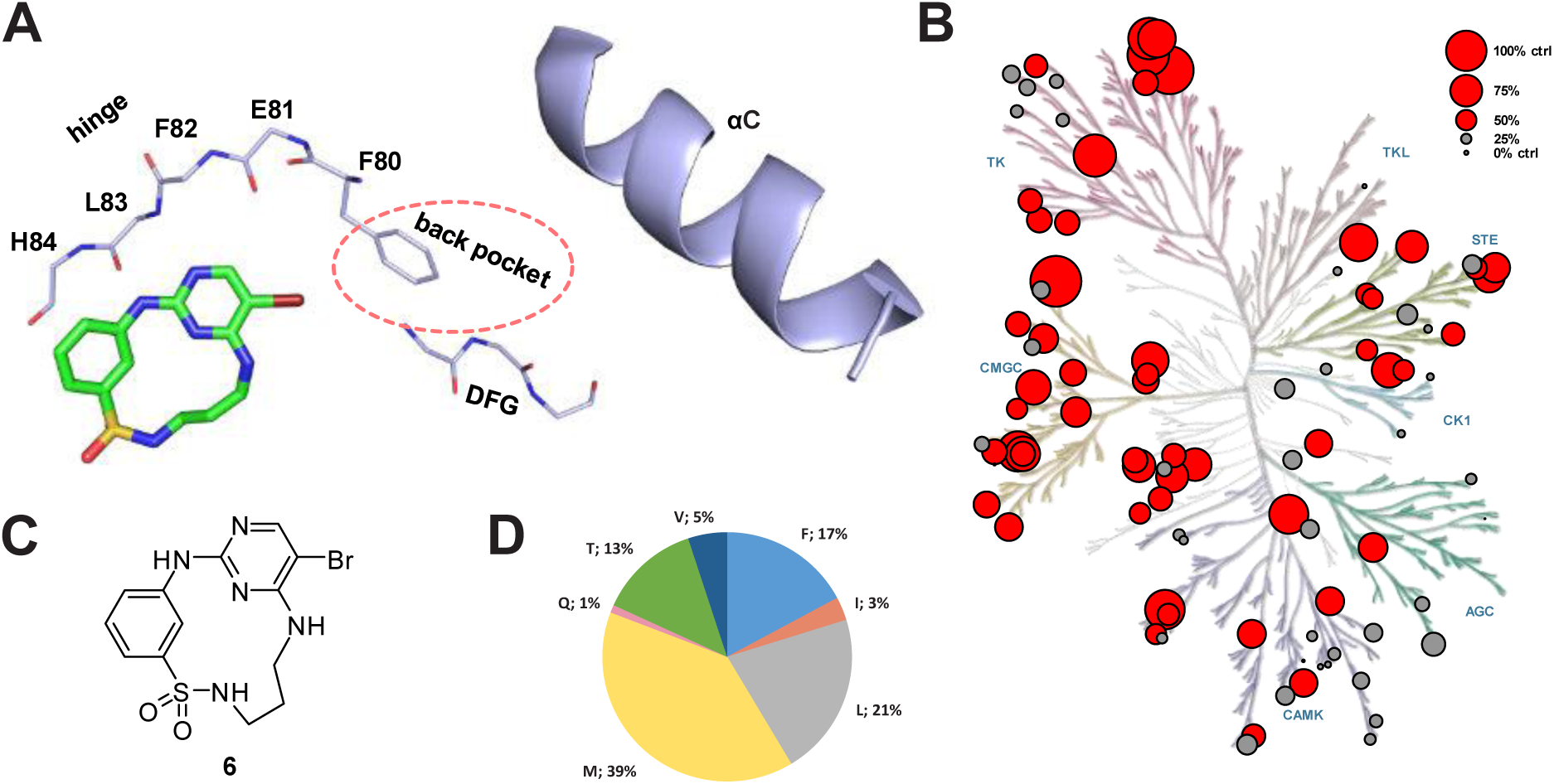
Starting point **6** is a promiscuous macrocyclic kinase inhibitor. **(A)** Macrocyclic compound **6** co-crystalized with CDK2 (PDB: 2J9M). Structure-guided design strategy illustrating the targeted back-pocket region, guarded by the gatekeeper residue F80. **(B)** Phylogenetic kinome tree depicting the screened kinases for **6** as dots of different size, relative to the thermal shift of a control compound in K set as 100%, hits lower than 50% control in gray, hits higher than 50% control in red. AGC (protein kinase A, G, C), CAMK (calcium/calmodulin-dependent kinases), CK1 (casein kinase 1), CMGC (cyclin-dependent kinases, MAP kinases, glycogen synthase kinases, casein kinase 2), STE (homologues of yeast sterile 7, 11, 20), TK (tyrosine kinases) and TKL (tyrosine kinase-like)-family. **(C)** Chemical structure of **6**. **(D)** Distribution of gatekeeper residues throughout the kinase panel used for screening as a pie chart, gatekeeper as single letter code next to the distribution in percent. The panel is representative of the whole kinome in terms of gatekeeper distribution.^31^

## RESULTS AND DISCUSSION

First, we resynthesized macrocycle **6** as a reference for our SAR studies. Except for the reduction of the aromatic nitro group, the published synthetic route already demonstrates very good to excellent yields. Lücking *et al.* used titanium(III) chloride in HCl as a reducing agent, which resulted in a yield of 49% of the desired aniline macrocyclization precursor (**5**).^30^ In order to improve the yield of this key reaction, we explored the Béchamp reduction process using iron under slightly acidic conditions as this reaction typically offers high selectivity for nitroaryls under relatively mild conditions.^32^ With this strategy, we were able to increase the yield of the aryl nitro reduction to an excellent yield of 90%. The synthetic route was conducted in six steps. In the first step, 5-bromo-2,4-dichloropyrimidine underwent a nucleophilic aromatic substitution at position 4 with *N*-Boc-1,3-diaminopropane. The intermediate was subsequently reacted with 4 N HCl in 1,4-dioxane to yield the hydrochloride **2** in quantitative yield. Next, the sulfonamide **4** was formed by the reaction of **2** with 3-nitrobenzenesulfonyl chloride (**3**) under basic conditions. The subsequent Béchamp reaction of **4** yielded aniline **5**. Macrocyclization was achieved by intramolecular nucleophilic aromatic substitution of **5**, yielding macrocycle **6** in an excellent yield of 90% even on gram scale. In the last step, **6** was subjected to a Suzuki cross-coupling reaction with the respective boronic acids or boron pinacol esters, yielding the final products **7-23** and **41-63**.

Because of its dual hydrogen-bond interaction with the amide backbone, the aminopyrimidine scaffold is a commonly used hinge-binding motif for the development of many kinase inhibitors, both in research and approved clinical drugs (e.g., brigatinib, osimertinib).^7,33–35^ However, an often-faced problem of this chemical scaffold is the lack of selectivity. To determine the selectivity profile of **6**, we screened this compound against our in-house panel of more than 100 kinases by differential scanning fluorimetry (DSF).^36^ A good stabilization in the DSF assay leads to a higher thermal shift, (Δ*T*_m_), which is usually related to a stronger binding. As expected, **6** showed promiscuous binding and interacted strongly with many members of the tested kinome (**Figure 1B**). Compound **6** stabilized 55 of 101 kinases, including our targets EPHA2 and GAK, by more than 50% of the thermal shift in K compared with a potent control compound. This promiscuous and strong binding of macrocycle **6** to a large number of the tested kinases may be due to its relatively small size and the strong hinge interaction.

Based on our initial data, we then installed various aromatic and heteroaromatic moieties in position 5 of the pyrimidine core to optimize the macrocycle towards the unoccupied back pocket, which then led to compounds **7-20 (Scheme 1B)**. Small unsubstituted (hetero-) cyclic residues such as 2-furyl (**7**), phenyl (**9**) and 3-pyridyl (**13**), or mono-substituted phenyls harboring functional groups in meta or para position (**8**, **10-12**, **14**), resulted in moderate to strong stabilization for fewer kinases than with compound **6**, indicating improved selectivity (**Figure 2**). Sterically demanding residues such as 5-(1-methylpyrazol)yl (**17**), 4-*tert*-butylphenyl (**18**) or biphenyls (**19, 20**) showed little to no temperature shifts (Δ*T*_m_) in our assay panel, suggesting that these moieties cannot be accommodated in the ATP pocket of kinases due to sterical clashes. Installing fluorine-methoxy-disubstituted phenyls **21-23 (Figure 2)** resulted in preferential binding to FGFR1-3, FLT1, ABL1, EPHA2/A4/A5/A7/B1/B3, and GAK. These kinases share a valine or threonine as a gatekeeper residue, confirming our hypothesis that targeting the small gatekeeper in these kinases should increase the selectivity of the scaffold. In particular, the 4-fluoro-3-methoxyphenyl substituted compound **23** showed a promising selectivity profile for the EPH kinases and GAK over FGFRs and FLT1. These findings encouraged us to further optimize the back-pocket interaction of **23**.

**Scheme 1.**
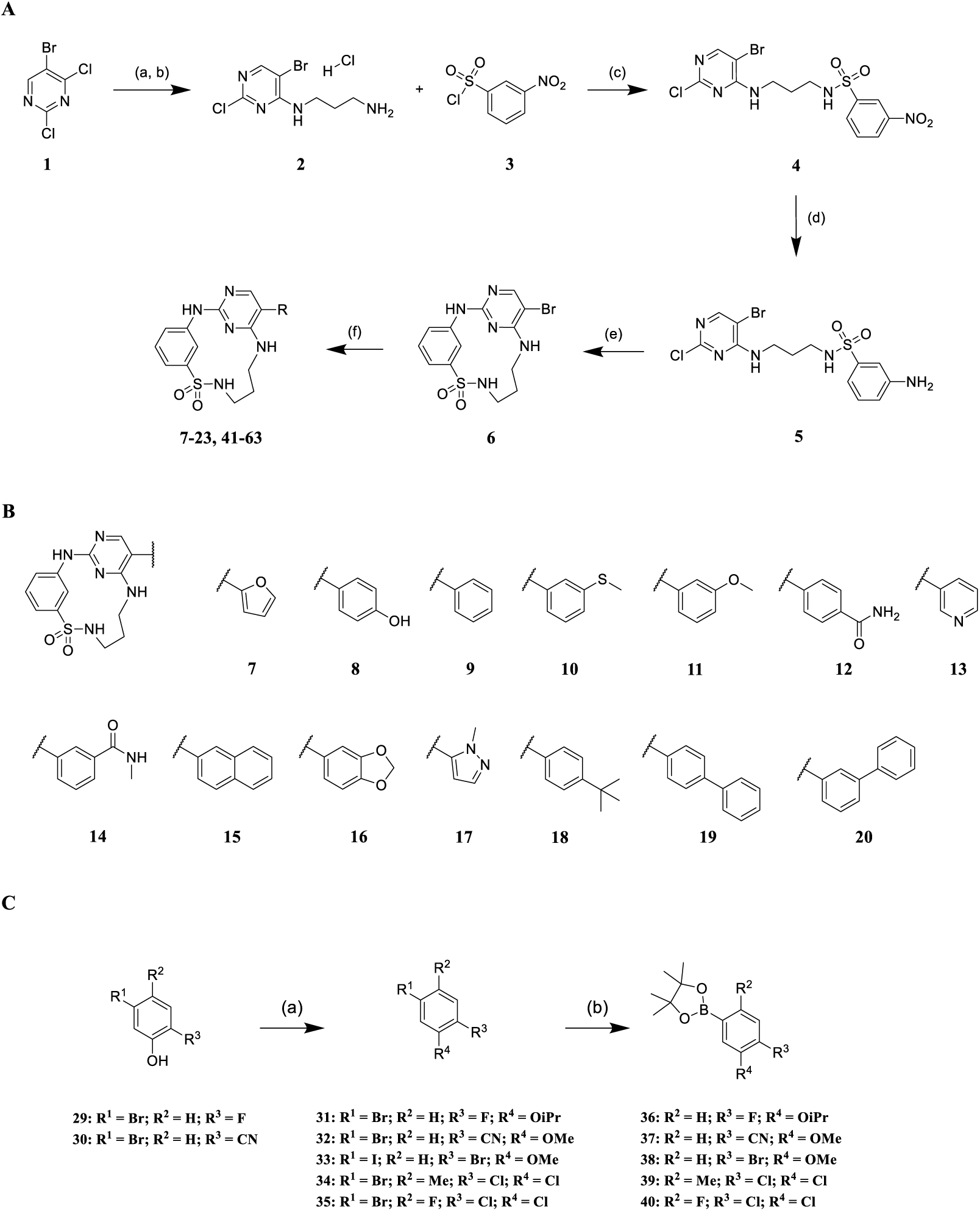
**(A):** Synthesis of macrocyclic inhibitors **7-23** and **41-63** according to Lücking *et al*.^30a^ **(B):** Back-pocket targeting moieties of derivatives **7-20**. **(C):** Synthesis of borpinacole esters **36-40**.^b^ ^a^Reagents and conditions: (a) *N*-Boc-1,3-diaminopropane, triethylamine, acetonitrile, rt, 3 h; (b) 4 N HCl in 1,4-dioxane, acetonitrile, rt, 3 h; (c) triethylamine, acetone/water (3:1), rt, 5 h; (d) iron, ammonium chloride, methanol/water (9:1), reflux, 3 h; (e) 4 N HCl in 1,4-dioxane, acetonitrile, water, 2-butanol, reflux, 6 h; (f) potassium carbonate, [1,1’-Bis(diphenylphosphino)-ferrocene]palladium(II) dichloride, boronic acid, 1,4-dioxane/dimethylformamide (1:1), 100 °C, µW, 2 h. ^b^Reagents and conditions: (a) 2-bromopropane (for 31) or iodomethane (for 32), cesium carbonate, dimethylformamide, rt, 16 h; (b) bis(pinacolato)diboron, [1,1’-Bis(diphenyl-phosphino)ferrocene]-palladium(II) dichloride, potassium acetate, 1,4-dioxane, 100 °C, µW, 2 h.

**Figure 2.**
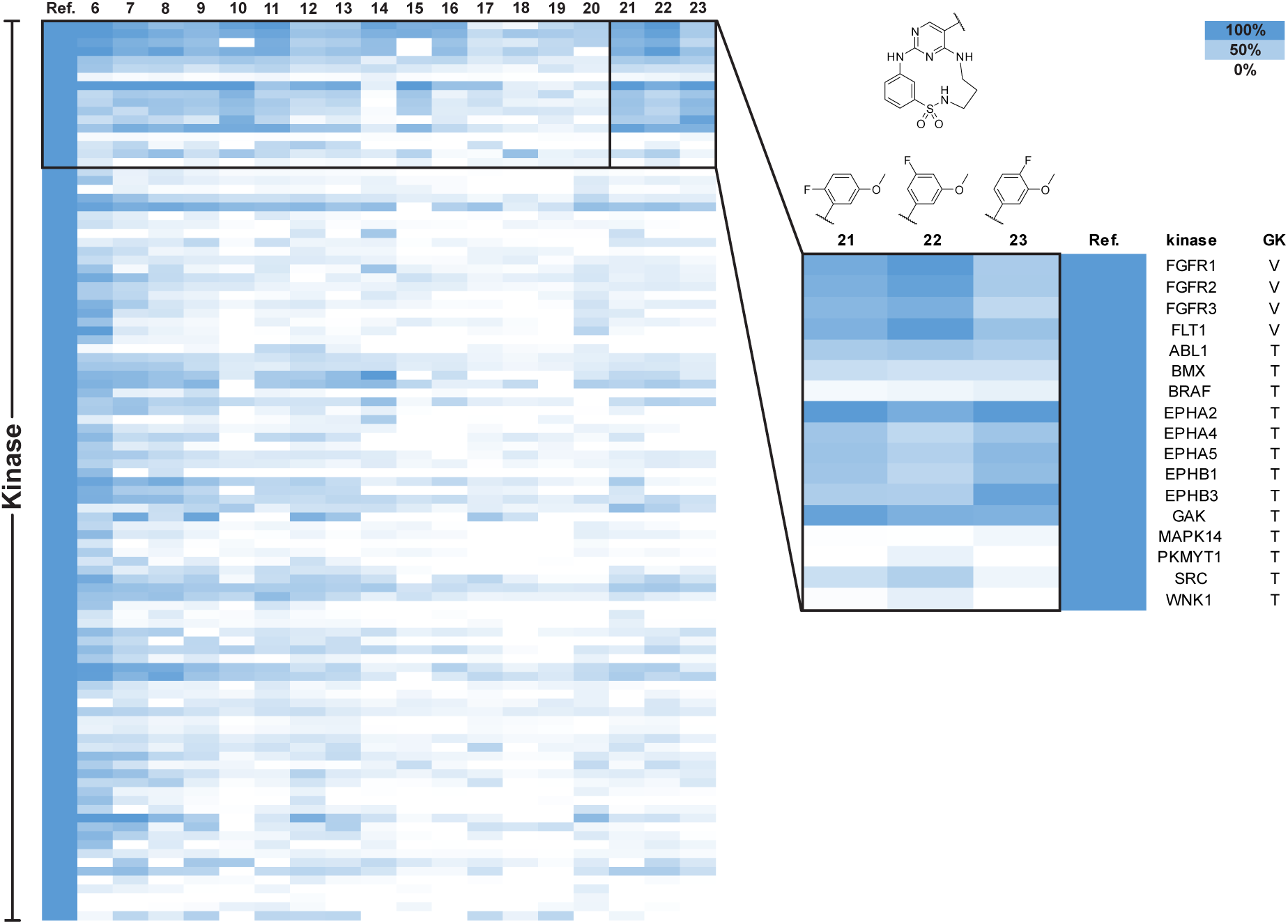
Screening data from initial SAR by differential scanning fluorimetry. Kinases are sorted by gatekeeper residues on the y-axis, against tested compounds (**6-23**) on the x-axis. Heatmap color code ranging from white (0% of reference thermal shift) to blue (100% of reference thermal shift). Disubstituted compounds **21-23** showed selectivity for several kinases with small gatekeeper residues (valine or threonine), which are highlighted in the black box. See Table S1 for the full set of data.

### Crystal structure of EPHA2 in complex with 23 revealed key interactions for further optimization

To gain detailed insights into the binding mode of the 2-aminopyrimidine containing macrocycles substituted at position 5, we determined a crystal structure of the EPHA2-**23** complex (**Figure 3A**). As expected, compound **23** bound to the ATP pocket of EPHA2 with canonical hinge interactions of the 2-aminopyrimidine motif. The sulfonamide N-H interacted with the amide carbonyl oxygen of I619 in the P-loop, while one of the sulfonamide oxygen atoms interacted with the amide N-H of A699 at the N-terminus of the αD-helix. This specific interaction with the front pocket was enhanced by macrocyclization since the rigidity of the linker locked the sulfonamide in its bioactive conformation. In comparison, the acyclic derivative **28**, which was synthesized as shown in **Scheme 2**, has a freely rotatable bond between the sulfur and the ring carbon, potentially accounting for the low Δ*T*_m_ (4.2 K) due to unfavorable entropic contribution to binding. Conversely, the macrocyclic compound **23** stabilized EPHA2 with a Δ*T*_m_ of 8.5 K, possibly due to its restrained bioactive conformation (**Figure 3B**).

**Figure 3.**
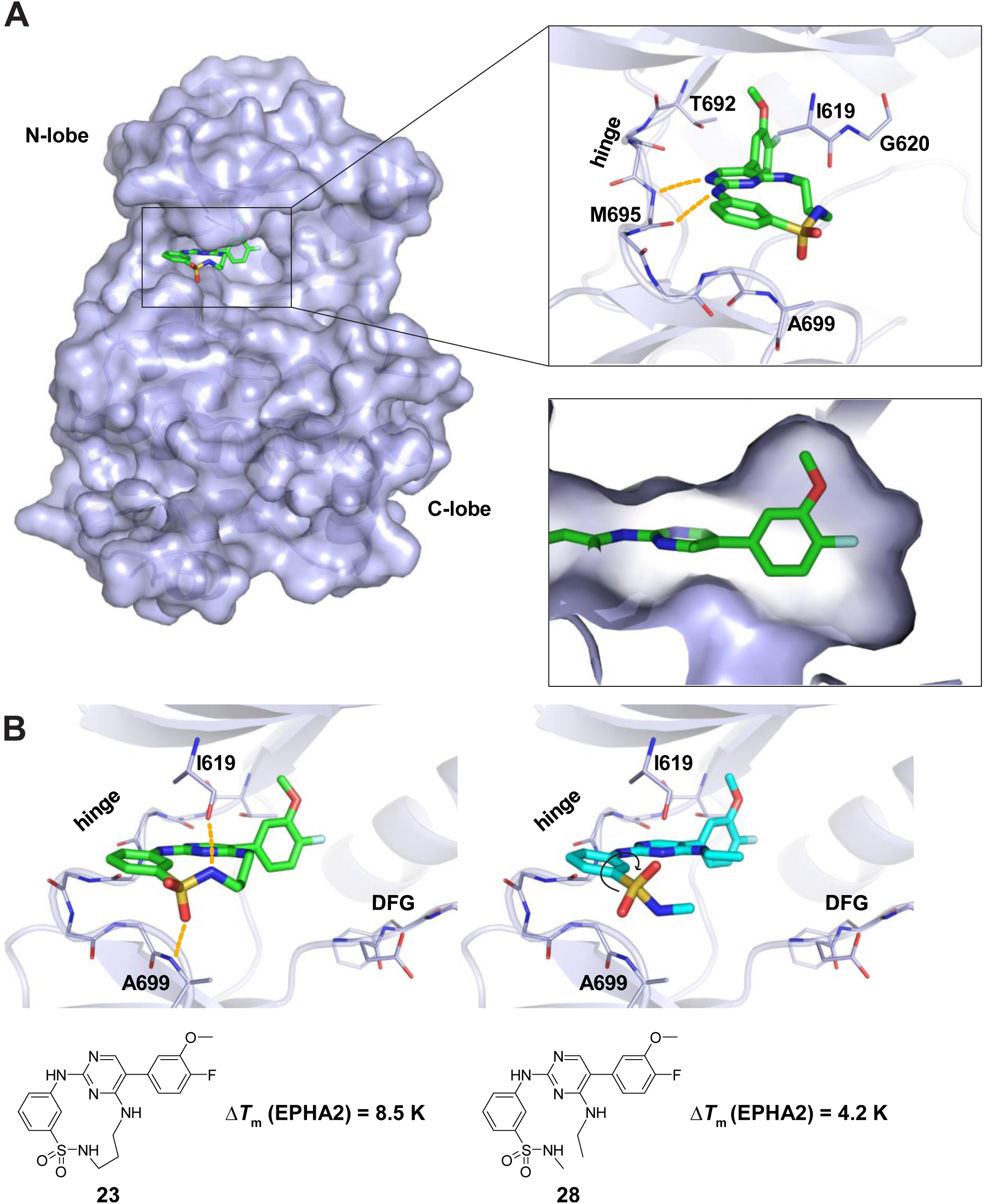
Binding mode of macrocycle **23**. **(A)** Crystal structure of EPHA2 with bound compound **23** in (PDB: 8QQY). Hydrogen bonds are shown as yellow dotted lines. The 5’-(4-fluoro-3-methoxyphenyl) substituent reached into the back pocket of the kinase. **(B)** Comparison of macrocycle **23** and acyclic analogue **28** highlighting the advantage gained by macrocyclization for EPHA2 potency. **28** was energetically minimized using ChemDraw 3D and aligned with **23**. Δ*T*_m_ values were determined by DSF.

**Scheme 2.**
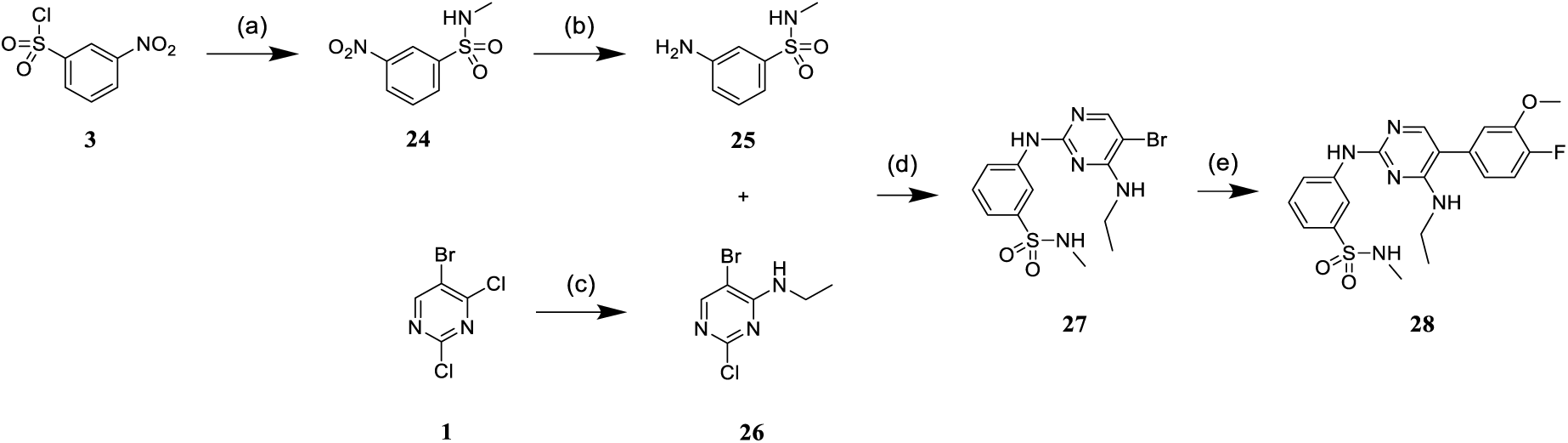
Synthesis of acyclic inhibitor **28**.^a^ ^a^Reagents and conditions: (a) 2 M methylamine in tetrahydrofurane, triethylamine, tetrahydrofurane, 1 h, rt; (b) iron, ammonium chloride, methanol/water (9:1), reflux, 3 h; (c) 2 M ethylamine in tetrahydrofurane, triethylamine, tetrahydrofurane, 1 h, rt; (d) 4 N HCl in 1,4-dioxane, acetonitrile/water, reflux, 24 h; (e) potassium carbonate, [1,1’-Bis(diphenyl-phosphino)ferrocene]-palladium(II) dichloride, 4-fluoro-3-methoxyphenylboronic acid, 1,4-dioxane/dimethylformamide (1:1), 100 °C, µW, 2 h.

The 4-fluoro-3-methoxyphenyl substituent in **23** was oriented perpendicular to the 2-aminopyrimidine motif, and this moiety was located in close proximity to the T692 gatekeeper. The methoxy group in 3’-position filled a cavity created by the β3 and β5 sheet, while the 4-fluorine was located at the end of the back pocket. A closer look at the back-pocket interactions revealed unoccupied space in the pocket targeted by the substituted phenyl, which provided an opportunity for further optimization of **23**. Therefore, the subsequent modifications explored back-pocket interactions in EPHA2 and assessed the impact of various substitutions on inhibitor potency and off-target profiles.

### Back-pocket substitution improved the potency and selectivity for EPHA2 and GAK

We incorporated different di- and trisubstituted aromatic ring systems in the 5’-position of the pyrimidine core and profiled the synthesized compounds with our in-house DSF panel. **Table 1** shows the thermal shifts in K for the target kinases EPHA2 and GAK, closely related off-targets within the EPH family (EPHA4/A5/B1/B3), as well as FGFR1/2 and FLT1. Di-substituted macrocycles (**Table 1**) bearing a 3-methoxy residue on the pendant ring showed considerable differences in Δ*T*_m_ when the 4’-position was modified. Introduction of polar groups such as the carboxymethyl (**41**) or methoxy (**44**) did not improve potency. Having a fluorine (**23**, **48**, **49**, **51**, **53**), chlorine (**42**, **54**), or bromine (**55**) substituent at the 4’-position correlated with high Δ*T*_m_ values, as long as the substituent in position 3 was not too bulky. For example, introduction of isopropoxy (**43**), trifluoromethyl (**47**), or carboxymethyl (**50**) at the 3-position resulted in almost complete loss of binding, highlighting the spatial limitations of the back pocket. The highest Δ*T*_m_ values for EPHA2 were observed for **23**, **48**, and **52**, which had all Δ*T*_m_ values >8 K. For GAK, compounds **42**, **52**, **53,** and **55** showed Δ*T*_m_ values of more than 7 K. Compounds **45**, **46**, **48,** and **49** showed the highest Δ*T*_m_ stabilization for the off-targets FGFR1/2 and FLT1. When comparing the selectivity of these di-substituted derivatives, **52** and **55** showed improved on-target potency and reduced off-target interactions. To further explore the EPHA2 back pocket, we also synthesized trisubstituted derivatives (**Table 2**), with additional substitution of the 4-fluoro-3-methoxy moiety by a fluorine (**56**, **58,** and **59**). The 2,4,5-substituted derivative **56** demonstrated improved potency and selectivity for EPHA2 and GAK. We then compared various substituents, including fluorine, chlorine, methoxy and methyl groups, all following the 2,4,5-substitution pattern. The 2-fluoro-4-chloro-5-methoxyphenyl derivative (**61**) showed the highest stabilization, with Δ*T*_m_ values of 9.8 K and 9.3 K for EPHA2 and GAK, respectively.

**Table 1.**
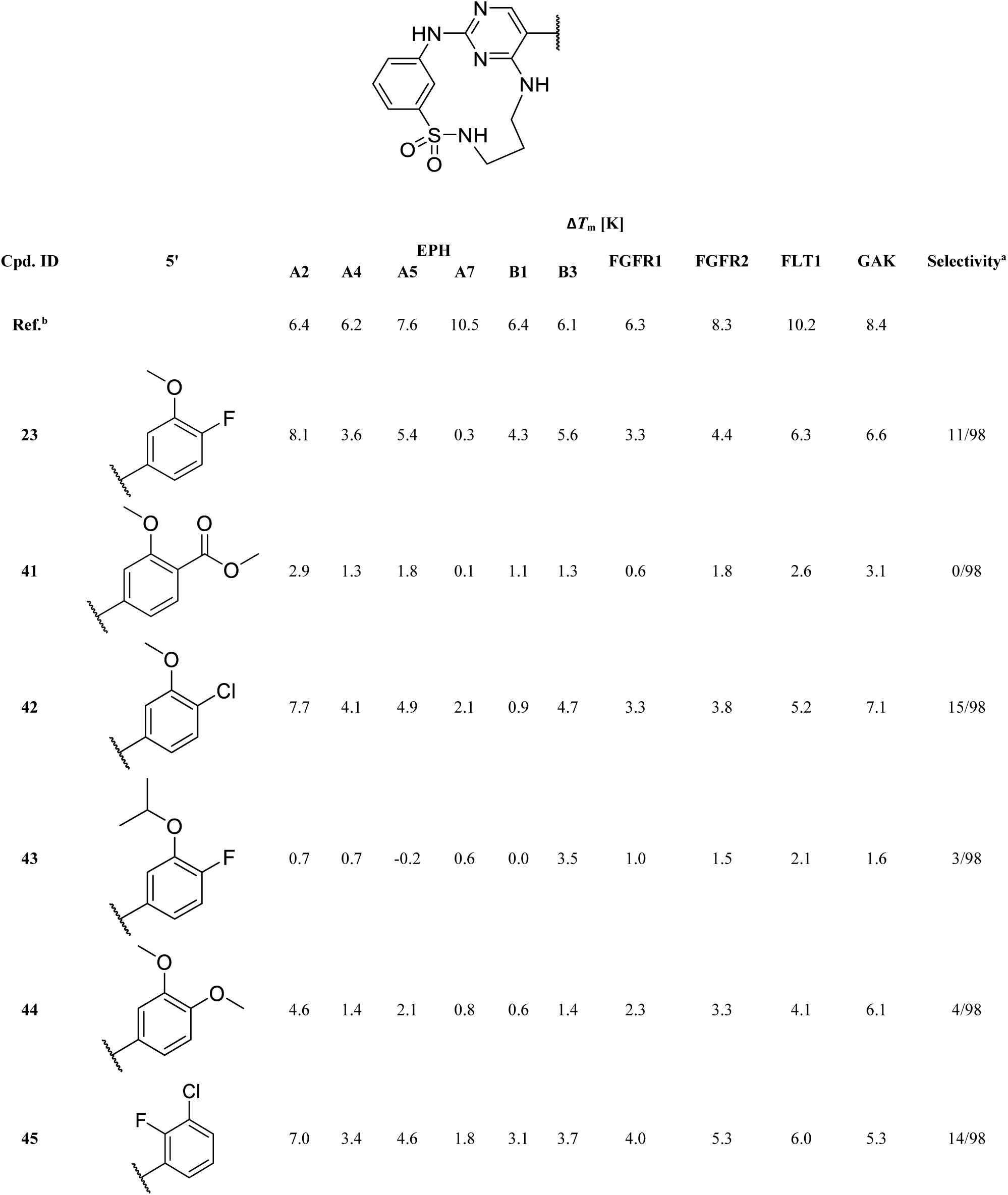
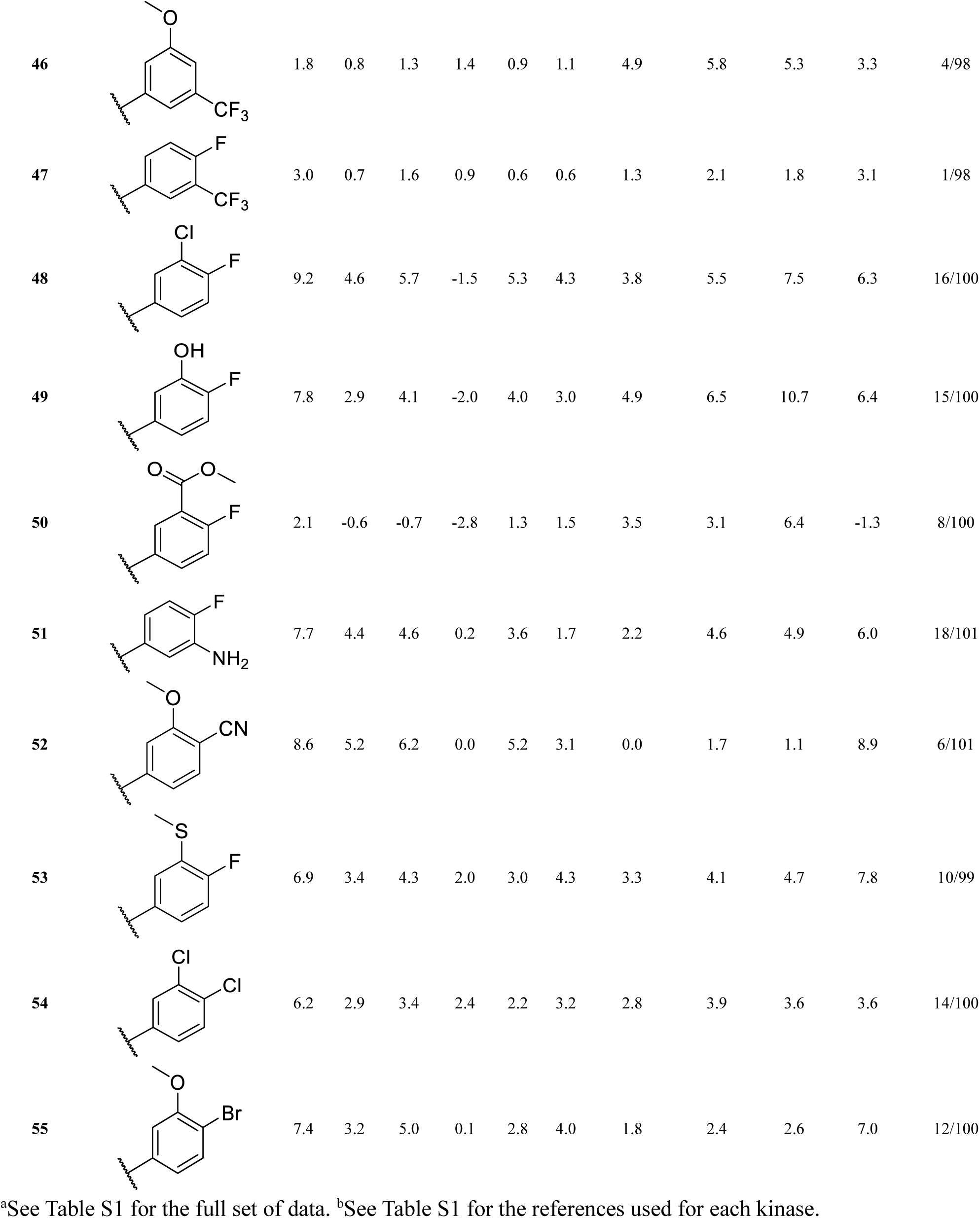
Thermal shift data of 5’-disubstituted compounds for selected kinases. The selectivity was determined by dividing the number of kinases with thermal shifts higher than 50% of that of the reference compound by the total number of kinases investigated.

**Table 2.**
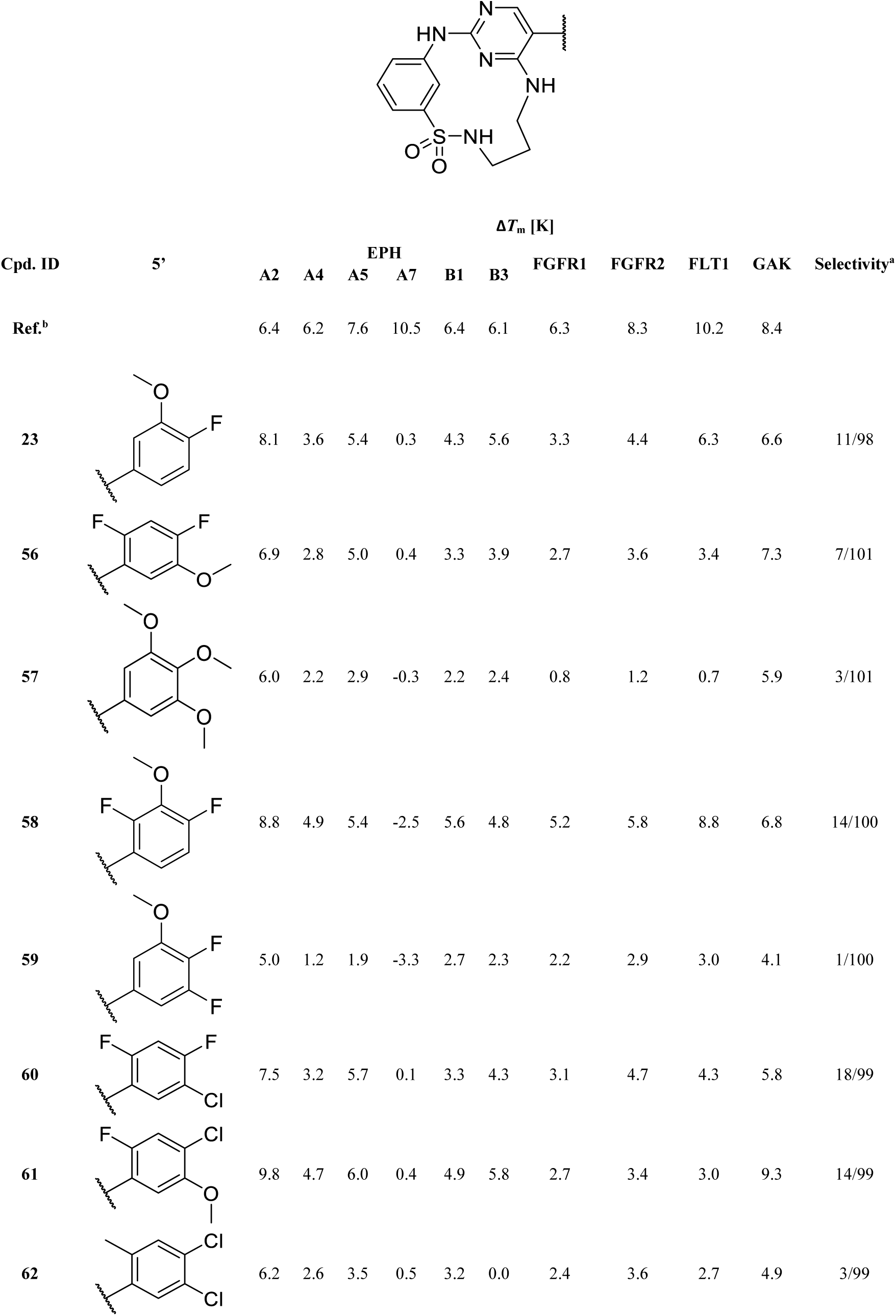
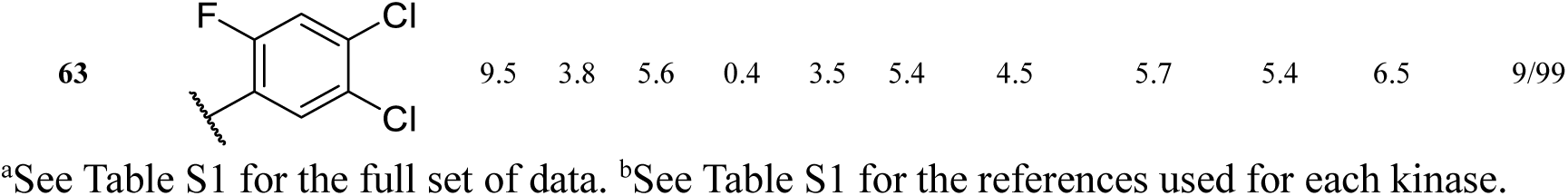
Thermal shift data of 5’-trisubstituted cpds for selected kinases. The selectivity was determined as in. **Table 1**.^a^

### Front-pocket derivatization leads to selectivity for GAK

The crystal structure of EPHA2 in complex with **23** suggested that this inhibitor series can be further optimized by targeting interactions in the solvent-exposed area of the ATP-binding site (front pocket). A sequence alignment of the identified on- and off-targets revealed significant differences in position 3 of the αD-helix, close to the binding site of the sulfonamide phenyl moiety (**Figure 4A**). To potentially exploit differences in the front-pocket region of our target kinases and the identified off-targets, different substituents were inserted in ortho position (synthetic route is shown in **Scheme 3**) for elongation towards the αD-helix. We introduced methyl, chlorine, and methoxy moieties by using the respective commercially available sulfonyl chlorides, and the derivatives **73-75** were synthesized according to the route described in **Scheme 1**. Suzuki coupling of **74** with 4-hydroxyphenylboronic acid yielded **76** (**Scheme 3B**). Demethylation of **72** with sodium ethanethiolate and subsequent Suzuki coupling led to **78**. The resulting hydroxy function of **78** was then utilized for a nucleophilic substitution reaction under basic conditions with either 2-bromo-1-methoxyethane (**79**), methyl 2-bromoacetate (**80**), 2-bromoacetamide (**81**) or 4-(2-bromoethyl)morpholine (**82**), yielding the respective front-pocket derivatives.

**Figure 4.**
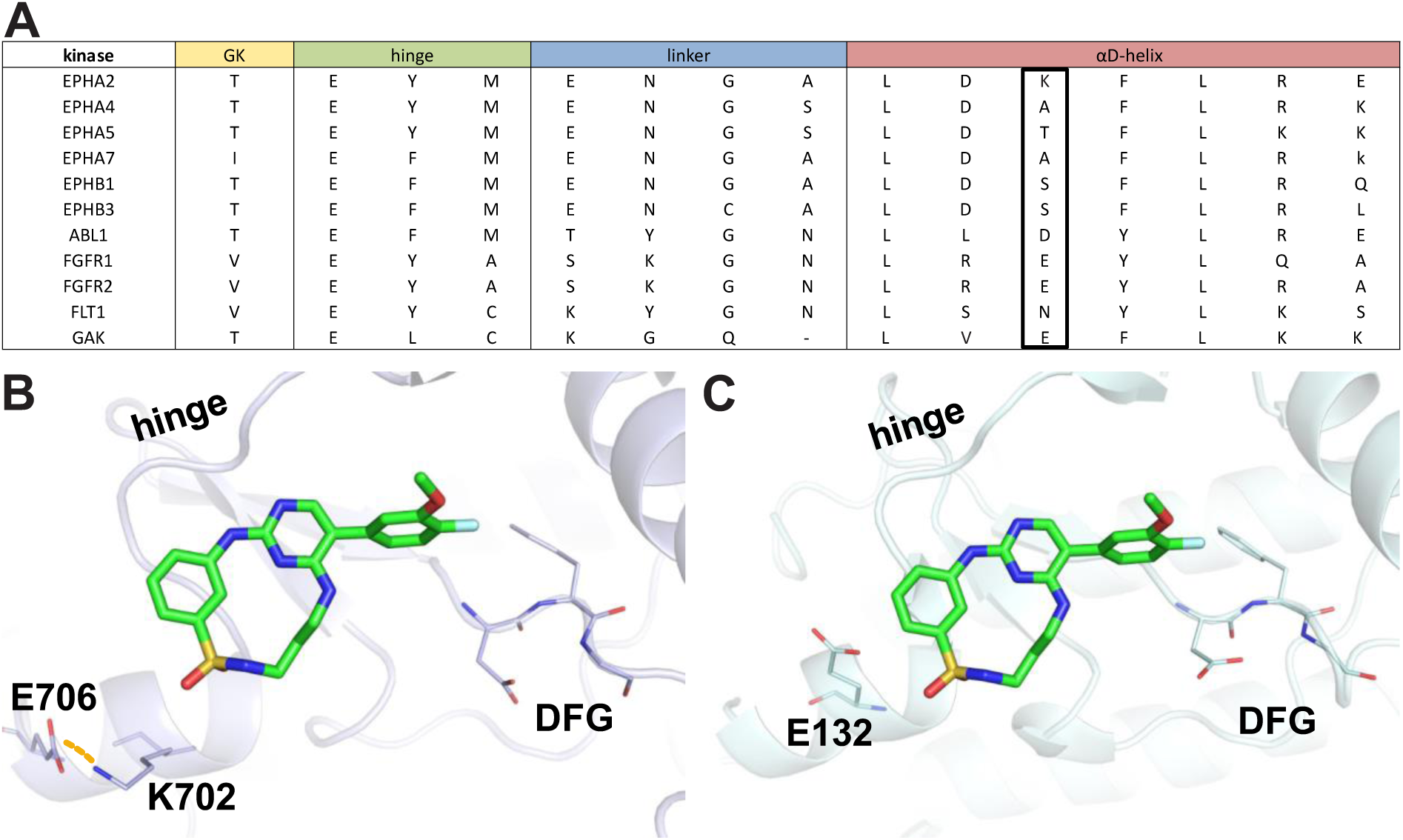
Sequence comparison of the front pocket in EPH family proteins, GAK, and related kinases. **(A)** Sequence alignment of identified on- and off-target kinases, highlighting the differences between gatekeeper (GK), hinge, linker and αD-helix residues. **(B)** Crystal structure of EPHA2 in complex with **23**. K702, an isoform-specific variation in the αD-helix close to the sulfonyl group of **23** is highlighted. The salt bridge with E706 is indicated by an orange dashed line. **(C)** Superimposition of the structures of the EPHA2-**23** complex and GAK (PDB: 5Y80) highlighting the acidic E132 residue at the equivalent position in the αD-helix in GAK. The protein chain of EPHA2 has been omitted for clarity.

**Scheme 3.**
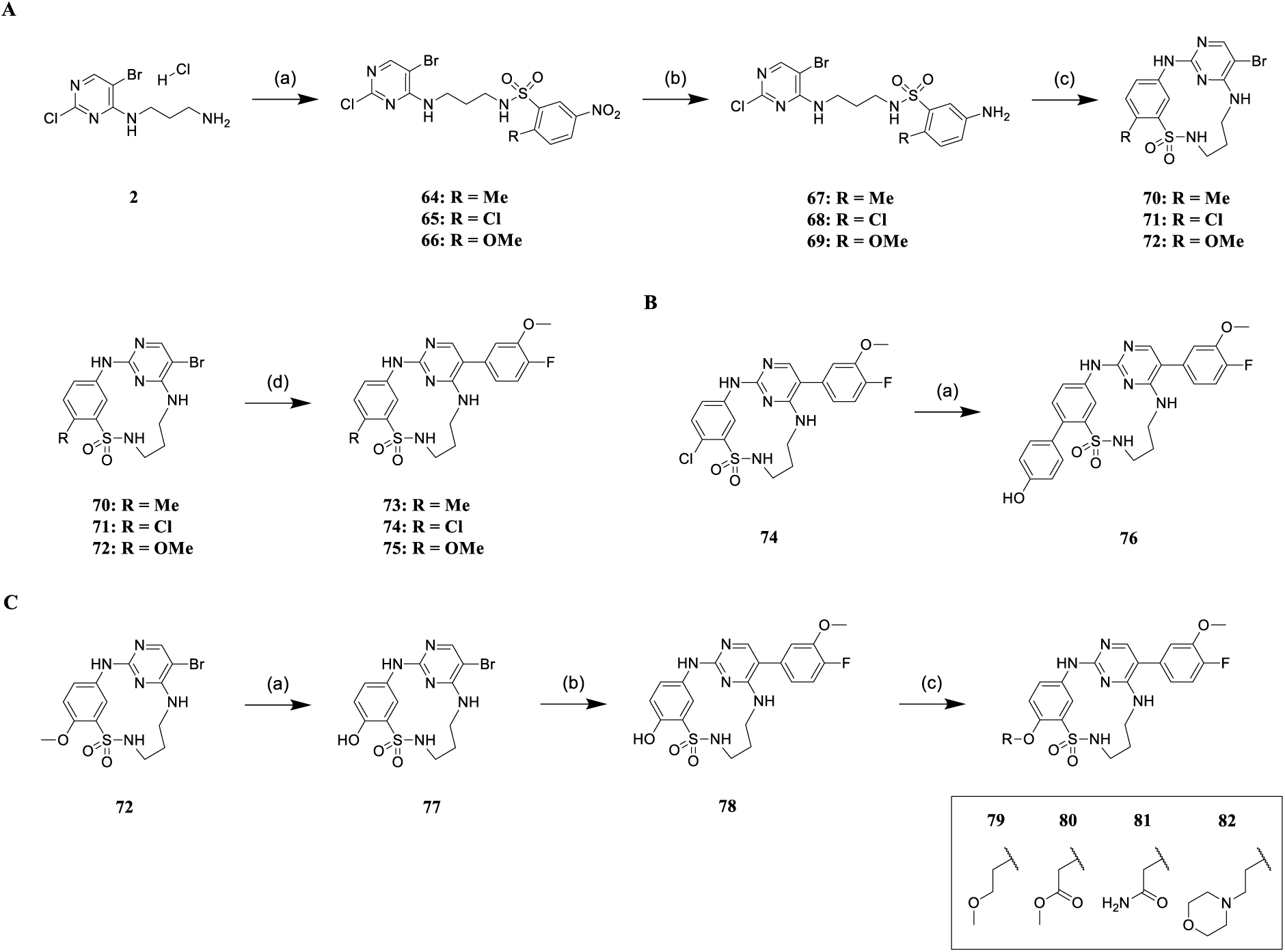
Synthesis of front-pocket derivatives. **(A)** Synthesis of inhibitors **73-75**.^a^ **(B)** Synthesis of inhibitor **76**.^b^ **(C)** Synthesis of inhibitors **79**-**82**.^c^ ^a^Reagents and conditions: (a) 2-R-5-nitrobenzenesulfonyl chloride, triethylamine, acetone / water (3:1), rt, 5 h; (b) iron, ammonium chloride, methanol/water (9:1), reflux, 3 h; (c) 4 N HCl in 1,4-dioxane, acetonitrile, water, 2-butanol, reflux, 6 h; (d) potassium carbonate, [1,1’-Bis(diphenyl-phosphino)ferrocene]palladium(II) dichloride, 4-fluoro-3methoxyphenylboronic acid, 1,4-dioxane / dimethylformamide (1:1), 100 °C, µW, 2 h. ^b^Reagents and conditions: (a) potassium carbonate, XPhos Pd G2, 4-hydroxyphenylboronic acid, 1,4-dioxane/dimethylformamide (1:1), 100 °C, µW, 2 h. ^c^Reagents and conditions: (a) sodium ethanethiolate, dimethylformamide, 100 °C, µW, 10 h; (b) potassium carbonate, [1,1’-Bis(diphenylphosphino)ferrocene]palladium(II) dichloride, 4-fluoro-3-methoxyphenylboronic acid, 1,4-dioxane/dimethylformamide (1:1), 100 °C, µW, 2 h; (c) potassium carbonate, R-Br, dimethylformamide, rt, 2-10 h.

Introduction of a methyl (**73**) or chlorine (**74**) resulted in decreased Δ*T*_m_ values for EPHA2 while increasing selectivity compared with the parent compound **23** (**Table 3**). In contrast, introduction of hydrophilic residues such as 4-hydroxyphenyl (**76**), hydroxy (**78**), and 2-methoxyethoxy (**79**) maintained inhibitor potency for EPHA2 and improved selectivity. Somewhat surprisingly, introduction of methyl 2-hydroxyacetate (**80**) and 2-hydroxyacetamide (**81**) at this position led to drastically reduced Δ*T*_m_ values for all tested EPH family kinases. The 2-morpholinoethan-1-olyl derivative (**82**) showed a slight decrease of Δ*T*_m_ for EPHA2 (6.9 K), coupled with an increased stabilization (Δ*T*_m_ = 9 K) for GAK. This may be due to its basic character as GAK harbors a glutamate residue at the beginning of the αD-helix, which is in proximity to the front-pocket substitution (**Figure 4C**). In contrast, EPHA2 has a lysine residue at this position (engaged in a salt bridge with E706), which is less favorable for interaction with **82**.

**Table 3.**
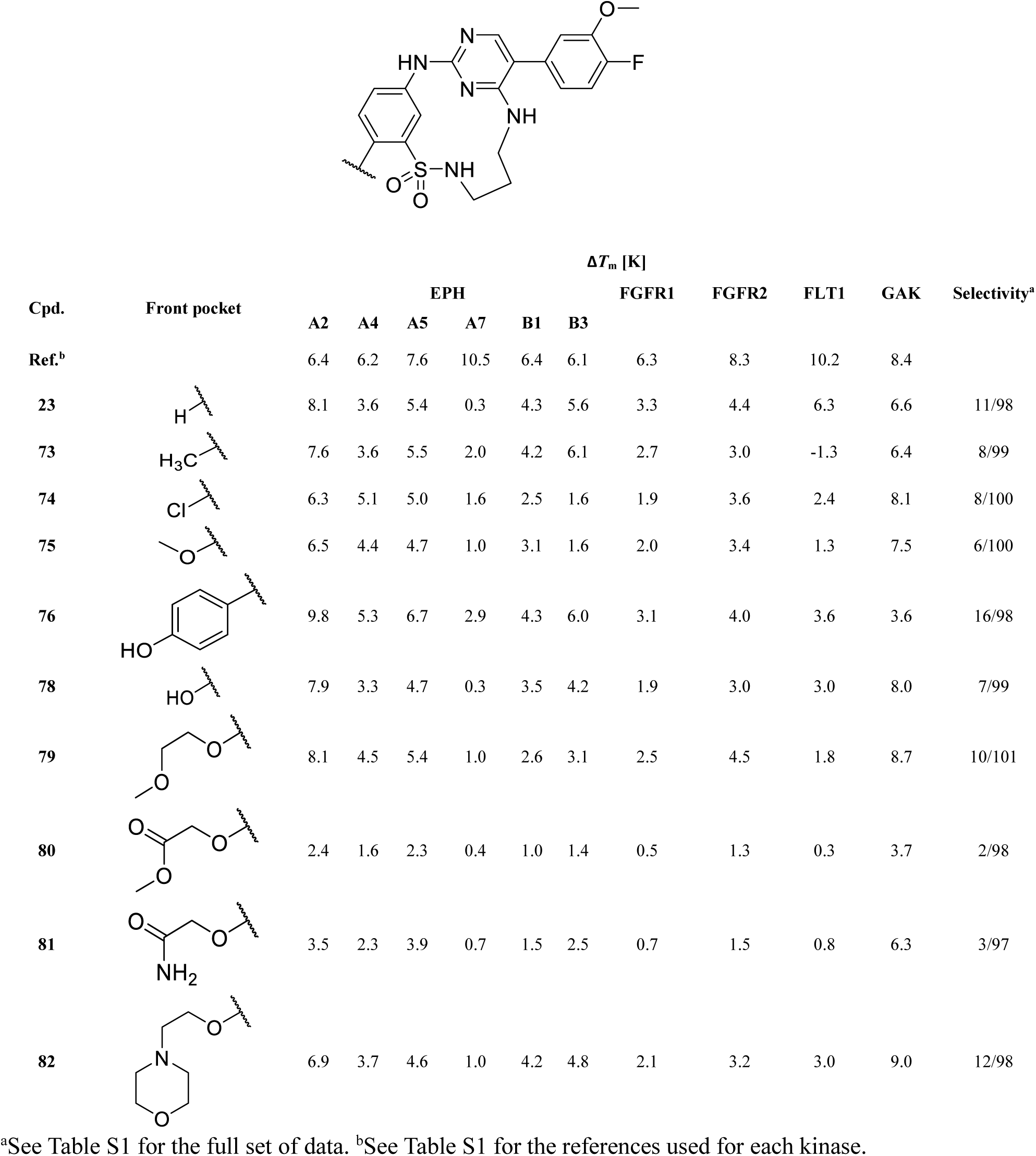
Thermal shift data of front-pocket SAR for selected kinases.^a^.

| Cpd. | Front pocket | $\Delta T_m$ [K] | | | | | | | | | | Selectivity<sup>a</sup> | |
| --- | --- | --- | --- | --- | --- | --- | --- | --- | --- | --- | --- | --- | --- |
|  |  | EPH |  |  |  |  |  | FGFR1 | FGFR2 | FLT1 | GAK |  |  |
|  |  | A2 | A4 | A5 | A7 | B1 | B3 |  |  |  |  |  |  |
| Ref.<sup>b</sup> |  | 6.4 | 6.2 | 7.6 | 10.5 | 6.4 | 6.1 | 6.3 | 8.3 | 10.2 | 8.4 |  |  |
| 23 |  | 8.1 | 3.6 | 5.4 | 0.3 | 4.3 | 5.6 | 3.3 | 4.4 | 6.3 | 6.6 | 11/98 |  |
| 73 |  | 7.6 | 3.6 | 5.5 | 2.0 | 4.2 | 6.1 | 2.7 | 3.0 | -1.3 | 6.4 | 8/99 |  |
| 74 |  | 6.3 | 5.1 | 5.0 | 1.6 | 2.5 | 1.6 | 1.9 | 3.6 | 2.4 | 8.1 | 8/100 |  |
| 75 |  | 6.5 | 4.4 | 4.7 | 1.0 | 3.1 | 1.6 | 2.0 | 3.4 | 1.3 | 7.5 | 6/100 |  |
| 76 |  | 9.8 | 5.3 | 6.7 | 2.9 | 4.3 | 6.0 | 3.1 | 4.0 | 3.6 | 3.6 | 16/98 |  |
| 78 |  | 7.9 | 3.3 | 4.7 | 0.3 | 3.5 | 4.2 | 1.9 | 3.0 | 3.0 | 8.0 | 7/99 |  |
| 79 |  | 8.1 | 4.5 | 5.4 | 1.0 | 2.6 | 3.1 | 2.5 | 4.5 | 1.8 | 8.7 | 10/101 |  |
| 80 |  | 2.4 | 1.6 | 2.3 | 0.4 | 1.0 | 1.4 | 0.5 | 1.3 | 0.3 | 3.7 | 2/98 |  |
| 81 |  | 3.5 | 2.3 | 3.9 | 0.7 | 1.5 | 2.5 | 0.7 | 1.5 | 0.8 | 6.3 | 3/97 |  |
| 82 |  | 6.9 | 3.7 | 4.6 | 1.0 | 4.2 | 4.8 | 2.1 | 3.2 | 3.0 | 9.0 | 12/98 |  |

<sup>a</sup>See Table S1 for the full set of data. <sup>b</sup>See Table S1 for the references used for each kinase.

### *In-vitro* characterization of the developed macrocycles by surface plasmon resonance spectroscopy reveals high binding affinity for EPHA2 and GAK

We used surface plasmon resonance (SPR) to gain a more comprehensive understanding of the binding affinity and kinetics of the developed macrocycles. The determined binding constants, *K*_D_ values, by SPR (affinity fit) were in good agreement with the DSF assay data for both EPHA2 and GAK. Notably, all compounds tested exhibited lower *K*_D_ values for GAK than for EPHA2, indicating that absolute Δ*T*_m_ values cannot be compared between different proteins. Compound **6** showed the weakest affinity for EPHA2 (*K*_D_ = 307.0 nM) and GAK (*K*_D_ = 112.4 nM), respectively. The key compound **23** had a significantly improved *K*_D_ of 120.4 nM for EPHA2 and 7.7 nM for GAK. SPR data for the acyclic derivative **28** on EPHA2 was not fitted within the covered concentration range (five-point dose-response: 4-1000 nM) due to its low affinity for EPHA2. In contrast, **28** bound to GAK with a *K*_D_ value of 19.7 nM, suggesting that macrocyclization preferentially enhanced the potency on EPHA2 but not GAK. Compound **55** gave *K*_D_ values of 180.5 nM for EPHA2 and 19.2 nM for GAK. The introduction of a front-pocket motif in **75** and **79** slightly increased the potency for GAK compared to **23**. Compound **75** gave *K*_D_ values of 95.3 nM for EPHA2 and 6.2 nM for GAK, while **79** showed 126.1 nM for EPHA2 and 5.0 nM for GAK. Data fitted via kinetic and steady-state affinity fit correlated with an R^2^ of 0.99 for both EPHA2 and GAK (see Table S4 and Figure S1). A closer look at the SPR sensorgrams revealed fast on-rates for all compounds for both EPHA2 and GAK. Interestingly, the dissociation from GAK proceeded more slowly than from EPHA2 for compounds **23** and **55**. The residence time, τ, which is calculated by inverting the dissociation rate constant *k*_d_ of a specific compound from a target protein (τ = 1/*k*_d_) allows easy comparison of binding kinetics.^37^ For compound **23**, the residence time on GAK was 7.6-fold longer than on EPHA2 (τ(GAK) = 85 s; τ(EPHA2) = 11 s), and for compound **55** 6.4-fold longer (τ(GAK) = 93 s; τ(EPHA2) = 15 s).

### Macrocyclic diaminopyrimidines are potent inhibitors in cellular assays

The most promising compounds from our SAR series were subjected to an *in cellulo* target engagement assay (NanoBRET).^38^ This assay is based on bioluminescence resonance energy transfer and enables a dose-dependent candidate evaluation by competition with a fluorescently labeled tracer molecule.^39^ Our objective was to validate the results from our previous *in vitro* experiments and thereby assess both potency and selectivity in a cellular context. **Table 4** shows the results of this experiment for selected compounds. Surprisingly, despite the high Δ*T*_m_ shifts observed for this series for ABL1, the data measured on the isolated kinase domain did not translate into a significant *in cellulo* potency on ABL1 full-length protein as the EC_50_ values for this kinase were mostly in the two-digit micromolar range. However, for EPHA2, EPHA4, and GAK, the cellular BRET data qualitatively correlated well with the DSF data. It is noteworthy that for GAK, most compounds had EC_50_ values below 100 nM in cells. The most potent compounds for EPHA2 were **55** and **58**, which had EC_50_ values of 260 nM and 250 nM, respectively, while compound **23** was found to be the most potent cellular inhibitor of GAK, with an EC_50_ of 20 nM. Interestingly, the potency on GAK remained high throughout this SAR series, whereas the values for EPHA2 varied considerably. Furthermore, the BRET assay data revealed that some members of our SAR series were potent EPHA4 inhibitors. A simple front-pocket modification in compound **75**, harboring an additional methoxy group in *ortho* position to the sulfonamide, led to potent EPHA4 inhibition, with an EC_50_ value of 1600 nM for EPHA2, compared to 37 nM for EPHA4. Compound **79** also demonstrated such a selectivity profile, favoring EPHA4 (EC_50_ = 60 nM) over EPHA2 (EC_50_ = 1400 nM). This suggested that structural variations in the front-pocket interacting residues significantly modulate the selectivity within the EPH family, which may be exploited for the future development of EPHA4-selective chemical probes. According to the SPR data, the front-pocket derivatized compounds **75** and **79** bound to EPHA2 with a *K*_D_ of 95.3 nM and 126.1 nM, respectively, but they showed only weak target engagement in the cellular NanoBRET assays. A possible explanation could be the difference between full-length proteins being assayed in cells, opposed to the isolated kinase domain *in vitro*. Compound **55** showed the most favorable selectivity profile against the off-targets ABL1, FGFR1, and FLT1. In contrast, compound **58**, which was as potent on EPHA2 as **55**, potently targeted FGFR1 (EC_50_ = 470 nM) and FLT1 (EC_50_ = 1100 nM) as well.

**Table 4.** EC_50_ values for prioritized compounds determined in a cellular NanoBRET target engagement assay. See Table S2 for complete data set.

| cpd ID | EC<sub>50</sub> [nM] |  |  |  |  |  |
| --- | --- | --- | --- | --- | --- | --- |
|  | EPHA2 | EPHA4 | GAK | ABL1 | FGFR1 | FLT1 |
| 23 | 470 ± 31 | 450 ± 140 | 20 ± 5.7 | 27000 ± 210 | 3100 ± 870 | 13000 ± 5400 |
| 54 | 370 ± 68 | 84 ± 32 | 160 ± 29 | 2700* | 1200 ± 230 | 13000 ± 130 |
| 55 | 260 ± 73 | 57 ± 23 | 27 ± 5.6 | >50000 | 18000 ± 1900 | 33000 ± 3600 |
| 56 | 440 ± 0.99 | 250 ± 57 | 40* | 30000 ± 2300 | 2000* | 7500* |
| 58 | 250* | 190 ± 45 | N/A. | 6400* | 470* | 1100* |
| 61 | 340 ± 81 | 37 ± 12 | 33 ± 4.9 | 8400* | 7600 ± 1100 | 17000 ± 2300 |
| 75 | 1600 ± 380 | 37 ± 12 | 24 ± 3.4 | >50000 | 34000 ± 11000 | 42000 ± 5700 |
| 78 | 470* | N/A | 30* | 17000* | N/A. | N/A |
| 79 | 1400 ± 290 | 60 ± 13 | 31 ± 2.9 | 32000* | 9800 ± 1900 | 23000 ± 2600 |

\*Mean of one experiment performed in technical duplicates

Encouraged by these findings, we evaluated the potency and selectivity of our most promising inhibitors across all EPH kinases, a TK subfamily with high sequence similarity (**Figure 5**) using NanoBRET assays. To the best of our knowledge, there is currently no inhibitor available that selectively targets only one of the EPH kinases and could therefore be considered an EPH isoform-selective chemical probe.^40^ Lead compound **23** exhibited slight selectivity for EPHA1, EPHA2, EPHA4 and EPHB1, whereas macrocycle **55** featured improved potency and selectivity for EPHA2, EPHA4 and EPHB1. Compound **75**, modified at the front-pocket region, was the most selective inhibitor within the EPH kinase family. It bound to EPHA4 with 37 nM and to EPHA5 with an EC_50_ value of 500 nM (see Table S2 for complete data set).

**Figure 5.**
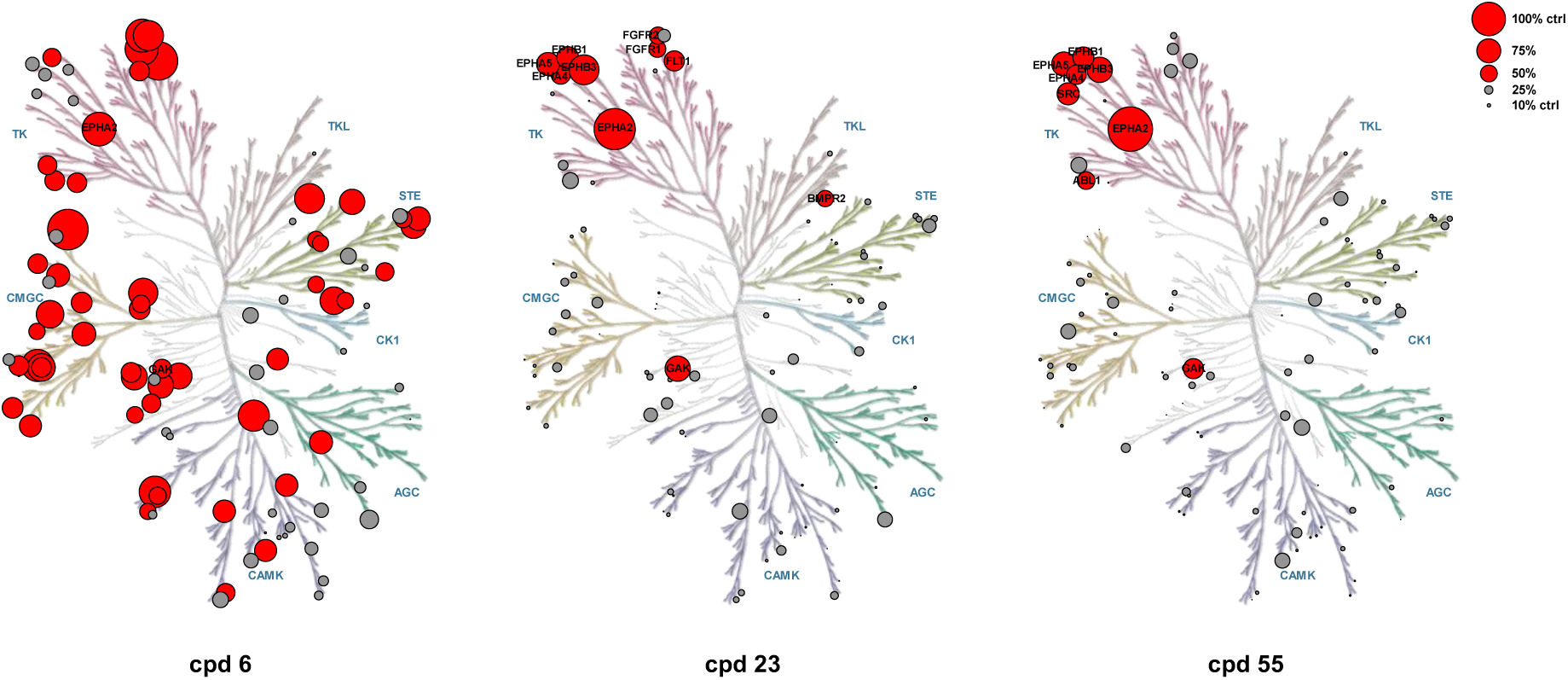
Kinome trees for compounds **6**, **23,** and **55** outlining the improved selectivity. Kinases are represented as dots of different sizes, correspoding to their thermal shift, Δ*T*_m_, relative to a control compound (set as 100%); hits with Δ*T*_m_ values lower than 50% of the Δ*T*_m_ of the control are shown in gray, hits higher than 50% of the control in red. Illustration reproduced courtesy of Cell Signaling Technology, Inc. (www.cellsignal.com).

### Macrocyclic inhibitors demonstrated good microsomal stability

In order to better assess the potential of our macrocycles in a biological context, we next investigated the pharmacokinetic properties of compounds **23** and **55** in a microsomal stability experiment. The compounds were incubated with Sprague-Dawley rat liver microsomes for 1 h at 37 °C using a NADPH regenerating system in a phosphate buffer.^41^ Data was collected at four different time points (0, 30, 45 and 60 min), by quenching the reaction with ice cooled DMF and measuring the samples by HPLC (UV/Vis detection). After 60 min, we found that 90% of **23** and 87% of **55** remained in the normalized area under the curve observed at 0 min. For **23**, we calculated a half-life of 6.6 h, and a half-life of 4.6 h for **55**, by extrapolating the linear regression. The solubility of compound **55** was determined by incubating a saturated solution of the compound in a shaking water bath at 37 °C, filtrating the suspension after 24 h and analyzing the samples by HPLC/MS. We measured a solubility of 2.24 µg/mL for **55**, which was according to the USP definition “slightly soluble”.^42^

Table 5 summarizes the *in vitro* and *in cellulo* experiments for the starting point **6**, initial hit **23**, acyclic derivative **28,** and selected compounds with derivatized back and front pocket motifs, **55**, **75,** and **79**. Compound **55** stands out as the most potent and selective dual EPHA2/GAK inhibitor, with an extrapolated half-life time of 4.6 h in a microsomal stability assay. Interestingly, front-pocket modified compounds **75** and **79** showed good *in vitro* potency for EPHA2 and GAK but were selective for GAK in cells. Within the EPH kinase family, both compounds showed an intriguing *in cellulo* selectivity for EPHA4 (see Table S2 for complete data set). A summary of the SAR is shown in **Figure 8**.

**Table 5.** Profiling of selected compounds.

| | $\Delta T_m$ [K] | | SPR $K_D$ [nM] | | NanoBRET $EC_{50}$ [nM] | | | $t_{1/2}$ [h] | solubility<br>[µg/mL] |
| --- | --- | --- | --- | --- | --- | --- | --- | --- | --- |
|  | EPHA2 | GAK | EPHA2 | GAK | EPHA2 | EPHA4 | GAK |  |  |
| **6** | 6.7 | 4.9 | 307.0 ± 3.8 | 112.4 | 2800 | N/A | N/A | N/A | N/A |
| **23** | 8.1 | 6.6 | 120.4 ± 4.6 | 7.7 | 470 ± 31 | 450 ± 140 | 20 ± 5.7 | 6.6 | N/A |
| **28** | 4.3 | 7.0 | N/A | 49.7 | N/A | N/A | N/A | N/A | N/A |
| **55** | 7.4 | 7.0 | 180.5 ± 3.5 | 19.2 | 260 ± 73 | 57 ± 23 | 27 ± 5.6 | 4.6 | 2.24 ± 0.12 |
| **75** | 6.5 | 7.5 | 95.3 ± 2.0 | 6.2 | 1600 ± 380 | 37 ± 12 | 24 ± 3.4 | N/A | N/A |
| **79** | 8.1 | 8.7 | 126.1 ± 9.9 | 5.0 | 1400 ± 290 | 60 ± 13 | 31 ± 2.9 | N/A | N/A |

**Figure 6.**
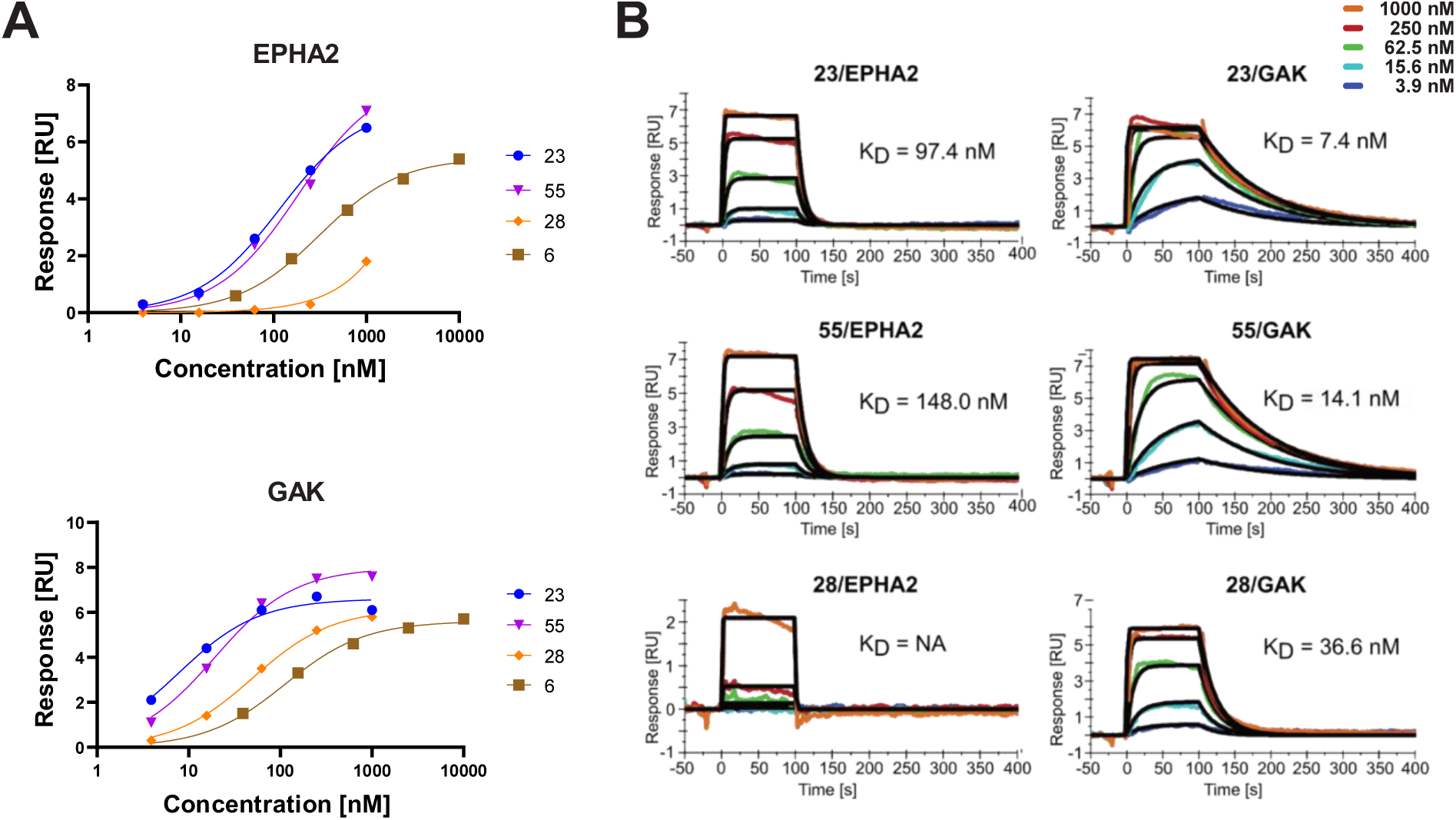
Binding of selected macrocycles to EPHA2 and GAK measured by SPR. **(A)** Affinity fits for compounds **6** (*K*_D_ (EPHA2) = 307.0 ± 5.4; *K*_D_ (GAK) = 112.4), **23** (*K*_D_ (EPHA2) = 120.4 ± 6.5; *K*_D_ (GAK) = 7.7), **28** (*K*_D_ (EPHA2) = NA; *K*_D_ (GAK) = 49.7), and **55** (*K*_D_ (EPHA2) = 180.5 ± 5.0; *K*_D_ (GAK) = 19.2) determined by SPR. (B): Sensorgram for compounds **23**, **28,** and **55** for EPHA2 and GAK with kinetic-fit *K*_D_ values. SPR experiments were conducted in duplicates for EPHA2 and as a single measurement for GAK (five point dose response).

**Figure 7.**
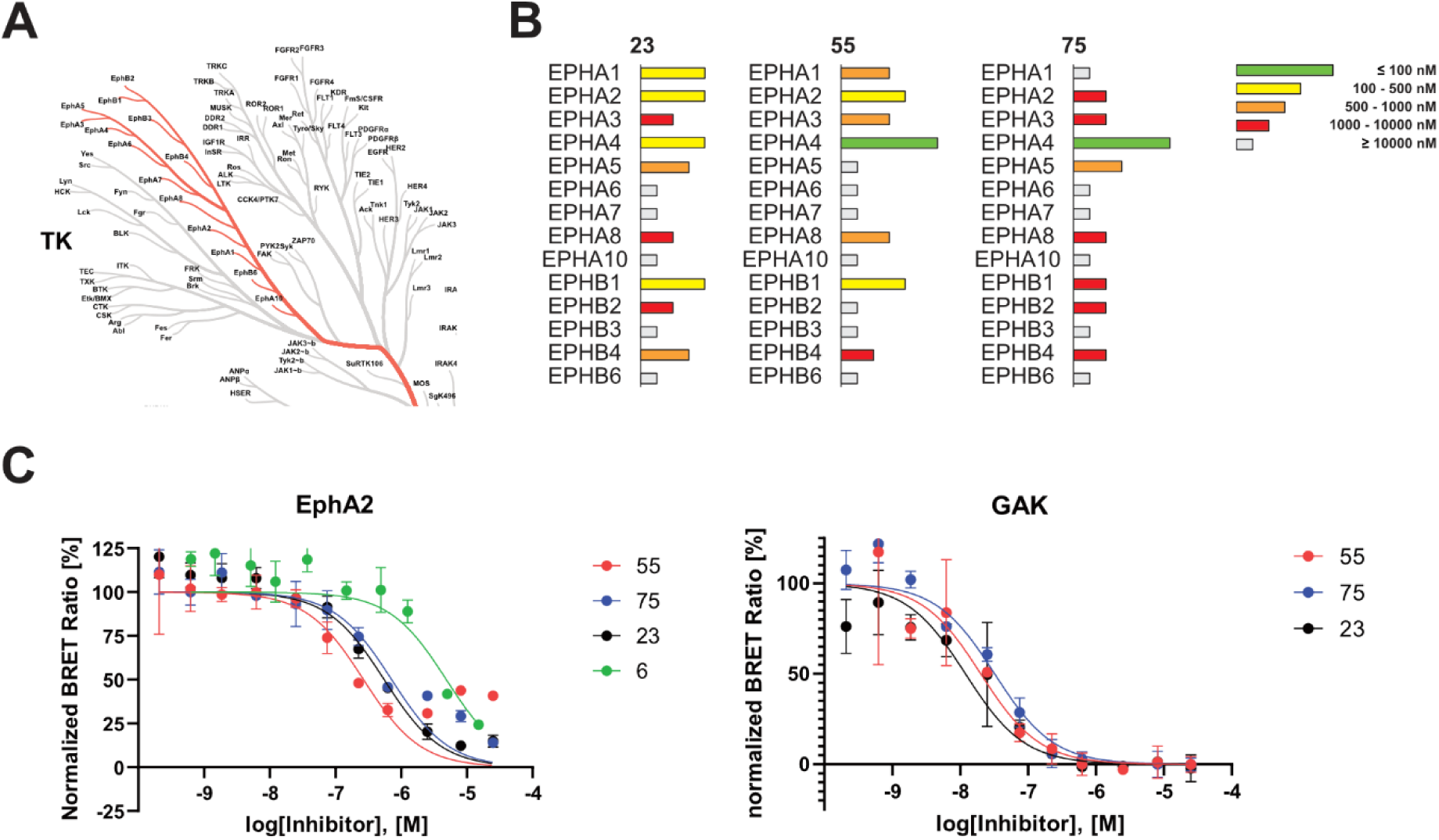
Evaluation of the key compounds from this SAR study in a cellular context. **(A)** Phylogenetic kinome tree showing TK-family members. The EPH subfamily is highlighted by red colored branches (figure was created using the CORAL web software). **(B)** NanoBRET EC_50_ values for compounds **23**, **55**, and **75** measured for all 14 EPHs. Values are depicted as horizontal bars and simplified by different lengths and colors (EC_50_ 100 nM (green), 100-500 nM (yellow), 500-1000 nM (orange), 1000-10000 nM (red), and ≥ 10000 nM (gray)). See Table S2 for comoplete data set. **(C)** Results from the cellular NanoBRET assay, shown as normalized BRET ratio [%] against log(inhibitor) [M] for EPHA2 and GAK. Mean and standard error of the mean (SEM) performed in technical duplicates. The exact number of repeats for each set of experiments is given in Table S2.

**Figure 8.**
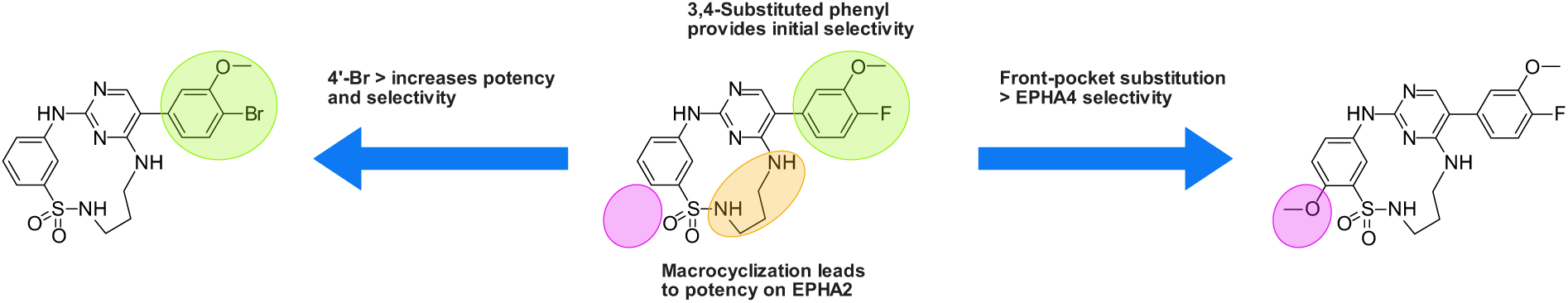
SAR summary of this study. Back-pocket optimization of 2-aminopyrimidine-based macrocycles with 3,4-substituted phenyl moieties eliminated kinases with large gatekeeper residues. Macrocyclization resulted in increased potency on EPHA2. A bromine residue in position 4 of the pendant ring led to increased potency and selectivity for the target kinases EPHA2 and GAK. Front-pocket substitution resulted in improved selectivity for EPHA4 over EPHA2.

### Dual EPHA2/GAK inhibitor 55 showed antiviral activity in DENV infection assay

Taking into account the *in vitro* and *in cellulo* activity data as well as their selectivity and pharmacokinetics, we decided to evaluate the most promising compounds in antiviral activity assays. Huh7 (human hepatoma) cells (Passage 14-15) were pre-treated with selected inhibitors for 1 h and then infected with DENV-2-Rluc virus (a wild-type DENV-2 virus carrying a Renilla luciferase reporter gene) for 45-48 h. Viral infection was measured via luciferase assays, while cell viability was measured via alamarBlue assays. The compounds were screened at three different concentrations (**Figure 9**). We observed antiviral activity for compounds **23**, **54**-**56** and **58**, whereas **75**, **78** and **79** showed no significant effect. This observation was in line with the *in cellulo* activity data showing strong potency against EPHA2. Macrocyclic compounds **23**, **54**-**56** and **58** ranged from 250 to 470 nM activity on EPHA2 and 20 to 160 nM activity on GAK and showed antiviral activity against DENV infection. Surprisingly, the GAK-selective derivatives **75**, **78,** and **79** did not reduce DENV infection at any of the tested concentrations (1, 3 or 10 µM). This is in contrast to the previous work of Wouters *et al.* or Martinez-Gualda *et al.* who were able to demonstrate antiviral activity of potent GAK inhibitors in a DENV infection assay.^22,43^ Other NAK-family members that have been associated with antiviral activity, such as AAK1, are not targeted by our compounds.^27^ Compounds **23** and **56**, failed to reduce DENV infection below 50% even at the highest concentration tested. The most potent EPHA2 inhibitors, **55** and **58,** exhibited the most substantial antiviral activity, both reducing DENV infection by 90% at a concentration of 10 µM. Compound **54** also reduced infection at 10 µM but was slightly less active compared to **55** and **58**. However, it is worth noting that compounds **54** and **58** significantly affected cell viability. This suggests that their antiviral effect may be contingent on a certain level of cellular toxicity. In contrast, candidate **55** had no discernible impact on cell viability at all three concentrations. With an estimated EC_50_ value of approximately 5 µM, compound **55** emerged as the most promising compound within our series, demonstrating both *in vitro* and *in cellulo* selectivity, potency, and efficacy in terms of antiviral activity. The observation that dual EPHA2/GAK inhibitors showed superior antiviral activity than the GAK-selective inhibitors suggests that the antiviral properties of the compounds are essentially due to the inhibition of EPHA2.

**Figure 9.**
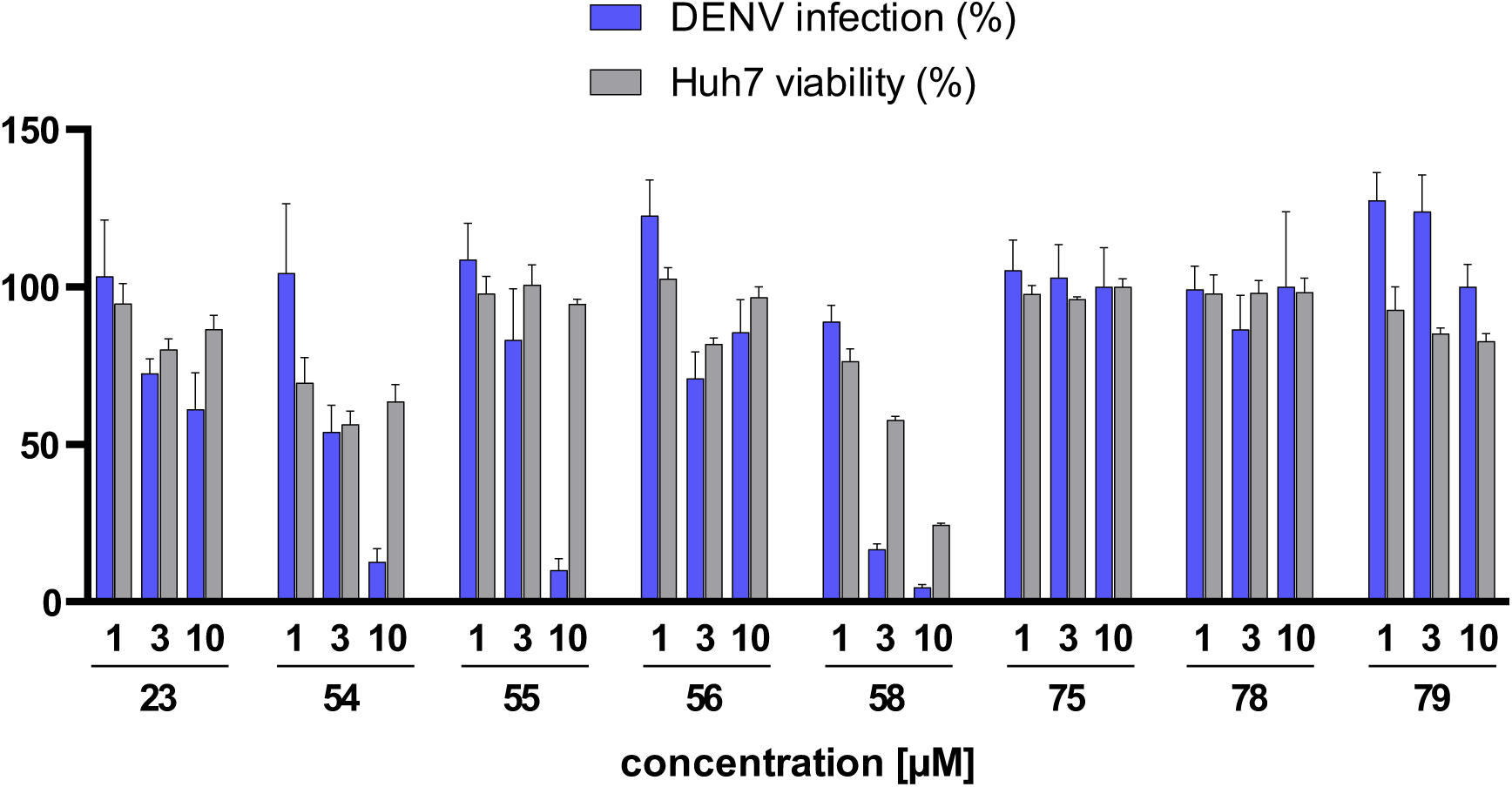
Antiviral activity of compounds **23**, **54**, **55**, **56**, **58**, **75**, **78**, and **79** in a DENV-2 infection assay with Huh7 liver cancer cells. Compounds were tested at three concentrations. Data are represented as bars depicting DENV infection (blue) and cell viability (gray), and mean plus standard deviation (n = 4).

## CONCLUSION

We initiated our GAK/EPHA2 inhibitor optimization study with the potent yet promiscuous published macrocycle **6.** Using a straightforward back-pocket modification at the 5’-position of the pyrimidine core, we were able to significantly improve both potency and selectivity. The 4-fluoro-3-methoxyphenyl substituted derivative **23** was identified as the most potent dual inhibitor from our first SAR series. It displayed excellent stabilization of our intended targets, EPHA2 and GAK, while also showing a promising selectivity profile. Our structure-guided design approach was further supported and validated by determining the crystal structure of EPHA2 in complex with **23**, further substantiating the superiority of the macrocycle structure compared with the acyclic derivative **28**. Subsequent optimization of the back-pocket targeting moiety led to compound **55**, which proved to be the best hit across the entire series. It exhibited favorable potency and selectivity for EPHA2 and GAK, both *in vitro* and *in cellulo*. Furthermore, compound **55** demonstrated good microsomal stability, with an extrapolated half-life of 4.6 h and a water solubility of 2.24 µg/mL. Screening in a DENV-2 infection assay revealed promising antiviral activity without compromising cell viability. In addition, we evaluated the effects of simple front-pocket optimization on the *in cellulo* selectivity profile of our compound series. Macrocycles **75** and **79** exhibited only micromolar EC_50_ values for EPHA2 in the NanoBRET assay but displayed significantly improved potency against EPHA4, whereas their potency against GAK remained unchanged. This observation could serve as a promising starting point for future inhibitor development, especially as EPHA4 represents an intriguing target not only in different types of cancers but also in neurological pathologies such as amyotrophic lateral sclerosis or Alzheimer’s disease.^44–48^ Notably, the *in cellulo* GAK-selective compounds **75** and **79** showed no antiviral activity against DENV infection, which was unexpected, given that GAK is a known antiviral target.^27^ To the best of our knowledge, inhibition of EPHA2 has not been reported as a strategy to combat DENV infection so far. Our inhibition studies, however, suggest that EPHA2 plays a significant role in DENV infection and could therefore be a promising new drug target for DENV therapy.

## MATERIALS AND METHODS

### Chemistry

Detailed information on compound synthesis, including analytical data for the final products, is provided in the Supporting Information. All commercial chemicals, obtained in reagent grade from common suppliers, were used without additional purification. For the purification of compounds through flash chromatography, a puriFlash XS 420 device with a UV-VIS multiwave detector (200–400 nm) from Interchim and pre-packed normal-phase PF-SIHP silica columns (30 μm particle size, Interchim) were employed. The synthesized compounds underwent characterization by NMR and mass spectrometry (ESI). Final inhibitors were identified using high-resolution mass spectrometry (HRMS), and their purity was assessed by HPLC. ^1^H and ^13^C NMR spectra were measured on an AV300, an AV400, or an AV500 HD AVANCE III spectrometer from Bruker. Chemical shifts (δ) are reported in parts per million (ppm), with DMSO-d6 as the solvent, and the spectra referenced to the residual solvent signal: 2.50 ppm (^1^H NMR) or 39.52 ppm (^13^C NMR). HRMS was measured on a ThermoScientific MALDI LTQ Orbitrap XL or a Bruker micOTOF. Compound purity determination by HPLC was carried out on an Agilent 1260 Infinity II device with a 1260 DAD HS detector (G7117C; 254 nm, 280 nm, 310 nm) and an LC/MSD device (G6125B, ESI pos. 100-1000). The compounds were analyzed on a Poroshell 120 EC-C18 (Agilent, 3 x 150 mm, 2.7 µm) reversed phase column with 0.1% formic acid in water (A) and 0.1% formic acid in acetonitrile (B) as eluents. The gradient used was as follows: Method 1 (M1): 0 min: 5% B - 8 min: 100% B – 10 min: 100% B (flow rate of 0.6 mL/min.). Method 2 (M2): 0 min: 5% B - 2 min: 80% B - 5 min: 95% B - 7 min: 95% B (flow rate of 0.5 mL/min.). UV-detection was performed at 320 nm (150 nm bandwidth), and all compounds for further biological characterization exhibited ≥ 95% purity unless stated otherwise.

### Differential scanning fluorimetry (DSF)

DSF was employed to determine the effects of inhibitor binding on the apparent melting temperature of recombinant kinases. This was done in a 96-well plate at a protein concentration of 2 μM with 10 μΜ compound in a buffer containing 25 mM HEPES, pH 7.5, 150 mM NaCl, and 0.5 mM TCEP. SYPRO Orange (5000×, Invitrogen) was added at a dilution of 1:1000 (final concentration of 5x). Protein unfolding profiles were recorded using an Agilent MX3005P real-time qPCR instrument while increasing the temperature from 25 to 95 °C at a heating rate of 3 °C/min. *T*_m_ values were calculated using the Boltzmann equation, and differences in melting temperature upon compound binding (Δ*T*_m_ = *T*_m_ (protein with inhibitor) - *T*_m_ (protein without inhibitor)) were determined. Measurements were performed in triplicates, and all kinase selectivity panel data are provided in **Supporting Table S1.**

### Surface plasmon resonance

SPR experiments were performed on a Biacore T200 at 25°C using 10 mM HEPES pH 7.5, 150 mM NaCl, 1 mM TCEP, 0.05% Tween 20 and 2% DMSO as the running buffer. A Series S Sensor Chip CM5 (Cytiva) was normalized with 70% glycerol, then recombinant proteins EPHA2-His and GAK-His diluted in acetate buffer pH 5.5 were covalently coupled, using a standard amine coupling procedure. During this process, the surfaces were activated for protein immobilization using a combination of 483 mM EDC and 10 mM NHS and subsequently deactivated with 1 M ethanolamine. Final responses of 1100 RU and 1300 RU were reached for EPHA2-His as well as 1800 RU for GAK-His. One flow channel was used as a blank reference surface without protein injection. Each compound was measured at a flow rate of 30 µl/min using multi-cycle conditions, serially injecting five single concentrations from 0.4 nM up to 10 µM over the reference and protein surfaces. The resulting sensorgrams were referenced, the blank subtracted and evaluated using the Biacore T200 evaluation software. The following steady-state fit affinity model was employed for plotting and assessing dose-response curves in GraphPad Prism: Req=(C×Rmax)/(K_D_+C). Here, C is the injected concentration, Rmax is the maximal response, and Req is the response at steady state to determine respective *K*_D_ values.

### NanoBRET assay

The assay was performed as described previously.^38^ In brief: full-length kinases were obtained as plasmids cloned in frame with a terminal NanoLuc-fusion (Promega) as specified in Supplementary Table S5. Plasmids were transfected into HEK293T cells using FuGENE HD (Promega, E2312), and proteins were allowed to express for 20 h. Serially diluted inhibitor and NanoBRET™ Kinase Tracer (Supplementary Table S5) at a concentration determined previously as the Tracer EC_50_ (Supplementary Table S5) were pipetted into white 384-well plates (Greiner 784075) using an Echo 550 acoustic dispenser (Labcyte). Corresponding protein-transfected cells were added and reseeded at a density of 2 x 10^5^ cells/mL after trypsinization and resuspending in Opti-MEM without phenol red (Life Technologies). The system was allowed to equilibrate for 2 h at 37 °C/5% CO2 prior to BRET measurements. To measure BRET, NanoBRET™ NanoGlo Substrate + Extracellular NanoLuc Inhibitor (Promega, N2540) was added as per the manufacturer’s protocol, and filtered luminescence was measured on a PHERAstar FSX plate reader (BMG Labtech) equipped with a luminescence filter pair (450 nm BP filter (donor) and 610 nm LP filter (acceptor)). Competitive displacement data were then graphed using GraphPad Prism 10 software using a normalized 3-parameter curve fit with the following equation: Y=100/(1+10^(X-LogIC50)).

### Protein expression and purification

EPHA2 was expressed and purified as previously described by Rak *et al.*^49^. Briefly, EPHA2 (D596-G900) was subcloned in a pET28a vector (N-terminal His_6_ tag, followed by a TEV cleavage site). The expression plasmids were transformed in *Escherichia coli* Rosetta BL21(D3)-R3-pRARE2 competent cells. Initially, cells were cultured in Terrific Broth (TB) media at 37 °C to an optical density (OD) of 3.2 prior to induction with 0.5 mM IPTG at 18 °C overnight. Cells were harvested and suspended in a buffer containing 50 mM Tris, pH 7.5, 500 mM NaCl, 5 mM imidazole, 5% glycerol, and 1 mM TCEP, and lysed by sonication. The recombinant protein was initially purified by Ni^2+^-affinity chromatography. The histidine tag was removed by TEV protease treatment, and the cleaved protein was separated by reverse Ni^2+^-affinity purification. The protein was further purified by size exclusion chromatography using a HiLoad 16/600 Superdex 200 with a buffer containing 30 mM Tris, pH 7.5, 150 mM NaCl, 0.5 mM TCEP, and 5% glycerol. Quality control was performed by SDS gel electrophoresis and ESI-MS, (EPHA2: expected 34.418.8 Da, observed 34.419.2 Da).

### Crystallization

Co-crystallization trials were performed using the sitting-drop vapor-diffusion method at 293 K with a mosquito crystallization robot (TTP Labtech, Royston UK). EPHA2 (10.5 mg/mL in 30 mM Tris pH 7.5, 150 mM NaCl, 0.5 mM TCEP). The protein was incubated with inhibitor **23** at a final concentration of 1 mM prior to setting up crystallization trials. Crystals grew under the condition 20% PEG 3350, 10% ethylene glycol, 0.2 M sodium iodide. Crystals were cryo protected with mother liquor supplemented with 20% ethylene glycol and flash-frozen in liquid nitrogen. X-ray diffraction data sets were collected at 100K at beamline X06SA of the Swiss Light Source, Villigen, Switzerland. Diffraction data were integrated with the program XDS^50^ and scaled with AIMLESS^51^, which is part of the CCP4 package.^52^ The ephrin receptor kinase structure was solved by molecular replacement using MOLREP^53^ with PDB entry 8BIN^49^ as a starting model. Structure refinement was then performed using iterative cycles of manual model building in COOT^54^ and refinement with REFMAC.^55^ The dictionary file for inhibitor **23** was generated using the Grade Web Server (http://grade.globalphasing.org). X-ray data collection and refinement statistics are shown in **Supplementary Table S3.**

### Microsomal stability assay

A mixture of 432 µL phosphate buffer (0.1 M, pH 7.4) and 50 µL NADPH regenerating system (30 mM glucose-6-phosphate, 4 U/mL glucose-6-phosphate dehydrogenase, 10 mM NADP, 30 mM MgCl_2_) was used for preincubation of solubilized test compound (5 µL, final concentration 10 µM) at 37 °C. The reaction was started after 5 min preincubation by adding 13 µL of a microsome mix from the liver of Sprague-Dawley rats (Sigma Aldrich, 20 mg protein/mL in 0.1 M phosphate buffer) in a shaking water bath at 37 °C. 500 µL ice cold methanol were used for stopping the reaction after 0, 30, 45 and 60 min, and the samples were centrifuged at 5000 g for 5 min at 4 °C, followed by analyzing the supernatant by HPLC for quantification of test compound. HPLC was carried out on an Agilent 1260 Infinity II device with a 1260 DAD HS detector (G7117C; 254 nm, 280 nm, 310 nm) and an LC/MSD device (G6125B, ESI pos. 100-1000). The test compounds were analyzed on a Poroshell 120 EC-C18 (Agilent, 3 x 150 mm, 2.7 µm) reversed phase column with 0.1% formic acid in water (A) and 0.1% formic acid in acetonitrile (B) as mobile phase. The gradient used was as follows: 0 min: 5% B - 2 min: 80% B - 5 min: 95% B - 7 min: 95% B (flow rate of 0.6 mL/min.). UV detection was performed at 320 nm (150 nm bandwidth). Different control samples were used as quality control. The first control lacked NADPH, essential for microsomal enzymatic activity. The second control involved inactivated microsomes (incubated for 20 min at 90 °C), and the third control lacked the test compound (serving as a baseline reference). Quantification of the test compound amounts was performed using an external calibration curve. The data is presented as the mean ± SEM of the remaining compound from three independent experiments.

### Solubility

1 mg of test compound was suspended in 2 mL of water and sonicated for 1 min to give a suspension, which was then incubated in a shaking water bath at 37 °C for 24 h. After this time the sample was filtered through a Whatman Uniprep filter with a pore size of 0.2 µM, and the filtrate was analyzed on a Poroshell 120 EC-C18 (Agilent, 3 x 150 mm, 2.7 µm) reversed phase column with 0.1% formic acid in water (A) and 0.1% formic acid in acetonitrile (B) as mobile phase. The gradient used was as follows: 0 min: 5% B - 2 min: 80% B - 5 min: 95% B - 7 min: 95% B (flow rate of 0.6 mL/min.). UV-detection was performed at 320 nm (150 nm bandwidth). A six-point compound titration, ranging from 10 µM to 500 µM, was used for linear regression and quantification of test compound solubility. The data is presented as the mean ± SEM of the remaining compound from three independent experiments.

### Cell lines for DENV and viability assays

Human hepatoma (Huh7, Apath, L.L.C, New York, NY, USA) and BHK-21 (ATCC) cells were grown in Dulbecco’s Modified Eagle’s medium DMEM (Gibco) supplemented with 10% FCS (Biowest), 1% nonessential amino acids (Corning), 1% L-glutamine (Gibco), and 1% penicillin-streptomycin (Gibco). All the cells were maintained in a humidified incubator with 5% CO_2_ at 37 °C and tested negative for mycoplasma by MycoAlert (Lonza, Morristown, NJ). Cells from passage 14-15 (P14-15) were used for this study.

### Virus construct

DENV-2 (New Guinea C strain) TSV01 Renilla reporter plasmid used for the production of (DENV2-Rluc) was a gift from Pei-Yong Shi (The University of Texas Medical Branch).^56^

### Virus production

DENV2-Rluc RNA was transcribed *in vitro* using mMessage/mMachine (Ambion) kits. DENV2-Rluc was produced by electroporating RNA into BHK-21 cells, supernatants were harvested on day 10 post-electroporation, clarified, and stored at −80°C. Viral titers were determined via standard plaque assays on BHK-21 cells.

### Infection assays and drug screening

Huh7 cells were pretreated with the compounds or DMSO for 1 h prior to infection with DENV2-Rluc at a multiplicity of infection (MOI) of 0.01 (n = 4). The inhibitors were maintained for the duration of the experiment. Overall viral infection was measured at 45-48 h post-infection using a Renilla luciferase substrate via luciferase assays. The relative light units (RLUs) were normalized to DMSO-treated cells (set as 100%).

### Cell viability assays

Cell viability was assessed using AlamarBlue® reagent (Invitrogen) according to the manufacturer’s protocol. Fluorescence was detected at 560 nm on GloMax Discover Microplate Reader (Promega). The raw fluorescence values were normalized to DMSO-treated cells (set as 100%).

## Supporting Information

Additional experimental details, materials and methods, DSF kinase selectivity panel, NanoBRET data in intact cells, X-ray data collection and refinement statistics, surface plasmon resonance data, and HPLC and NMR spectra for all compounds.

## Acknowledgments

The authors acknowledge support by the Structural Genomics Consortium (SGC), a registered charity (no: 1097737) that receives funds from Bayer AG, Boehringer Ingelheim, Bristol Meyer Squibb, Genentech, Genome Canada through Ontario Genomics Institute [OGI-196], EU/EFPIA/OICR/McGill/KTH/Diamond Innovative Medicines Initiative 2 Joint Undertaking [EUbOPEN grant 875510], Janssen, Pfizer and Takeda. S.K. is funded by the German Cancer Research Center (DKTK) and the Frankfurt Cancer Institute (FCI). S.K. and B.T.B. also received support from the Collaborative Research Center CRC1399 “Mechanism of drug sensitivity and resistance in small cell lung cancer”. A.C.J. was supported by the German Research Foundation (DFG) grant JO 1473/1-3. This work was supported by awards W81XWH2210283 and W81XWH-16-1-0691 from the Department of Defense/Congressionally Directed Medical Research Programs, award HDTRA11810039 from the Defense Threat Reduction Agency/Fundamental Research to Counter Weapons of Mass Destruction and grant 1R01AI158569-01 from the National Institutes of Health (to SE). S.E. is a Chan Zuckerberg Biohub investigator. M.K. was supported by a PhRMA Foundation Postdoctoral Fellowship in Translational Medicine. We thank the staff at the Swiss Light Source for assistance during x-ray data collection.

## Data availability

The coordinates and structure factors of the EPHA2 complex with macrocyclic inhibitor **23** have been deposited in the Protein Data Bank (PDB) under accession code 8QQY.

## Author contributions

J.G., T.H., and S.K. designed the project; J.G. and C.S. synthesized the compounds; R.Z. expressed, purified and co-crystallized EPHA2; R.Z., A.K., and A.C.J. refined and analyzed the structures of EPHA2 inhibitor complexes; A.K., L.E., and J.G. performed DSF assays; L.M.B., B.T.B. and T.A.L.E. performed NanoBRET assays; C.L. and K.S. performed SPR experiments, J.G. performed microsomal stability and solubility assays; D.T., M.K., and S.E. performed antiviral and cell viability assays, S.K. supervised the research. The manuscript was written by J.G., T.H., and S.K. with contributions from all authors.

