## Supplementary Material for "Back-pocket optimization of 2-aminopyrimidine-based macrocycles leads to potent dual EPHA2/GAK kinase inhibitors with antiviral activity"

#### Synthesis

*N1-(5-bromo-2-chloropyrimidin-4-yl)propane-1,3-diamine hydrochloride* (**2**)

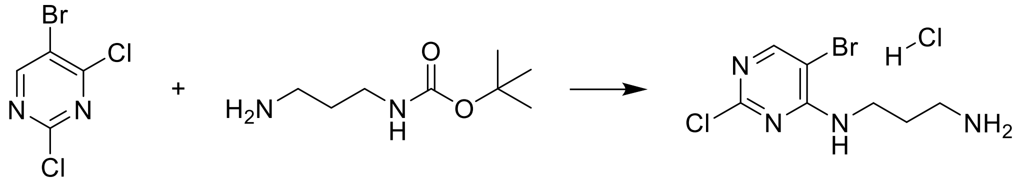

5-bromo-2,4-dichloropyrimidine (**1**) was dissolved in acetonitrile (0.33 M). Triethylamine (1.25 eq.) and tert-butyl (3-aminopropyl)carbamate (1.05 eq.) were added. The reaction mixture was stirred at room temperature for 4 h. The crude reaction mixture was diluted with ethyl acetate and washed with brine, 10% aq. citric acid and sat. aq. $NaHCO_3$, dried over $MgSO_4$, filtered, and concentrated under reduced pressure. The crude product was dissolved in acetonitrile (0.1 M), 4 N HCl in dioxane (4 eq.) was added and stirred like this for 3.5 h. The resulting precipitate was filtered off, washed with acetonitrile, and dried to afford the product as a white crystalline solid (6.6 g, 100%).

**$^1$H-NMR** (400 MHz, DMSO-$d_6$) δ 8.25 (s, 1H), 8.11 (s, 3H), 7.92 (t, J = 5.9 Hz, 1H), 3.44 (q, J = 6.3 Hz, 2H), 2.78 (h, J = 6.9 Hz, 2H), 1.86 (p, J = 6.6 Hz, 2H).

**$^{13}$C NMR** (101 MHz, DMSO) δ 159.48, 158.14, 156.67, 102.79, 37.85, 36.43, 26.13.

**MS (ESI+)** *m/z* $[M+H]^+$: calc. 266.98, found 266.90 (free base)

**HPLC**: $t_R$ = 3.860 min (M1)

*N-(3-((5-bromo-2-chloropyrimidin-4-yl)amino)propyl)-3-nitrobenzenesulfonamide* (**4**)

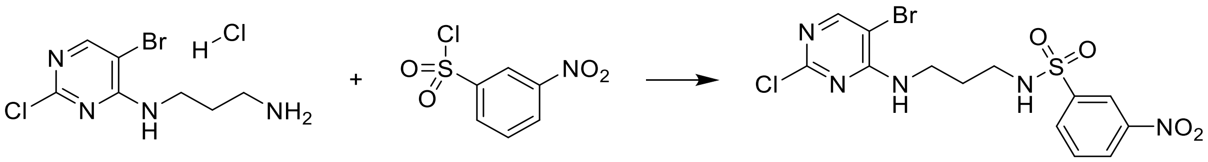

**2** (1.1 eq.) was dissolved in acetone/water 3:1 (0.1 M) and triethylamine (5 eq.) and 3-nitrobenzenesulfonyl chloride (**3**) (1 eq.) were added on ice. The reaction mixture was stirred for 3 h at room temperature after which time the acetone was removed *in vacuo*. The mixture was then diluted with water and extracted 2x with ethyl acetate. The combined organic phases were washed with 10% aq. citric acid, sat. aq. $NaHCO_3$ and brine, then dried over $MgSO_4$, filtered over cotton, and concentrated under reduced pressure to afford the product as a yellow crystalline solid (3.9 g, 96%).

**$^1$H-NMR** (400 MHz, DMSO-$d_6$) δ 8.50 (ddd, J = 2.3, 1.7, 0.5 Hz, 1H), 8.45 (ddd, J = 8.2, 2.3, 1.0 Hz, 1H), 8.21 – 8.17 (m, 2H), 7.99 (t, J = 5.8 Hz, 1H), 7.88 (ddd, J = 8.3, 7.8, 0.5 Hz, 1H), 7.63 (t, J = 5.7 Hz, 1H), 3.33 – 3.28 (m, 2H), 2.87 – 2.80 (m, 2H), 1.66 (p, J = 7.0 Hz, 2H).

**$^{13}$C NMR** (101 MHz, DMSO) δ 159.33, 158.12, 156.61, 147.88, 142.00, 132.53, 131.33, 127.04, 121.26, 102.59, 40.39, 38.35, 28.16.

**MS (ESI+)** *m/z* $[M+H]^+$: calc. 451.95, found 451.90

**HPLC**: $t_R$ = 7.389 min (M1)

*3-Amino-N-(3-((5-bromo-2-chloropyrimidin-4-yl)amino)propyl)benzenesulfonamide* (**5**)

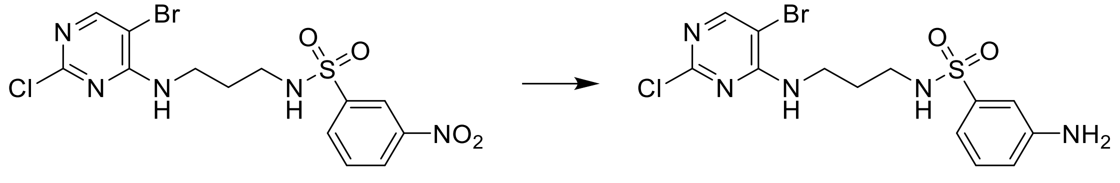

**4** was dissolved in MeOH/water 9:1 (0.07 M), iron (7 eq.) and ammonium chloride (7 eq.) were added, and the resulting mixture was heated under reflux for 4 h. The reaction crude was filtered over Celite and concentrated under reduced pressure. The residue was diluted with ethyl acetate and washed with 10% aq. citric acid, sat. aq. $NaHCO_3$ and brine, then dried over $MgSO_4$, filtered over cotton, and purified by flash column chromatography on silica gel using dichloromethane / methanol as an eluent (95:5) to afford the product as a white crystalline solid (3.1 g, 85%).

**$^1$H-NMR** (400 MHz, DMSO-$d_6$) δ 8.22 (s, 1H), 7.64 (t, J = 5.8 Hz, 1H), 7.39 (t, J = 6.0 Hz, 1H), 7.15 (t, J = 7.8 Hz, 1H), 6.97 (t, J = 2.0 Hz, 1H), 6.85 (ddd, J = 7.7, 1.8, 0.9 Hz, 1H), 6.73 (ddd, J = 8.1, 2.3, 0.9 Hz, 1H), 5.54 (s, 2H), 3.38 – 3.29 (m, 2H), 2.75 (q, J = 6.7 Hz, 2H), 1.66 (p, J = 7.0 Hz, 2H).

**$^{13}$C NMR** (101 MHz, DMSO) δ 159.35, 158.18, 156.59, 149.35, 140.78, 129.46, 117.10, 113.07, 111.06, 102.63, 38.45, 28.34.

**$^{13}$C-dept NMR** (101 MHz, DMSO) δ 156.33, 129.20, 116.84, 112.81, 110.80, 40.10, 38.19, 28.08.

**MS (ESI+)** *m/z* $[M+H]^+$: calc. 421.98, found 421.90

**HPLC**: $t_R$ = 6.421 min (M1)

*$1^5$-Bromo-4-thia-2,5,9-triaza-1(2,4)-pyrimidina-3(1,3)-benzenacyclononaphane 4,4-dioxide* (**6**)

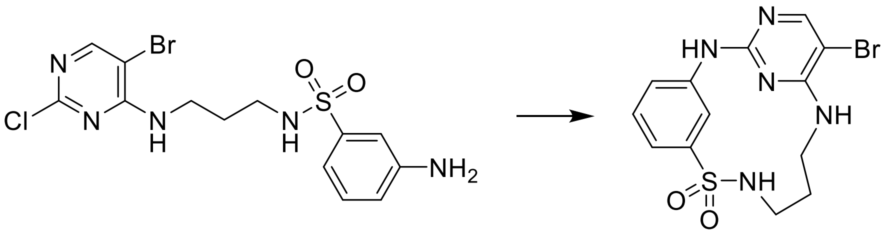

**5** was dissolved in acetonitrile/water/2-butanol 9:1:0.3 (0.05 M) and added dropwise to a refluxing solution of acetonitrile/water 9:1 and HCl in dioxane (4 eq.) over 3 h, after which time it was stirred for another 3 h. The reaction mixture was allowed to cool down, the precipitate was filtered off and washed with water to afford the product as a white solid (850 mg, 85%).

**$^{1}$H-NMR** (400 MHz, DMSO-d$_6$) δ 10.86 (s, 1H), 9.03 (t, J = 1.4 Hz, 1H), 8.64 (t, J = 5.8 Hz, 1H), 8.27 (s, 1H), 7.83 (t, J = 5.8 Hz, 1H), 7.52 – 7.46 (m, 2H), 7.36 (dt, J = 6.8, 2.3 Hz, 1H), 3.45 (dt, J = 11.3, 5.3 Hz, 2H), 3.30 (q, J = 5.6 Hz, 2H), 1.82 (dt, J = 14.5, 5.3 Hz, 2H).

**$^{13}$C NMR** (101 MHz, DMSO) δ 158.40, 152.36, 146.31, 143.78, 138.74, 129.38, 122.67, 120.13, 118.67, 92.56, 28.05.

**$^{13}$C-dept NMR** (101 MHz, DMSO) δ 146.05, 129.12, 122.42, 119.87, 118.42, 40.02, 39.38, 27.80.

**MS (ESI+)** *m/z* [M+H]$^+$: calc. 386.00, found 385.95

**HPLC**: t$_R$ = 3.210 min (M2), purity ≧ 95% (UV: 254/280 nm)

*1$^5$-(Furan-2-yl)-4-thia-2,5,9-triaza-1(2,4)-pyrimidina-3(1,3)-benzenacyclononaphane 4,4-dioxide* (**7**)

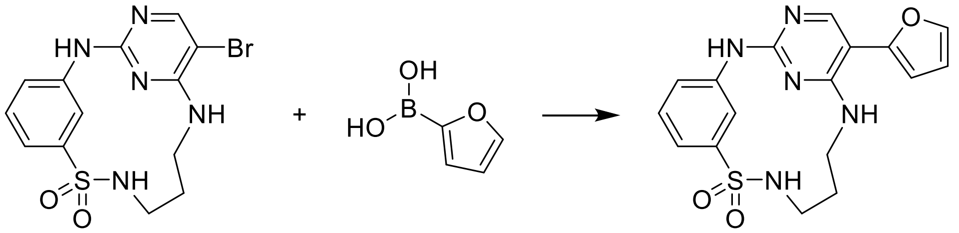

**6** (1 eq.), [1,1′-Bis-(diphenylphosphino)-ferrocen]-dichloro-palladium(II) (0.05 eq.), 2 M aq. potassium carbonate (2 eq.) and 2-furylboronic acid (1.2 eq.), were dissolved in 1,4-dioxane and DMF (1:1, 0.125 M) and degassed with Argon. The reaction mixture was then heated under µW irradiation at 100 °C for 2 h. The crude product was purified by flash column chromatography on silica gel using dichloromethane / ethyl acetate as an eluent to afford the product as a white solid (60 mg, 62%).

**$^{1}$H-NMR** (400 MHz, DMSO-d$_6$) δ 9.75 (s, 1H), 9.42 (t, J = 2.0 Hz, 1H), 8.23 (s, 1H), 7.78 (t, J = 6.0 Hz, 1H), 7.71 (dd, J = 1.9, 0.7 Hz, 1H), 7.42 – 7.37 (m, 1H), 7.31 (ddd, J = 8.3, 2.0, 1.1 Hz, 2H), 7.18 (t, J = 5.5 Hz, 1H), 6.71 (dd, J = 3.4, 0.7 Hz, 1H), 6.60 (dd, J = 3.4, 1.9 Hz, 1H), 3.52 (dt, J = 11.6, 5.5 Hz, 2H), 3.33 – 3.26 (m, 2H), 1.94 – 1.83 (m, 2H).

**$^{13}$C NMR** (101 MHz, DMSO) δ 158.36, 157.34, 153.76, 149.05, 144.07, 141.89, 141.46, 128.96, 121.04, 117.52, 116.47, 111.53, 105.47, 101.17, 28.76.

**MS (ESI+)** *m/z* [M+H]$^+$: calc. 372.11, found 372.05

**HRMS** calculated *m/z* for $[C_{17}H_{17}N_5O_3S+H]^+$: 372.11249, found: 372.11386

**HPLC**: t$_R$ = 3.326 min (M2), purity ≧ 95% (UV: 254/280 nm)

*$1^5$-(4-Hydroxyphenyl)-4-thia-2,5,9-triaza-1(2,4)-pyrimidina-3(1,3)-benzenacyclononaphane 4,4-dioxide* (**8**)

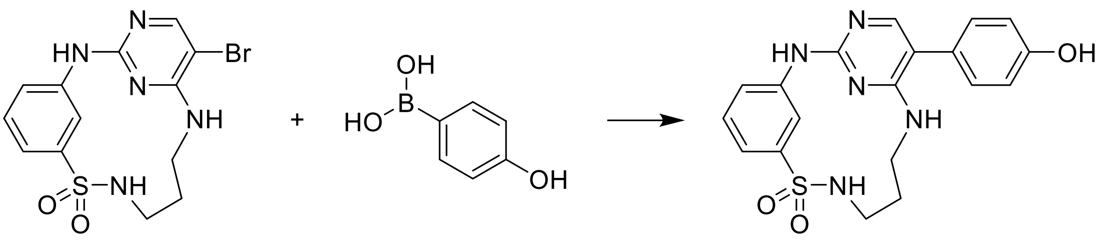

**8** was prepared according to the method for **7**, using 4-hydroxyphenylboronic acid and DME as solvent. The product was obtained as a white solid (27 mg, 26%).

**$^1$H-NMR** (400 MHz, DMSO-d$_6$) δ 9.52 (s, 1H), 9.51 (s, 1H), 9.46 (t, J = 2.0 Hz, 1H), 7.74 (t, J = 6.0 Hz, 1H), 7.72 (s, 1H), 7.40 – 7.34 (m, 1H), 7.30 – 7.26 (m, 2H), 7.20 – 7.15 (m, 2H), 6.87 – 6.82 (m, 2H), 6.79 (t, J = 5.5 Hz, 1H), 3.46 – 3.36 (m, 2H), 3.27 (q, J = 5.6 Hz, 2H), 1.83 (m, 2H).

**$^{13}$C NMR** (101 MHz, DMSO) δ 159.27, 158.36, 156.74, 154.47, 144.04, 141.93, 129.80, 128.85, 125.36, 120.76, 117.04, 116.38, 115.82, 111.16, 28.83.

**MS (ESI+)** *m/z* [M+H]$^+$: calc. 398.12, found 398.10

**HRMS** calculated *m/z* for [$C_{19}H_{19}N_5O_3S$+H]$^+$: 398.12814, found: 398.12837

**HPLC**: $t_R$ = 4.957 min (M1), purity ≧ 95% (UV: 254/280 nm)

*$1^5$-Phenyl-4-thia-2,5,9-triaza-1(2,4)-pyrimidina-3(1,3)-benzenacyclononaphane 4,4-dioxide* (**9**)

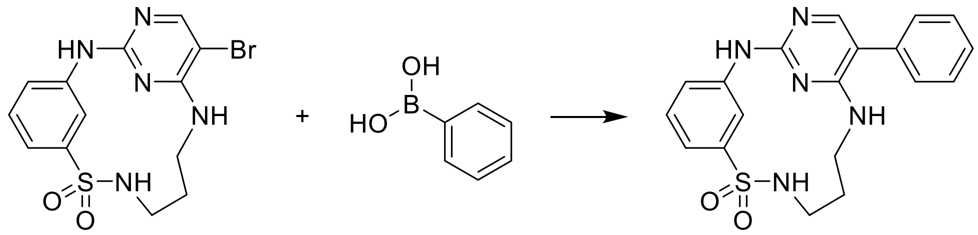

**9** was prepared according to the method for **7**, using benzeneboronic acid. The product was obtained as a white solid (53 mg, 64%).

**$^1$H-NMR** (400 MHz, DMSO-d$_6$) δ 9.60 (s, 1H), 9.46 (t, J = 2.0 Hz, 1H), 7.80 (s, 1H), 7.76 (t, J = 6.0 Hz, 1H), 7.50 – 7.33 (m, 7H), 7.30 (ddd, J = 8.8, 4.7, 1.7 Hz, 2H), 6.94 (t, J = 5.5 Hz, 1H), 3.46 – 3.37 (m, 2H), 3.28 (q, J = 5.8 Hz, 2H), 1.85 (q, J = 9.0, 8.5 Hz, 2H).

**$^{13}$C NMR** (101 MHz, DMSO) δ 159.04, 158.64, 155.02, 144.05, 141.80, 135.11, 129.01, 128.88, 128.55, 127.13, 120.87, 117.21, 116.47, 110.96, 28.79.

**MS (ESI+)** *m/z* [M+H]$^+$: calc. 382.13, found 382.05

**HRMS** calculated *m/z* for [$C_{19}H_{19}N_5O_2S$+H]$^+$: 382.13322, found: 382.13287

**HPLC**: $t_R$ = 5.017 min (M1), purity ≧ 95% (UV: 254/280 nm)

*1$^{5}$-(3-(Methylthio)phenyl)-4-thia-2,5,9-triaza-1(2,4)-pyrimidina-3(1,3)-benzenacyclononaphane 4,4-dioxide* (**10**)

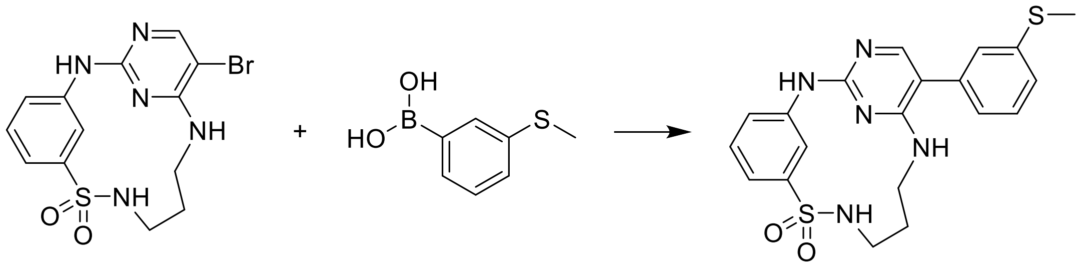

**10** was prepared according to the method for **7**, using 3-(methylthio)phenylboronic acid. The product was obtained as a white solid (78 mg, 74%).

**$^{1}$H-NMR** (400 MHz, DMSO-d$_6$) δ 9.62 (s, 1H), 9.45 (t, J = 2.0 Hz, 1H), 7.82 (s, 1H), 7.76 (t, J = 6.0 Hz, 1H), 7.42 – 7.36 (m, 2H), 7.33 – 7.27 (m, 2H), 7.26 – 7.20 (m, 2H), 7.15 (dt, J = 7.6, 1.4 Hz, 1H), 6.99 (t, J = 5.5 Hz, 1H), 3.46 – 3.36 (m, 2H), 3.28 (q, J = 5.8 Hz, 2H), 1.91 – 1.80 (m, 2H).

**$^{13}$C NMR** (101 MHz, DMSO) δ 158.99, 158.71, 155.09, 144.04, 141.77, 138.94, 135.91, 129.41, 128.87, 125.48, 125.03, 124.70, 120.88, 117.23, 116.47, 110.57, 28.80, 14.47.

**MS (ESI+)** *m/z* [M+H]$^+$: calc. 428.11, found 428.10

**HRMS** calculated *m/z* for [$C_{20}H_{21}N_5O_2S_2$+H]$^+$: 428.12094, found: 428.12063

**HPLC**: $t_R$ = 5.769 min (M1), purity ≧ 95% (UV: 254/280 nm)

*1$^{5}$-(3-Methoxyphenyl)-4-thia-2,5,9-triaza-1(2,4)-pyrimidina-3(1,3)-benzenacyclononaphane 4,4-dioxide* (**11**)

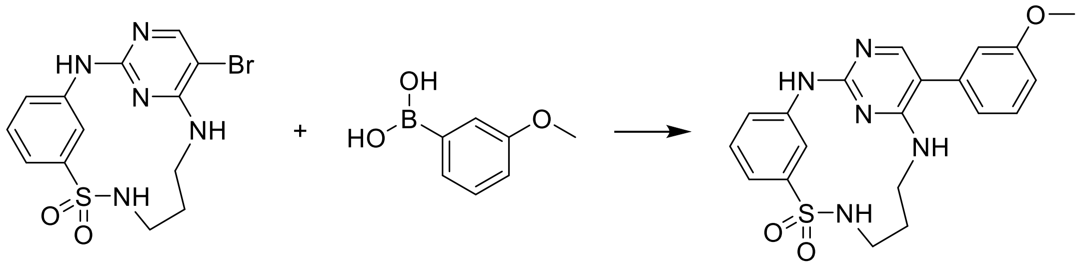

**11** was prepared according to the method for **7**, using 3-methoxyphenylboronic acid. The product was obtained as a white solid (45 mg, 42%).

**$^{1}$H-NMR** (400 MHz, DMSO-d$_6$) δ 9.60 (s, 1H), 9.46 (t, J = 2.0 Hz, 1H), 7.82 (s, 1H), 7.76 (t, J = 6.0 Hz, 1H), 7.42 – 7.34 (m, 2H), 7.32 – 7.27 (m, 2H), 6.99 – 6.89 (m, 4H), 3.80 (s, 3H), 3.46 – 3.37 (m, 2H), 3.28 (q, J = 5.7 Hz, 2H), 1.90 – 1.79 (m, 2H).

**$^{13}$C NMR** (101 MHz, DMSO) δ 159.62, 158.98, 158.64, 154.94, 144.04, 141.79, 136.44, 130.05, 128.87, 120.86, 120.72, 117.20, 116.45, 113.84, 112.94, 110.85, 55.01, 28.80.

**MS (ESI+)** *m/z* [M+H]$^+$: calc. 412.14, found 412.10

**HRMS** calculated *m/z* for [$C_{20}H_{21}N_5O_3S$+H]$^+$: 412.14379, found: 412.14404

**HPLC**: $t_R$ = 5.542 min (M1), purity ≧ 95% (UV: 254/280 nm)

*4-(4,4-Dioxido-4-thia-2,5,9-triaza-1(2,4)-pyrimidina-3(1,3)-benzenacyclononaphane-15-yl)-benzamide* (**12**)

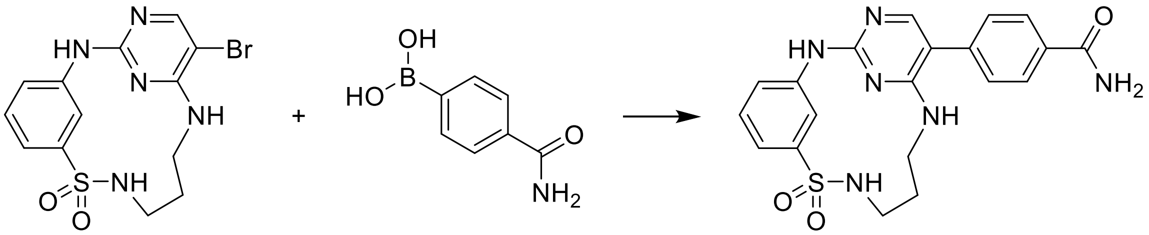

**12** was prepared according to the method for **7**, using (4-carbamoylphenyl)boronic acid. The product was obtained as a white solid (26 mg, 23%).

**$^1$H-NMR** (400 MHz, DMSO-$d_6$) δ 9.66 (s, 1H), 9.45 (t, J = 2.0 Hz, 1H), 8.07 – 7.92 (m, 3H), 7.86 (s, 1H), 7.77 (t, J = 6.0 Hz, 1H), 7.47 (d, J = 7.9 Hz, 2H), 7.43 – 7.25 (m, 4H), 7.06 (t, J = 5.5 Hz, 1H), 3.46 – 3.37 (m, 2H), 3.31 – 3.24 (m, 2H), 1.92 – 1.79 (m, 2H).

**$^{13}$C NMR** (101 MHz, DMSO) δ 167.42, 158.96, 158.81, 155.28, 144.06, 141.70, 138.15, 132.65, 128.91, 128.26, 128.17, 120.94, 117.32, 116.52, 110.23, 28.76.

**MS (ESI+)** *m/z* $[M+H]^+$: calc. 425.13, found 425.10

**HRMS** calculated *m/z* for $[C_{20}H_{20}N_6O_3S+H]^+$: 425.13904, found: 425.13894

**HPLC**: $t_R$ = 3.150 min (M2), purity ≧ 95% (UV: 254/280 nm)

*1$^5$-(Pyridin-3-yl)-4-thia-2,5,9-triaza-1(2,4)-pyrimidina-3(1,3)-benzenacyclononaphane 4,4-dioxide* (**13**)

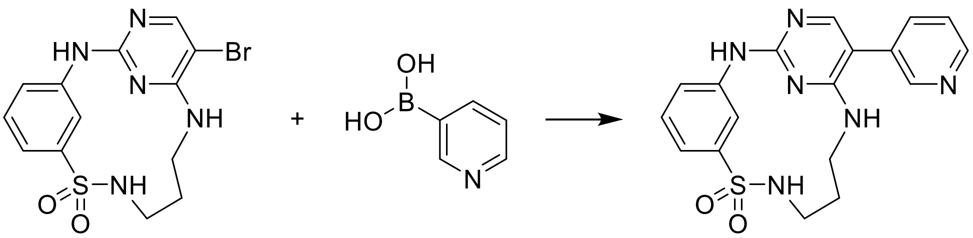

**13** was prepared according to the method for **7**, using (pyridin-3-yl)boronic acid. The product was obtained as a white solid (36 mg, 42%).

**$^1$H-NMR** (400 MHz, DMSO-$d_6$) δ 9.68 (s, 1H), 9.44 (s, 1H), 8.67 – 8.49 (m, 2H), 7.89 – 7.73 (m, 3H), 7.47 (dd, J = 7.8, 4.8 Hz, 1H), 7.40 (t, J = 7.8 Hz, 1H), 7.31 (t, J = 6.5 Hz, 2H), 7.16 (d, J = 5.6 Hz, 1H), 3.44 – 3.37 (m, 2H), 3.31 – 3.26 (m, 2H), 1.92 – 1.79 (m, 2H).

**$^{13}$C NMR** (101 MHz, DMSO) δ 159.25, 158.99, 155.48, 149.28, 148.12, 144.04, 141.65, 136.32, 131.11, 128.92, 123.91, 121.01, 117.42, 116.62, 107.72, 28.75.

**MS (ESI+)** *m/z* $[M+H]^+$: calc. 383.12, found 383.10

**HRMS** calculated *m/z* for $[C_{18}H_{18}N_6O_2S+H]^+$: 383.12847, found: 383.12686

**HPLC**: $t_R$ = 3.012 min (M2), purity ≧ 95% (UV: 254/280 nm)

*3-(4,4-Dioxido-4-thia-2,5,9-triaza-1(2,4)-pyrimidina-3(1,3)-benzenacyclononaphane-15-yl)-N-methylbenzamide* (**14**)

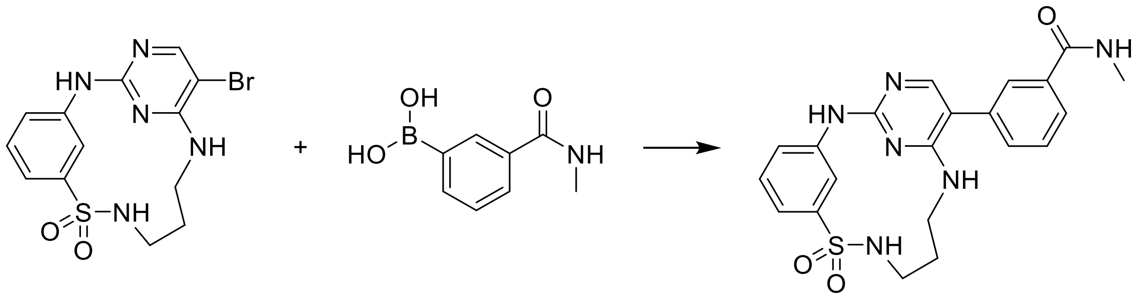

**14** was prepared according to the method for **7**, using (3-methylcarbamoyl)phenyl)boronic acid. The product was obtained as a white solid (11 mg, 9%).

**$^{1}$H-NMR** (400 MHz, DMSO-$d_6$) δ 9.66 (s, 1H), 9.47 (t, J = 2.1 Hz, 1H), 8.48 (q, J = 4.5 Hz, 1H), 7.87 (s, 1H), 7.84 – 7.76 (m, 3H), 7.56 – 7.52 (m, 2H), 7.43 – 7.38 (m, 1H), 7.31 (ddt, J = 7.5, 6.5, 1.3 Hz, 2H), 7.04 (t, J = 5.5 Hz, 1H), 3.47 – 3.38 (m, 2H), 3.30 (q, J = 5.7 Hz, 2H), 2.81 (d, J = 4.5 Hz, 3H), 1.91 – 1.82 (m, 2H).

**$^{13}$C NMR** (101 MHz, DMSO) δ 166.57, 159.01, 158.76, 155.32, 144.06, 141.74, 135.30, 135.12, 131.16, 129.02, 128.91, 127.09, 126.00, 120.89, 117.27, 116.45, 110.43, 28.78, 26.26.

**MS (ESI+)** *m/z* $[M+H]^+$: calc. 439.15, found 439.15

**HRMS** calculated *m/z* for $[C_{21}H_{22}N_6O_3S+H]^+$: 439.15472, found: 439.15445

**HPLC**: $t_R$ = 4.690 min (M1), purity ≧ 95% (UV: 254/280 nm)

*$1^5$-(Naphthalen-2-yl)-4-thia-2,5,9-triaza-1(2,4)-pyrimidina-3(1,3)-benzenacyclononaphane 4,4-dioxide* (**15**)

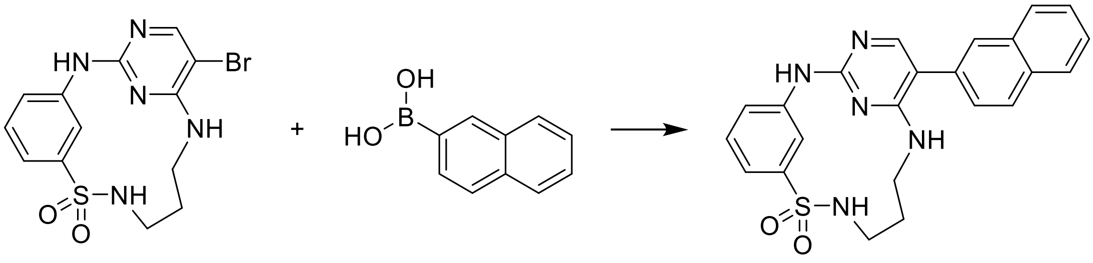

**15** was prepared according to the method for **7**, using 2-naphthalenboronic acid. The product was obtained as a white solid (50 mg, 45%).

**$^{1}$H-NMR** (400 MHz, DMSO-$d_6$) δ 9.66 (s, 1H), 9.49 (t, J = 2.0 Hz, 1H), 7.99 (d, J = 8.5 Hz, 1H), 7.97 – 7.91 (m, 4H), 7.78 (t, J = 6.0 Hz, 1H), 7.57 – 7.50 (m, 3H), 7.41 (t, J = 8.1 Hz, 1H), 7.32 (ddt, J = 8.8, 7.5, 1.3 Hz, 2H), 7.09 (t, J = 5.4 Hz, 1H), 3.43 (dt, J = 11.4, 5.5 Hz, 2H), 3.29 (q, J = 5.7 Hz, 2H), 1.95 – 1.81 (m, 2H).

**$^{13}$C NMR** (101 MHz, DMSO) δ 159.26, 158.68, 155.24, 144.07, 141.79, 133.34, 132.61, 132.11, 128.90, 128.43, 127.99, 127.47, 127.21, 127.02, 126.24, 126.02, 120.89, 117.23, 116.49, 110.96, 28.80.

**MS (ESI+)** *m/z* $[M+H]^+$: calc. 432.14, found 432.15

**HRMS** calculated *m/z* for $[C_{23}H_{21}N_5O_2S+H]^+$: 432.14887, found: 432.14862

**HPLC**: $t_R$ = 6.200 min (M1), purity ≧ 95% (UV: 254/280 nm)

*1⁵-(Benzo[d][1,3]dioxol-5-yl)-4-thia-2,5,9-triaza-1(2,4)-pyrimidina-3(1,3)-benzenacyclononaphane 4,4-dioxide* (**16**)

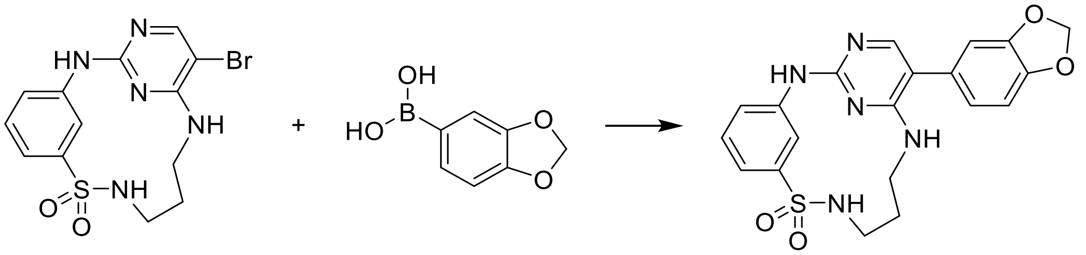

**16** was prepared according to the method for **7**, using (1,3-dioxaindan-5-yl)boronic acid. The product was obtained as a white solid (74 mg, 67%).

**$^1$H-NMR** (400 MHz, DMSO-$d_6$) δ 9.56 (s, 1H), 9.45 (t, J = 2.0 Hz, 1H), 7.78 – 7.72 (m, 2H), 7.42 – 7.35 (m, 1H), 7.29 (ddt, J = 7.5, 5.9, 1.3 Hz, 2H), 6.99 (d, J = 8.0 Hz, 1H), 6.93 (d, J = 1.7 Hz, 1H), 6.89 (t, J = 5.5 Hz, 1H), 6.82 (dd, J = 8.0, 1.8 Hz, 1H), 6.05 (s, 2H), 3.45 – 3.35 (m, 2H), 3.27 (q, J = 5.7 Hz, 2H), 1.89 – 1.78 (m, 2H).

**$^{13}$C-NMR** (101 MHz, DMSO) δ 159.18, 158.51, 154.69, 147.61, 146.51, 144.03, 141.84, 128.84, 128.73, 122.19, 120.81, 117.13, 116.42, 110.87, 109.21, 108.83, 101.05, 28.79.

**MS (ESI+)** *m/z* $[M+H]^+$: calc. 426.12, found 426.15

**HRMS** calculated *m/z* for $[C_{20}H_{19}N_5O_4S+H]^+$: 426.12305, found: 426.12328

**HPLC**: $t_R$ = 5.398 min (M1), purity ≧ 95% (UV: 254/280 nm)

*1⁵-(1-Methyl-1H-pyrazol-5-yl)-4-thia-2,5,9-triaza-1(2,4)-pyrimidina-3(1,3)-benzenacyclononaphane 4,4-dioxide* (**17**)

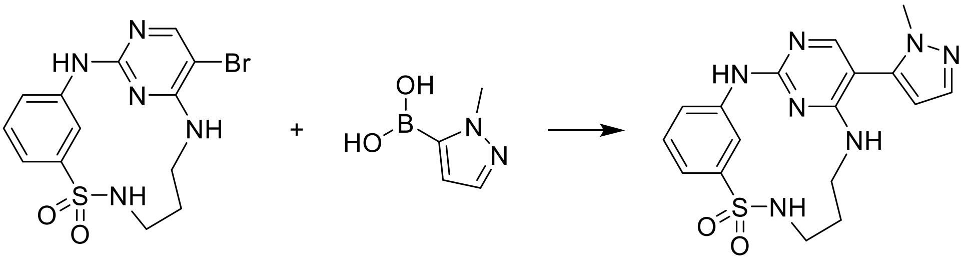

**17** was prepared according to the method for **7**, using (1-methyl-1H-pyrazol-5-yl)boronic acid. The product was obtained as a white solid (31 mg, 34%).

**$^1$H-NMR** (400 MHz, DMSO-$d_6$) δ 9.79 (s, 1H), 9.41 (t, J = 2.0 Hz, 1H), 7.83 (s, 1H), 7.76 (t, J = 5.9 Hz, 1H), 7.51 (d, J = 1.8 Hz, 1H), 7.41 (dd, J = 8.7, 6.9 Hz, 1H), 7.32 (dd, J = 7.0, 1.8 Hz, 2H), 6.98 (t, J = 5.5 Hz, 1H), 6.31 (d, J = 1.8 Hz, 1H), 3.65 (s, 3H), 3.42 – 3.35 (m, 2H), 3.27 (q, J = 5.7 Hz, 2H), 1.82 (q, J = 8.5, 7.5 Hz, 2H).

**$^{13}$C-NMR** (101 MHz, DMSO) δ 159.42, 159.05, 155.80, 143.98, 141.34, 138.24, 135.99, 128.96, 121.27, 117.76, 117.01, 107.20, 99.92, 36.54, 28.72.

**MS (ESI+)** *m/z* $[M+H]^+$: calc. 386.13, found 386.10

**HRMS** calculated *m/z* for $[C_{17}H_{19}N_7O_2S+H]^+$: 386.13937, found: 386.13921

**HPLC**: $t_R$ = 3.089 min (M2), purity ≧ 95% (UV: 254/280 nm)

*1$^5$-(4-(Tert-butyl)phenyl)-4-thia-2,5,9-triaza-1(2,4)-pyrimidina-3(1,3)-benzenacyclononaphane 4,4-dioxide* (**18**)

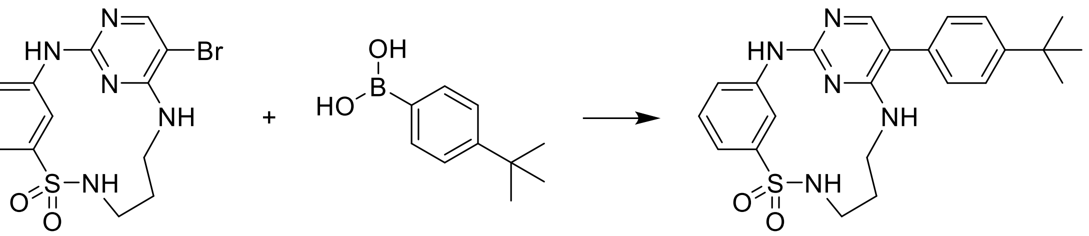

**18** was prepared according to the method for **7**, using (1,3-dioxaindan-5-yl)boronic acid. The product was obtained as a white solid (67 mg, 57%).

**$^1$H-NMR** (400 MHz, DMSO-$d_6$) δ 9.57 (s, 1H), 9.46 (t, J = 2.0 Hz, 1H), 7.79 (s, 1H), 7.75 (t, J = 6.0 Hz, 1H), 7.50 – 7.46 (m, 2H), 7.41 – 7.36 (m, 1H), 7.35 – 7.27 (m, 4H), 6.93 (t, J = 5.5 Hz, 1H), 3.46 – 3.37 (m, 2H), 3.27 (q, J = 5.7 Hz, 2H), 1.90 – 1.80 (m, 2H), 1.32 (s, 9H).

**$^{13}$C-NMR** (101 MHz, DMSO) δ 159.06, 158.54, 154.95, 149.42, 144.04, 141.83, 132.17, 128.85, 128.13, 125.80, 120.83, 117.15, 116.43, 110.77, 34.29, 31.14, 28.80.

**MS (ESI+)** *m/z* $[M+H]^+$: calc. 438.19, found 438.15

**HRMS** calculated *m/z* for $[C_{23}H_{27}N_5O_2S+H]^+$: 438.19582, found: 438.19548

**HPLC**: $t_R$ = 6.185 min (M1), purity ≧ 95% (UV: 254/280 nm)

*1$^5$-([1,1'-Biphenyl]-4-yl)-4-thia-2,5,9-triaza-1(2,4)-pyrimidina-3(1,3)-benzenacyclononaphane 4,4-dioxide* (**19**)

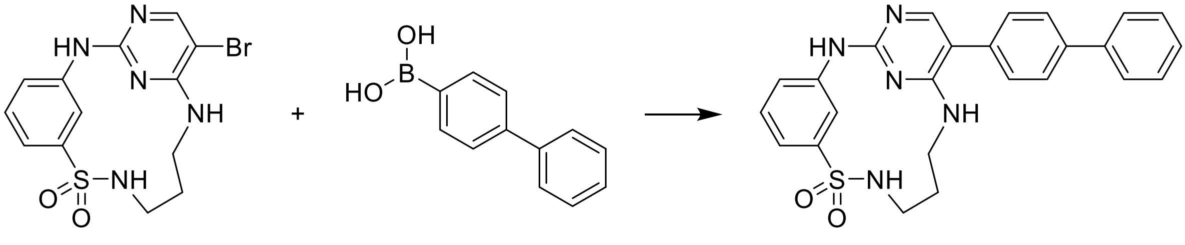

**19** was prepared according to the method for **7**, using [1,1'-biphenyl]-4-ylboronic acid. The product was obtained as a white solid (42 mg, 35%).

**$^1$H-NMR** (400 MHz, DMSO-d$_6$) δ 10.74 (s, 1H), 9.17 (t, J = 1.9 Hz, 1H), 8.18 (t, J = 5.5 Hz, 1H), 7.93 (s, 1H), 7.87 – 7.79 (m, 3H), 7.75 – 7.71 (m, 2H), 7.55 – 7.48 (m, 6H), 7.43 – 7.35 (m, 2H), 3.50 – 3.39 (m, 2H), 3.30 (q, J = 5.7 Hz, 2H), 1.91 – 1.79 (m, 2H).

**$^{13}$C-NMR** (101 MHz, DMSO) δ 159.98, 152.67, 152.64, 143.83, 140.13, 139.48, 139.14, 131.35, 129.58, 129.41, 129.11, 127.78, 127.39, 126.64, 122.63, 119.96, 118.69, 111.94, 28.20.

**MS (ESI+)** *m/z* $[M+H]^+$: calc. 458.16, found 458.10

**HRMS** calculated *m/z* for $[C_{25}H_{23}N_5O_2S+H]^+$: 458.16452, found: 458.16405

**HPLC**: $t_R$ = 3.569 min (M2), purity ≧ 95% (UV: 254/280 nm)

*$1^5$-([1,1'-Biphenyl]-3-yl)-4-thia-2,5,9-triaza-1(2,4)-pyrimidina-3(1,3)-benzenacyclononaphane 4,4-dioxide* (**20**)

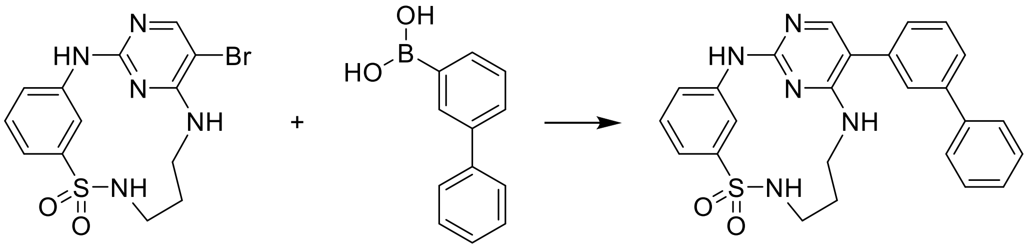

**20** was prepared according to the method for **7**, using [1,1'-biphenyl]-3-ylboronic acid. The product was obtained as a white solid (16 mg, 13%).

**$^1$H-NMR** (400 MHz, DMSO-d$_6$) δ 9.63 (s, 1H), 9.48 (s, 1H), 7.90 (s, 1H), 7.80 – 7.71 (m, 3H), 7.66 – 7.61 (m, 2H), 7.55 (t, J = 7.7 Hz, 1H), 7.48 (t, J = 7.5 Hz, 2H), 7.43 – 7.36 (m, 3H), 7.30 (t, J = 8.0 Hz, 2H), 7.07 (s, 1H), 3.49 – 3.38 (m, 2H), 3.31 – 3.24 (m, 2H), 1.93 – 1.82 (m, 2H).

**$^{13}$C-NMR** (101 MHz, DMSO) δ 159.10, 158.72, 155.29, 144.07, 141.81, 141.00, 140.17, 135.80, 129.62, 128.93, 128.92, 127.65, 127.56, 126.99, 126.98, 125.62, 120.88, 117.23, 116.47, 110.89, 28.82.

**MS (ESI+)** *m/z* $[M+H]^+$: calc. 458.16, found 458.15

**HRMS** calculated *m/z* for $[C_{25}H_{23}N_5O_2S+H]^+$: 458.16452, found: 458.16399

**HPLC**: $t_R$ = 6.750 min (M1), purity ≧ 95% (UV: 254/280 nm)

*1⁵-(2-Fluoro-5-methoxyphenyl)-4-thia-2,5,9-triaza-1(2,4)-pyrimidina-3(1,3)-benzenacyclononaphane 4,4-dioxide* (**21**)

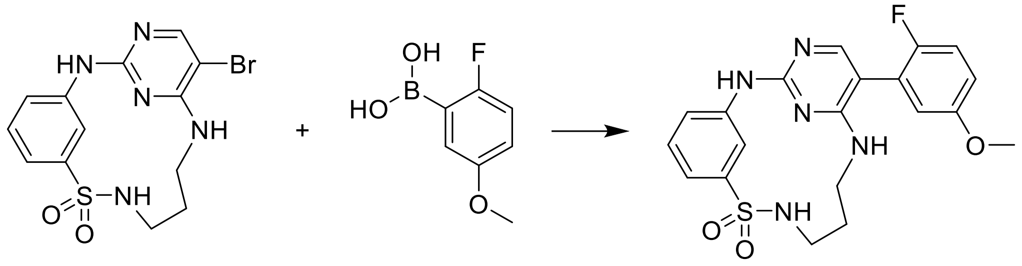

**21** was prepared according to the method for **7**, using 2-fluoro-5-methoxyphenylboronic acid. The product was obtained as a white solid (18 mg, 16%).

**¹H-NMR** (400 MHz, DMSO-d$_6$) δ 9.64 (s, 1H), 9.45 (t, J = 2.0 Hz, 1H), 7.79 (s, 1H), 7.76 (t, J = 6.0 Hz, 1H), 7.39 (t, J = 7.7 Hz, 1H), 7.34 – 7.27 (m, 2H), 7.21 (t, J = 9.2 Hz, 1H), 6.99 – 6.93 (m, 1H), 6.92 – 6.86 (m, 2H), 3.77 (s, 3H), 3.42 – 3.35 (m, 2H), 3.27 (q, J = 5.7 Hz, 2H), 1.88 – 1.78 (m, 2H).

**¹³C-NMR** (101 MHz, DMSO) δ 159.08, 155.79, 155.63 (d, J = 1.6 Hz), 155.26, 152.91, 142.85 (d, J = 231.6 Hz), 128.87, 122.91 (d, J = 17.9 Hz), 120.94, 117.31, 116.61, 116.58 (d, J = 24.1 Hz), 116.18 (d, J = 3.2 Hz), 114.73 (d, J = 7.9 Hz), 105.03, 55.59, 28.74.

**MS (ESI+)** *m/z* $[M+H]^+$: calc. 430.13, found 430.10

**HRMS** calculated *m/z* for $[C_{20}H_{20}FN_5O_3S+H]^+$: 430.13436, found: 430.13388

**HPLC**: $t_R$ = 5.726 min (M1), purity ≧ 95% (UV: 254/280 nm)

*1⁵-(3-Fluoro-5-methoxyphenyl)-4-thia-2,5,9-triaza-1(2,4)-pyrimidina-3(1,3)-benzenacyclononaphane 4,4-dioxide* (**22**)

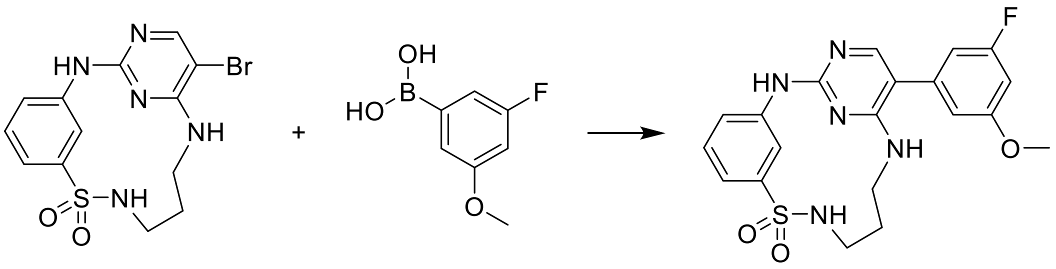

**22** was prepared according to the method for **7**, using 3-fluoro-5-methoxyphenylboronic acid. The product was obtained as a white solid (47 mg, 42%).

**¹H-NMR** (400 MHz, DMSO-d$_6$) δ 9.65 (s, 1H), 9.44 (t, J = 2.1 Hz, 1H), 7.86 (s, 1H), 7.76 (t, J = 6.0 Hz, 1H), 7.39 (t, J = 7.8 Hz, 1H), 7.33 – 7.27 (m, 2H), 7.07 (t, J = 5.5 Hz, 1H), 6.82 – 6.77 (m, 3H), 3.81 (s, 3H), 3.44 – 3.36 (m, 2H), 3.28 (q, J = 5.9 Hz, 2H), 1.90 – 1.80 (m, 2H).

**¹³C-NMR** (101 MHz, DMSO) δ 164.44, 162.03, 161.05, 160.93, 158.81 (d, J = 2.2 Hz), 155.15, 142.86 (d, J = 237.3 Hz), 137.91, 128.89, 120.92, 117.32, 116.49, 110.36 (d, J = 2.0 Hz), 109.91 (d, J = 2.5 Hz), 107.43 (d, J = 21.9 Hz), 100.39 (d, J = 24.9 Hz), 55.61, 28.77.

**MS (ESI+)** *m/z* $[M+H]^+$: calc. 430.13, found 430.05

**HRMS** calculated *m/z* for $[C_{20}H_{20}FN_5O_3S+H]^+$: 430.13436, found: 430.13409

**HPLC**: $t_R$ = 5.922 min (M1), purity ≧ 95% (UV: 254/280 nm)

*1⁵-(4-Fluoro-3-methoxyphenyl)-4-thia-2,5,9-triaza-1(2,4)-pyrimidina-3(1,3)-benzenacyclononaphane 4,4-dioxide* (**23**)

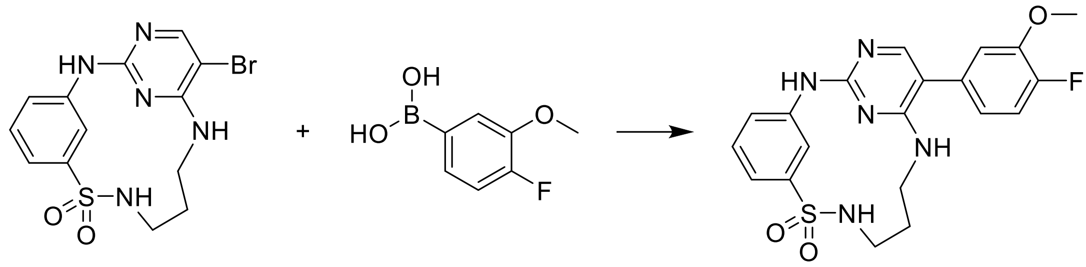

**23** was prepared according to the method for **7**, using 4-fluoro-3-methoxyphenylboronic acid. The product was obtained as a white solid (21 mg, 19%).

**$^1$H-NMR** (400 MHz, DMSO-d$_6$) δ 9.61 (s, 1H), 9.46 (t, J = 2.0 Hz, 1H), 7.83 (s, 1H), 7.76 (t, J = 6.0 Hz, 1H), 7.39 (t, J = 7.8 Hz, 1H), 7.32 – 7.24 (m, 3H), 7.13 (dd, J = 8.5, 2.1 Hz, 1H), 6.99 (t, J = 5.4 Hz, 1H), 6.92 (ddd, J = 8.3, 4.4, 2.1 Hz, 1H), 3.88 (s, 3H), 3.45 – 3.37 (m, 2H), 3.28 (q, J = 5.7 Hz, 2H), 1.89 – 1.80 (m, 2H).

**$^{13}$C-NMR** (101 MHz, DMSO) δ 159.06, 158.69, 154.97, 150.83 (d, J = 243.7 Hz), 147.26 (d, J = 10.7 Hz), 144.05, 141.79, 131.81 (d, J = 3.6 Hz), 128.88, 121.07 – 120.77 (m), 117.21, 116.44, 116.18 (d, J = 18.0 Hz), 114.33, 110.38, 55.88, 28.83.

**MS (ESI+)** *m/z* $[M+H]^+$: calc. 430.13, found 430.10

**HRMS** calculated *m/z* for $[C_{20}H_{20}FN_5O_3S+H]^+$: 430.13436, found: 430.13415

**HPLC**: $t_R$ = 3.297 min (M2), purity ≧ 95% (UV: 254/280 nm)

*N-methyl-3-nitrobenzenesulfonamide* (**24)**

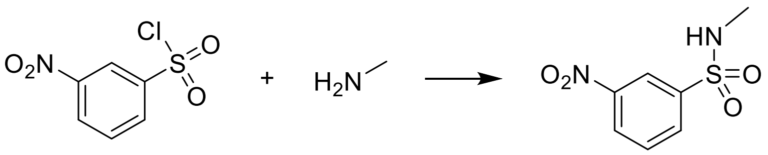

2 M methylamine in THF (1.1 eq.) was added to a solution of 3-nitrobenzenesulfonyl chloride (1 eq.) and triethylamine (2.2 eq.) in THF (0.1 M) and was stirred for 1 h at rt. After completion, THF was removed *in vacuo* and the residue was diluted with ethyl acetate. The organic phase was washed with 10% citric acid, sat. aq. $NaHCO_3$, and brine, dried over $Na_2SO_4$ and concentrated under reduced pressure, to yield the desired product as a white solid in quantitative yield (489 mg).

**$^1$H-NMR** (250 MHz, DMSO-d6) δ 8.53 – 8.46 (m, 2H), 8.23 – 8.17 (m, 1H), 7.96 – 7.88 (m, 1H), 7.83 (q, J = 4.9 Hz, 1H), 2.46 (d, J = 4.9 Hz, 3H).

*3-Amino-N-methylbenzenesulfonamide* **(25)**

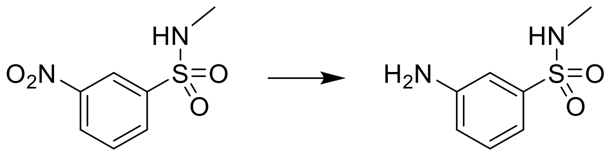

**24** was dissolved in MeOH/water 9:1 (0.11 M), iron (7 eq.) and ammonium chloride (7 eq.) were added, and the resulting mixture was heated under reflux for 4 h. The reaction crude was filtered over Celite and concentrated under reduced pressure. The residue was diluted with ethyl acetate and washed with 10% aq. citric acid, sat. aq. $NaHCO_3$ and brine, then dried over $MgSO_4$, filtered over cotton, the product was used without further purification.

**$^1$H-NMR** (250 MHz, DMSO-d6) δ 7.25 – 7.15 (m, 2H), 6.98 – 6.95 (m, 1H), 6.85 (dd, J = 7.6, 1.8 Hz, 1H), 6.75 (dd, J = 8.0, 2.3 Hz, 1H), 5.55 (s, 2H), 2.39 (d, J = 5.0 Hz, 3H).

*5-Bromo-2-chloro-N-ethylpyrimidin-4-amine* **(26)**

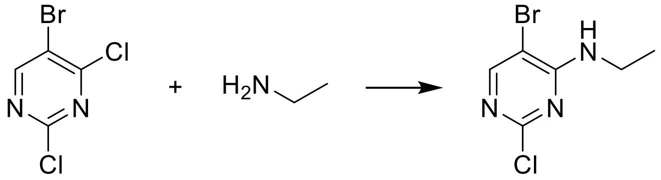

2 M ethylamine in THF (1.1 eq.) was added to a solution of **1** (1 eq.) and triethylamine (2.2 eq.) in THF (0.1 M) and was stirred for 1 h at rt. After completion, THF was removed *in vacuo* and the residue was diluted with ethyl acetate. The organic phase was washed with 10% citric acid, sat. aq. $NaHCO_3$, and brine, dried over $Na_2SO_4$ and concentrated under reduced pressure, to yield the desired product in quantitative yield (473 mg).

**$^1$H-NMR** (250 MHz, DMSO-d6) δ 8.21 (s, 1H), 7.73 (t, J = 5.7 Hz, 1H), 3.39 (p, 2H), 1.12 (t, 3H).

*3-((5-Bromo-4-(ethylamino)pyrimidin-2-yl)amino)-N-methylbenzenesulfonamide* **(27)**

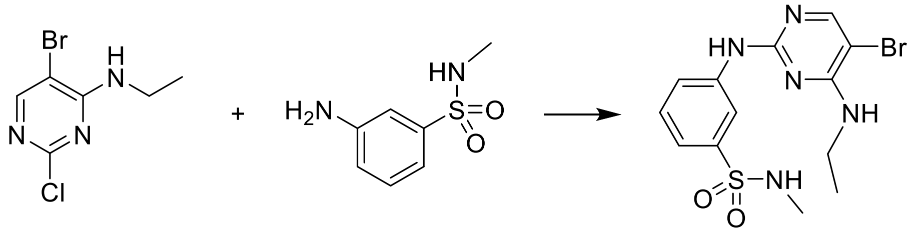

4 N HCl in 1,4-dioxane (4 eq.) was added to solution of **25** and **26** in acetonitrile/water 9:1 (0.06 M). The mixture was heated under reflux for 24 h after which time it was quenched with $NaHCO_3$ and concentrated under reduced pressure. The crude product was purified by reverse phase flash column chromatography, using acetonitrile/water as eluent. The desired product was obtained as a white solid (319 mg, 48%).

**$^{1}$H-NMR** (400 MHz, DMSO-d6) δ 9.58 (s, 1H), 8.48 (t, J = 2.0 Hz, 1H), 8.04 (s, 1H), 7.79 (ddd, J = 8.3, 2.3, 1.0 Hz, 1H), 7.45 (t, J = 8.0 Hz, 1H), 7.33 – 7.26 (m, 2H), 7.07 (t, J = 5.8 Hz, 1H), 3.48 (p, 2H), 2.42 (d, J = 5.0 Hz, 3H), 1.17 (t, J = 7.1 Hz, 3H).

**$^{13}$C-NMR** (101 MHz, DMSO) δ 158.21, 157.93, 155.73, 155.00, 141.52, 139.55, 129.16, 121.76, 118.48, 116.25, 35.57, 28.73, 14.60.

**MS (ESI+)** *m/z* $[M+H]^+$: calc. 386.05, found 386.00

**HRMS** calculated *m/z* for $[C_{13}H_{16}BrN_5O_2S+H]^+$: 386.02808,found: 386.02932

**HPLC**: $t_R$ = 5.545 min (M1), purity ≧ 95% (UV: 254/280 nm)

*3-((4-(Ethylamino)-5-(4-fluoro-3-methoxyphenyl)pyrimidin-2-yl)amino)-N-methylbenzene-sulfonamide* (**28**)

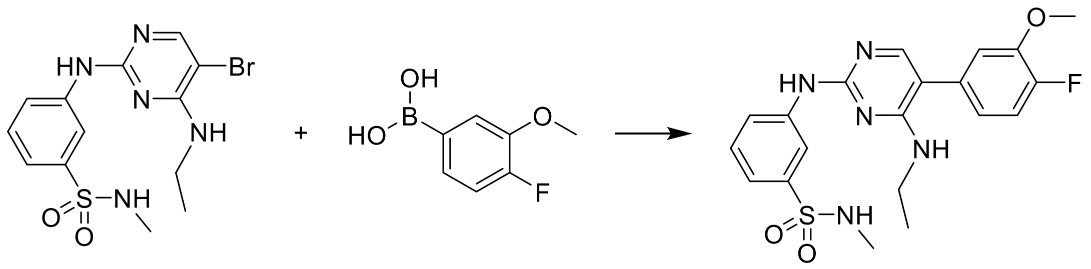

**28** was prepared according to the method for **7**, using **27** and 4-fluoro-3-methoxyphenylboronic acid. The product was obtained as a white solid (42 mg, 70%).

**$^{1}$H-NMR** (400 MHz, DMSO-d6) δ 9.51 (s, 1H), 8.58 (s, 1H), 7.89 – 7.76 (m, 2H), 7.45 (td, J = 8.0, 3.0 Hz, 1H), 7.28 (td, J = 8.3, 3.9 Hz, 3H), 7.18 – 7.09 (m, 1H), 6.95 – 6.89 (m, 1H), 6.64 (d, J = 6.6 Hz, 1H), 3.88 (t, J = 3.0 Hz, 3H), 3.48 (p, J = 6.3 Hz, 2H), 2.44 (t, J = 3.7 Hz, 3H), 1.14 (td, J = 7.2, 2.9 Hz, 3H).

**$^{13}$C-NMR** (101 MHz, DMSO-d6) δ 159.44, 158.60, 154.27, 150.86 (d, J = 243.9 Hz), 147.27 (d, J = 10.7 Hz), 141.93, 139.55, 131.84 (d, J = 3.7 Hz), 129.12, 121.58, 121.05 (d, J = 7.0 Hz), 118.13, 116.31, 116.15, 114.56 – 114.38 (m), 110.86, 55.85, 35.35, 28.75, 14.68.

**MS (ESI+)** *m/z* $[M+H]^+$: calc. 432.14, found 432.05

**HRMS** calculated *m/z* for $[C_{20}H_{22}FN_5O_3S+H]^+$: 432.15001,found: 432.14949

**HPLC**: $t_R$ = 3.436 min (M2), purity ≧ 95% (UV: 254/280 nm)

*4-Bromo-1-fluoro-2-isopropoxybenzene* (**31**)

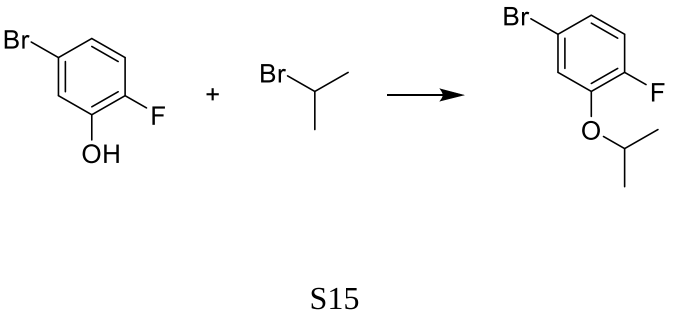

A round-bottom flask was charged with 5-bromo-2-fluorophenol (**29**) (1 eq.), which was dissolved with DMF (0.5 M), followed by addition of cesium carbonate (2 eq.) and 2-bromopropane (2 eq.). The reaction mixture was stirred for 24 h at rt. After completion the reaction mixture was diluted with water and 3x extracted with ethyl acetate. The combined organic phases were washed with brine and dried over $MgSO_4$ before purification by reverse phase flash column chromatography with water / acetonitrile as eluent. The product was afforded as a colourless oil (398 mg, 65%).

**$^1$H-NMR** (400 MHz, DMSO-d6) δ 7.37 (dd, J = 7.6, 2.3 Hz, 1H), 7.19 (dd, J = 11.2, 8.7 Hz, 1H), 7.10 (ddd, J = 8.7, 4.1, 2.3 Hz, 1H), 4.69 (hept, J = 6.0 Hz, 1H), 1.27 (d, J = 6.1 Hz, 6H).

**$^{13}$C NMR** (101 MHz, DMSO) δ 153.11, 150.68, 146.35, 146.24, 123.68, 123.61, 119.58, 119.56, 118.00, 117.80, 116.09, 116.05.

**HPLC**: $t_R$ = 6.357 min (M1), purity ≧ 95% (UV: 254/280 nm)

*4-Bromo-2-methoxybenzonitrile* (**32**)

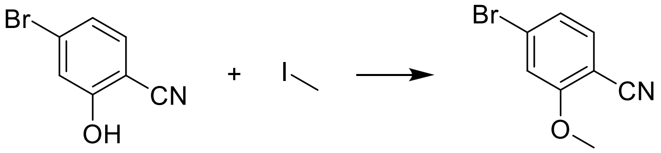

**32** was prepared according to the method for **31**, using 4-bromo-2-hydroxybenzonitrile (**30**) and iodmethane solution… The product was obtained as a … (533 mg, 91%)

**$^1$H-NMR** (250 MHz, DMSO-d6) δ 7.68 (d, J = 8.4 Hz, 1H), 7.50 (d, J = 1.7 Hz, 1H), 7.31 (dd, J = 8.2, 1.7 Hz, 1H), 3.94 (s, 3H).

**MS (ESI+)** *m/z* $[M+H]^+$: calc. 211.96, found: 211.90

**HPLC**: $t_R$ = 4.341 min (M2), purity ≧ 95% (UV: 254/280 nm)

*2-(4-Fluoro-3-isopropoxyphenyl)-4,4,5,5-tetramethyl-1,3,2-dioxaborolane* (**36**)

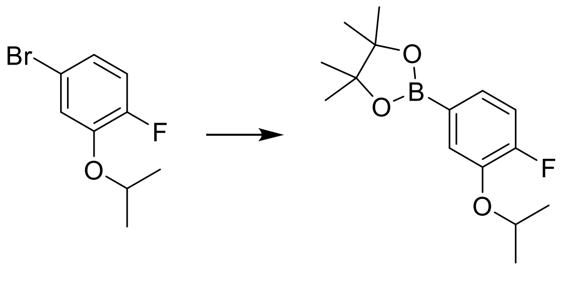

A µW vial was charged with **31** (1 eq.), bis(pinacolato)diboron (1.2 eq.), [1,1′-Bis-(diphenylphosphino)-ferrocen]-dichloro-palladium(II) (0.05 eq.), potassium acetate (3 eq.) and dissolved in 1,4-dioxane (0.2 M). The reaction mixture was degassed with argon and heated under µW irradiation at 100 °C for 2 h. The crude product was filtered over a pad of Celite, diluted with ethyl acetate, washed with water and brine, dried over $MgSO_4$ and then purified by flash column chromatography on silica gel using n-hexane / ethyl acetate as an eluent. The product was afforded as a colourless oil (87 mg, 40%).

**$^{1}$H-NMR** (400 MHz, DMSO-d6) δ 7.31 (dd, J = 8.9, 1.4 Hz, 1H), 7.26 (ddd, J = 6.7, 5.2, 1.4 Hz, 1H), 7.20 (dd, J = 11.4, 8.0 Hz, 1H), 4.62 (hept, J = 6.0 Hz, 1H), 1.29 (s, 12H), 1.27 (d, J = 6.1 Hz, 6H).

**MS (ESI+)** *m/z* $[M+H]^+$: calc. 281.16, found: 281.10

**HPLC**: $t_R$ = 7.116 min (M1)

*2-Methoxy-4-(4,4,5,5-tetramethyl-1,3,2-dioxaborolan-2-yl)benzonitrile* (**37)**

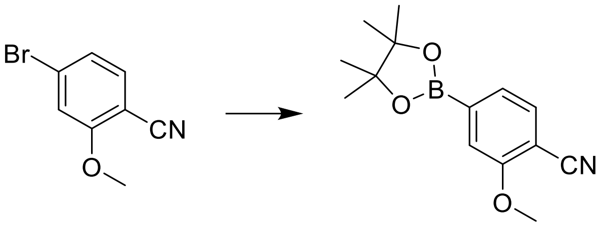

**37** was prepared to according to the method for **36**, using **32**. The product was obtained as a white solid, in a mixture of borpinacole ester and boronic acid (183 mg).

**$^{1}$H-NMR** (250 MHz, DMSO-d6) δ 8.42 (s, 2H), 7.72 (d, J = 7.7 Hz, 1H), 7.67 (d, J = 7.6 Hz, 1H), 7.58 (s, 2H), 7.46 (dd, J = 7.6, 0.9 Hz, 1H), 7.37 – 7.32 (m, 1H), 3.94 (s, 3H), 3.92 (s, 3H), 1.31 (s, 12H).

**MS (ESI+)** *m/z* $[M+H]^+$: calc. 178.06, 260.14; found: 178.00, 260.00

**HPLC**: $t_R$ = 3.511 min, 4.980 min (M2); (UV: 254/280 nm)

*2-(4-Bromo-3-methoxyphenyl)-4,4,5,5-tetramethyl-1,3,2-dioxaborolane* (**38)**

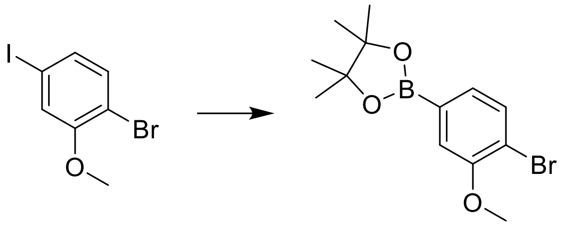

**38** was prepared to according to the method for **36**, using 1-bromo-4-iodo-2-methoxybenzene (**33**). The product was obtained as a white solid (256 mg, 26%).

**$^{1}$H-NMR** (400 MHz, DMSO-d6) δ 7.59 (d, J = 7.7 Hz, 1H), 7.23 (d, J = 1.3 Hz, 1H), 7.17 (dd, J = 7.8, 1.3 Hz, 1H), 3.87 (s, 3H), 1.30 (s, 12H).

**$^{13}$C NMR** (101 MHz, DMSO) δ 154.97, 132.83, 127.96, 117.08, 114.67, 84.01, 56.05, 24.64.

*2-(4,5-dichloro-2-methylphenyl)-4,4,5,5-tetramethyl-1,3,2-dioxaborolane* (**39**)

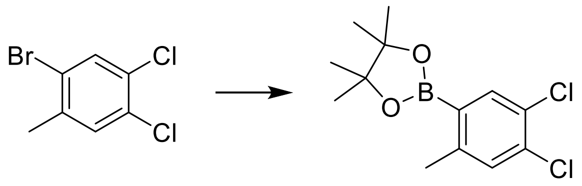

**39** was prepared to according to the method for **36**, using 2-bromo-4,5-dichlorotoluene (**34**). The product was obtained as a white solid (222 mg, 68%).

**$^{1}$H-NMR** (400 MHz, DMSO-d6) δ 7.68 (s, 1H), 7.49 (d, J = 0.8 Hz, 1H), 2.43 (d, J = 0.7 Hz, 3H), 1.30 (s, 12H).

**$^{13}$C NMR** (101 MHz, DMSO) δ 145.04, 136.54, 133.39, 131.80, 127.72, 83.98, 24.58, 20.82.

**HPLC**: $t_R$ = 9.150 min (M1), (UV: 254/280 nm)

*2-(4,5-dichloro-2-fluorophenyl)-4,4,5,5-tetramethyl-1,3,2-dioxaborolane* (**40**)

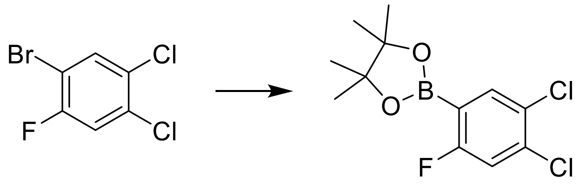

**40** was prepared to according to the method for **36**, using 4,5-Dichloro-2-fluorobromobenzene (**35**). The product was obtained as a white solid (131 mg, 35%).

**$^{1}$H-NMR** (400 MHz, DMSO-d6) δ 7.72 (d, J = 5.6 Hz, 1H), 7.65 (d, J = 8.6 Hz, 1H), 1.30 (s, 12H).

**HPLC**: $t_R$ = 4.066 min (M2), (UV: 254/280 nm)

*Methyl 4-(4,4-dioxido-4-thia-2,5,9-triaza-1(2,4)-pyrimidina-3(1,3)-benzenacyclononaphane-1$^{5}$-yl)-2-methoxybenzoate* (**41**)

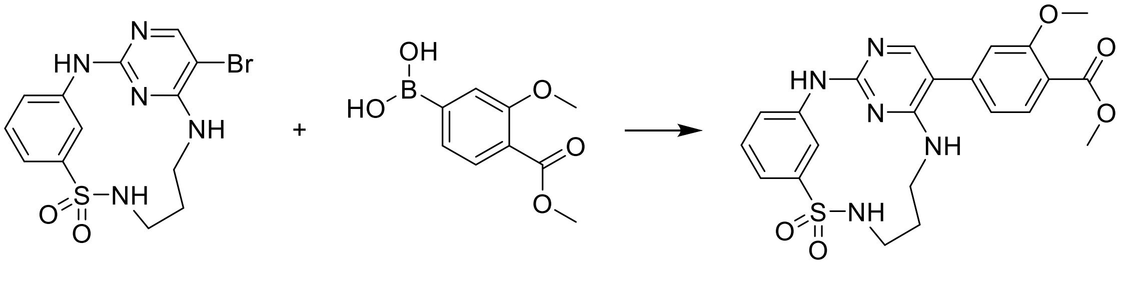

**41** was prepared according to the method for **7**, using 3-methoxy-4-methoxycarbonylphenylboronic acid. The product was obtained as a white solid (25 mg, 21%).

**$^{1}$H-NMR** (400 MHz, DMSO-$d_6$) δ 9.69 (s, 1H), 9.45 (t, J = 2.0 Hz, 1H), 7.92 (s, 1H), 7.76 (t, J = 6.0 Hz, 1H), 7.72 (d, J = 7.9 Hz, 1H), 7.42 – 7.36 (m, 1H), 7.34 – 7.28 (m, 2H), 7.16 (t, J =

5.5 Hz, 1H), 7.12 (d, J = 1.5 Hz, 1H), 7.04 (dd, J = 7.9, 1.5 Hz, 1H), 3.88 (s, 3H), 3.80 (s, 3H), 3.46 – 3.37 (m, 2H), 3.29 (q, J = 5.8 Hz, 2H), 1.91 – 1.80 (m, 2H).

**$^{13}$C-NMR** (101 MHz, DMSO) δ 166.00, 158.88, 158.84, 158.60, 155.33, 144.06, 141.65, 140.65, 131.39, 128.91, 120.95, 120.01, 118.27, 117.36, 116.50, 112.51, 110.08, 55.72, 51.87, 28.79.

**MS (ESI+)** *m/z* $[M+H]^+$: calc. 470.14, found 470.10

**HRMS** calculated *m/z* for $[C_{22}H_{23}N_5O_5S+H]^+$: 470.14927, found: 470.14887

**HPLC**: $t_R$ = 3.459 min (M2), purity ≧ 95% (UV: 254/280 nm)

*1$^5$-(4-Chloro-3-methoxyphenyl)-4-thia-2,5,9-triaza-1(2,4)-pyrimidina-3(1,3)-benzenacyclononaphane 4,4-dioxide* (**42**)

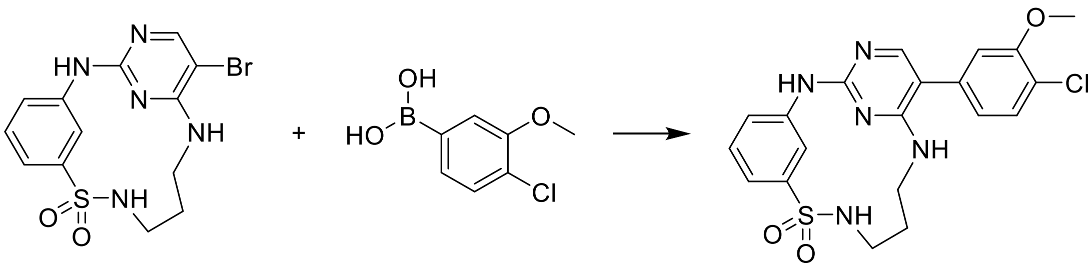

**42** was prepared according to the method for **7**, using 4-chloro-3-methoxyphenylboronic acid. The product was obtained as a white solid (28 mg, 24%).

**$^1$H-NMR** (400 MHz, DMSO-d$_6$) δ 9.64 (s, 1H), 9.46 (t, J = 2.0 Hz, 1H), 7.86 (s, 1H), 7.76 (t, J = 6.0 Hz, 1H), 7.47 (d, J = 8.1 Hz, 1H), 7.39 (t, J = 7.8 Hz, 1H), 7.30 (ddt, J = 8.6, 6.1, 1.3 Hz, 2H), 7.11 (d, J = 1.9 Hz, 1H), 7.07 (t, J = 5.4 Hz, 1H), 6.96 (dd, J = 8.1, 1.9 Hz, 1H), 3.91 (s, 3H), 3.46 – 3.37 (m, 2H), 3.28 (q, J = 5.7 Hz, 2H), 1.90 – 1.78 (m, 2H).

**$^{13}$C-NMR** (101 MHz, DMSO) δ 158.95, 158.77, 155.03, 154.64, 144.05, 141.73, 135.38, 130.14, 128.88, 121.49, 120.89, 119.75, 117.26, 116.46, 113.14, 110.18, 55.98, 28.81.

**MS (ESI+)** *m/z* $[M+H]^+$: calc. 446.10, found 446.05

**HRMS** calculated *m/z* for $[C_{20}H_{20}ClN_5O_3S+H]^+$: 446.10481, found: 446.10457

**HPLC**: $t_R$ = 6.425 min (M1), purity ≧ 95% (UV: 254/280 nm)

*1⁵-(4-Fluoro-3-isopropoxyphenyl)-4-thia-2,5,9-triaza-1(2,4)-pyrimidina-3(1,3)-benzenacyclo-nonaphane 4,4-dioxide* (**43**)

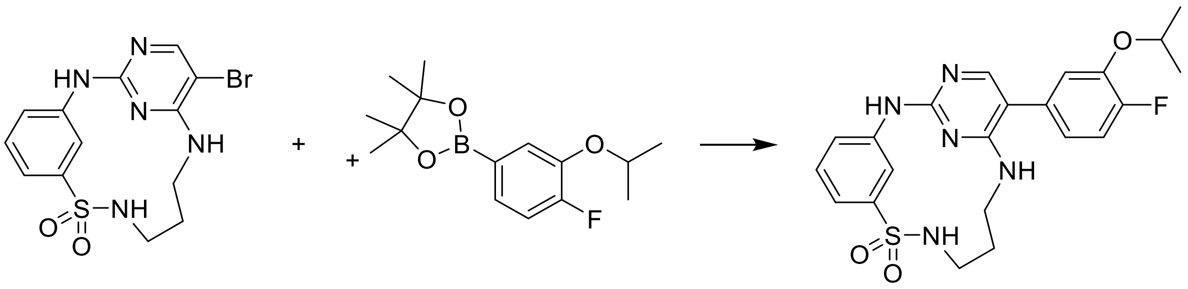

**43** was prepared according to the method for **7**, using 4-fluoro-3-isopropoxyphenylboronic acid. The product was obtained as a white solid (70 mg, 61%).

**$^1$H-NMR** (400 MHz, DMSO-d$_6$) δ 9.61 (s, 1H), 9.45 (t, J = 2.0 Hz, 1H), 7.80 (s, 1H), 7.75 (t, J = 6.0 Hz, 1H), 7.41 – 7.36 (m, 1H), 7.33 – 7.22 (m, 3H), 7.10 (dd, J = 8.3, 2.1 Hz, 1H), 6.96 (t, J = 5.4 Hz, 1H), 6.91 (ddd, J = 8.3, 4.3, 2.1 Hz, 1H), 4.69 (hept, J = 6.0 Hz, 1H), 3.39 (dt, J = 11.6, 5.5 Hz, 2H), 3.28 (q, J = 5.7 Hz, 2H), 1.91 – 1.77 (m, 2H), 1.30 (d, J = 6.0 Hz, 6H).

**$^{13}$C-NMR** (101 MHz, DMSO) δ 159.12, 158.66, 154.95, 151.84 (d, J = 243.4 Hz), 145.49 (d, J = 10.7 Hz), 144.04, 141.79, 131.78 (d, J = 3.6 Hz), 128.86, 121.26 (d, J = 6.9 Hz), 120.85, 117.19, 117.15 (d, J = 2.1 Hz), 116.61, 116.45, 116.43 (d, J = 2.3 Hz), 110.40, 71.17, 28.83, 21.87.

**MS (ESI+)** *m/z* [M+H]$^+$: calc. 458.16, found 458.15

**HRMS** calculated *m/z* for [$C_{22}H_{24}FN_5O_3S$+H]$^+$: 458.16567, found: 458.16536

**HPLC**: $t_R$ = 6.274 min (M1), purity ≧ 95% (UV: 254/280 nm)

*1⁵-(3,4-Dimethoxyphenyl)-4-thia-2,5,9-triaza-1(2,4)-pyrimidina-3(1,3)-benzenacyclonona-phane 4,4-dioxide* (**44**)

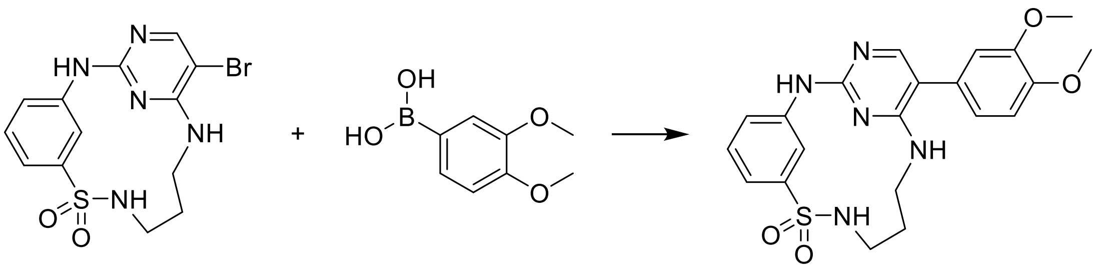

**44** was prepared according to the method for **7**, using 3,4-dimethoxyphenylboronic acid. The product was obtained as a white solid (66 mg, 54%).

**$^1$H-NMR** (400 MHz, DMSO-d$_6$) δ 9.65 (s, 1H), 9.43 (t, J = 2.0 Hz, 1H), 7.80 (s, 1H), 7.76 (t, J = 6.0 Hz, 1H), 7.42 – 7.37 (m, 1H), 7.33 – 7.28 (m, 2H), 7.04 (d, J = 8.2 Hz, 2H), 6.95 – 6.87 (m, 2H), 3.79 (d, J = 4.5 Hz, 6H), 3.41 (dt, J = 10.8, 5.5 Hz, 2H), 3.28 (q, J = 5.8 Hz, 2H), 1.90 – 1.79 (m, 2H).

**$^{13}$C-NMR** (101 MHz, DMSO) δ 159.27, 157.84, 153.53, 148.98, 148.19, 144.02, 141.61, 128.91, 127.13, 120.96, 120.78, 117.36, 116.60, 112.39, 112.35, 111.16, 55.61, 55.39, 28.79.

**MS (ESI+)** *m/z* $[M+H]^+$: calc. 442.15, found 442.05

**HRMS** calculated *m/z* for $[C_{21}H_{23}N_5O_4S+H]^+$: 442.15435, found: 442.15434

**HPLC**: $t_R$ = 5.275 min (M1), purity ≧ 95% (UV: 254/280 nm)

*$1^5$-(3-Chloro-2-fluorophenyl)-4-thia-2,5,9-triaza-1(2,4)-pyrimidina-3(1,3)-benzenacyclononaphane 4,4-dioxide* (**45**)

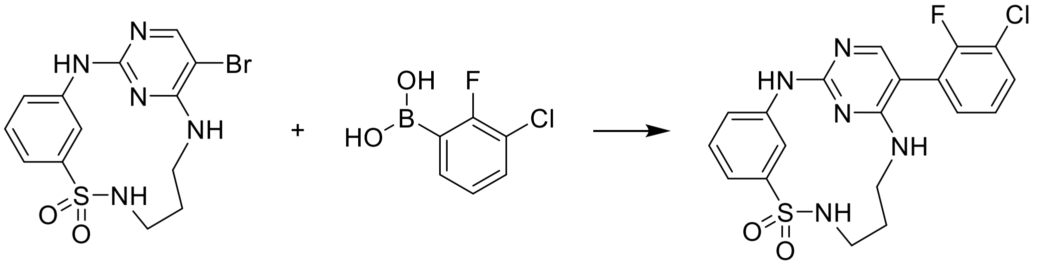

**45** was prepared according to the method for **7**, using 3-chloro-2-fluorophenylboronic acid. The product was obtained as a white solid (51 mg, 45%).

**$^1$H-NMR** (400 MHz, DMSO-d$_6$) δ 9.75 (s, 1H), 9.40 (d, J = 2.1 Hz, 1H), 7.84 – 7.74 (m, 2H), 7.60 (td, J = 7.4, 2.0 Hz, 1H), 7.46 – 7.26 (m, 5H), 7.15 (t, J = 5.5 Hz, 1H), 3.42 – 3.34 (m, 2H), 3.28 (q, J = 5.7 Hz, 2H), 1.82 (d, J = 7.4 Hz, 2H).

**$^{13}$C-NMR** (101 MHz, DMSO) δ 159.10, 158.71, 155.18 (d, J = 246.6 Hz), 155.00, 143.97, 141.36, 130.91 (d, J = 2.9 Hz), 130.07, 128.94, 125.69 (d, J = 4.6 Hz), 124.17 (d, J = 16.0 Hz), 121.24, 120.35 (d, J = 18.2 Hz), 117.71, 116.95, 104.37, 28.68.

**MS (ESI+)** *m/z* $[M+H]^+$: calc. 434.08, found 434.05

**HRMS** calculated *m/z* for $[C_{19}H_{18}ClFN_5O_2S+H]^+$: 434.08483, found: 434.08477

**HPLC**: $t_R$ = 3.585 min (M2), purity ≧ 95% (UV: 254/280 nm)

*$1^5$-(3-Methoxy-5-(trifluoromethyl)phenyl)-4-thia-2,5,9-triaza-1(2,4)-pyrimidina-3(1,3)benzenacyclononaphane 4,4-dioxide* (**46**)

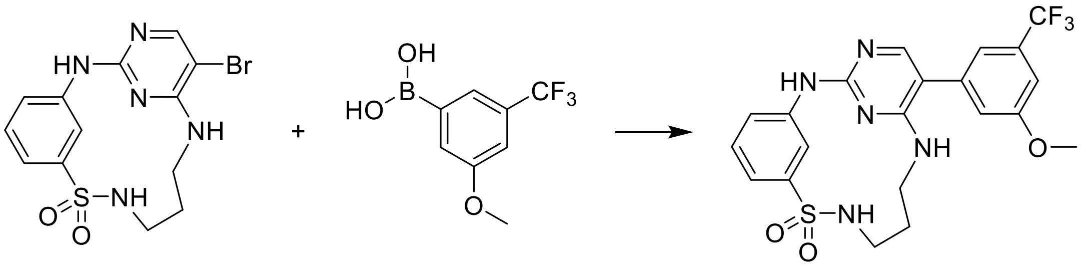

**46** was prepared according to the method for **7**, using 3-methoxy-5-(trifluoromethyl)phenylboronic acid. The product was obtained as a white solid (102 mg, 82%).

**$^1$H-NMR** (400 MHz, DMSO-d$_6$) δ 9.69 (s, 1H), 9.45 (t, J = 2.0 Hz, 1H), 7.88 (s, 1H), 7.77 (t, J = 6.0 Hz, 1H), 7.42 – 7.37 (m, 1H), 7.34 – 7.28 (m, 2H), 7.26 – 7.19 (m, 3H), 7.14 (t, J = 5.4

Hz, 1H), 3.88 (s, 3H), 3.40 (dt, J = 11.3, 5.4 Hz, 2H), 3.29 (q, J = 6.9, 5.6 Hz, 2H), 1.91 – 1.80 (m, 2H).

**$^{13}$C-NMR** (101 MHz, DMSO) δ 159.93, 158.93, 155.46, 144.05, 141.66, 137.84, 130.91 (q, J = 31.7 Hz), 128.90, 123.97 (d, J = 272.8 Hz), 120.95, 118.24, 117.44 – 117.17 (m), 116.52, 109.61, 109.44 (d, J = 3.5 Hz), 55.66, 28.78.

**MS (ESI+)** *m/z* $[M+H]^+$: calc. 480.12, found 480.15

**HRMS** calculated *m/z* for $[C_{21}H_{20}F_3N_5O_3S+H]^+$: 480.13117, found: 480.13116

**HPLC**: $t_R$ = 6.419 min (M1), purity ≧ 95% (UV: 254/280 nm)

*1$^5$-(4-Fluoro-3-(trifluoromethyl)phenyl)-4-thia-2,5,9-triaza-1(2,4)-pyrimidina-3(1,3)benzeneacyclononaphane 4,4-dioxide* (**47**)

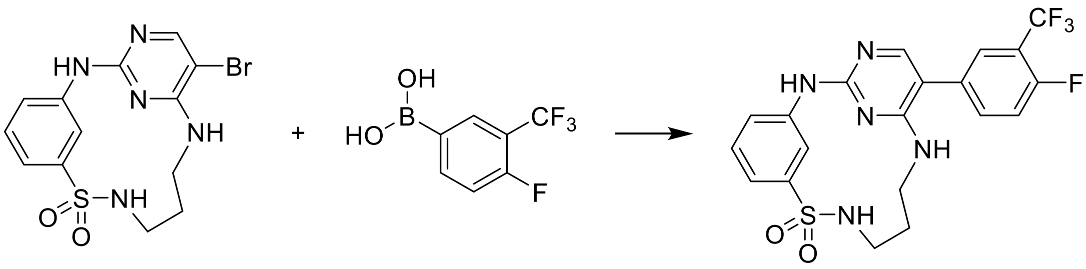

**47** was prepared according to the method for **7**, using 4-fluoro-3-(trifluoromethyl)-phenylboronic acid. The product was obtained as a white solid (82 mg, 67%).

**$^1$H-NMR** (400 MHz, DMSO-$d_6$) δ 9.68 (s, 1H), 9.44 (t, J = 2.0 Hz, 1H), 7.84 (s, 1H), 7.77 (t, J = 6.1 Hz, 1H), 7.75 – 7.69 (m, 2H), 7.58 (dd, J = 10.8, 8.4 Hz, 1H), 7.42 – 7.37 (m, 1H), 7.33 – 7.28 (m, 2H), 7.13 (t, J = 5.4 Hz, 1H), 3.39 (p, J = 5.3 Hz, 2H), 3.29 (q, J = 5.8 Hz, 2H), 1.89 – 1.80 (m, 2H).

**$^{13}$C-NMR** (101 MHz, DMSO) δ 159.04 (d, J = 11.2 Hz), 156.84 (d, J = 2.4 Hz), 155.54, 144.04, 141.66, 135.74 (d, J = 8.9 Hz), 132.23 (d, J = 3.6 Hz), 128.90, 127.56 (q, J = 4.6, 3.6 Hz), 122.62 (d, J = 271.9 Hz), 120.96, 117.66 (d, J = 20.3 Hz), 117.36, 117.30 – 116.78 (m), 116.56, 108.87, 28.78, .

**MS (ESI+)** *m/z* $[M+H]^+$: calc. 468.10, found 468.10

**HRMS** calculated *m/z* for $[C_{20}H_{17}F_4N_5O_2S+H]^+$: 468.11118, found: 468.11128

**HPLC**: $t_R$ = 6.222 min (M1), purity ≧ 95% (UV: 254/280 nm)

*1$^{5}$-(3-Chloro-4-fluorophenyl)-4-thia-2,5,9-triaza-1(2,4)-pyrimidina-3(1,3)benzenacyclononaphane 4,4-dioxide* (**48**)

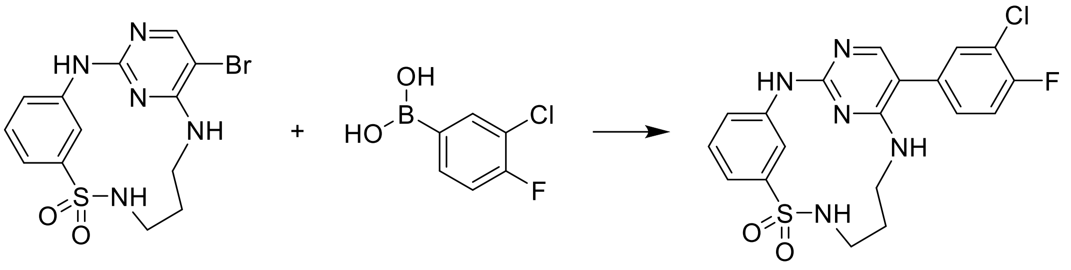

**48** was prepared according to the method for **7**, using 3-chloro-4-fluorophenylboronic acid. The product was obtained as a white solid (43 mg, 35%).

**$^{1}$H-NMR** (400 MHz, DMSO-d$_6$) δ 9.65 (s, 1H), 9.44 (t, J = 2.1 Hz, 1H), 7.81 (s, 1H), 7.77 (t, J = 5.9 Hz, 1H), 7.57 (dd, J = 7.2, 2.2 Hz, 1H), 7.48 (t, J = 9.0 Hz, 1H), 7.42 – 7.35 (m, 2H), 7.33 – 7.28 (m, 2H), 7.07 (t, J = 5.4 Hz, 1H), 3.38 (p, J = 6.7 Hz, 2H), 3.31 – 3.25 (m, 2H), 1.90 – 1.78 (m, 2H).

**$^{13}$C-NMR** (101 MHz, DMSO) δ 159.44 (d, J = 17.5 Hz), 155.76, 143.34 (d, J = 237.2 Hz), 133.47 (d, J = 3.8 Hz), 131.33, 130.14 (d, J = 7.4 Hz), 129.38, 121.43, 120.33 (d, J = 17.9 Hz), 117.83, 117.63, 117.04, 109.52, 29.25.

**MS (ESI+)** *m/z* $[M+H]^+$: calc. 434.08, found 434.05

**HRMS** calculated *m/z* for $[C_{19}H_{17}ClFN_5O_2S+H]^+$: 434.08492, found: 434.08404

**HPLC**: $t_R$ = 6.014 min (M1), purity ≧ 95% (UV: 254/280 nm)

*1$^{5}$-(4-Fluoro-3-hydroxyphenyl)-4-thia-2,5,9-triaza-1(2,4)-pyrimidina-3(1,3)benzenacyclononaphane 4,4-dioxide* (**49**)

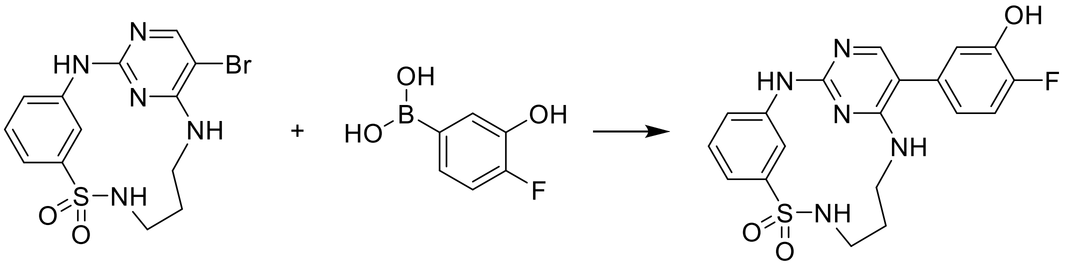

**49** was prepared according to the method for **7**, using 4-fluoro-3-hydroxyphenylboronic acid. The product was obtained as a white solid (51 mg, 46%).

**$^{1}$H-NMR** (400 MHz, DMSO-d$_6$) δ 9.97 (s, 1H), 9.58 (s, 1H), 9.43 (t, J = 2.0 Hz, 1H), 7.75 (d, J = 5.6 Hz, 2H), 7.41 – 7.36 (m, 1H), 7.29 (ddt, J = 7.6, 4.6, 1.3 Hz, 2H), 7.19 (dd, J = 11.4, 8.3 Hz, 1H), 6.96 – 6.90 (m, 2H), 6.76 (ddd, J = 8.3, 4.3, 2.2 Hz, 1H), 3.43 – 3.35 (m, 3H), 3.27 (q, J = 5.7 Hz, 2H), 1.89 – 1.78 (m, 2H).

**$^{13}$C-NMR** (101 MHz, DMSO) δ 159.13, 158.59, 154.58, 150.45 (d, J = 241.0 Hz), 145.12 (d, J = 12.4 Hz), 144.02, 141.77, 131.54 (d, J = 3.5 Hz), 128.87, 120.89, 119.70 (d, J = 6.4 Hz), 118.01 (d, J = 2.9 Hz), 117.22, 116.54 (d, J = 18.2 Hz), 116.52, 110.48, 28.78.

**MS (ESI+)** *m/z* $[M+H]^+$: calc. 416.11, found 416.10

**HRMS** calculated *m/z* for $[C_{19}H_{18}FN_5O_3S+H]^+$: 416.11872, found: 416.11824

**HPLC**: $t_R$ = 5.098 min (M1), purity ≧ 95% (UV: 254/280 nm)

*Methyl 5-(4,4-dioxido-4-thia-2,5,9-triaza-1(2,4)-pyrimidina-3(1,3)-benzenacyclononaphane-1⁵-yl)-2-fluorobenzoate* **(50)**

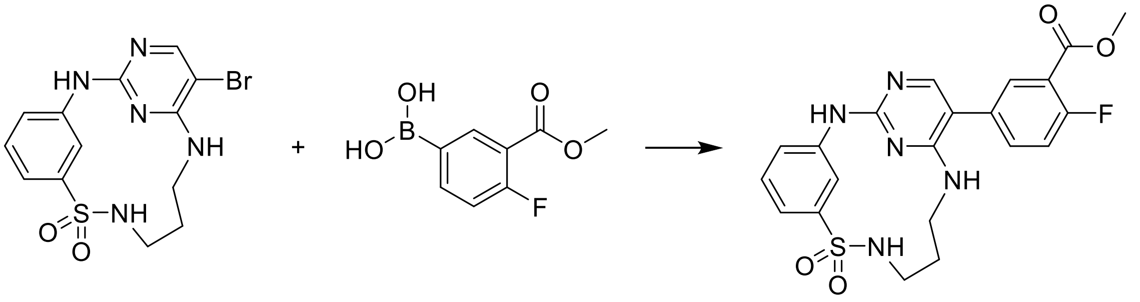

**50** was prepared according to the method for **7**, using 4-fluoro-3-methoxycarbonylphenylboronic acid. The product was obtained as a white solid (16 mg, 14%).

**¹H-NMR** (400 MHz, DMSO-$d_6$) δ 9.66 (s, 1H), 9.44 (t, J = 2.0 Hz, 1H), 7.85 – 7.80 (m, 2H), 7.77 (t, J = 6.0 Hz, 1H), 7.65 (ddd, J = 8.6, 4.7, 2.5 Hz, 1H), 7.46 – 7.36 (m, 2H), 7.33 – 7.28 (m, 2H), 7.07 (t, J = 5.4 Hz, 1H), 3.87 (s, 3H), 3.43 – 3.37 (m, 2H), 3.28 (q, J = 5.7 Hz, 2H), 1.90 – 1.79 (m, 2H).

**¹³C-NMR** (101 MHz, DMSO) δ 163.96 (d, J = 3.6 Hz), 161.44, 159.01 (d, J = 31.0 Hz), 158.88, 155.12, 144.04, 141.67, 135.66 (d, J = 9.2 Hz), 132.01, 131.59 (d, J = 3.7 Hz), 128.93, 120.99, 118.56 (d, J = 10.6 Hz), 117.72 (d, J = 22.5 Hz), 117.38, 116.60, 109.21, 52.47, 28.79.

**MS (ESI+)** *m/z* $[M+H]^+$: calc. 458.12, found 458.15

**HRMS** calculated *m/z* for $[C_{21}H_{20}FN_5O_4S+H]^+$: 458.12928, found: 458.12892

**HPLC**: $t_R$ = 5.165 min (M1), purity ≧ 95% (UV: 254/280 nm)

*1⁵-(3-Amino-4-fluorophenyl)-4-thia-2,5,9-triaza-1(2,4)-pyrimidina-3(1,3)benzenacyclono-naphane 4,4-dioxide* (**51**)

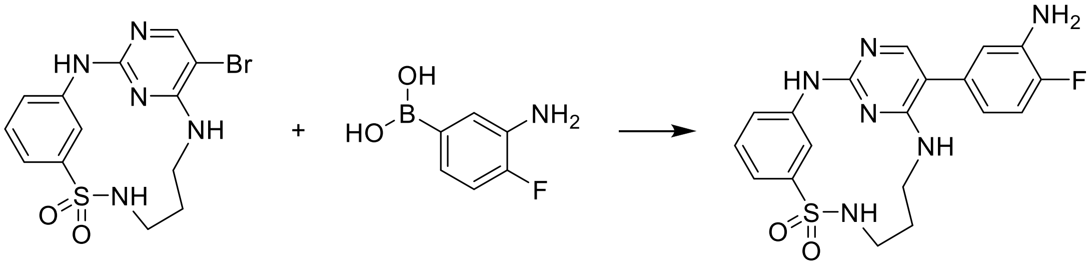

**51** was prepared according to the method for **7**, using 3-amino-4-fluorophenylboronic acid. The product was obtained as a white solid (28 mg, 26%).

**$^{1}$H-NMR** (400 MHz, DMSO-$d_6$) δ 9.55 (s, 1H), 9.44 (t, J = 2.0 Hz, 1H), 7.75 (t, J = 6.0 Hz, 1H), 7.72 (s, 1H), 7.38 (t, J = 7.8 Hz, 1H), 7.31 – 7.25 (m, 2H), 7.04 (dd, J = 11.6, 8.2 Hz, 1H), 6.81 (t, J = 5.5 Hz, 1H), 6.75 (dd, J = 8.8, 2.2 Hz, 1H), 6.48 (ddd, J = 8.2, 4.4, 2.2 Hz, 1H), 5.21 (s, 2H), 3.43 – 3.36 (m, 2H), 3.30 – 3.24 (m, 2H), 1.88 – 1.78 (m, 2H).

**$^{13}$C-NMR** (101 MHz, DMSO) δ 159.16, 158.52, 154.42, 150.09 (d, J = 237.3 Hz), 144.02, 141.85, 136.76 (d, J = 13.2 Hz), 131.29 (d, J = 3.1 Hz), 128.87, 120.82, 117.13, 116.44, 116.35 (d, J = 4.9 Hz), 116.22 (d, J = 6.8 Hz), 115.31 (d, J = 18.5 Hz), 111.03, 28.79.

**MS (ESI+)** *m/z* $[M+H]^+$: calc. 415.13, found 415.10

**HRMS** calculated *m/z* for $[C_{19}H_{19}FN_6O_2S+H]^+$: 415.13470, found: 415.13448

**HPLC**: $t_R$ = 3.193 min (M2), purity ≧ 95% (UV: 254/280 nm)

*4-(4,4-Dioxido-4-thia-2,5,9-triaza-1(2,4)-pyrimidina-3(1,3)-benzenacyclononaphane-1$^5$-yl)-2-methoxybenzonitrile* (**52**)

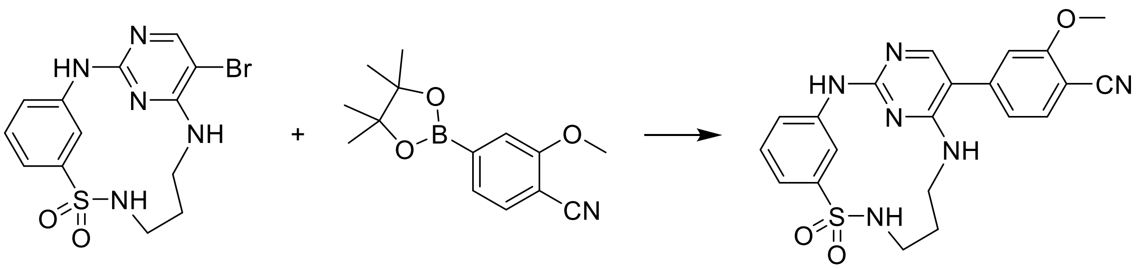

**52** was prepared according to the method for **6**, using **37**. The product was obtained as a white solid (33 mg, 30%).

**$^{1}$H-NMR** (400 MHz, DMSO-$d_6$) δ 9.73 (s, 1H), 9.44 (t, J = 2.0 Hz, 1H), 7.95 (s, 1H), 7.80 – 7.74 (m, 2H), 7.40 (t, J = 7.7 Hz, 1H), 7.34 – 7.29 (m, 2H), 7.24 – 7.19 (m, 2H), 7.12 (dd, J = 8.0, 1.4 Hz, 1H), 3.97 (s, 3H), 3.40 (q, J = 5.7 Hz, 2H), 3.28 (q, J = 4.9 Hz, 2H), 1.93 – 1.79 (m, 2H).

**$^{13}$C-NMR** (101 MHz, DMSO) δ 161.13, 159.02, 158.72, 155.64, 144.06, 142.34, 141.55, 133.95, 128.94, 121.03, 117.49, 116.57, 112.14, 109.67, 98.44, 56.24, 28.77.

**MS (ESI+)** *m/z* $[M+H]^+$: calc. 437.13, found 437.10

**HRMS** calculated *m/z* for $[C_{19}H_{19}N_6O_2S+H]^+$: 437.13904, found: 437.13870

**HPLC**: $t_R$ = 3.381 min (M2), purity ≧ 95% (UV: 254/280 nm)

*1^5-(4-Fluoro-3-(methylthio)phenyl)-4-thia-2,5,9-triaza-1(2,4)-pyrimidina-3(1,3)-benzena-cyclononaphane 4,4-dioxide* (**53**)

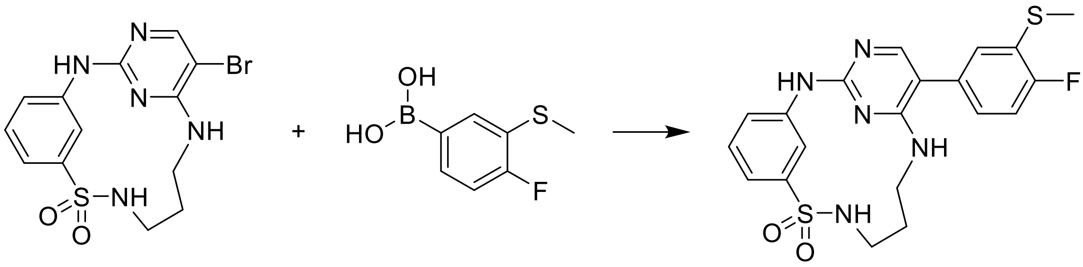

**53** was prepared according to the method for **7**, using 4-fluoro-3-(methylthio)phenylboronic acid. The product was obtained as a white solid (36 mg, 69%).

**$^1$H-NMR** (400 MHz, DMSO-d$_6$) δ 10.35 (s, 1H), 9.25 (t, J = 2.0 Hz, 1H), 7.88 (s, 1H), 7.83 – 7.73 (m, 2H), 7.48 (t, J = 7.8 Hz, 1H), 7.42 (dt, J = 7.8, 1.4 Hz, 1H), 7.37 – 7.28 (m, 3H), 7.21 (ddd, J = 8.4, 5.0, 2.2 Hz, 1H), 3.45 – 3.36 (m, 2H), 3.29 (q, J = 5.7 Hz, 2H), 2.52 (s, 3H), 1.88 – 1.78 (m, 2H).

**$^{13}$C-NMR** (101 MHz, DMSO) δ 159.74 – 159.65 (m), 157.27, 154.83 – 154.62 (m), 143.89, 140.00, 130.24, 129.23, 127.52, 127.08 (d, J = 8.0 Hz), 126.27 (d, J = 17.4 Hz), 122.04, 119.06, 117.94, 115.72 – 115.33 (m), 110.87, 28.43, 13.60.

**MS (ESI+)** *m/z* $[M+H]^+$: calc. 446.10, found 446.10

**HRMS** calculated *m/z* for $[C_{20}H_{20}FN_5O_2S+H]^+$: 446.11152, found: 446.11085

**HPLC**: $t_R$ = 3.937 min (M2), purity ≧ 95% (UV: 254/280 nm)

*1^5-(3,4-Dichlorophenyl)-4-thia-2,5,9-triaza-1(2,4)-pyrimidina-3(1,3)-benzenacyclononaphane 4,4-dioxide* (**54**)

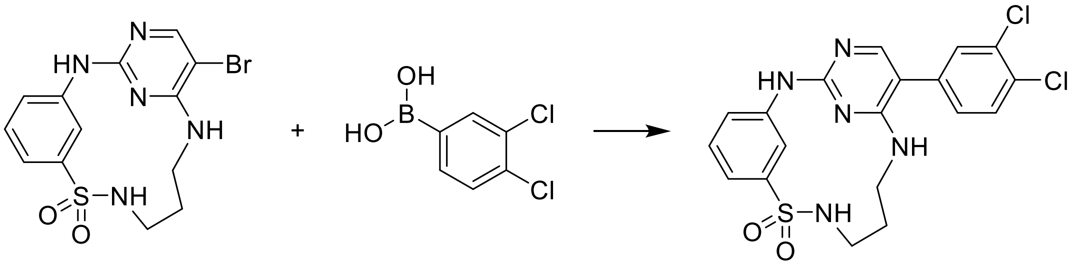

**54** was prepared according to the method for **7**, using 3,4-dichlorophenylboronic acid. The product was obtained as a white solid (36 mg, 34%).

**$^1$H-NMR** (400 MHz, DMSO-d$_6$) δ 10.41 (s, 1H), 9.22 (t, J = 2.0 Hz, 1H), 7.90 (s, 1H), 7.89 – 7.85 (m, 1H), 7.81 (t, J = 5.9 Hz, 1H), 7.74 (d, J = 8.3 Hz, 1H), 7.67 (d, J = 2.0 Hz, 1H), 7.51 – 7.32 (m, 4H), 3.46 – 3.34 (m, 2H), 3.29 (q, J = 5.7 Hz, 2H), 1.90 – 1.76 (m, 2H).

**$^{13}$C-NMR** (101 MHz, DMSO) δ 159.52, 154.73, 148.15, 143.87, 139.82, 133.94, 131.62, 131.10, 130.69, 129.53, 129.26, 122.18, 119.24, 118.08, 109.77, 28.36.

**MS (ESI+)** *m/z* $[M+H]^+$: calc. 450.05, found 450.05

**HRMS** calculated *m/z* for $[C_{19}H_{17}Cl_2N_5O_2S+H]^+$: 450.0553, found: 450.0566

**HPLC**: $t_R$ = 3.485 min (M2), purity ≧ 95% (UV: 254/280 nm)

*$1^5$-(4-Bromo-3-methoxyphenyl)-4-thia-2,5,9-triaza-1(2,4)-pyrimidina-3(1,3)-benzenacyclo-nonaphane 4,4-dioxide* (**55**)

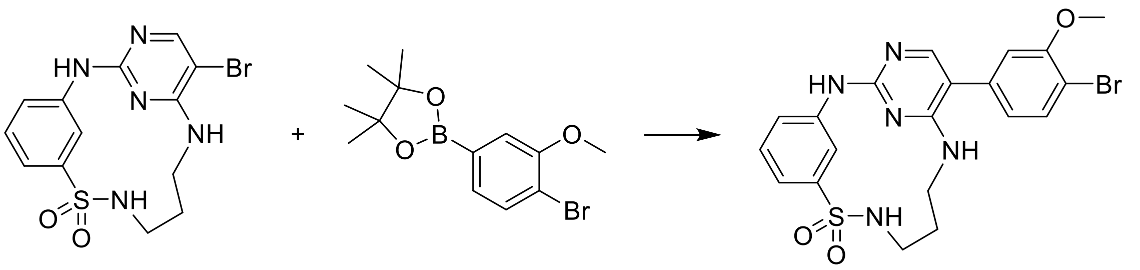

**55** was prepared according to the method for **7**, using 4-bromo-3-methoxyphenylboronic acid. The product was obtained as a white solid (11 mg, 11%).

**$^1$H-NMR** (400 MHz, DMSO-d$_6$) δ 10.59 (s, 1H), 9.20 (s, 1H), 8.01 (s, 1H), 7.92 (s, 1H), 7.83 (d, J = 6.0 Hz, 1H), 7.68 (d, J = 8.1 Hz, 1H), 7.54 – 7.42 (m, 2H), 7.37 (d, J = 7.7 Hz, 1H), 7.12 (s, 1H), 6.92 (d, J = 8.1 Hz, 1H), 3.90 (s, 3H), 3.45 – 3.36 (m, 2H), 3.32 – 3.24 (m, 2H), 1.93 – 1.70 (m, 2H).

**$^{13}$C-NMR** (101 MHz, DMSO) δ 159.63, 155.64, 153.60, 145.85, 143.88, 139.43, 133.49, 133.44, 129.34, 122.40, 122.32, 119.57, 118.27, 113.44, 111.30, 110.56, 56.22, 28.28.

**MS (ESI+)** *m/z* $[M+H]^+$: calc. 490.05, found 490.00

**HRMS** calculated *m/z* for $[C_{20}H_{20}BrN_5O_3S+H]^+$: 490.0543, found: 490.0557

**HPLC**: $t_R$ = 3.765 min (M2), purity ≧ 95% (UV: 254/280 nm)

*$1^5$-(2,4-Difluoro-5-methoxyphenyl)-4-thia-2,5,9-triaza-1(2,4)-pyrimidina-3(1,3)-benzena-cyclononaphane 4,4-dioxide* (**56**)

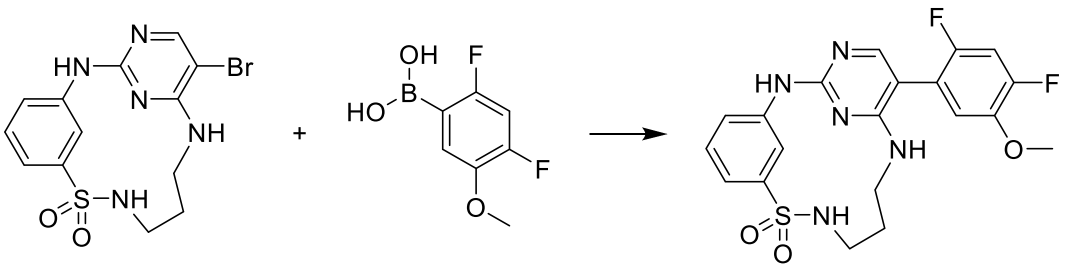

**56** was prepared according to the method for **7**, using 2,4-difluoro-5-methoxyphenylboronic acid. The product was obtained as a white solid (75 mg, 66%).

**$^1$H-NMR** (400 MHz, DMSO-d$_6$) δ 9.66 (s, 1H), 9.45 (t, J = 2.0 Hz, 1H), 7.80 (s, 1H), 7.76 (t, J = 6.0 Hz, 1H), 7.42 – 7.34 (m, 2H), 7.33 – 7.28 (m, 2H), 7.13 (dd, J = 9.6, 7.1 Hz, 1H), 6.94 (t, J = 5.4 Hz, 1H), 3.86 (s, 3H), 3.42 – 3.35 (m, 2H), 3.28 (q, J = 5.9 Hz, 2H), 1.87 – 1.78 (m, 2H).

**$^{13}$C-NMR** (101 MHz, DMSO) δ 159.22, 159.14, 155.85, 154.18 – 151.87 (m), 150.60 (dd, J = 218.3, 11.3 Hz), 144.06 – 143.86 (m), 141.69, 128.89, 120.96, 118.00 (dd, J = 17.5, 4.0 Hz), 117.35, 116.63, 116.08 (t, J = 3.5 Hz), 105.14 (dd, J = 28.4, 22.2 Hz), 104.41, 56.47, 28.76.

**MS (ESI+)** *m/z* $[M+H]^+$: calc. 448.12, found 448.10

**HRMS** calculated *m/z* for $[C_{20}H_{19}F_2N_5O_3S+H]^+$: 448.12494, found: 448.12458

**HPLC**: $t_R$ = 5.652 min (M1), purity ≧ 95% (UV: 254/280 nm)

*1<sup></sup>*

<u>*$1^5$-(3,4,5-Trimethoxyphenyl)-4-thia-2,5,9-triaza-1(2,4)-pyrimidina-3(1,3)benzenacyclononaphane 4,4-dioxide*</u> (**57**)

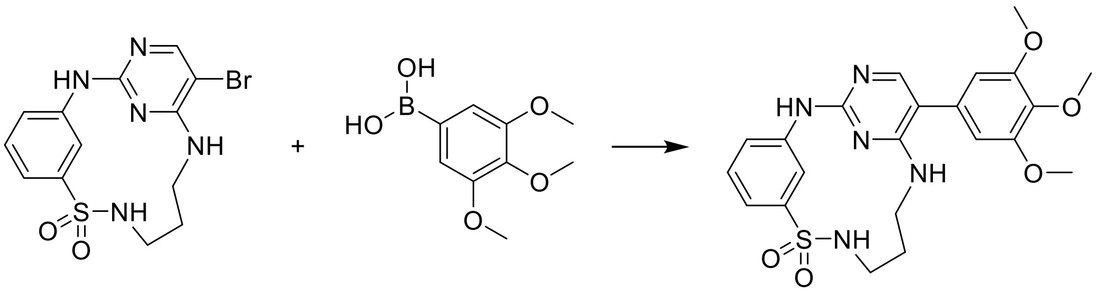

**57** was prepared according to the method for **7**, using 3,4,5-trimethoxyphenylboronic acid. The product was obtained as a white solid (43 mg, 35%).

**$^{1}$H-NMR** (400 MHz, DMSO-$d_6$) δ 9.59 (s, 1H), 9.47 (t, J = 2.0 Hz, 1H), 7.85 (s, 1H), 7.76 (t, J = 6.0 Hz, 1H), 7.39 (t, J = 7.9 Hz, 1H), 7.32 – 7.27 (m, 2H), 7.02 (t, J = 5.5 Hz, 1H), 6.64 (s, 2H), 3.82 (s, 6H), 3.69 (s, 3H), 3.42 (dt, J = 11.6, 5.5 Hz, 2H), 3.28 (q, J = 5.9 Hz, 2H), 1.90 – 1.80 (m, 2H).

**$^{13}$C-NMR** (101 MHz, DMSO) δ 158.96, 158.54, 154.79, 153.17, 144.06, 141.84, 136.58, 130.46, 128.87, 120.79, 117.13, 116.35, 111.11, 105.82, 59.92, 55.80, 28.86.

**MS (ESI+)** *m/z* $[M+H]^+$: calc. 472.16, found 472.15

**HRMS** calculated *m/z* for $[C_{22}H_{25}N_5O_5S+H]^+$: 472.16492, found: 472.16489

**HPLC**: $t_R$ = 5.428 min (M1), purity ≧ 95% (UV: 254/280 nm)

<u>*$1^5$-(2,4-Difluoro-3-methoxyphenyl)-4-thia-2,5,9-triaza-1(2,4)-pyrimidina-3(1,3)-benzenacyclononaphane 4,4-dioxide*</u> (**58**)

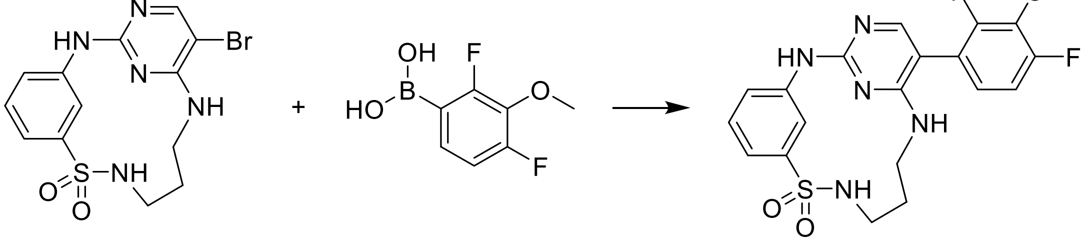

**58** was prepared according to the method for **7**, using 2,4-difluoro-3-methoxyphenylboronic acid. The product was obtained as a white solid (78 mg, 63%).

**$^1$H-NMR** (400 MHz, DMSO-$d_6$) δ 9.65 (s, 1H), 9.43 (t, J = 2.1 Hz, 1H), 7.80 – 7.74 (m, 2H), 7.39 (t, J = 7.9 Hz, 1H), 7.34 – 7.28 (m, 2H), 7.20 (ddd, J = 10.6, 8.7, 1.7 Hz, 1H), 7.10 – 7.03 (m, 1H), 6.98 (t, J = 5.5 Hz, 1H), 3.96 (s, 3H), 3.43 – 3.33 (m, 2H), 3.28 (q, J = 5.7 Hz, 2H), 1.90 – 1.76 (m, 2H).

**$^{13}$C-NMR** (101 MHz, DMSO) δ 159.24, 156.10 (d, J = 5.5 Hz), 155.80, 154.58 (d, J = 6.1 Hz), 153.66 (d, J = 5.0 Hz), 152.13 (d, J = 5.7 Hz), 144.00, 141.67, 136.18 (t, J = 14.6 Hz), 128.89, 121.00, 119.95 (dd, J = 14.9, 3.4 Hz), 117.38, 116.68, 112.41 (dd, J = 18.9, 3.5 Hz), 104.25, 61.75 (t, J = 3.0 Hz), 28.74.

**MS (ESI+)** *m/z* $[M+H]^+$: calc. 448.12, found 448.10

**HRMS** calculated *m/z* for $[C_{20}H_{19}F_2N_5O_3S+H]^+$: 448.12494, found: 448.12472

**HPLC**: $t_R$ = 5.374 min (M1), purity ≧ 95% (UV: 254/280 nm)

*1$^5$-(3,4-Difluoro-5-methoxyphenyl)-4-thia-2,5,9-triaza-1(2,4)-pyrimidina-3(1,3)-benzenacyclononaphane 4,4-dioxide* (**59**)

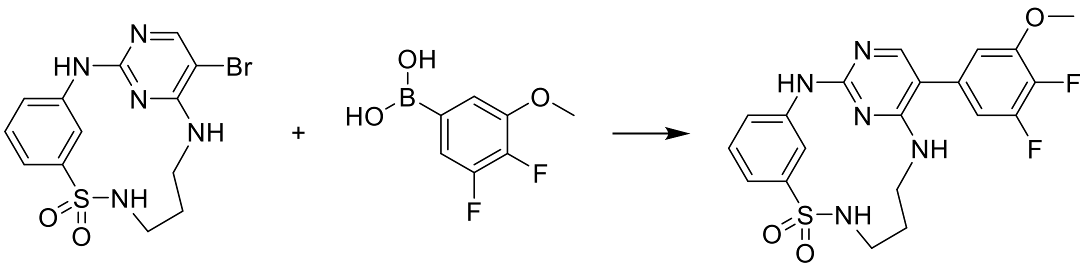

**59** was prepared according to the method for **7**, using 3,4-difluoro-5-methoxyphenylboronic acid. The product was obtained as a white solid (11 mg, 14%).

**$^1$H-NMR** (400 MHz, DMSO-$d_6$) δ 9.66 (s, 1H), 9.45 (t, J = 2.1 Hz, 1H), 7.86 (s, 1H), 7.76 (t, J = 6.0 Hz, 1H), 7.39 (t, J = 7.8 Hz, 1H), 7.34 – 7.27 (m, 2H), 7.10 (t, J = 5.4 Hz, 1H), 7.05 – 6.96 (m, 2H), 3.92 (s, 3H), 3.40 (q, J = 5.7 Hz, 2H), 3.29 (q, J = 5.6 Hz, 2H), 1.92 – 1.79 (m, 2H).

**$^{13}$C-NMR** (101 MHz, DMSO) δ 158.89 (d, J = 5.6 Hz), 155.20, 150.31 (dd, J = 243.6, 10.1 Hz), 148.75 (dd, J = 7.5, 3.9 Hz), 144.06, 141.71, 139.10 (dd, J = 245.1, 14.6 Hz), 131.35 (dd, J = 8.8, 4.6 Hz), 128.91, 120.93, 117.33, 116.50, 110.02, 109.57, 109.09 (d, J = 17.9 Hz), 56.59, 28.81.

**MS (ESI+)** *m/z* $[M+H]^+$: calc. 448.12, found 448.15

**HRMS** calculated *m/z* for $[C_{20}H_{19}F_2N_5O_3S+H]^+$: 448.12494, found: 448.12458

**HPLC**: $t_R$ = 3.369 min (M2), purity ≧ 95% (UV: 254/280 nm)

*1⁵-(5-Chloro-2,4-difluorophenyl)-4-thia-2,5,9-triaza-1(2,4)-pyrimidina-3(1,3)-benzenacyclononaphane 4,4-dioxide* (**60**)

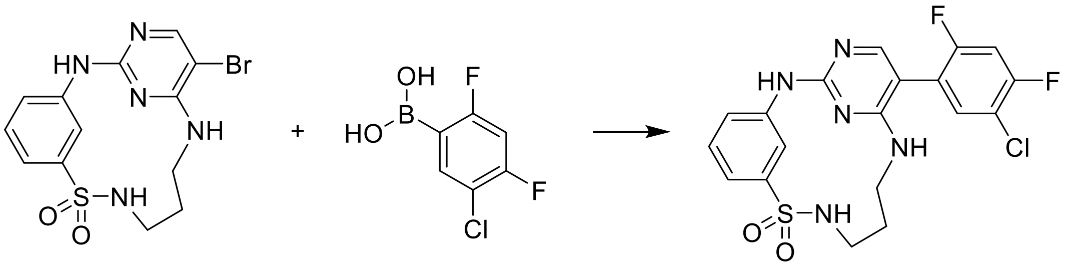

**60** was prepared according to the method for **7**, using 5-chloro-2,4-difluorophenylboronic acid. The product was obtained as a white solid (25 mg, 25%).

**$^1$H-NMR** (400 MHz, DMSO-d$_6$) δ 10.56 (s, 1H), 9.17 (s, 1H), 8.01 (s, 1H), 7.93 (s, 1H), 7.86 – 7.78 (m, 1H), 7.75 – 7.65 (m, 2H), 7.54 – 7.43 (m, 2H), 7.37 (d, J = 7.2 Hz, 1H), 3.43 – 3.33 (m, 2H), 3.32 – 3.24 (m, 2H), 1.88 – 1.72 (m, 2H).

**$^{13}$C-NMR** (101 MHz, DMSO) δ 160.25, 159.88, 159.66, 158.74, 154.27, 143.78, 139.43, 133.20 (d, J = 3.4 Hz), 129.31, 122.53, 119.68, 118.57, 117.95 (d, J = 24.6 Hz), 115.65 (d, J = 20.1 Hz), 106.64 – 105.97 (m), 104.24, 28.29.

**MS (ESI+)** *m/z* [M+H]$^+$: calc. 452.07, found 452.05

**HRMS** calculated *m/z* for [$C_{19}H_{16}ClF_2N_5O_2S$+H]$^+$: 452.07541, found: 452.07521

**HPLC**: $t_R$ = 3.414 min (M2), purity ≧ 95% (UV: 254/280 nm)

*1⁵-(4-Chloro-2-fluoro-5-methoxyphenyl)-4-thia-2,5,9-triaza-1(2,4)-pyrimidina-3(1,3)-benzenacyclononaphane 4,4-dioxide* (**61**)

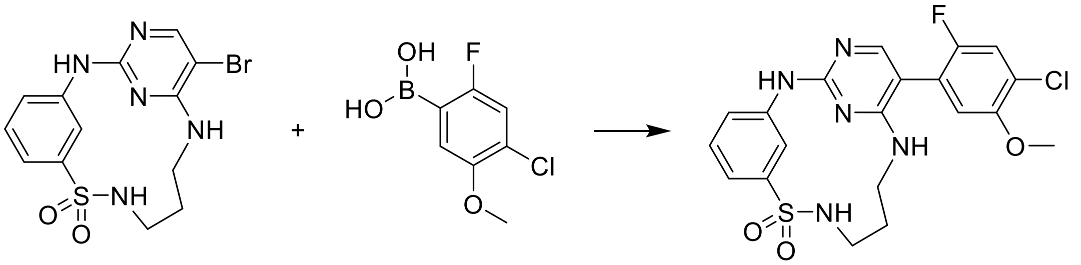

**61** was prepared according to the method for **7**, using 4-chloro-2-fluoro-5-methoxyphenylboronic acid. The product was obtained as a white solid (32 mg, 31%).

**$^1$H-NMR** (400 MHz, DMSO-d$_6$) δ 10.39 (s, 1H), 9.24 (s, 1H), 7.93 (s, 1H), 7.84 – 7.74 (m, 2H), 7.56 (d, J = 9.0 Hz, 1H), 7.50 – 7.40 (m, 2H), 7.35 (d, J = 7.9 Hz, 1H), 7.15 (d, J = 6.5 Hz, 1H), 3.87 (s, 3H), 3.39 (q, J = 6.0, 5.6 Hz, 2H), 3.29 (q, J = 5.6 Hz, 2H), 1.88 – 1.75 (m, 2H).

**$^{13}$C-NMR** (101 MHz, DMSO) δ 159.43, 155.21, 154.54, 152.15, 151.42 (d, J = 2.3 Hz), 143.87, 139.94, 129.24, 122.13, 121.54 (d, J = 10.7 Hz), 119.94 (d, J = 17.3 Hz), 119.15, 118.07, 117.74 (d, J = 27.0 Hz), 115.17 (d, J = 3.1 Hz), 105.25, 56.67, 28.40.

**MS (ESI+)** *m/z* [M+H]$^+$: calc. 464.09, found 464.05

**HRMS** calculated *m/z* for $[C_{20}H_{19}ClFN_5O_3S+H]^+$: 464.09539, found: 464.09443

**HPLC**: $t_R$ = 3.706 min (M2), purity ≧ 95% (UV: 254/280 nm)

*1[5]-(4,5-Dichloro-2-methylphenyl)-4-thia-2,5,9-triaza-1(2,4)-pyrimidina-3(1,3)-benzenacyclo-nonaphane 4,4-dioxide* (**62**)

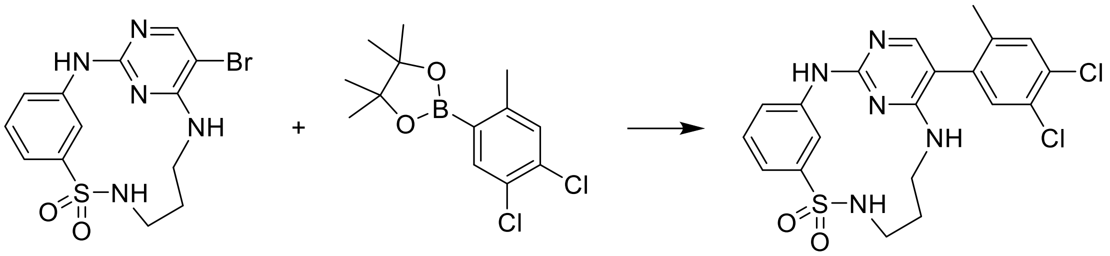

**62** was prepared according to the method for **7**, using **39**. The product was obtained as a white solid (38 mg, 44%).

**$^1$H-NMR** (400 MHz, DMSO-d$_6$) δ 9.63 (s, 1H), 9.45 (t, J = 2.0 Hz, 1H), 7.74 (t, J = 6.0 Hz, 1H), 7.70 (s, 1H), 7.61 (s, 1H), 7.42 (s, 1H), 7.41 – 7.36 (m, 1H), 7.32 – 7.27 (m, 2H), 6.74 (t, J = 5.4 Hz, 1H), 3.45 – 3.35 (m, 2H), 3.31 – 3.19 (m, 2H), 2.09 (s, 3H), 1.91 – 1.70 (m, 2H).

**$^{13}$C-NMR** (101 MHz, DMSO) δ 159.12, 158.86, 155.00, 143.98, 141.76, 138.72, 135.30, 132.27, 131.81, 130.14, 128.86, 128.29, 120.94, 117.28, 116.66, 108.57, 28.81, 18.61.

**MS (ESI+)** *m/z* $[M+H]^+$: calc. 464.06, found 464.05

**HRMS** calculated *m/z* for $[C_{20}H_{19}Cl_2N_5O_3S+H]^+$: 464.07093, found: 464.07026

**HPLC**: $t_R$ = 3.757 min (M2), purity ≧ 95% (UV: 254/280 nm)

*1[5]-(4,5-Dichloro-2-fluorophenyl)-4-thia-2,5,9-triaza-1(2,4)-pyrimidina-3(1,3)-benzenacyclo-nonaphane 4,4-dioxide* (**63**)

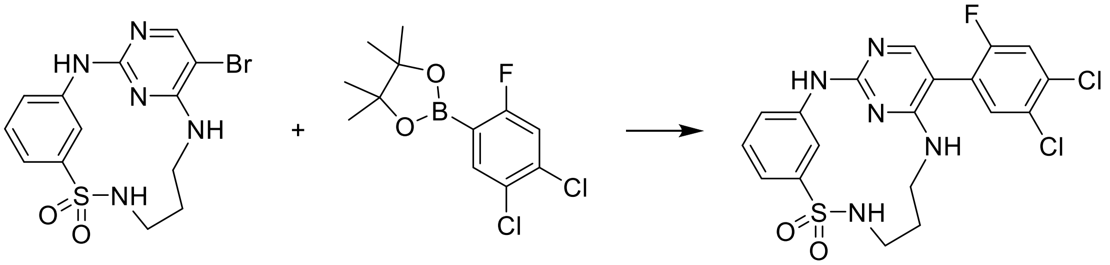

**63** was prepared according to the method for **7**, using **40**. The product was obtained as a white solid (11 mg, 10%).

**$^1$H-NMR** (400 MHz, DMSO-d$_6$) δ 9.73 (s, 1H), 9.41 (t, J = 2.1 Hz, 1H), 7.83 – 7.75 (m, 3H), 7.68 (d, J = 7.2 Hz, 1H), 7.43 – 7.36 (m, 1H), 7.33 – 7.29 (m, 2H), 7.13 (t, J = 5.5 Hz, 1H), 3.42 – 3.36 (m, 2H), 3.31 – 3.25 (m, 2H), 1.87 – 1.76 (m, 2H).

$^{13}$**C-NMR** (101 MHz, DMSO) δ 159.22, 159.08, 158.48 (d, J = 248.5 Hz), 155.77, 143.99, 141.49, 133.00 (d, J = 4.1 Hz), 131.13 (d, J = 11.0 Hz), 128.94, 127.07 (d, J = 3.8 Hz), 123.56 (d, J = 18.0 Hz), 121.14, 118.48 (d, J = 27.7 Hz), 117.59, 116.83, 103.05, 28.72.

**MS (ESI+)** *m/z* $[M+H]^+$: calc. 468.04, found 468.00

**HRMS** calculated *m/z* for $[C_{19}H_{16}Cl_2FN_5O_2S+H]^+$: 468.04586, found: 468.04590

**HPLC**: $t_R$ = 3.858 min (M2), purity = 68% (UV: 254/280 nm)

*N-(3-((5-bromo-2-chloropyrimidin-4-yl)amino)propyl)-2-methyl-5-nitrobenzenesulfonamide* (**64**)

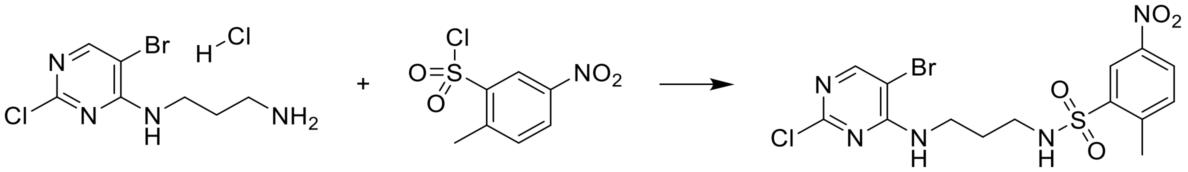

**64** was prepared according to the method for **4**, using 2-methyl-5-nitrobenzenesulfonyl chloride. The product was obtained as a white solid (1.64 g, 83%).

$^{1}$**H-NMR** (400 MHz, DMSO-$d_6$) δ 8.51 (d, J = 2.5 Hz, 1H), 8.30 (dd, J = 8.3, 2.5 Hz, 1H), 8.18 (s, 1H), 8.10 (t, J = 5.8 Hz, 1H), 7.66 (d, J = 8.4 Hz, 1H), 7.58 (t, J = 5.7 Hz, 1H), 3.27 (q, J = 6.5 Hz, 2H), 2.87 (dt, J = 7.7, 6.1 Hz, 2H), 2.69 (s, 3H), 1.70 – 1.58 (m, 2H).

$^{13}$**C-NMR** (101 MHz, DMSO) δ 159.26, 158.08, 156.56, 145.38, 144.54, 139.82, 134.11, 126.72, 123.22, 102.55, 40.15, 38.22, 28.12, 19.92.

**MS (ESI+)** *m/z* $[M+H]^+$: calc. 465.97, found 465.95

**HPLC**: $t_R$ = 7.564 min (M1), purity ≧ 95% (UV: 254/280 nm)

*N-(3-((5-bromo-2-chloropyrimidin-4-yl)amino)propyl)-2-chloro-5-nitrobenzenesulfonamide* (**65**)

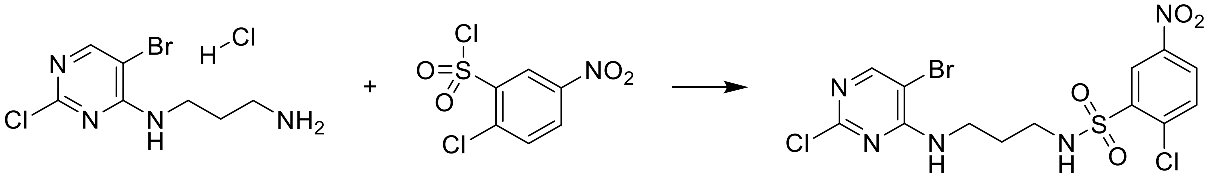

**65** was prepared according to the method for **4**, using 2-chloro-5-nitrobenzenesulfonyl chloride. The product was obtained as a white solid (864 mg, 91%).

$^{1}$**H-NMR** (400 MHz, DMSO-$d_6$) δ 8.60 (d, J = 2.7 Hz, 1H), 8.39 (dd, J = 8.7, 2.8 Hz, 1H), 8.31 (t, J = 5.8 Hz, 1H), 8.18 (s, 1H), 7.93 (d, J = 8.7 Hz, 1H), 7.60 (t, J = 5.7 Hz, 1H), 3.29 (q, J = 6.5 Hz, 2H), 2.95 (q, J = 6.5 Hz, 2H), 1.66 (p, J = 6.7 Hz, 2H).

$^{13}$**C-NMR** (101 MHz, DMSO) δ 159.30, 158.11, 156.58, 145.96, 139.13, 137.36, 133.46, 128.26, 125.14, 102.59, 40.19, 38.20, 28.18.

**MS (ESI+)** *m/z* $[M+H]^+$: calc. 485.91, found 485.85

**HPLC**: $t_R$ = 4.288 min (M2), purity = 94% (UV: 254/280 nm)

*N-(3-((5-bromo-2-chloropyrimidin-4-yl)amino)propyl)-2-methoxy-5-nitrobenzenesulfonamide* (**66**)

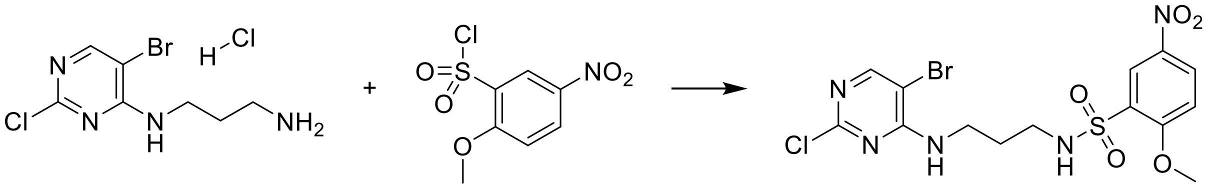

**66** was prepared according to the method for **4**, using 2-methoxy-5-nitrobenzenesulfonyl chloride. The product was obtained as a white solid (4,22 g, 97%).

**$^1$H-NMR** (400 MHz, DMSO-$d_6$) δ 8.46 (d, J = 2.9 Hz, 1H), 8.42 (ddd, J = 9.1, 2.9, 0.7 Hz, 1H), 8.18 (d, J = 0.7 Hz, 1H), 7.72 (t, J = 5.8 Hz, 1H), 7.59 (t, J = 5.7 Hz, 1H), 7.41 (d, J = 9.2 Hz, 1H), 4.05 (s, 3H), 3.27 (q, J = 6.4 Hz, 2H), 2.88 (d, J = 7.8 Hz, 2H), 1.62 (p, J = 6.8 Hz, 2H).

**$^{13}$C-NMR** (101 MHz, DMSO) δ 161.03, 159.26, 158.09, 156.53, 139.67, 129.81, 128.59, 125.07, 113.69, 102.58, 57.42, 40.24, 38.22, 28.08.

**MS (ESI+)** *m/z* $[M+H]^+$: calc. 481.96, found 481.90

**HPLC**: $t_R$ = 4.145 min, purity = 88% (UV: 254/280 nm), FAST_Nonpolar_General

*5-Amino-N-(3-((5-bromo-2-chloropyrimidin-4-yl)amino)propyl)-2-methylbenzenesulfonamide* (**67**)

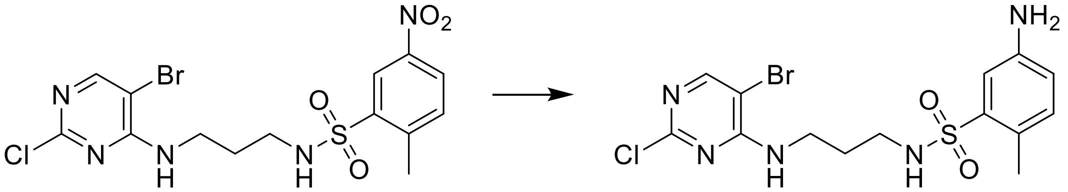

**67** was prepared according to the method for **5**. The product was obtained as a white solid (781 mg, 65%).

**$^1$H-NMR** (400 MHz, DMSO-$d_6$) δ 8.22 (s, 1H), 7.60 (t, J = 5.8 Hz, 1H), 7.41 (t, J = 6.0 Hz, 1H), 7.10 (d, J = 2.5 Hz, 1H), 6.98 (d, J = 8.1 Hz, 1H), 6.65 (dd, J = 8.1, 2.5 Hz, 1H), 5.42 (s, 2H), 3.32 (q, J = 6.5 Hz, 2H), 2.78 (q, J = 6.7 Hz, 2H), 2.36 (s, 3H), 1.65 (p, J = 6.9 Hz, 2H).

**$^{13}$C-NMR** (101 MHz, DMSO) δ 159.33, 158.17, 156.58, 146.46, 138.47, 132.90, 122.23, 117.49, 114.05, 102.65, 39.88, 38.36, 28.41, 18.75.

**MS (ESI+)** *m/z* $[M+H]^+$: calc. 436.00, found 436.00

**HPLC**: $t_R$ = 3.959 min (M2), purity = 93% (UV: 254/280 nm)

*5-Amino-N-(3-((5-bromo-2-chloropyrimidin-4-yl)amino)propyl)-2-chlorobenzenesulfonamide* (**68**)

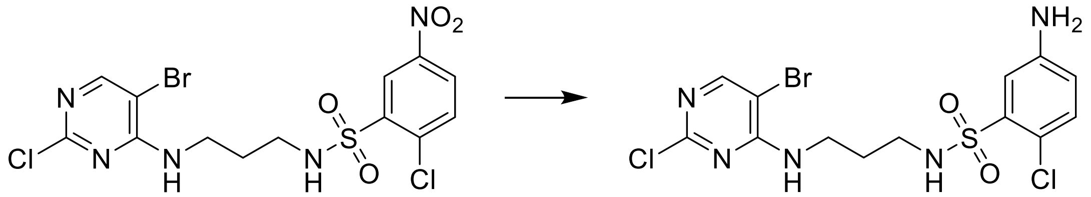

**68** was prepared according to the method for **5**. The product was obtained as a white solid (1.77 g, 95%).

**$^1$H-NMR** (400 MHz, DMSO-$d_6$) δ 8.22 (d, J = 0.6 Hz, 1H), 7.61 (t, J = 5.8 Hz, 1H), 7.56 (t, J = 6.0 Hz, 1H), 7.21 – 7.16 (m, 2H), 6.69 (dd, J = 8.6, 2.8 Hz, 1H), 5.68 (s, 2H), 3.38 – 3.30 (m, 2H), 2.84 (q, J = 6.7 Hz, 2H), 1.65 (p, J = 6.8 Hz, 2H).

**$^{13}$C-NMR** (101 MHz, DMSO) δ 159.35, 158.17, 156.58, 148.11, 137.64, 131.83, 117.97, 115.07, 114.88, 102.66, 38.24, 28.39.

**MS (ESI+)** *m/z* $[M+H]^+$: calc. 455.94, found 455.95

**HPLC**: $t_R$ = 3.986 min (M2), purity = 88% (UV: 254/280 nm)

*5-Amino-N-(3-((5-bromo-2-chloropyrimidin-4-yl)amino)propyl)-2-methoxybenzenesulfon-amide* (**69**)

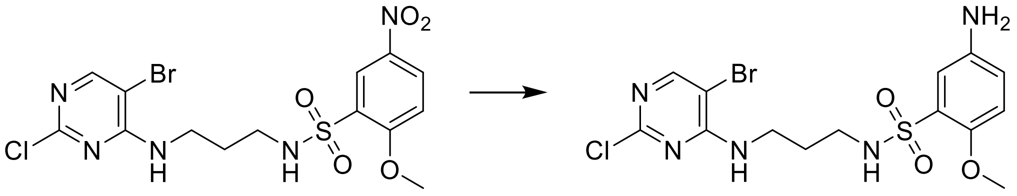

**69** was prepared according to the method for **5**. The product was obtained as a white solid (1.85 g, 85%).

**$^1$H-NMR** (400 MHz, DMSO-$d_6$) δ 8.22 (s, 1H), 7.61 (t, J = 5.8 Hz, 1H), 7.02 (d, J = 2.8 Hz, 1H), 6.98 (t, J = 6.0 Hz, 1H), 6.90 (d, J = 8.8 Hz, 1H), 6.74 (dd, J = 8.7, 2.9 Hz, 1H), 5.04 (s, 2H), 3.75 (s, 3H), 3.36 – 3.28 (m, 2H), 2.78 (q, J = 6.7 Hz, 2H), 1.62 (p, J = 6.9 Hz, 2H).

**$^{13}$C-NMR** (101 MHz, DMSO) δ 159.35, 158.18, 156.57, 147.07, 142.18, 128.20, 118.93, 114.42, 114.29, 102.65, 56.45, 38.24, 28.30.

**MS (ESI+)** *m/z* $[M+H]^+$: calc. 451.99, found 451.95

**HPLC**: $t_R$ = 3.476 min (M2), purity = ≧ 95% (UV: 254/280 nm)

*1⁵-Bromo-3⁴-methyl-4-thia-2,5,9-triaza-1(2,4)-pyrimidina-3(1,3)-benzenacyclononaphane 4,4-dioxide* (**70**)

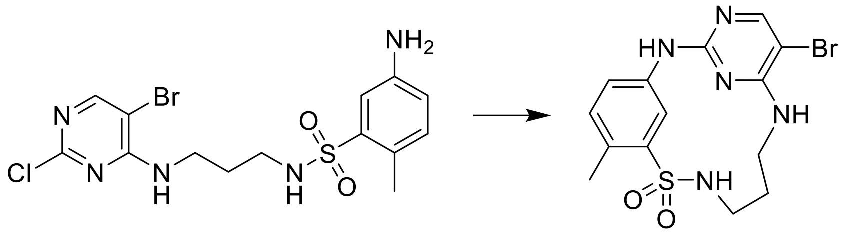

**70** was prepared according to the method for **6**. The product was obtained as a white solid (616 mg, 65%).

**$^{1}$H-NMR** (400 MHz, DMSO-d$_6$) δ 10.55 (s, 1H), 9.32 (d, J = 2.3 Hz, 1H), 8.46 (s, 1H), 8.23 (s, 1H), 8.00 (t, J = 6.1 Hz, 1H), 7.30 (d, J = 8.2 Hz, 1H), 7.23 (dd, J = 8.2, 2.3 Hz, 1H), 3.53 (dt, J = 11.6, 5.6 Hz, 2H), 3.33 (q, J = 5.8 Hz, 2H), 2.51 (s, 3H), 1.92 – 1.80 (m, 2H).

**$^{13}$C-NMR** (101 MHz, DMSO) δ 158.50, 153.04, 147.95, 143.12, 137.15, 132.39, 129.18, 121.11, 116.25, 92.09, 40.55, 27.17, 18.96.

**MS (ESI+)** *m/z* $[M+H]^+$: calc. 398.02, found 398.00

**HRMS** calculated *m/z* for $[C_{14}H_{16}BrN_5O_2S+H]^+$: 398.02808, found: 398.02802

**HPLC**: $t_R$ = 3.328 min (M2), purity = ≧ 95% (UV: 254/280 nm)

*1⁵-Bromo-3⁴-chloro-4-thia-2,5,9-triaza-1(2,4)-pyrimidina-3(1,3)-benzenacyclononaphane 4,4-dioxide* (**71**)

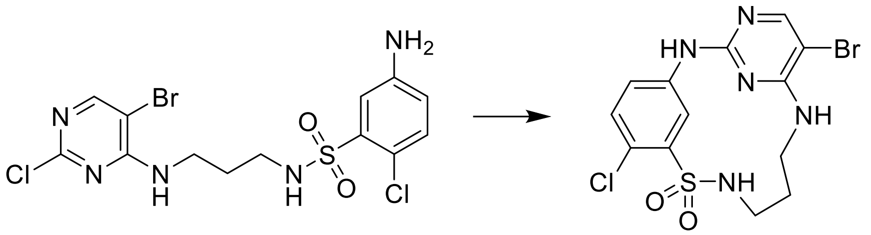

**71** was prepared according to the method for **6**. The product was obtained as a white solid (532 mg, 88%).

**$^{1}$H-NMR** (400 MHz, DMSO-d$_6$) δ 9.85 (s, 1H), 9.59 (d, J = 2.7 Hz, 1H), 8.13 (t, J = 6.1 Hz, 1H), 8.07 (s, 1H), 7.52 (t, J = 5.4 Hz, 1H), 7.45 (d, J = 8.6 Hz, 1H), 7.30 (dd, J = 8.7, 2.6 Hz, 1H), 3.60 – 3.42 (m, 2H), 3.36 – 3.31 (m, 2H), 2.02 – 1.69 (m, 2H).

**$^{13}$C-NMR** (101 MHz, DMSO) δ 158.00, 157.85, 155.93, 141.89, 140.37, 131.46, 121.56, 120.80, 116.65, 92.70, 27.77.

**MS (ESI+)** *m/z* $[M+H]^+$: calc. 419.96, found 419.90

**HRMS** calculated *m/z* for $[C_{13}H_{13}BrClN_5O_2S+H]^+$: 398.02808, found: 398.02802

**HPLC**: $t_R$ = 5.628 min (M1), purity = ≧ 95% (UV: 254/280 nm)

*1⁵-Bromo-3⁴-methoxy-4-thia-2,5,9-triaza-1(2,4)-pyrimidina-3(1,3)-benzenacyclononaphane 4,4-dioxide* (**72**)

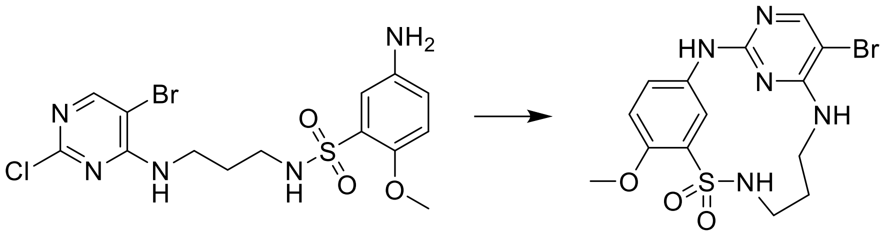

**72** was prepared according to the method for **6**. The product was obtained as a white solid (828 mg, 84%).

**$^1$H-NMR** (400 MHz, DMSO-d$_6$) δ 9.47 (s, 1H), 9.39 (d, J = 2.7 Hz, 1H), 8.00 (s, 1H), 7.80 (t, J = 6.1 Hz, 1H), 7.38 (t, J = 5.3 Hz, 1H), 7.29 (dd, J = 8.8, 2.7 Hz, 1H), 7.09 (d, J = 8.9 Hz, 1H), 3.80 (s, 3H), 3.51 – 3.40 (m, 2H), 3.31 – 3.25 (m, 2H), 1.94 – 1.73 (m, 2H).

**$^{13}$C-NMR** (101 MHz, DMSO) δ 158.16, 157.98, 155.97, 150.61, 134.15, 131.97, 122.00, 116.90, 113.65, 91.71, 56.31, 27.93.

**MS (ESI+)** *m/z* [M+H]$^+$: calc. 416.01, found 415.90

**HRMS** calculated *m/z* for [$C_{14}H_{16}BrN_5O_3S$+H]$^+$: 414.02300, found: 414.02238

**HPLC**: $t_R$ = 4.514 min (M1), purity = ≧ 95% (UV: 254/280 nm)

*1⁵-(4-fluoro-3-methoxyphenyl)-3⁴-methyl-4-thia-2,5,9-triaza-1(2,4)-pyrimidina-3(1,3)-benzenacyclononaphane 4,4-dioxide* (**73**)

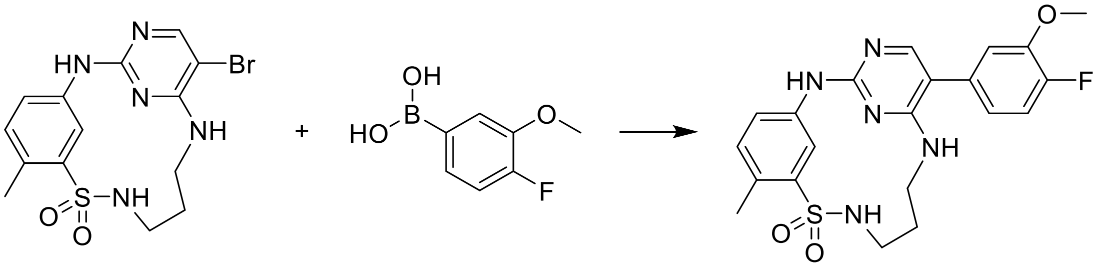

**73** was prepared according to the method for **7**, using **70** and 4-fluoro-3-methoxyphenylboronic acid. The product was obtained as a white solid (33 mg, 30%).

**$^1$H-NMR** (400 MHz, DMSO-d$_6$) δ 9.56 (d, J = 2.1 Hz, 1H), 9.50 (s, 1H), 7.88 (t, J = 6.2 Hz, 1H), 7.81 (s, 1H), 7.27 (dd, J = 11.6, 8.3 Hz, 1H), 7.24 – 7.17 (m, 2H), 7.13 (dd, J = 8.4, 2.1 Hz, 1H), 6.99 (t, J = 5.4 Hz, 1H), 6.92 (ddd, J = 8.3, 4.3, 2.1 Hz, 1H), 3.88 (s, 3H), 3.51 – 3.39 (m, 2H), 3.32 – 3.26 (m, 2H), 2.49 (s, 3H), 1.95 – 1.83 (m, 2H).

**$^{13}$C-NMR** (101 MHz, DMSO) δ 159.27, 158.73, 154.83, 152.01, 149.59, 147.25 (d, J = 10.7 Hz), 143.05, 139.65, 131.94 – 131.84 (m), 126.53, 120.97 (d, J = 6.8 Hz), 120.04, 116.17 (d, J = 17.8 Hz), 115.19, 114.34 (d, J = 1.4 Hz), 109.91, 55.87, 28.20, 18.90.

**MS (ESI+)** *m/z* [M+H]$^+$: calc. 444.14, found 444.10

**HRMS** calculated *m/z* for [$C_{21}H_{23}FN_5O_3S$+H]$^+$: 444.15001, found: 444.14968

**HPLC**: $t_R$ = 3.360 min (M2), purity ≧ 95% (UV: 254/280 nm)

*3⁴-chloro-1⁵-(4-fluoro-3-methoxyphenyl)-4-thia-2,5,9-triaza-1(2,4)-pyrimidina-3(1,3)-benzenacyclononaphane 4,4-dioxide* (**74**)

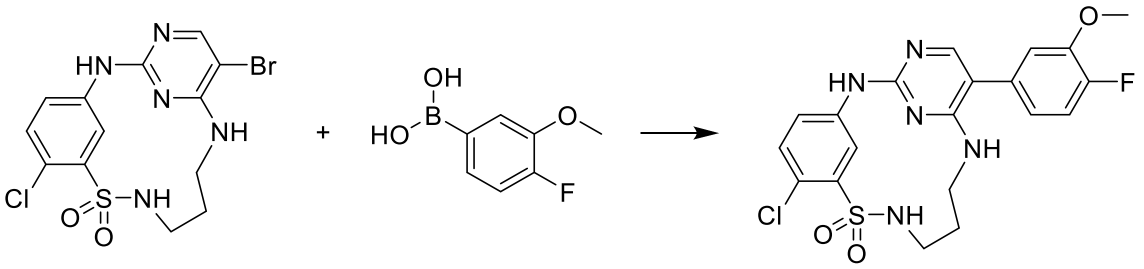

**74** was prepared according to the method for **7**, using **71** and 4-fluoro-3-methoxyphenylboronic acid. The product was obtained as a white solid (27 mg, 29%).

**$^1$H-NMR** (400 MHz, DMSO-d$_6$) δ 10.29 (s, 1H), 9.65 (d, J = 2.7 Hz, 1H), 8.19 (t, J = 6.1 Hz, 1H), 7.87 (s, 1H), 7.67 – 7.59 (m, 1H), 7.52 (d, J = 8.6 Hz, 1H), 7.37 – 7.27 (m, 2H), 7.17 (dd, J = 8.3, 2.1 Hz, 1H), 6.95 (ddd, J = 8.3, 4.3, 2.1 Hz, 1H), 3.88 (s, 3H), 3.51 – 3.41 (m, 2H), 3.39 – 3.30 (m, 2H), 1.95 – 1.80 (m, 2H).

**$^{13}$C-NMR** (101 MHz, DMSO) δ 159.74, 155.34 (d, J = 1.4 Hz), 152.45, 148.66 (d, J = 273.2 Hz), 147.41, 141.99, 139.59, 131.69, 130.25, 121.97, 121.73, 121.34 (d, J = 7.1 Hz), 117.22, 116.35 (d, J = 18.0 Hz), 114.68 – 114.60 (m), 110.99, 55.96, 27.61.

**MS (ESI+)** *m/z* [M+H]$^+$: calc. 464.09, found 464.00

**HRMS** calculated *m/z* for $[C_{20}H_{20}ClFN_5O_3S+H]^+$: 464.09539, found: 464.09500

**HPLC**: $t_R$ = 3.425 min (M2), purity ≧ 95% (UV: 254/280 nm)

*1⁵-(4-Fluoro-3-methoxyphenyl)-3⁴-chloro-4-thia-2,5,9-triaza-1(2,4)-pyrimidina-3(1,3)-benzenacyclononaphane 4,4-dioxide* (**75**)

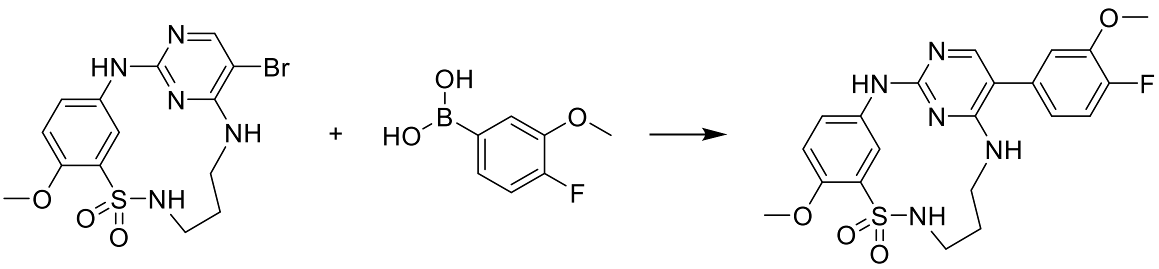

**75** was prepared according to the method for **7**, using **72** and 4-fluoro-3-methoxyphenylboronic acid. The product was obtained as a white solid (27 mg, 29%).

**$^1$H-NMR** (500 MHz, DMSO-d6) δ 9.51 (d, J = 2.7 Hz, 1H), 9.38 (s, 1H), 7.82 – 7.76 (m, 2H), 7.31 (dd, J = 8.8, 2.7 Hz, 1H), 7.26 (dd, J = 11.6, 8.3 Hz, 1H), 7.11 (dd, J = 8.5, 2.1 Hz, 1H), 7.09 (d, J = 8.9 Hz, 1H), 6.97 (t, J = 5.4 Hz, 1H), 6.91 (ddd, J = 8.2, 4.3, 2.1 Hz, 1H), 3.88 (s, 3H), 3.81 (s, 3H), 3.51 – 3.35 (m, 2H), 3.31 – 3.21 (m, 2H), 1.96 – 1.72 (m, 2H).

**$^{13}$C-NMR** (126 MHz, DMSO-d6) δ 159.75, 159.13, 155.16, 151.25 (d, J = 243.7 Hz), 150.86, 147.72 (d, J = 10.7 Hz), 135.06, 132.52, 132.42 (d, J = 3.6 Hz), 122.24, 121.43 (d, J = 6.7 Hz), 117.34, 116.64 (d, J = 17.9 Hz), 114.83 – 114.78 (m), 114.11, 110.13, 56.80, 56.35, 28.70.

**MS (ESI+)** *m/z* $[M+H]^+$: calc. 460.14, found 460.10

**HRMS** calculated *m/z* for $[C_{21}H_{22}FN_5O_4S+H]^+$: 460.14493, found: 460.144333

**HPLC**: $t_R$ = 3.002 min (M2), purity ≧ 95% (UV: 254/280 nm)

*$1^5$-(4-Fluoro-3-methoxyphenyl)-$3^4$-(4-hydroxyphenyl)-4-thia-2,5,9-triaza-1(2,4)-pyrimidina-3(1,3)-benzenacyclononaphane 4,4-dioxide* (**76**)

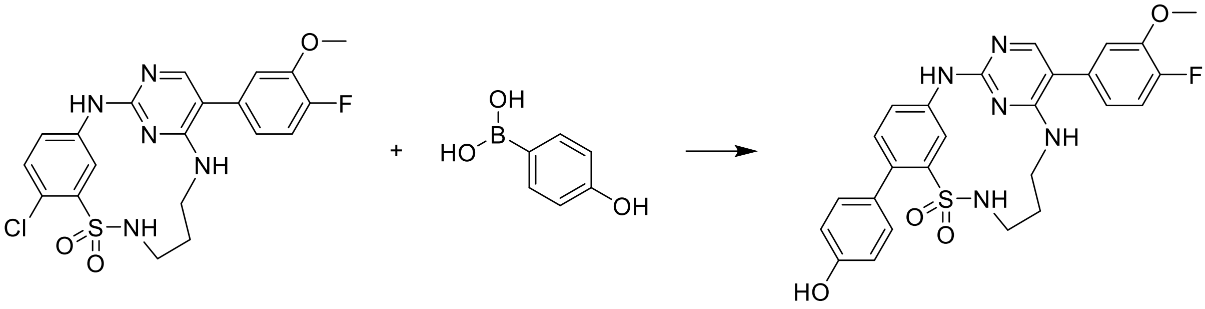

**74** (1 eq.), XPhos Pd G2 (0.05 eq.), 2 M aq. potassium carbonate (3 eq.) and 4-hydroxyphenylboronic acid (1.5 eq.), were dissolved in 1,4-dioxane and DMF (1:1, 0.125 M) and degassed with Argon (g). The reaction mixture was then heated under µW irradiation at 100 °C for 2 h. The crude product was purified by flash column chromatography on silica gel using dichloromethane / ethyl acetate as an eluent to afford the product as a white solid (41 mg, 65%).

**$^1$H-NMR** (400 MHz, DMSO-$d_6$) δ 10.74 (s, 1H), 9.48 (d, J = 2.2 Hz, 1H), 8.12 (s, 1H), 7.98 – 7.82 (m, 3H), 7.29 (ddt, J = 34.7, 18.2, 8.2 Hz, 7H), 6.98 (t, J = 6.4 Hz, 1H), 6.76 (d, J = 8.1 Hz, 2H), 3.89 (s, 2H), 3.54 – 3.42 (m, 2H), 3.34 – 3.25 (m, 2H), 1.96 – 1.80 (m, 2H).

**$^{13}$C NMR** (101 MHz, DMSO) δ 162.78, 160.64, 157.14, 150.76, 147.87 (d, J = 11.0 Hz), 143.14, 138.36, 134.96, 133.70, 131.25, 130.83, 129.46, 122.05 (d, J = 7.0 Hz), 121.76 – 121.57 (m), 117.34, 117.00, 116.82, 115.32, 114.81, 111.72, 56.46, 40.83, 27.82.

**MS (ESI+)** *m/z* $[M+H]^+$: calc. 522.15, found 522.15

**HRMS** calculated *m/z* for $[C_{26}H_{24}FN_5O_4S+H]^+$: 522.16058, found: 522.15880

**HPLC**: $t_R$ = 3.660 min (M2), purity ≧ 95% (UV: 254/280 nm)

*1^5-Bromo-3^4-hydroxy-4-thia-2,5,9-triaza-1(2,4)-pyrimidina-3(1,3)-benzenacyclononaphane 4,4-dioxide* (**77**)

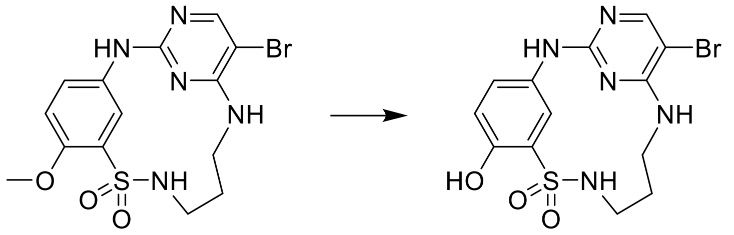

**72** (1 eq.) and sodium ethanethiolate (1.5 eq.) were dissolved in DMF (0.21 M). The reaction mixture was then heated under µW irradiation at 80 °C for 10 h. The crude product was purified by reverse phase flash column chromatography on silica gel using water / acetonitrile as an eluent to afford the product as a white solid (45 mg, 54%).

**$^1$H-NMR** (400 MHz, DMSO-d$_6$) δ 9.67 (s, 1H), 9.35 (s, 1H), 9.24 (d, J = 2.7 Hz, 1H), 7.98 (s, 1H), 7.73 (t, J = 6.1 Hz, 1H), 7.33 (t, J = 5.4 Hz, 1H), 7.15 (dd, J = 8.7, 2.7 Hz, 1H), 6.83 (d, J = 8.7 Hz, 1H), 3.43 (dt, J = 11.2, 5.7 Hz, 2H), 3.28 (q, J = 5.8 Hz, 2H), 1.89 – 1.78 (m, 2H).

**$^{13}$C NMR** (101 MHz, DMSO) δ 158.21, 157.92, 155.97, 148.94, 132.64, 129.42, 122.58, 117.19, 117.05, 91.41, 28.09.

**MS (ESI+)** *m/z* [M+H]$^+$: calc. 402.00, found 401.95

**HPLC**: $t_R$ = 3.010 min (M2), purity ≧ 95% (UV: 254/280 nm)

*1^5-(4-Fluoro-3-methoxyphenyl)-3^4-hydroxy-4-thia-2,5,9-triaza-1(2,4)-pyrimidina-3(1,3)-benzenacyclononaphane 4,4-dioxide* (**78**)

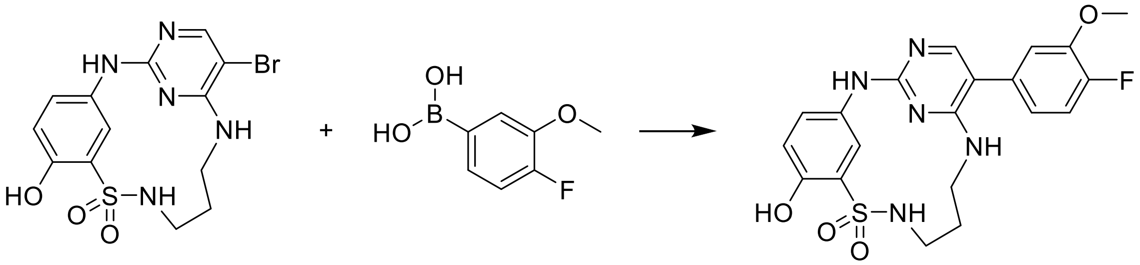

**78** was prepared according to the method for **7**, using **77** and 4-fluoro-3-methoxyphenylboronic acid. The product was obtained as a white solid (56 mg, 74%).

**$^1$H-NMR** (400 MHz, DMSO-d6) δ 10.79 (s, 1H), 10.17 (s, 1H), 9.21 (d, J = 2.7 Hz, 1H), 8.25 (s, 1H), 7.92 – 7.78 (m, 2H), 7.34 (dd, J = 11.5, 8.2 Hz, 1H), 7.25 – 7.15 (m, 2H), 7.02 – 6.92 (m, 2H), 3.88 (s, 3H), 3.46 (q, J = 5.5 Hz, 2H), 3.29 (q, J = 5.8 Hz, 2H), 1.92 – 1.79 (m, 2H).

**$^{13}$C-NMR** (101 MHz, DMSO-d6) δ 160.30, 151.60 (d, J = 245.3 Hz), 151.26, 150.91, 147.41 (d, J = 10.7 Hz), 142.45, 129.67 (d, J = 22.5 Hz), 128.54 (d, J = 3.0 Hz), 123.75, 121.73 – 121.53 (m), 118.73, 117.64, 116.45 (d, J = 17.8 Hz), 114.95, 114.94, 110.92, 56.01, 40.64, 27.24.

**MS (ESI+)** *m/z* [M+H]$^+$: calc. 446.12, found 446.05

**HRMS** calculated *m/z* for [$C_{20}H_{20}FN_5O_4S$+H]$^+$: 446.12928, found: 446.12767

**HPLC**: $t_R$ = 3.262 min (M2), purity ≧ 95% (UV: 254/280 nm)

*1⁵-(4-Fluoro-3-methoxyphenyl)-3⁴-(2-methoxyethoxy)-4-thia-2,5,9-triaza-1(2,4)-pyrimidina-3(1,3)-benzenacyclononaphane 4,4-dioxide* (**79**)

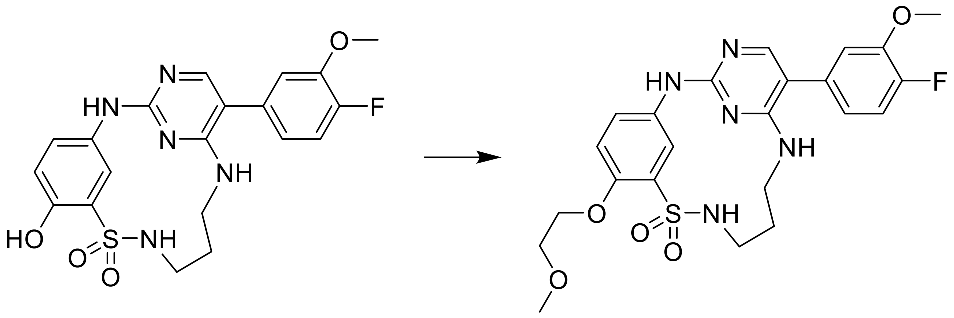

**78** (1 eq.) and potassium carbonate (3 eq.) were dissolved in DMF (0.05 M), 1-bromo-2-methoxyethane (1 eq.) was added and the resulting mixture was stirred for 2 h at room temperature. The crude product was purified by reverse flash column chromatography, using $H_2O$ / acetonitrile as eluent. The product was obtained as a white solid (6 mg, 25%).

**$^1$H-NMR** (400 MHz, DMSO-d6) δ 9.52 – 9.44 (m, 2H), 7.84 – 7.75 (m, 2H), 7.34 – 7.22 (m, 2H), 7.17 – 7.02 (m, 3H), 6.96 – 6.87 (m, 1H), 4.14 (t, J = 4.9 Hz, 2H), 3.88 (s, 3H), 3.67 (t, J = 4.8 Hz, 2H), 3.48 – 3.39 (m, 2H), 3.35 (s, 3H), 3.34 – 3.22 (m, 2H), 1.86 (s, 2H).

**$^{13}$C-NMR** (101 MHz, DMSO-d6) δ 159.34, 158.08, 153.89, 150.83 (d, J = 243.8 Hz), 149.85, 147.25 (d, J = 10.6 Hz), 134.68, 132.83, 131.70, 121.86, 121.00 (d, J = 6.9 Hz), 116.87, 116.19 (d, J = 18.2 Hz), 115.61, 114.36, 109.78, 70.44, 69.38, 58.52, 55.88, 28.13.

**MS (ESI+)** *m/z* $[M+H]^+$: calc. 504.16, found 504.10

**HRMS** calculated *m/z* for $[C_{23}H_{26}FN_5O_5S+H]^+$: 504.17114, found: 504.17020

**HPLC**: $t_R$ = 3.286 min (M2), purity ≧ 95% (UV: 254/280 nm)

*Methyl 2-((1⁵-(4-fluoro-3-methoxyphenyl)-4,4-dioxido-4-thia-2,5,9-triaza-1(2,4)-pyrimidina-3(1,3)-benzenacyclononaphane-3⁴-yl)oxy)acetate* (**80**)

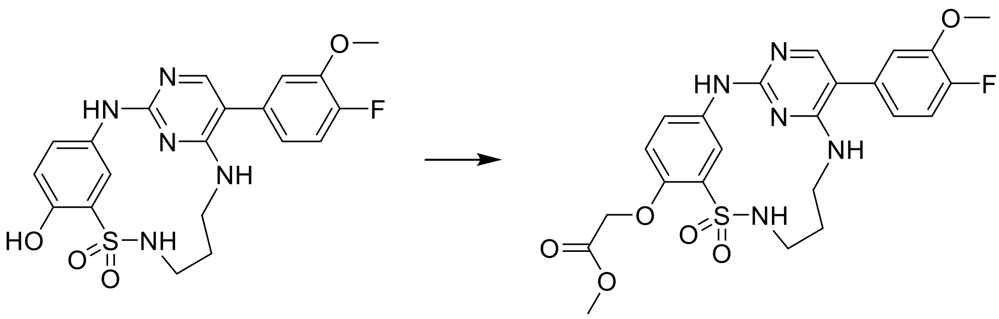

**80** was prepared according to the method for **79**, using methyl bromoacetate. The product was obtained as a white solid (10 mg, 41%).

**¹H-NMR** (400 MHz, DMSO-d6) δ 10.75 (s, 1H), 9.39 (d, J = 2.7 Hz, 1H), 8.20 (s, 1H), 7.95 (t, J = 6.0 Hz, 1H), 7.89 (s, 1H), 7.38 – 7.27 (m, 2H), 7.20 (dd, J = 8.3, 2.1 Hz, 1H), 7.09 (d, J = 8.9 Hz, 1H), 7.01 – 6.94 (m, 1H), 4.90 (s, 2H), 3.88 (s, 3H), 3.72 (s, 3H), 3.54 – 3.43 (m, 2H), 3.37 – 3.26 (m, 2H), 1.94 – 1.80 (m, 2H).

**¹³C-NMR** (101 MHz, DMSO-d6) δ 168.89, 160.26, 151.57 (d, J = 245.5 Hz), 150.55, 147.41 (d, J = 10.9 Hz), 143.51, 132.46, 132.38, 128.74, 122.96, 121.63 (d, J = 7.2 Hz), 118.29, 116.45 (d, J = 18.5 Hz), 115.02 (d, J = 20.8 Hz), 114.90, 111.07, 65.88, 56.01, 51.89, 40.62, 27.17.

**MS (ESI+)** *m/z* $[M+H]^+$: calc. 518.14, found 518.15

**HRMS** calculated *m/z* for $[C_{23}H_{24}FN_5O_6S+H]^+$: 518.15041,found: 518.14970

**HPLC**: $t_R$ = 3.313 min (M2), purity ≧ 95% (UV: 254/280 nm)

*2-((1⁵-(4-Fluoro-3-methoxyphenyl)-4,4-dioxido-4-thia-2,5,9-triaza-1(2,4)-pyrimidina-3(1,3)-benzenacyclononaphane-3⁴-yl)oxy)acetamide* (**81**)

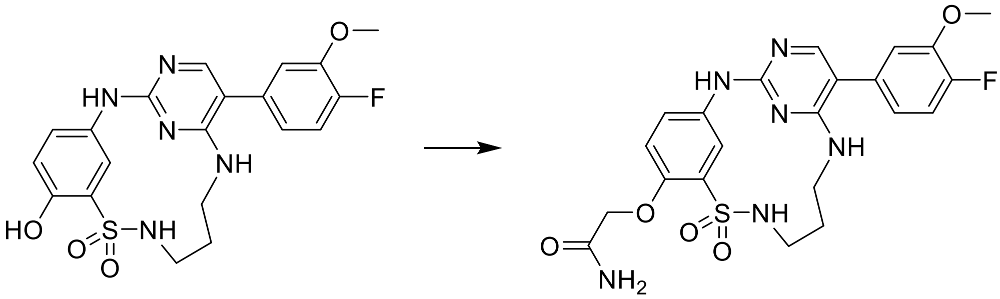

**81** was prepared according to the method for **79**, using 2-bromoacetamide. The product was obtained as a white solid (20 mg, 71%).

**¹H-NMR** (400 MHz, DMSO-d6) δ 10.75 (s, 1H), 9.41 (d, J = 2.7 Hz, 1H), 8.16 (t, J = 5.7 Hz, 2H), 7.90 (s, 1H), 7.71 – 7.64 (m, 1H), 7.37 – 7.31 (m, 2H), 7.28 – 7.24 (m, 2H), 7.20 (dd, J = 8.3, 2.1 Hz, 1H), 6.97 (ddd, J = 8.4, 4.4, 2.1 Hz, 1H), 4.60 (s, 2H), 3.88 (s, 3H), 3.55 – 3.42 (m, 2H), 3.34 (q, J = 5.8 Hz, 2H), 1.94 – 1.81 (m, 2H).

**¹³C-NMR** (101 MHz, DMSO-d6) δ 169.74, 160.23, 152.06, 151.55 (d, J = 245.3 Hz), 149.91, 147.41 (d, J = 10.8 Hz), 143.90, 132.61, 132.02, 128.79, 123.36, 121.60 (d, J = 7.2 Hz), 118.27, 116.45 (d, J = 18.0 Hz), 114.94 (d, J = 9.5 Hz), 114.87, 67.68, 56.00, 40.59, 27.20.

**MS (ESI+)** *m/z* $[M+H]^+$: calc. 503.14, found 503.15

**HRMS** calculated *m/z* for $[C_{22}H_{23}FN_6O_5S+H]^+$: 503.15074,found: 503.15159

**HPLC**: $t_R$ = 3.193 min (M2), purity ≧ 95% (UV: 254/280 nm)

*1[5]-(4-Fluoro-3-methoxyphenyl)-3[4]-(2-morpholinoethoxy)-4-thia-2,5,9-triaza-1(2,4)-pyrimidina-3(1,3)-benzenacyclononaphane 4,4-dioxide 2,2,2-trifluoroacetate* (**82**)

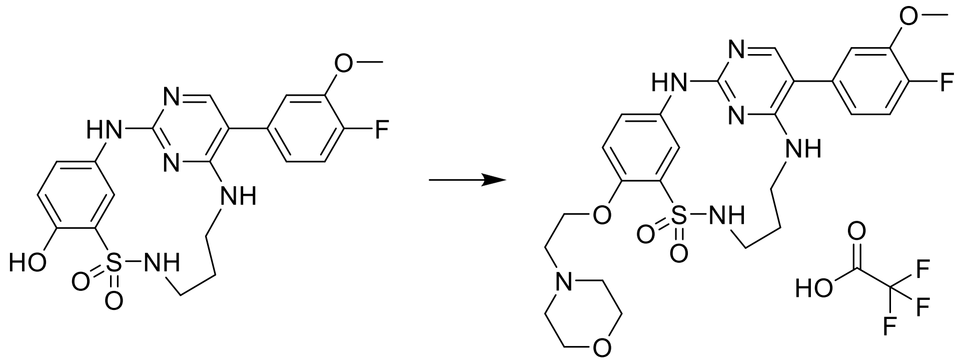

**82** was prepared according to the method for **79**, using 4-(2-bromoethyl)morpholine. The crude product was purified by reverse flash column chromatography, using $H_2O$ / acetonitrile (+ 0.1% TFA) as eluent. The product was obtained as a white solid (33 mg, 55%).

**$^1$H-NMR** (400 MHz, DMSO-d6) δ 11.19 (s, 1H), 10.09 (s, 1H), 9.42 (s, 1H), 8.37 (s, 1H), 8.13 (d, J = 6.3 Hz, 1H), 7.95 (s, 1H), 7.37 (dd, J = 18.6, 8.9 Hz, 3H), 7.21 (d, J = 8.2 Hz, 1H), 6.97 (d, J = 8.6 Hz, 1H), 4.56 – 4.36 (m, 2H), 4.04 – 3.93 (m, 2H), 3.89 (s, 3H), 3.76 – 3.26 (m, 12H), 1.99 – 1.78 (m, 2H).

**$^{13}$C-NMR** (101 MHz, DMSO-d6) δ 160.38, 158.84 (d, J = 35.4 Hz), 151.05, 150.32 (d, J = 25.5 Hz), 147.44 (d, J = 11.0 Hz), 132.51, 132.36, 128.32, 123.32, 121.69 (d, J = 5.4 Hz), 121.64 (d, J = 4.9 Hz), 118.29, 116.40, 115.26, 114.95 (d, J = 3.0 Hz), 111.25, 64.52, 63.44, 56.01, 55.23, 52.43, 40.71, 27.01.

**MS (ESI+)** *m/z* $[M+H]^+$: calc. 559.21, found 559.20

**HRMS** calculated *m/z* for $[C_{26}H_{31}FN_6O_5S+H]^+$: 559.21334,found: 559.21279

**HPLC**: $t_R$ = 3.251 min (M2), purity ≧ 95% (UV: 254/280 nm)

### Analytical data of compounds

$^{1}H$ and $^{13}C$ {H} NMR, HRMS (Orbitrap) and HPLC/MS (ESI) data of compound **6**

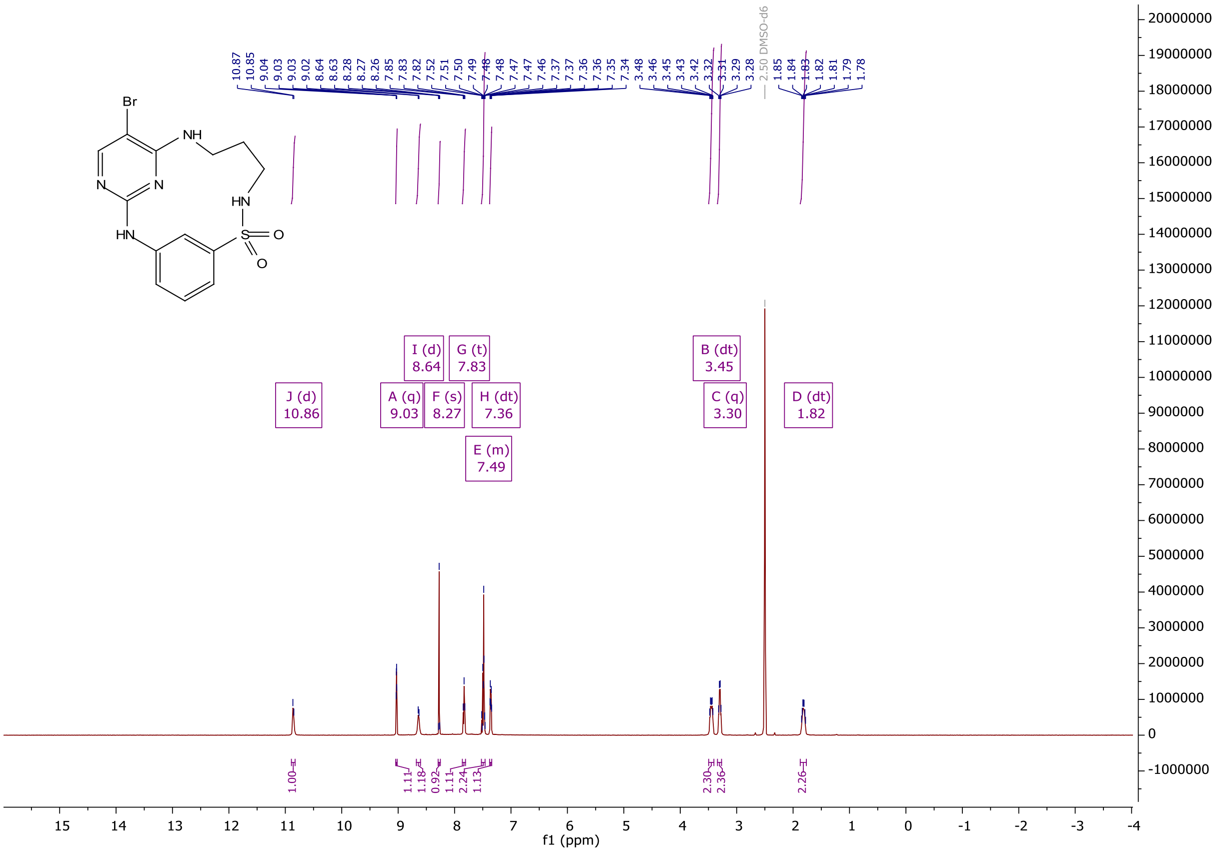

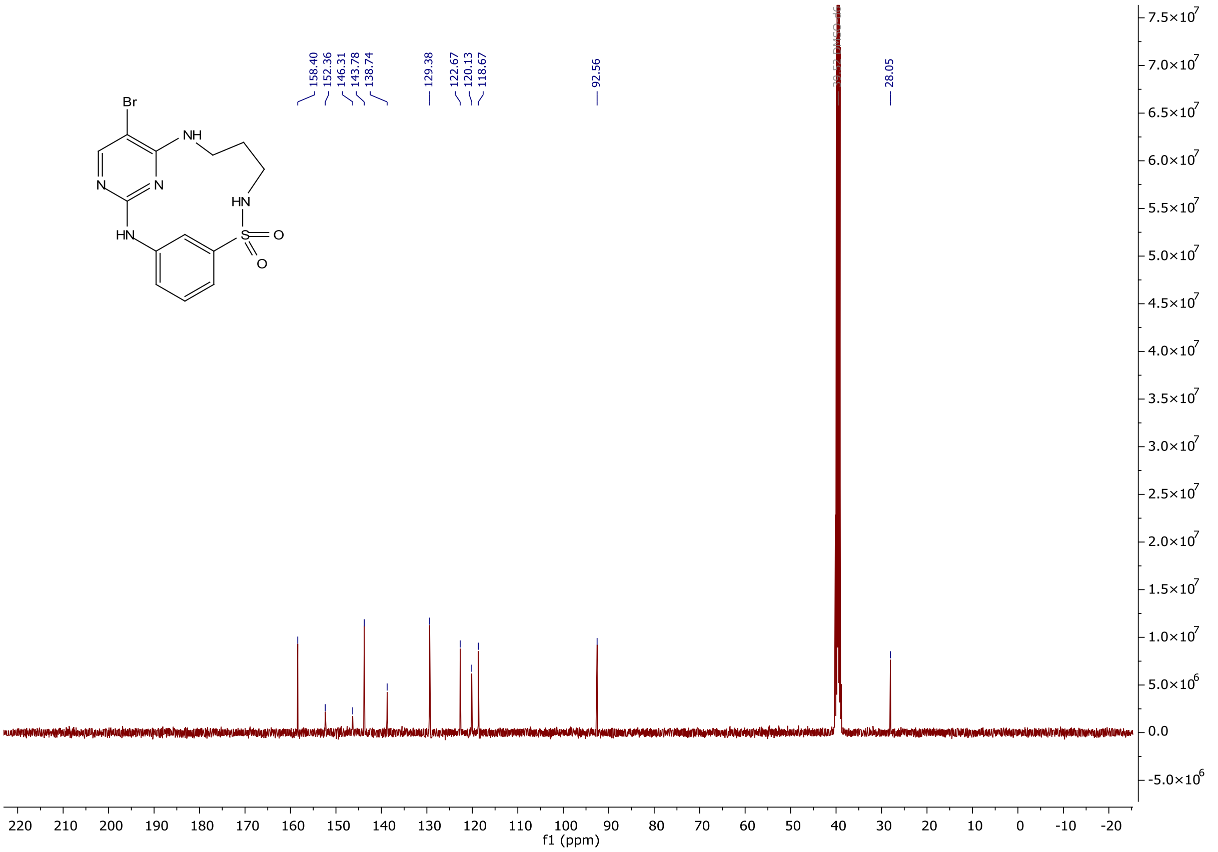

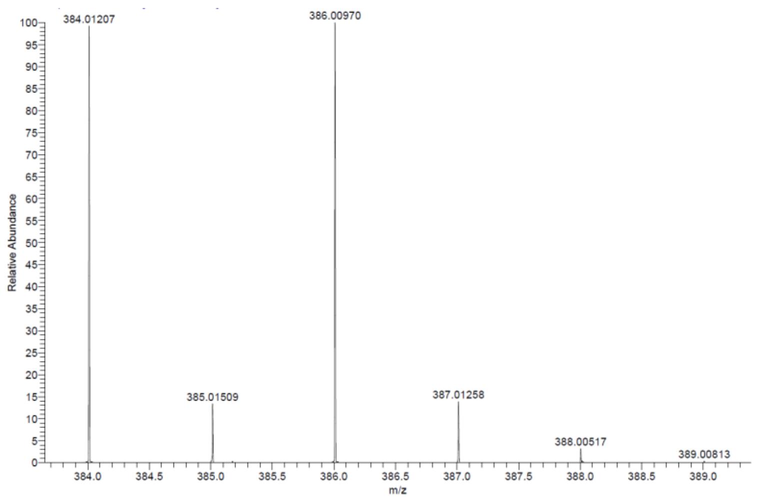

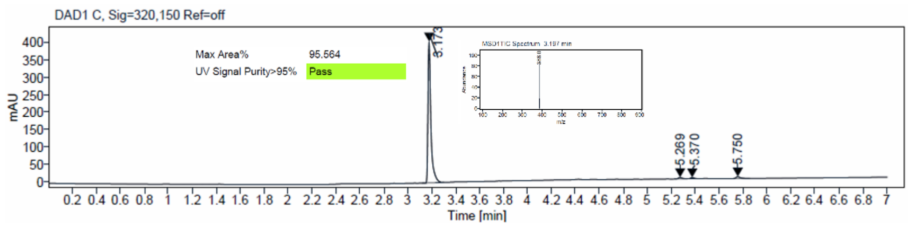

$^1$H and $^{13}$C {H} NMR, HRMS (Orbitrap) and HPLC/MS (ESI) data of compound **7**

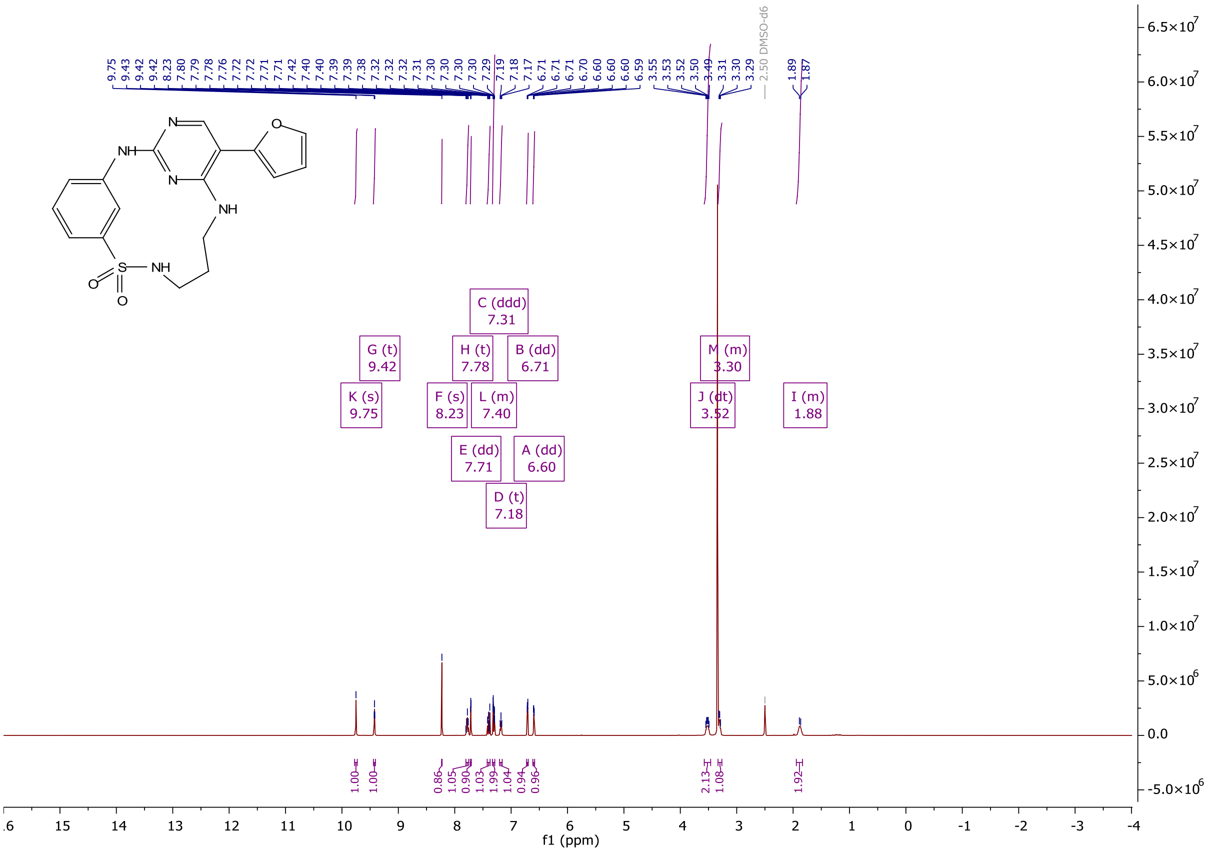

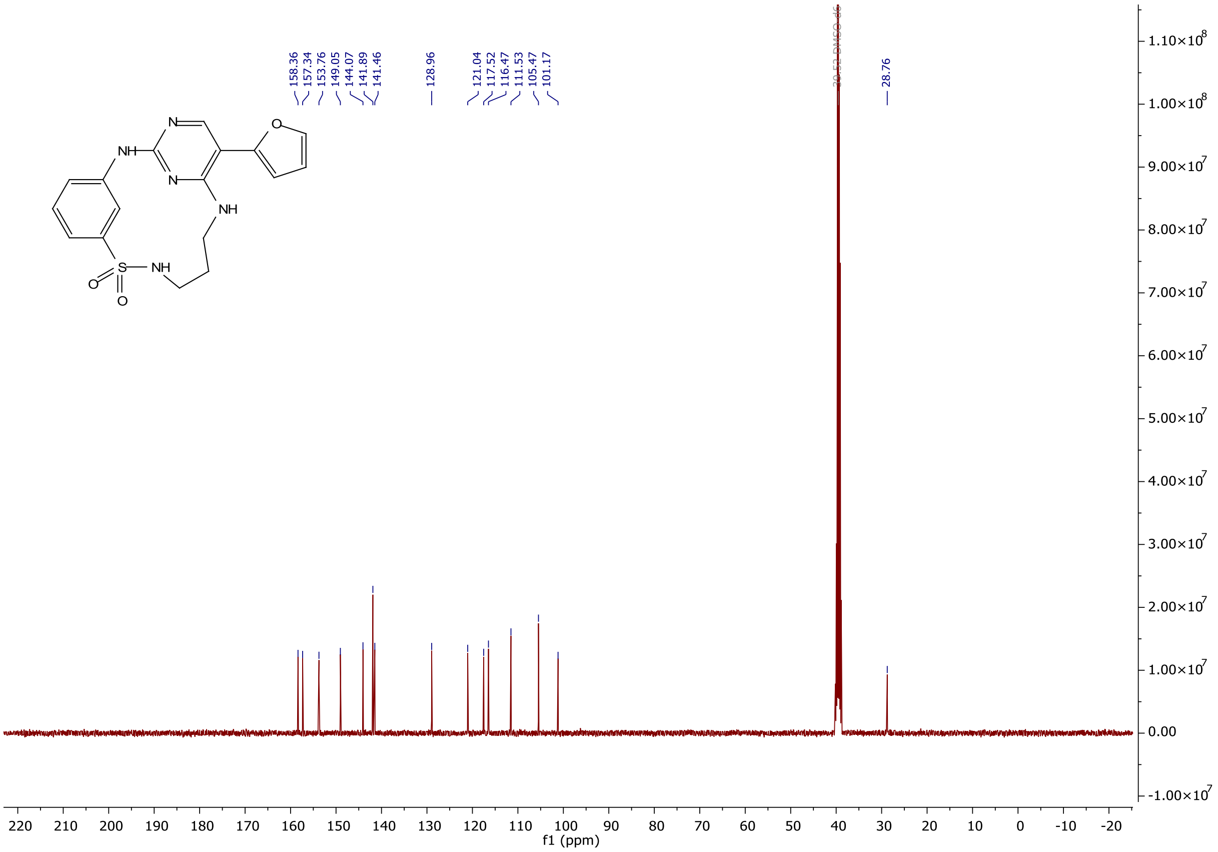
158.36
157.34
153.76
149.05
144.07
141.89
141.46
128.96
121.04
117.52
116.47
111.53
105.47
101.17
28.76
f1 (ppm)

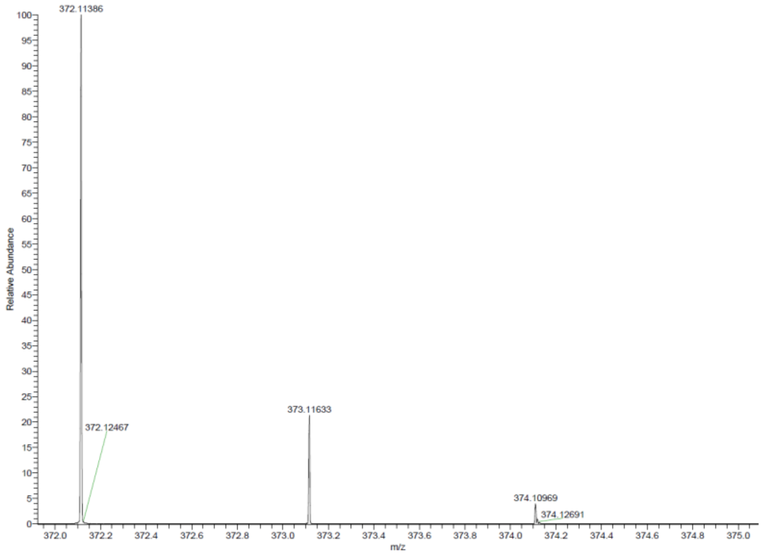
372.11386
372.12467
373.11633
374.10969
374.12691
Relative Abundance
m/z

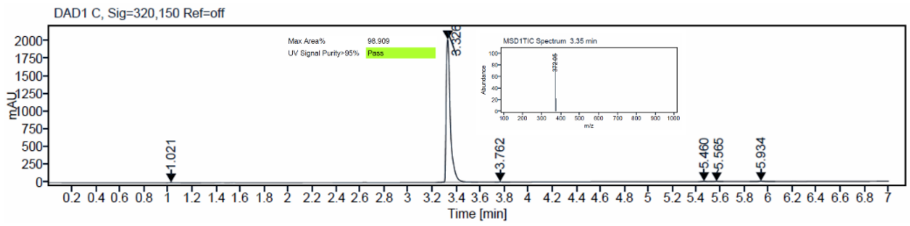
DAD1 C, Sig=320,150 Ref=off
Max Area% 98.909
UV Signal Purity>95% Pass
MSD1TIC Spectrum 3.35 min
1.021
3.762
5.460
5.565
5.934
mAU
Time [min]

$^{1}$H and $^{13}$C {H} NMR, HRMS (Orbitrap) and HPLC/MS (ESI) data of compound **8**

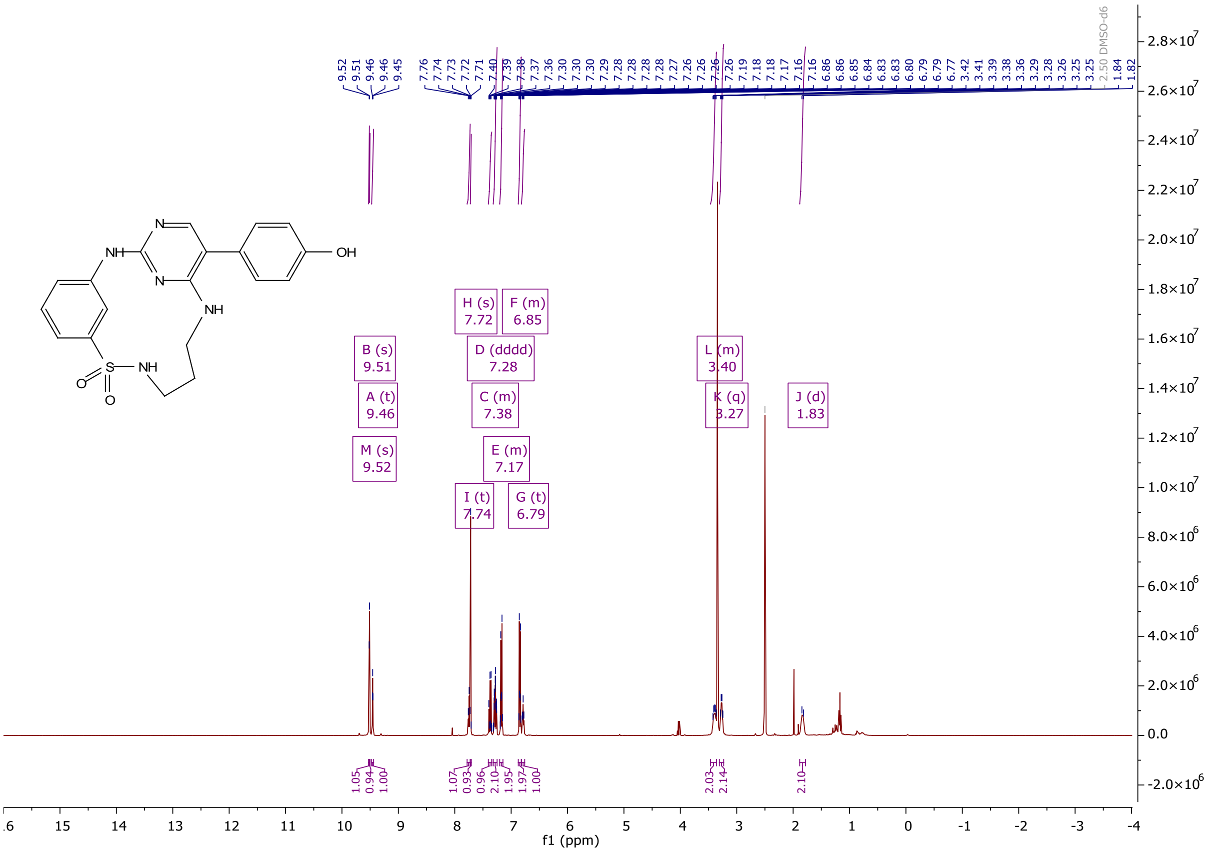

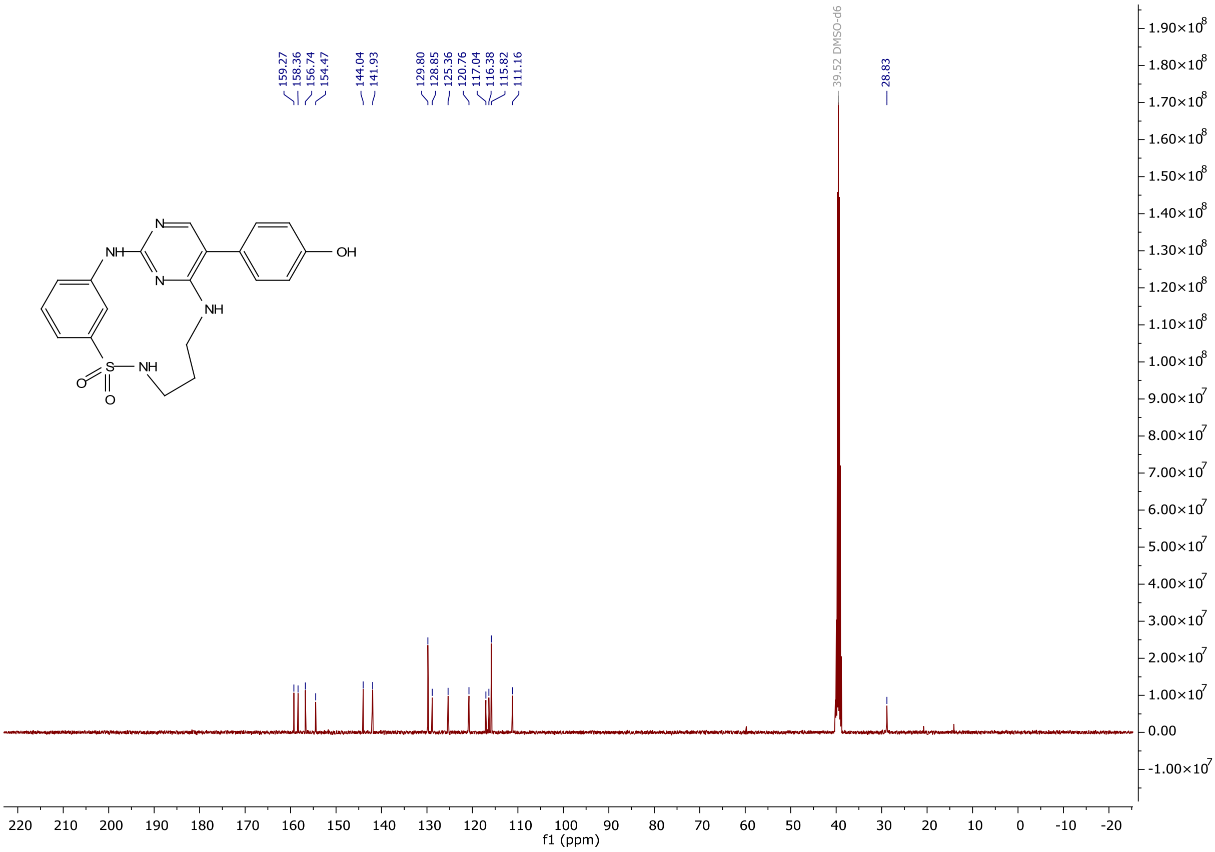

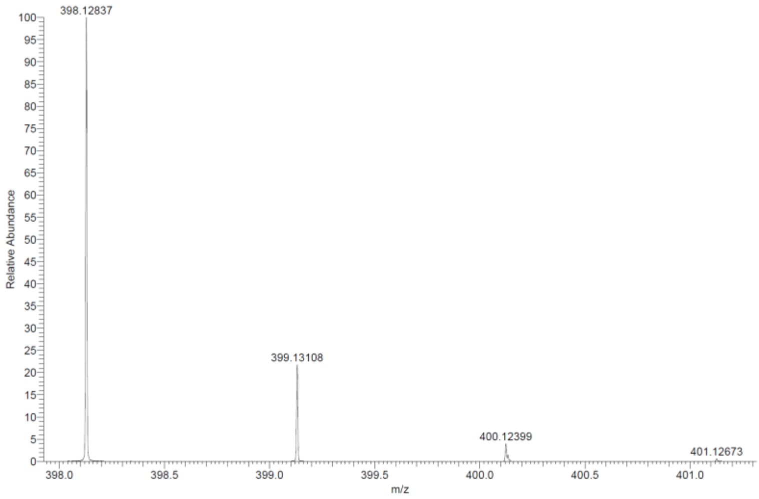

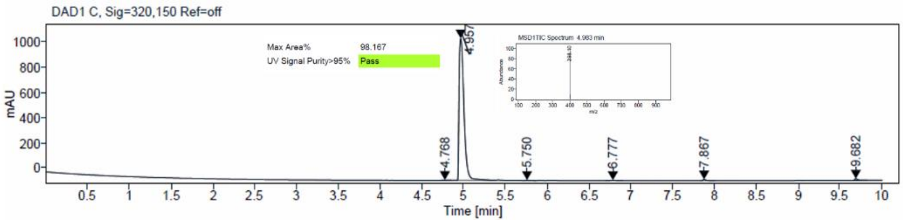

$^{1}H$ and $^{13}C$ {H} NMR, HRMS (Orbitrap) and HPLC/MS (ESI) data of compound **9**

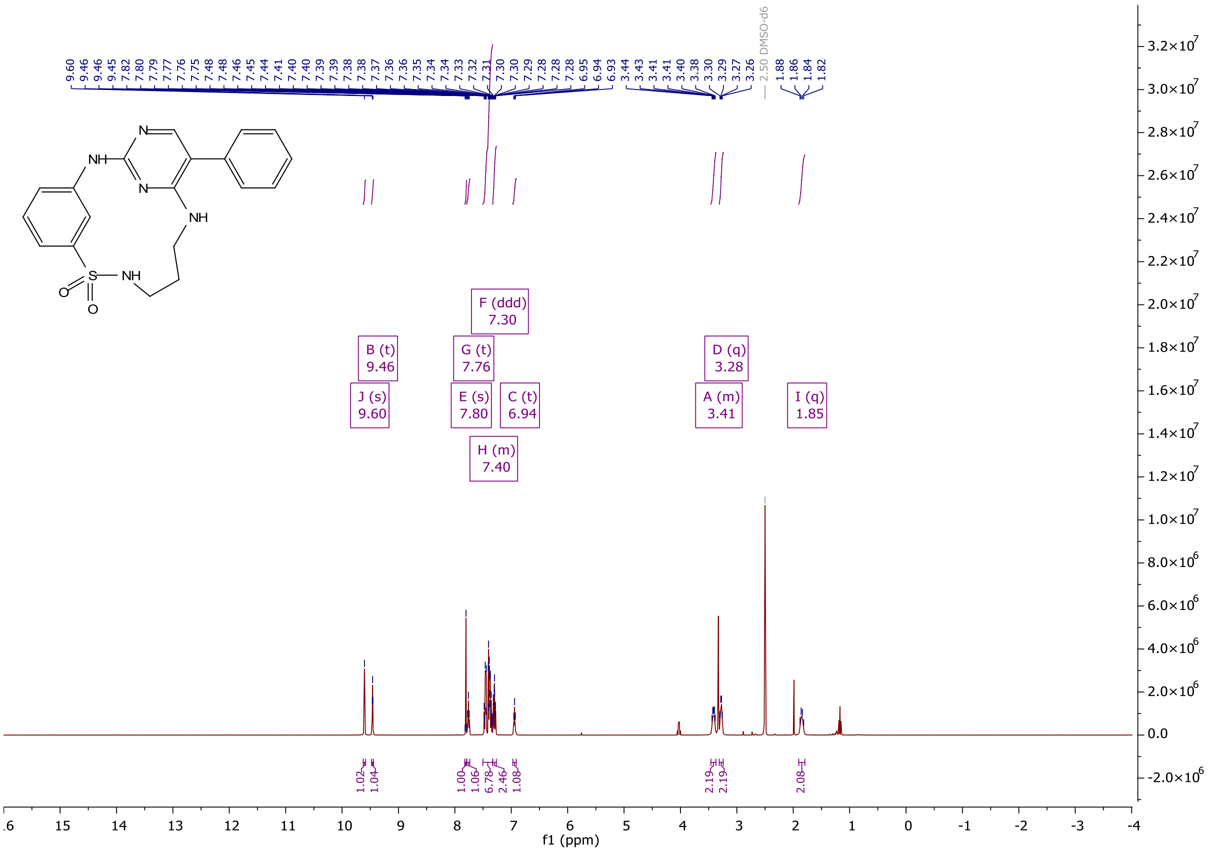

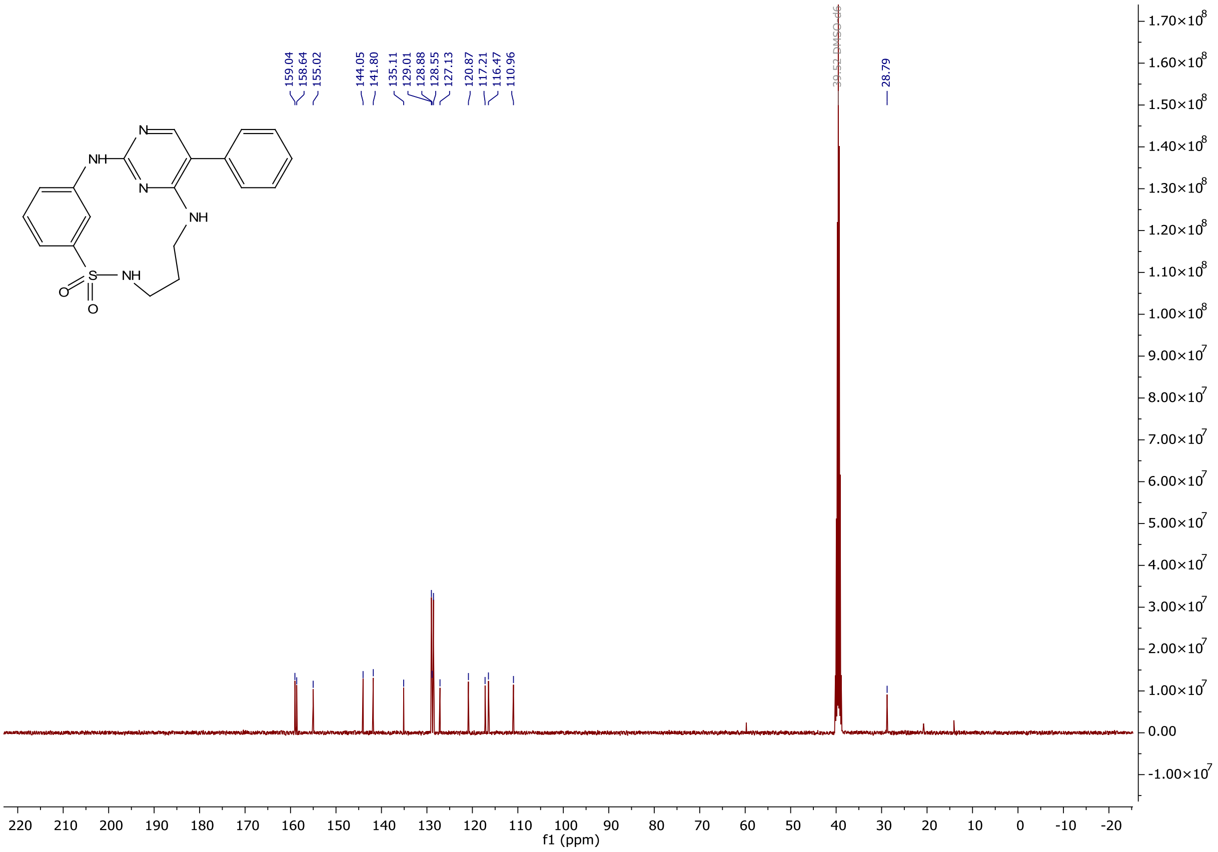
159.04
158.64
155.02
144.05
141.80
135.11
129.01
128.88
128.55
127.13
120.87
117.21
116.47
110.96
39.52 DMSO-d6
28.79
f1 (ppm)

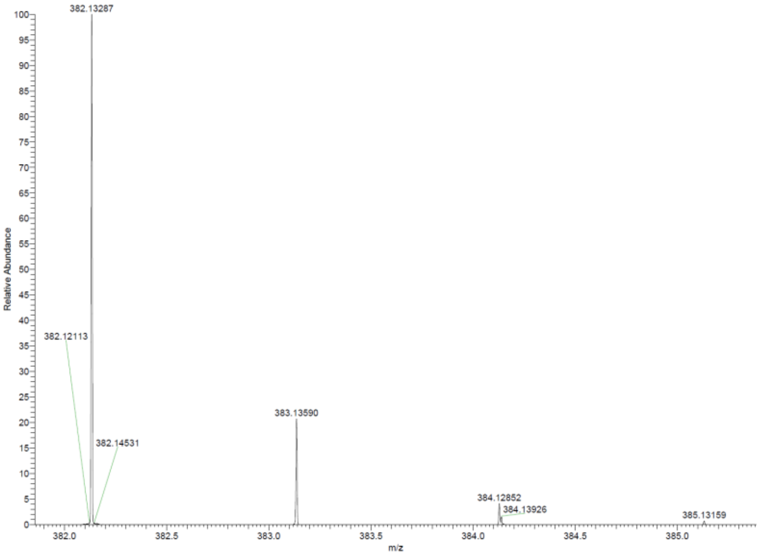
382.13287
382.12113
382.14531
383.13590
384.12852
384.13926
385.13159
Relative Abundance
m/z

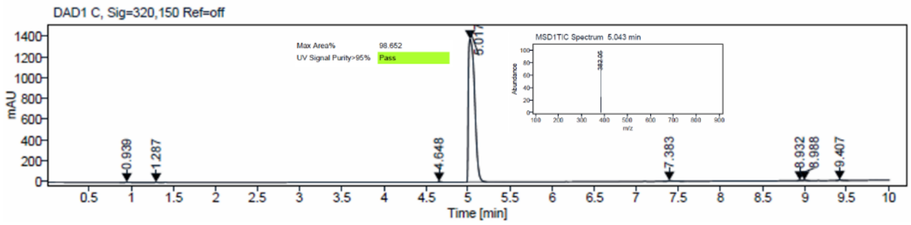
DAD1 C, Sig=320,150 Ref=off
Max Area% 98.652
UV Signal Purity>95% Pass
MSD1TIC Spectrum 5.043 min
mAU
Time [min]

[1]H and [13]C {H} NMR, HRMS (Orbitrap) and HPLC/MS (ESI) data of compound **10**

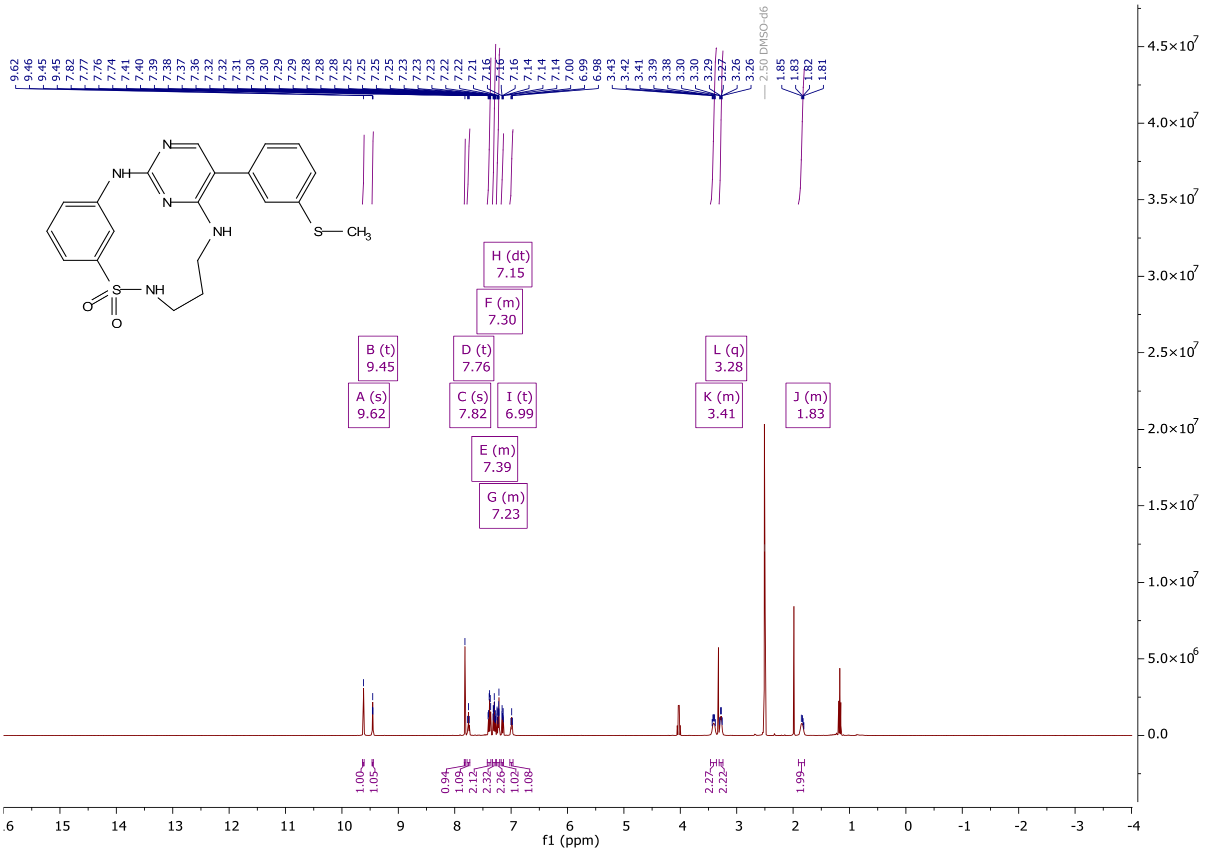

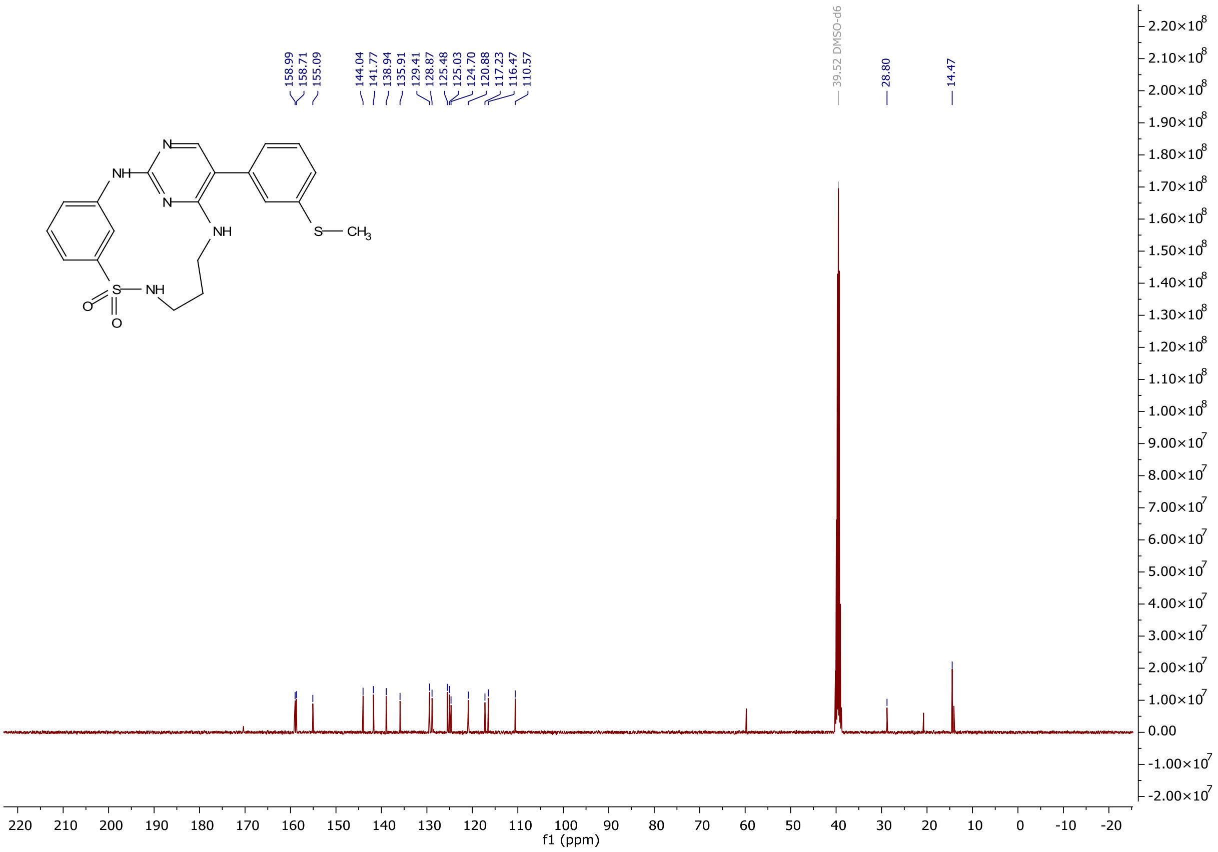

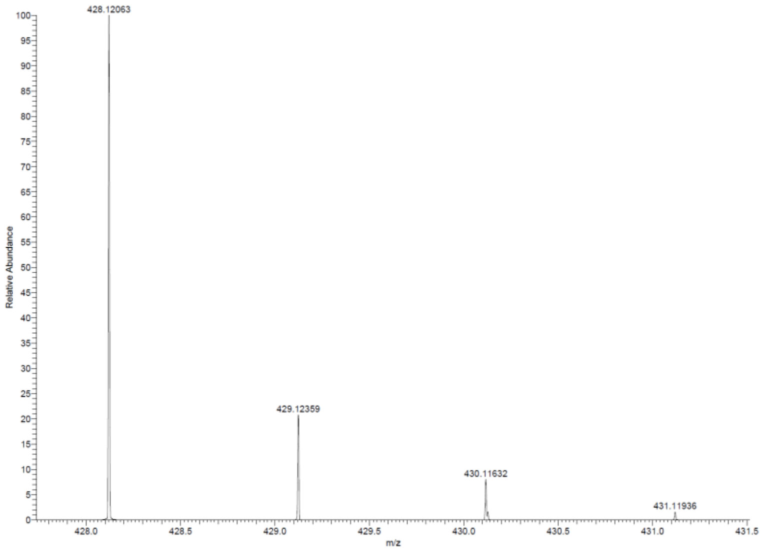

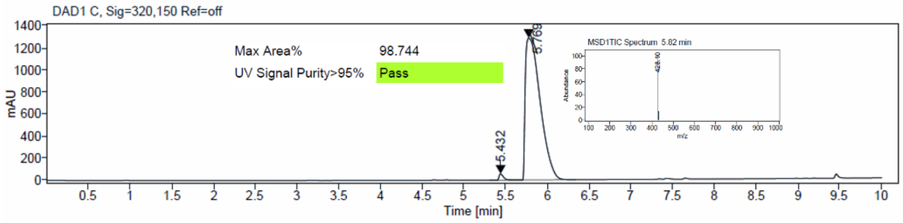

$^{1}H$ and $^{13}C$ {H} NMR, HRMS (Orbitrap) and HPLC/MS (ESI) data of compound **11**

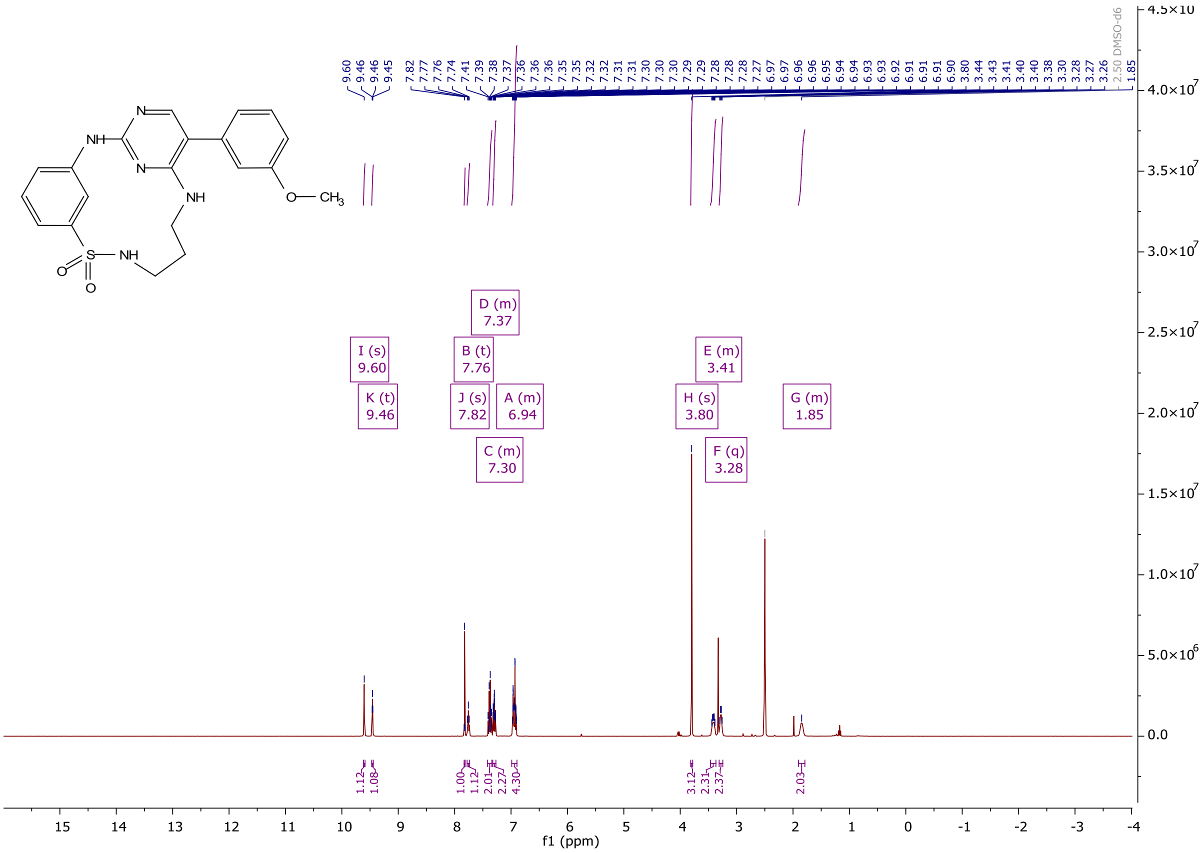

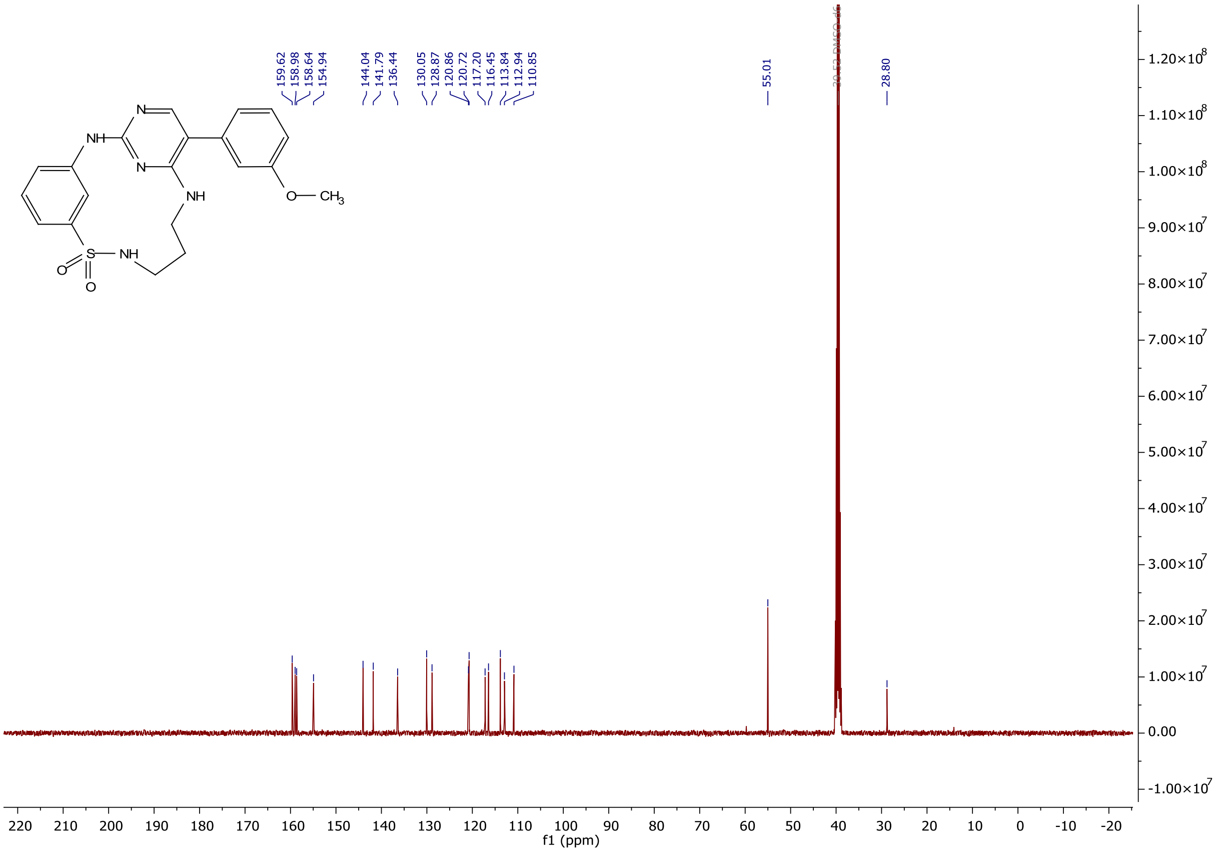
159.62
158.98
158.64
154.94
144.04
141.79
136.44
130.05
128.87
120.86
120.72
117.20
116.45
113.84
112.94
110.85
55.01
28.80
NH
N
N
NH
O—CH3
S
NH
O
O
f1 (ppm)

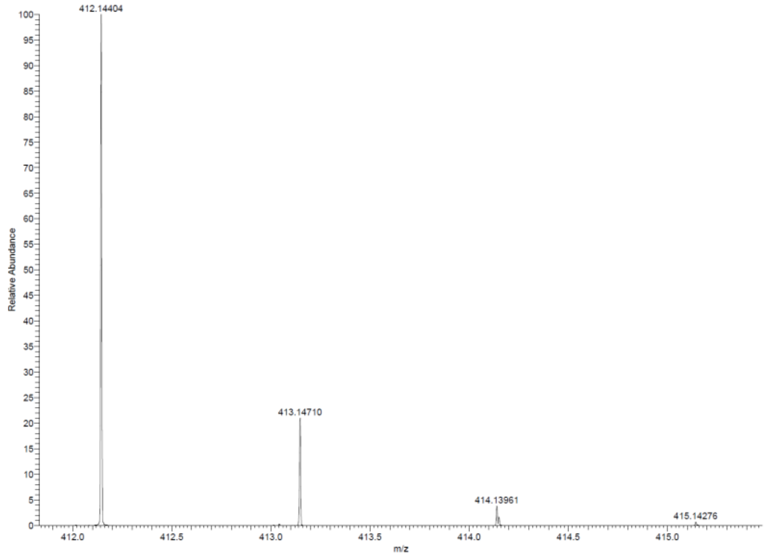
412.14404
413.14710
414.13961
415.14276
Relative Abundance
m/z

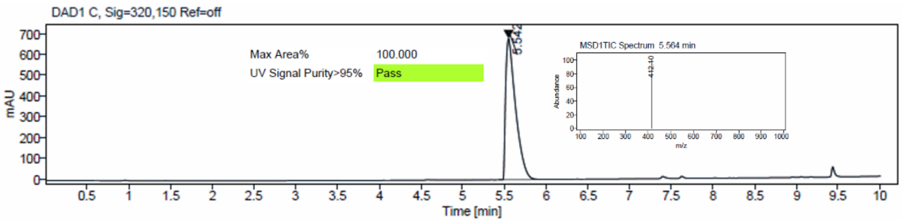
DAD1 C, Sig=320,150 Ref=off
Max Area%
100.000
UV Signal Purity>95%
Pass
MSD1TIC Spectrum 5.564 min
mAU
Time [min]

[1]H and [13]C {H} NMR, HRMS (oTOF) and HPLC/MS (ESI) data of compound **12**

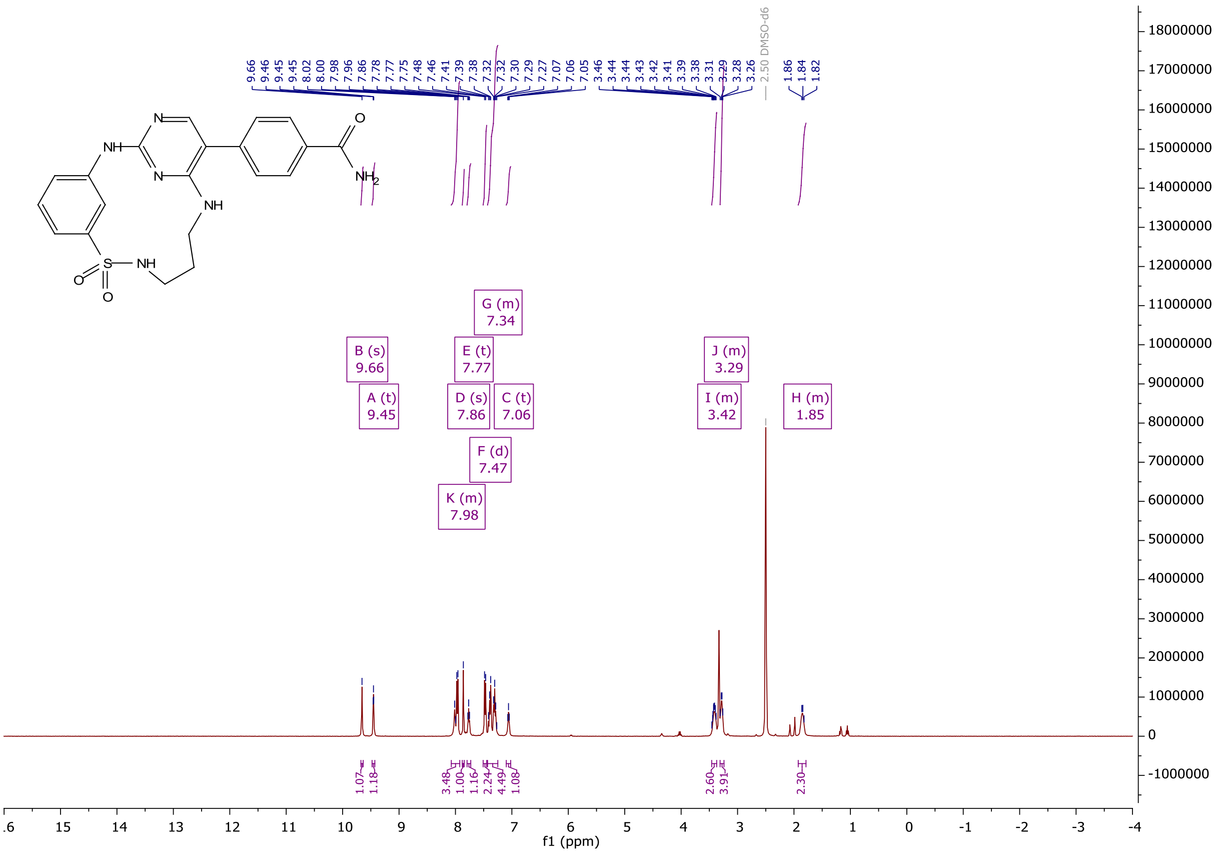

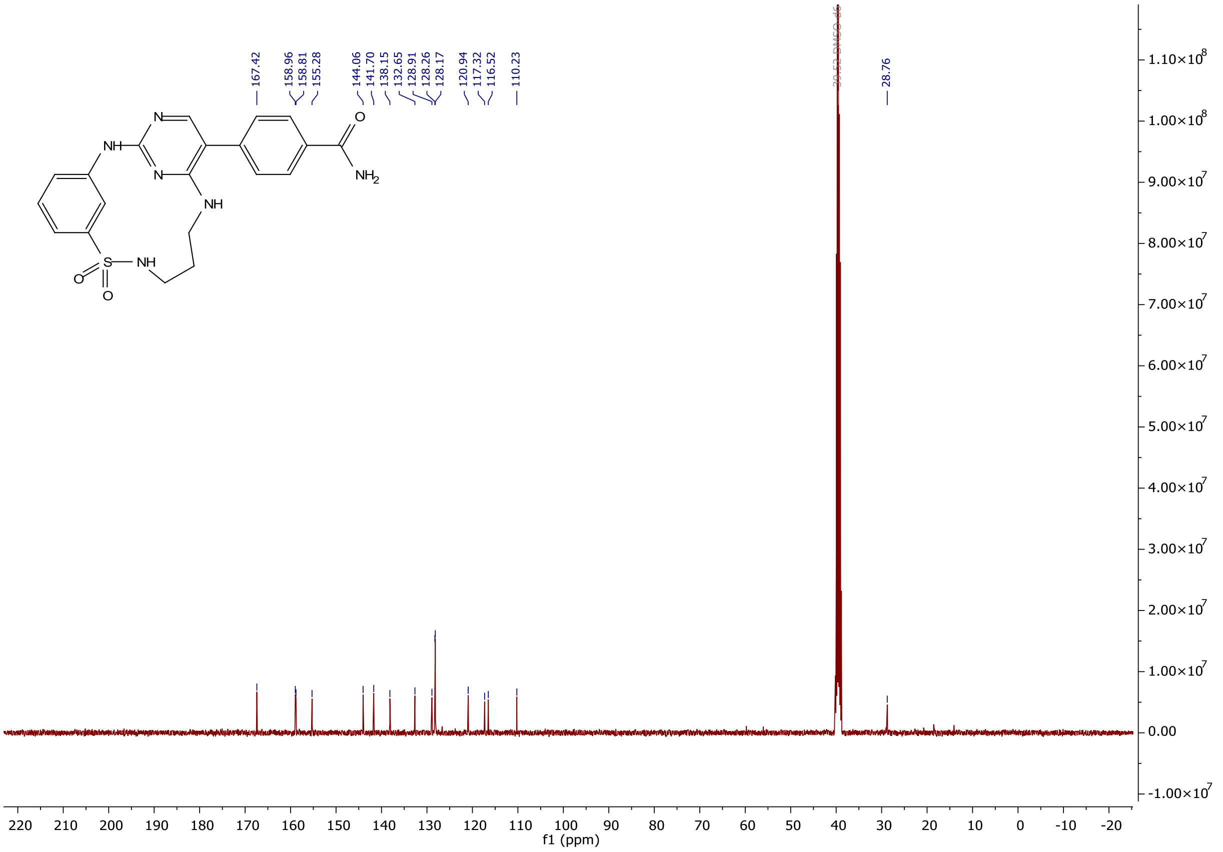

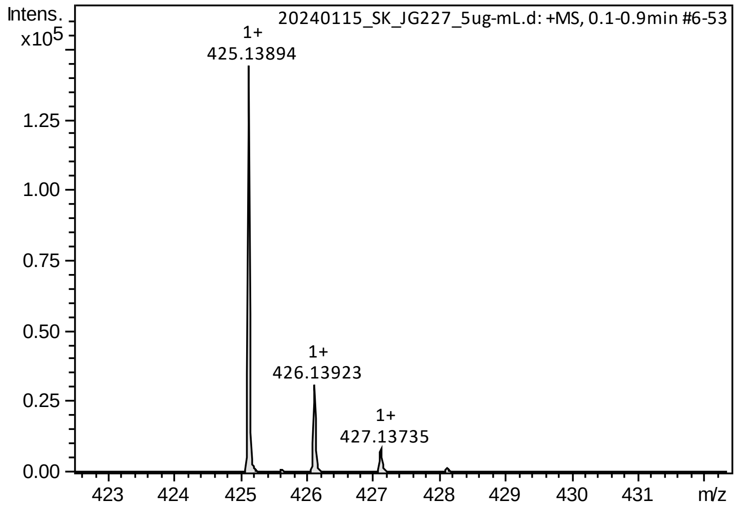

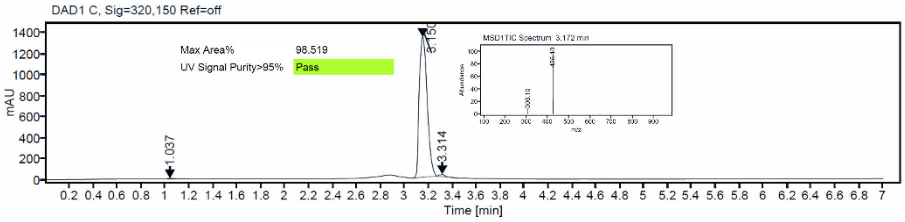

$^{1}$H and $^{13}$C {H} NMR, HRMS (oTOF) and HPLC/MS (ESI) data of compound **13**

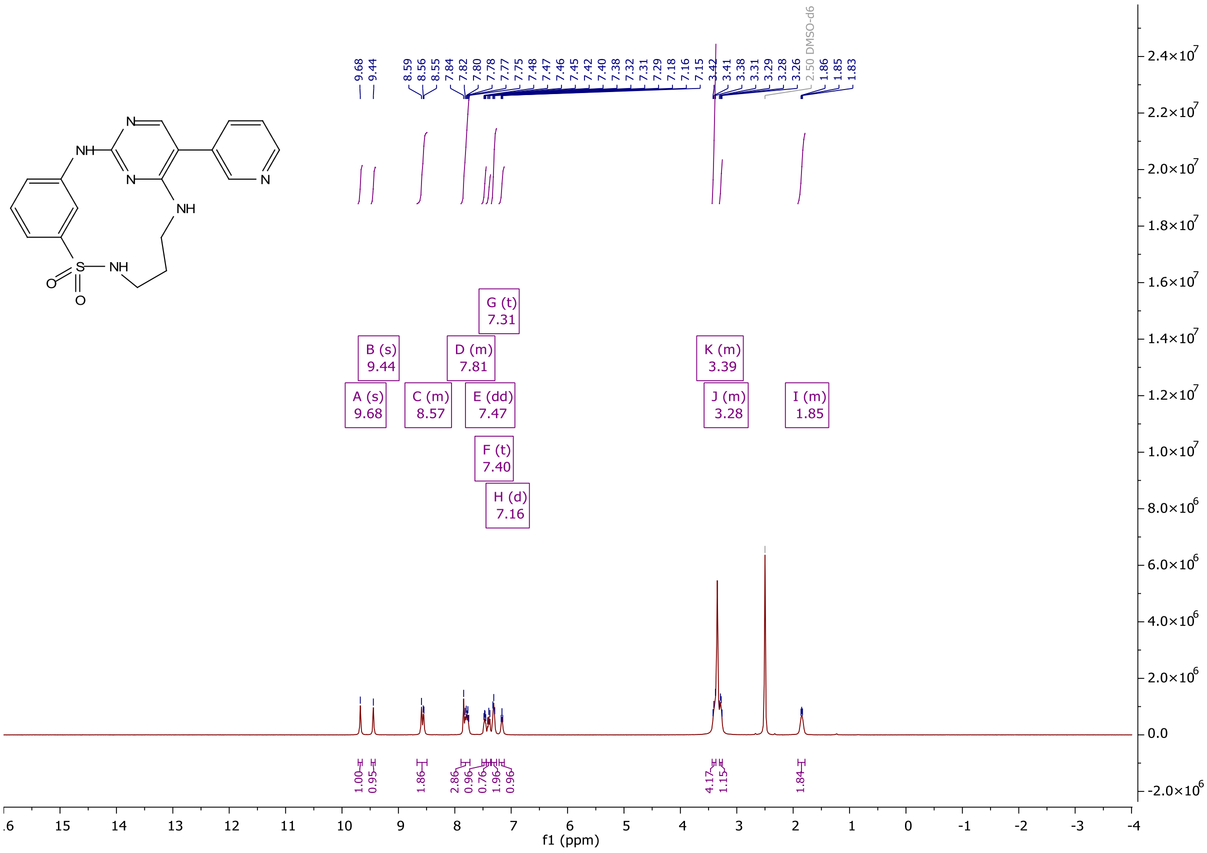

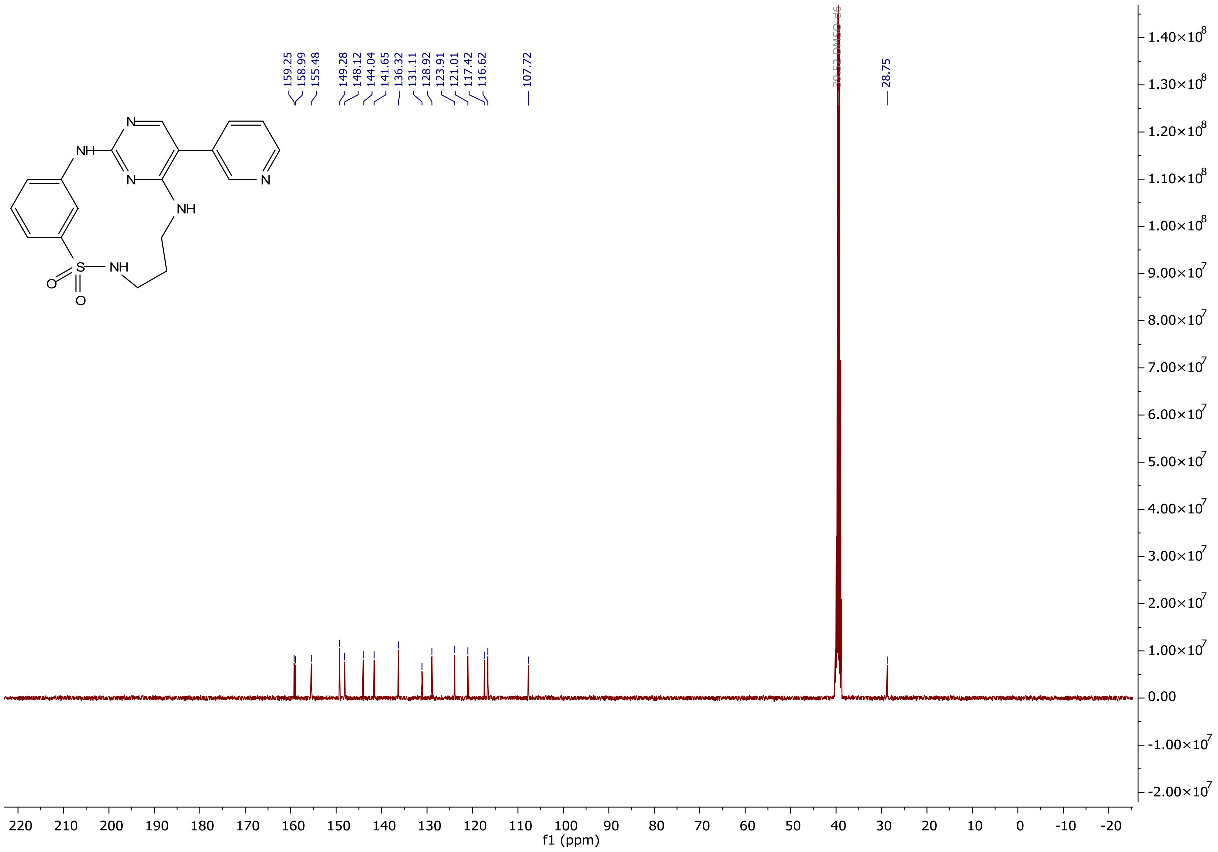
159.25
158.99
155.48
149.28
148.12
144.04
141.65
136.32
131.11
128.92
123.91
121.01
117.42
116.62
107.72
28.75
f1 (ppm)

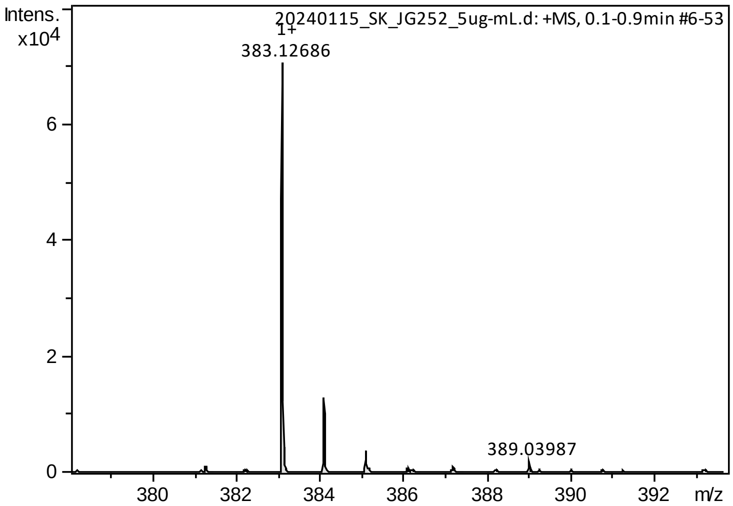
Intens.
x10^4
20240115_SK_JG252_5ug-mL.d: +MS, 0.1-0.9min #6-53
1+
383.12686
389.03987
m/z

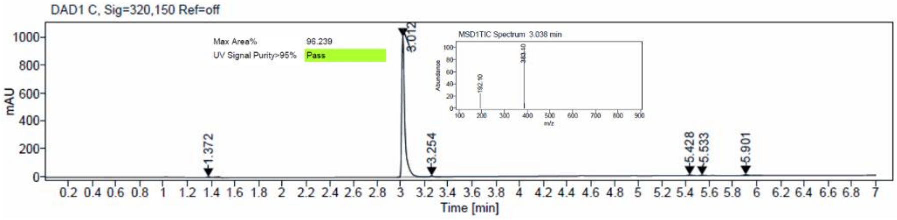
DAD1 C, Sig=320,150 Ref=off
Max Area%
96.239
UV Signal Purity>95%
Pass
MSD1TIC Spectrum 3.038 min
mAU
3.012
1.372
3.254
5.428
5.533
5.901
Time [min]

<sup></sup>

### [1]H and [13]C {H} NMR, HRMS (oTOF) and HPLC/MS (ESI) data of compound **14**

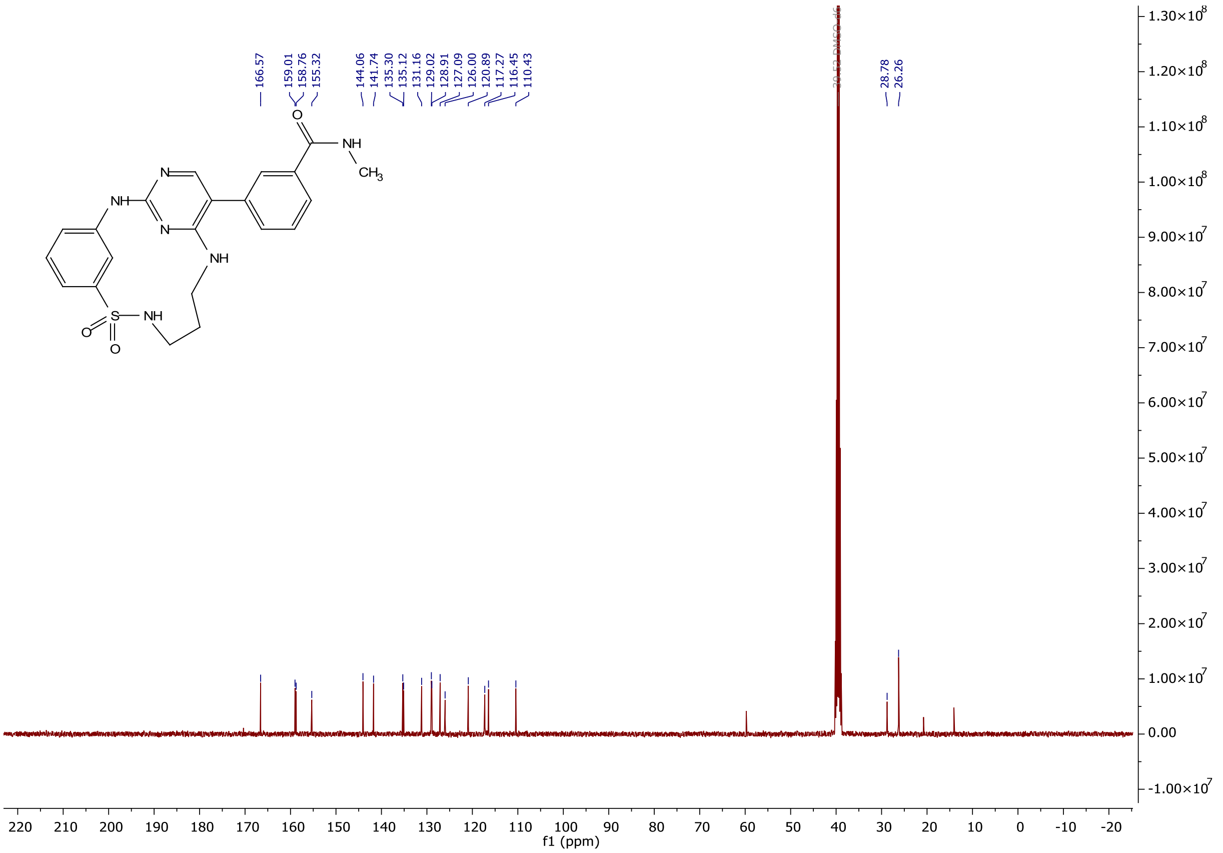

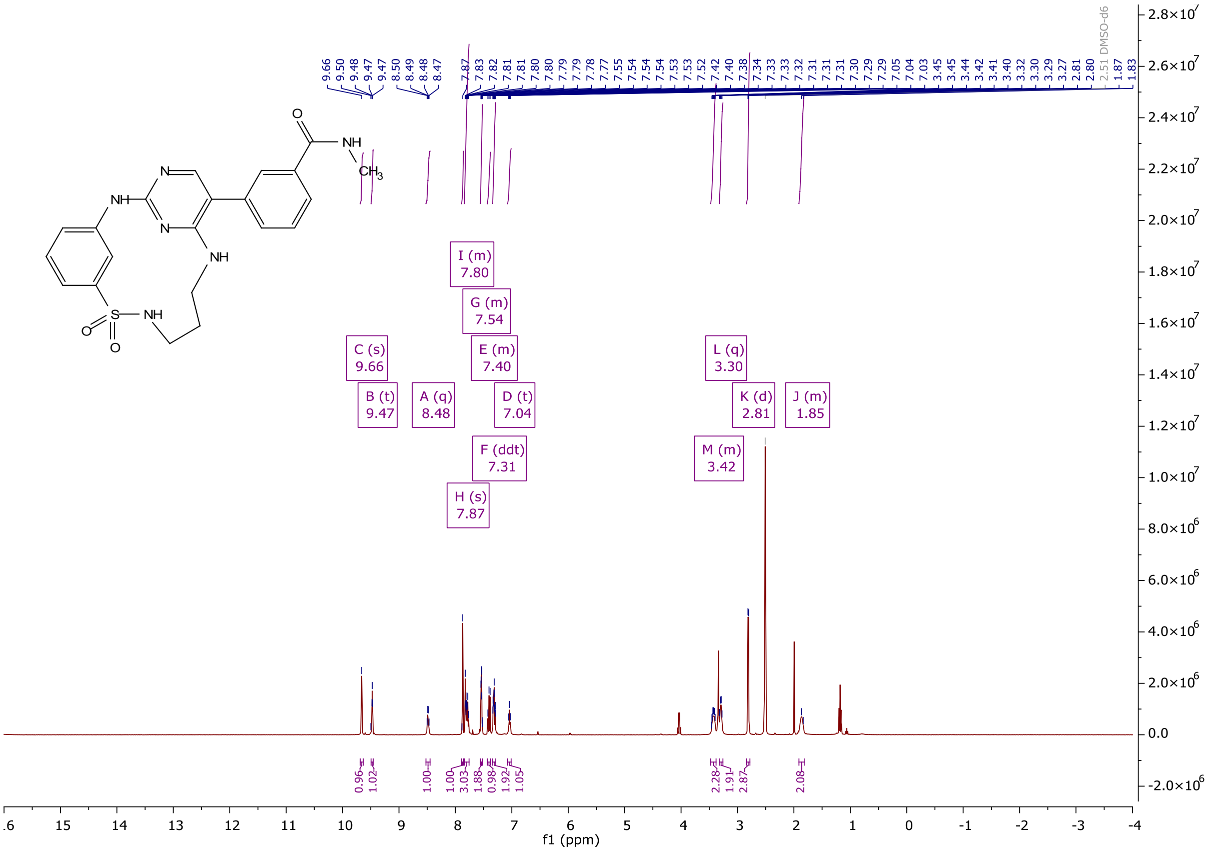

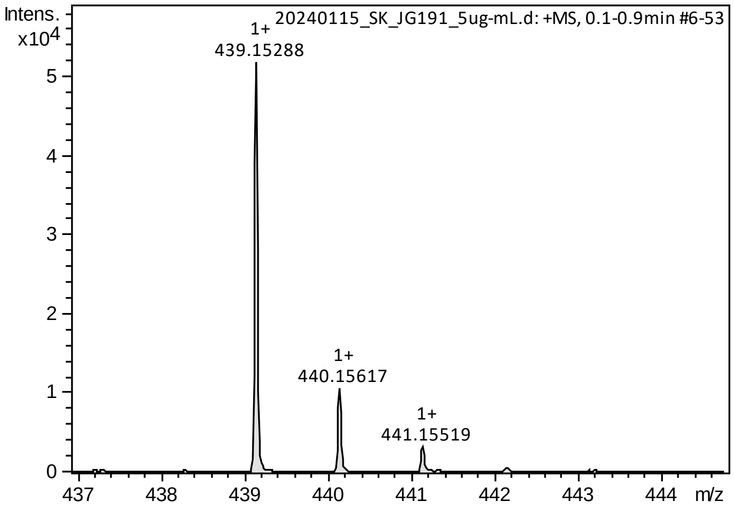

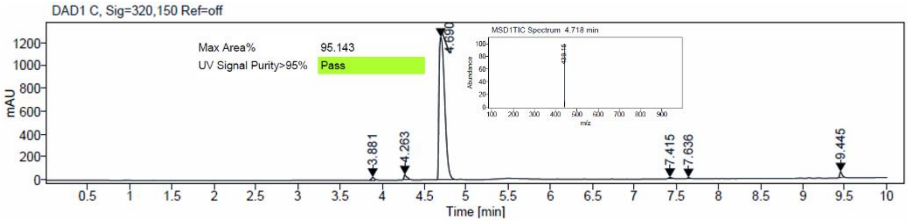

[1]H and [13]C {H} NMR, HRMS (Orbitrap) and HPLC/MS (ESI) data of compound **15**

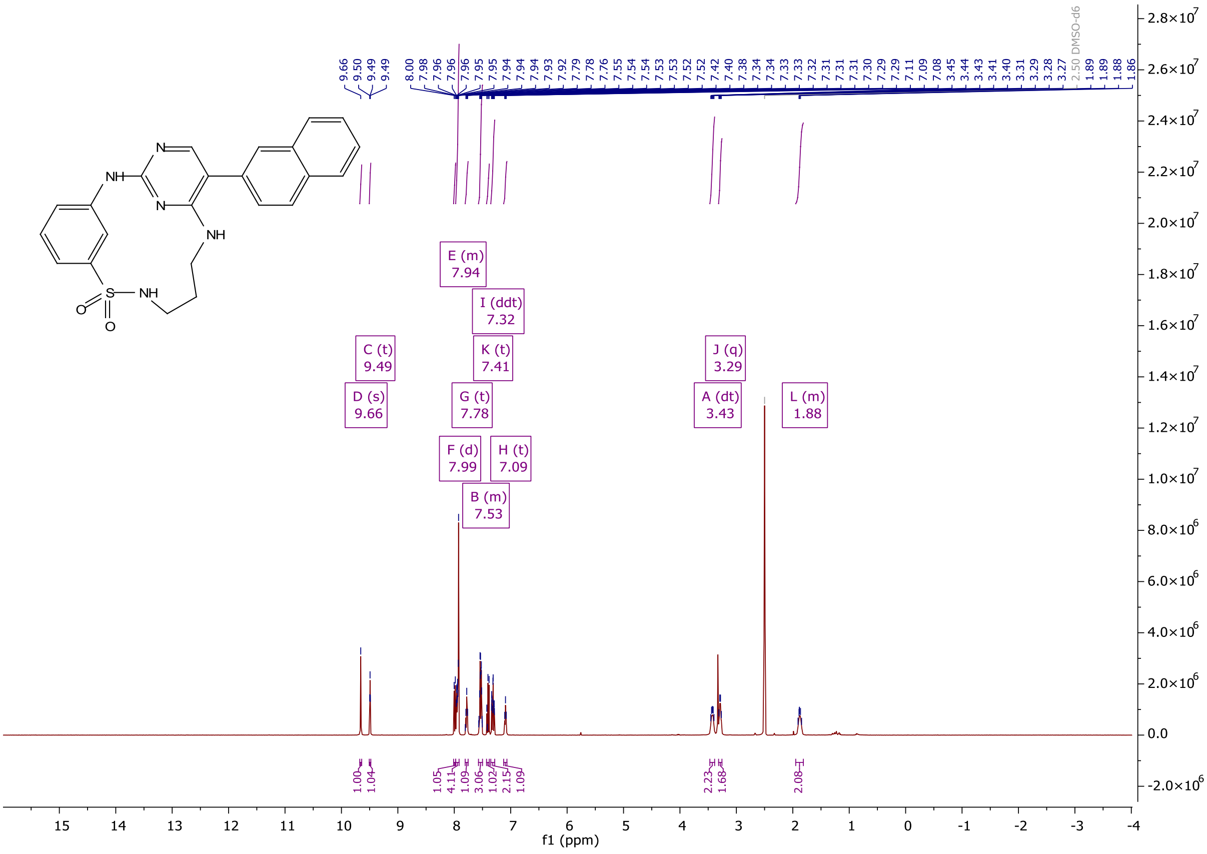

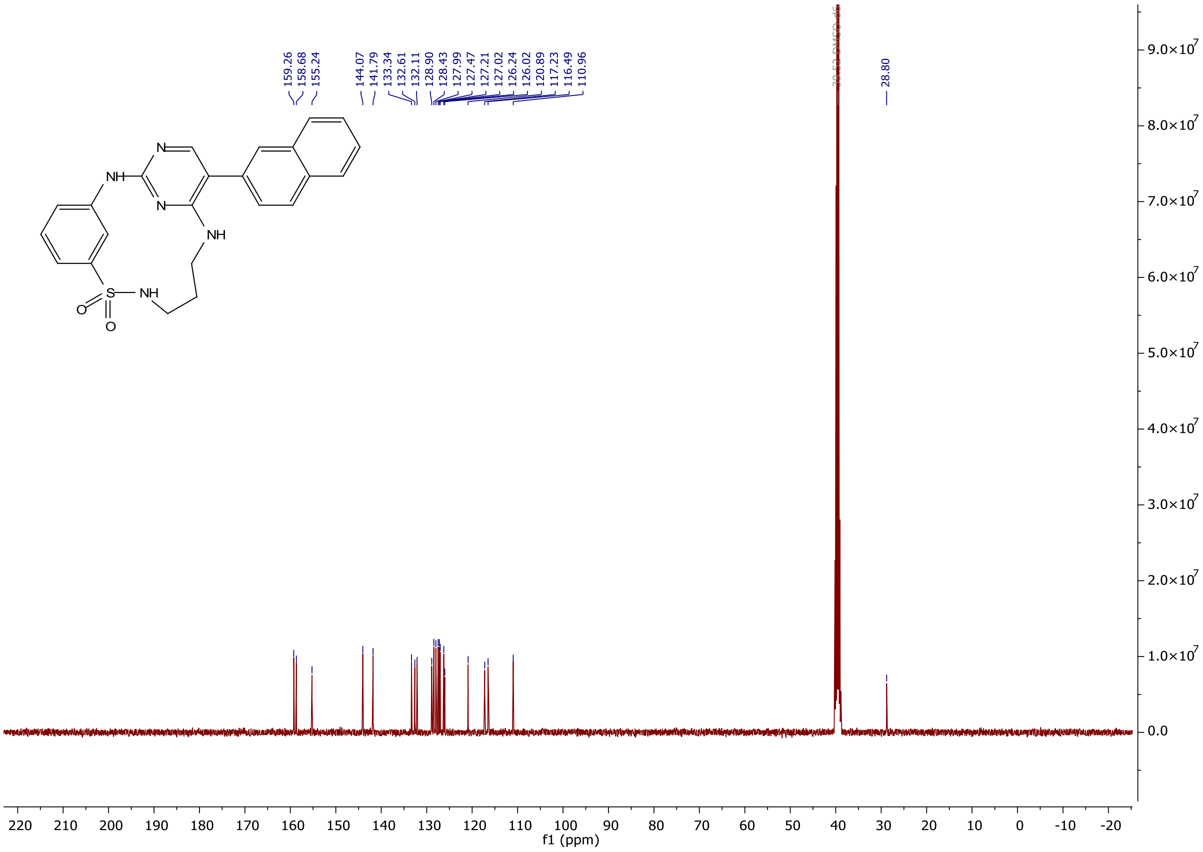

159.26
158.68
155.24
144.07
141.79
133.34
132.61
132.11
128.90
128.43
127.99
127.47
127.21
127.02
126.24
126.02
120.89
117.23
116.49
110.96
28.80
NH
N
N
NH
S
NH
O
O
9.0×10^7
8.0×10^7
7.0×10^7
6.0×10^7
5.0×10^7
4.0×10^7
3.0×10^7
2.0×10^7
1.0×10^7
0.0
f1 (ppm)

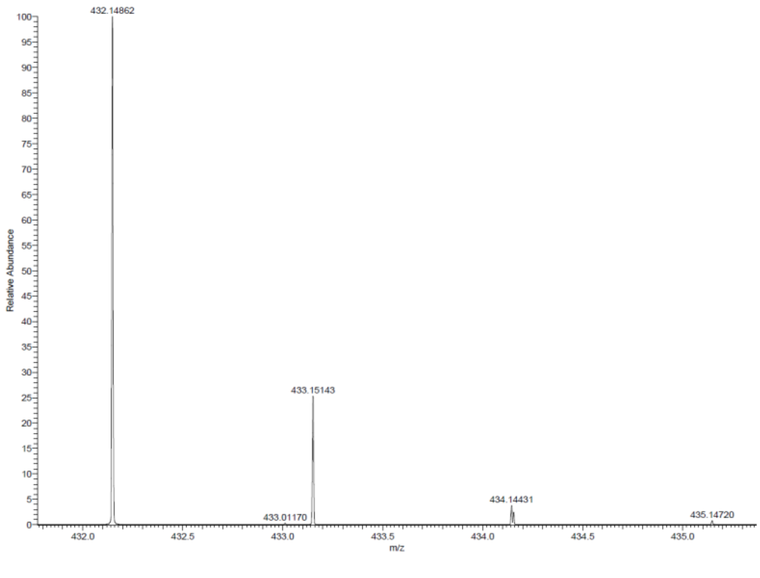

432.14862
433.15143
433.01170
434.14431
435.14720
Relative Abundance
m/z

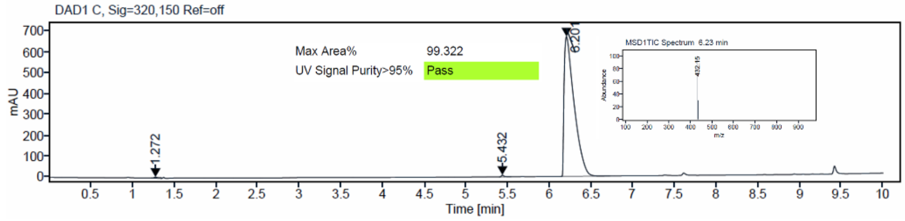

DAD1 C, Sig=320,150 Ref=off
Max Area%
99.322
UV Signal Purity>95%
Pass
mAU
1.272
5.432
6.201
Time [min]

$^{1}H$ and $^{13}C$ {H} NMR, HRMS (Orbitrap) and HPLC/MS (ESI) data of compound **16**

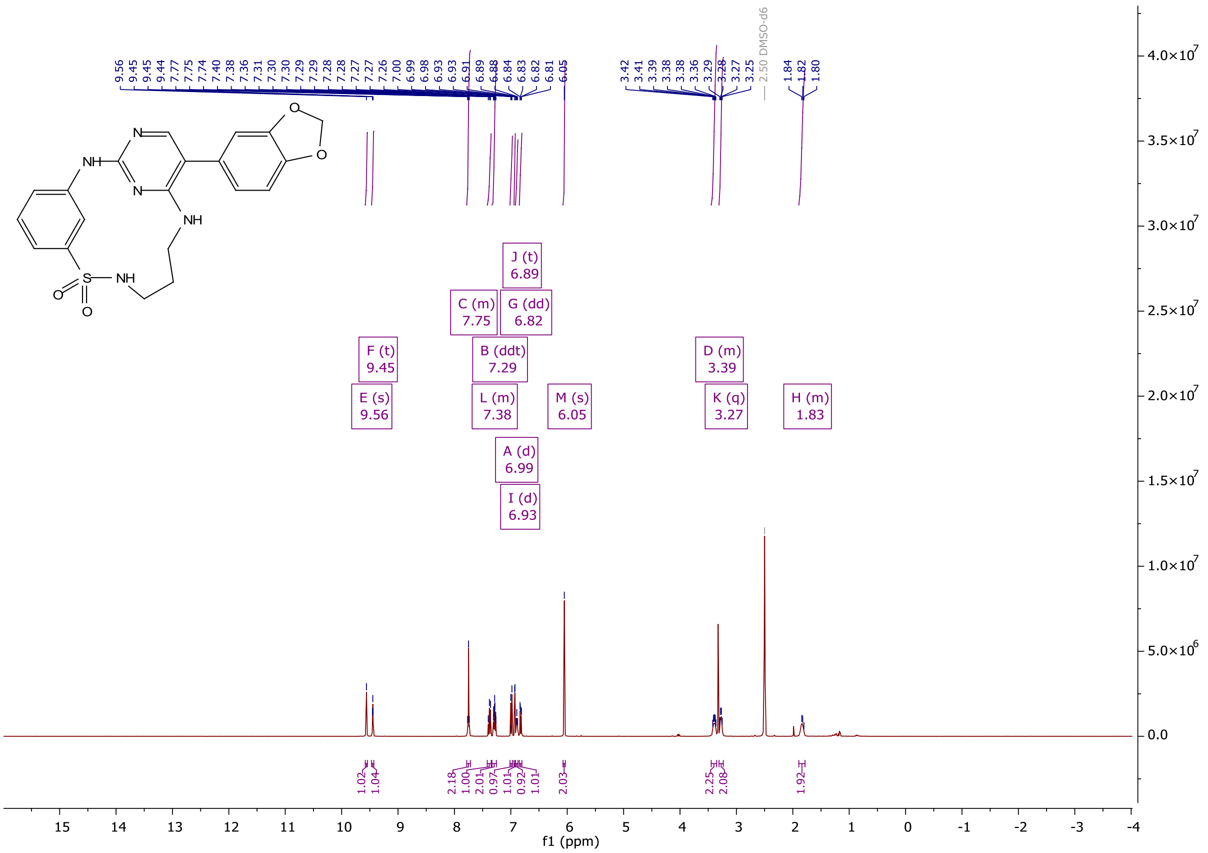

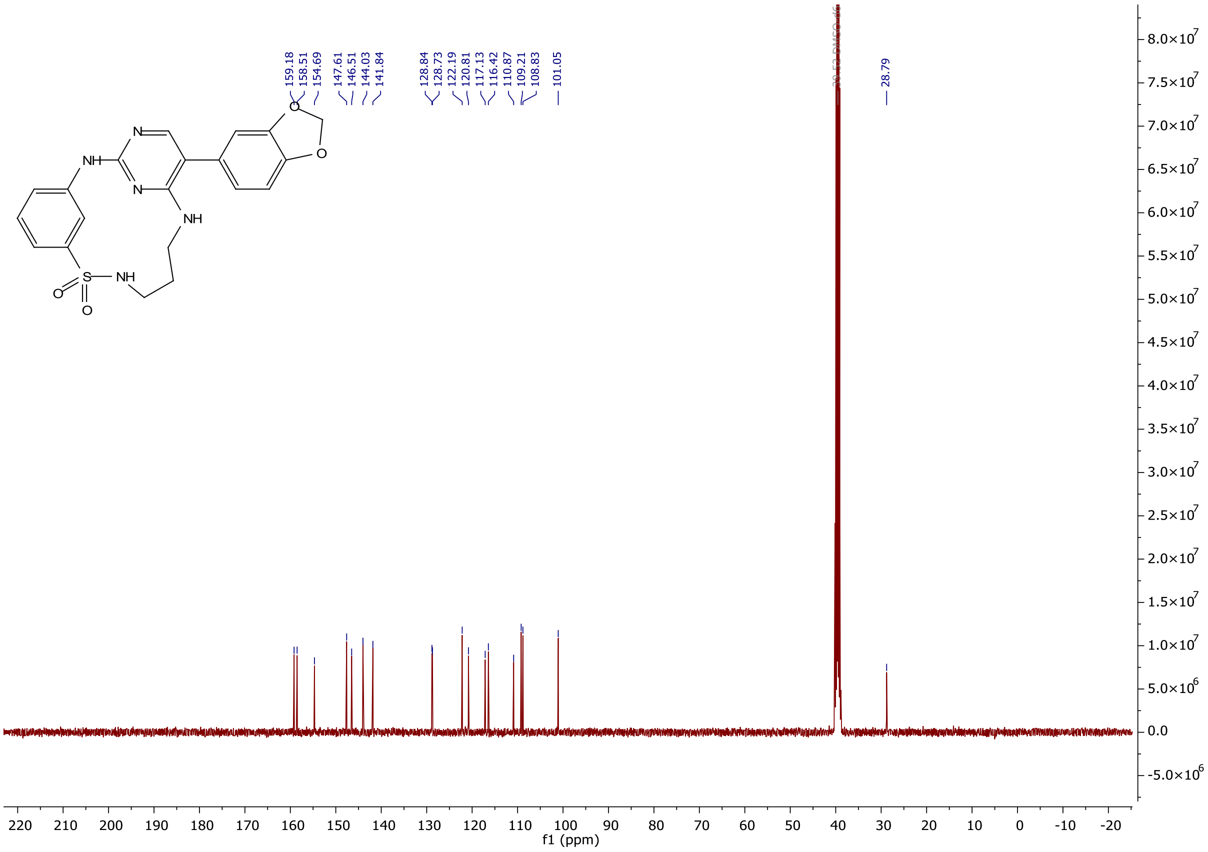

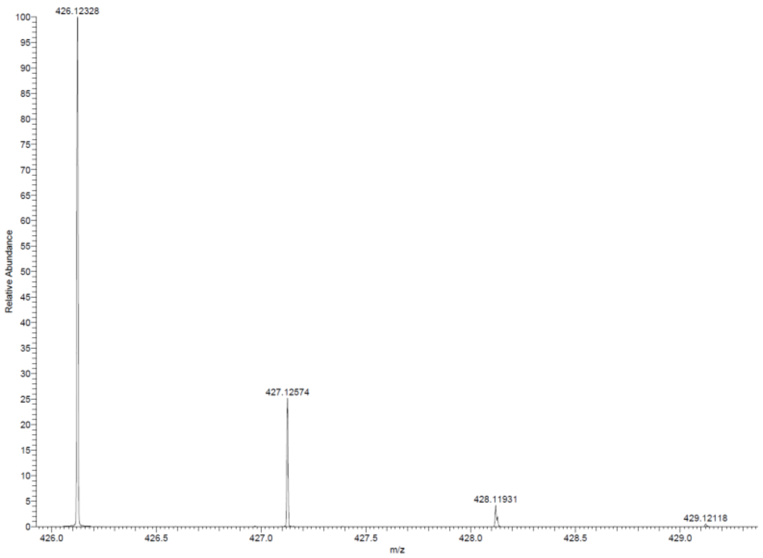

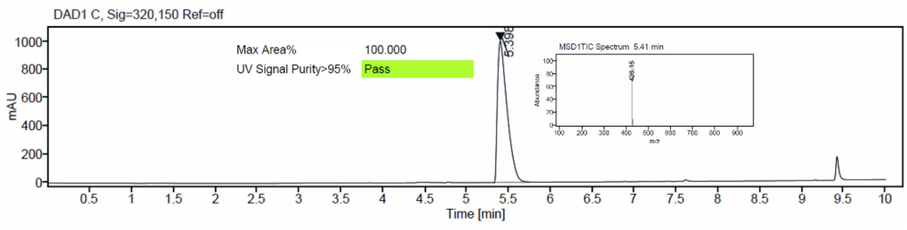

$^{1}H$ and $^{13}C$ {H} NMR, HRMS (Orbitrap) and HPLC/MS (ESI) data of compound **17**

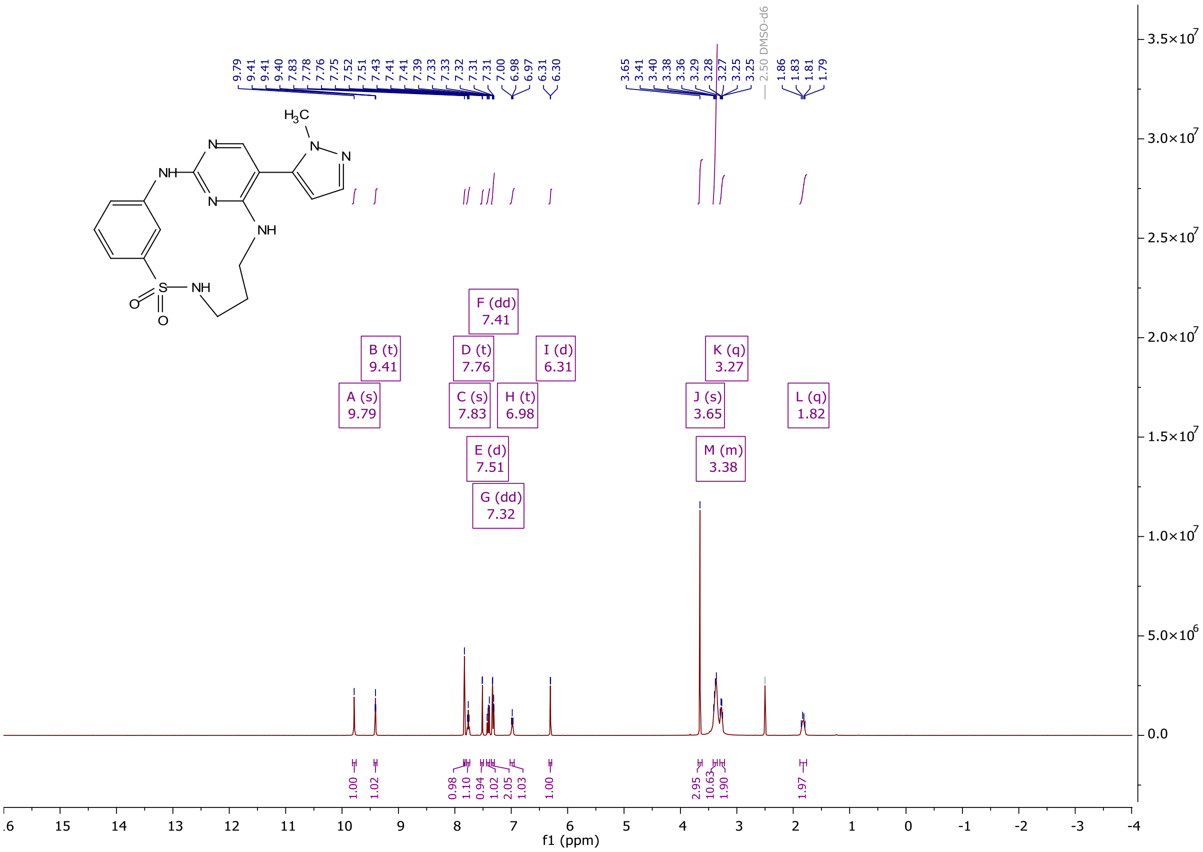

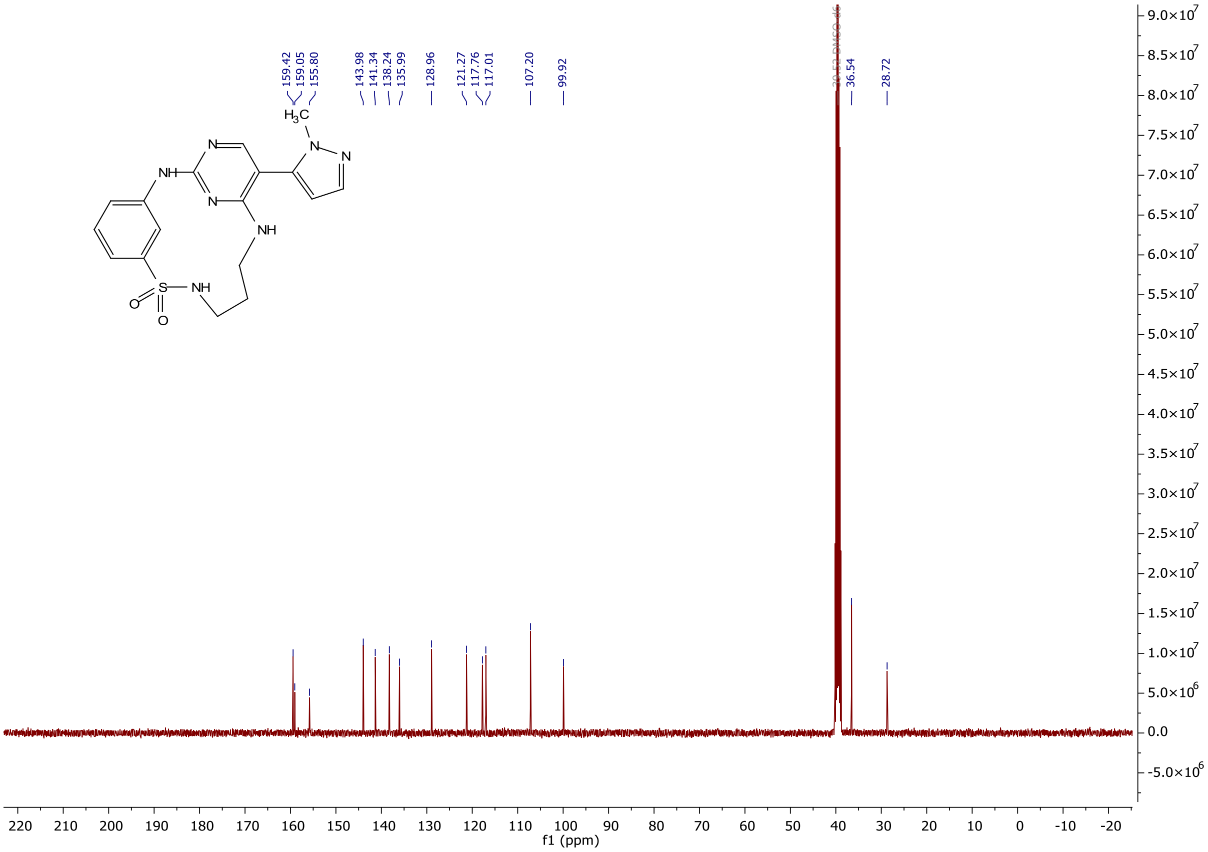
159.42
159.05
155.80
143.98
141.34
138.24
135.99
128.96
121.27
117.76
117.01
107.20
99.92
36.54
28.72
f1 (ppm)

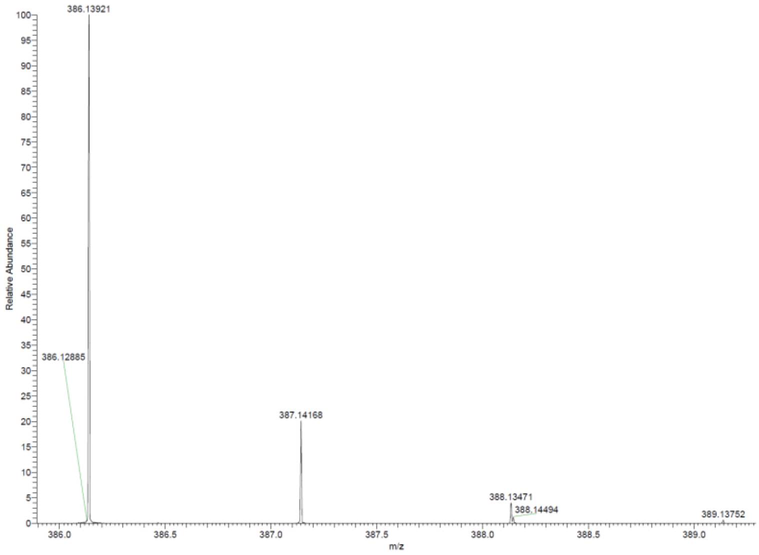
386.13921
386.12885
387.14168
388.13471
388.14494
389.13752
Relative Abundance
m/z

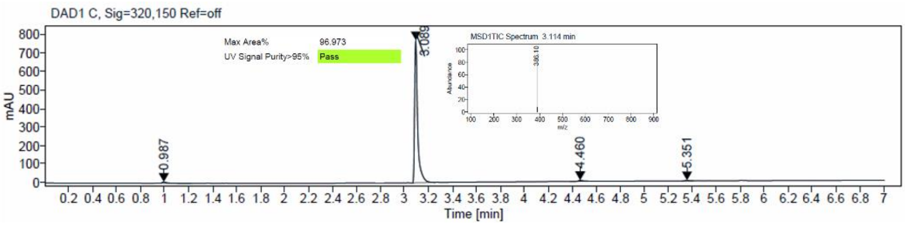
DAD1 C, Sig=320,150 Ref=off
Max Area% 96.973
UV Signal Purity>95% Pass
MSD1TIC Spectrum 3.114 min
3.089
0.987
4.460
5.351
mAU
Time [min]

$^{1}$H and $^{13}$C {H} NMR, HRMS (Orbitrap) and HPLC/MS (ESI) data of compound **18**

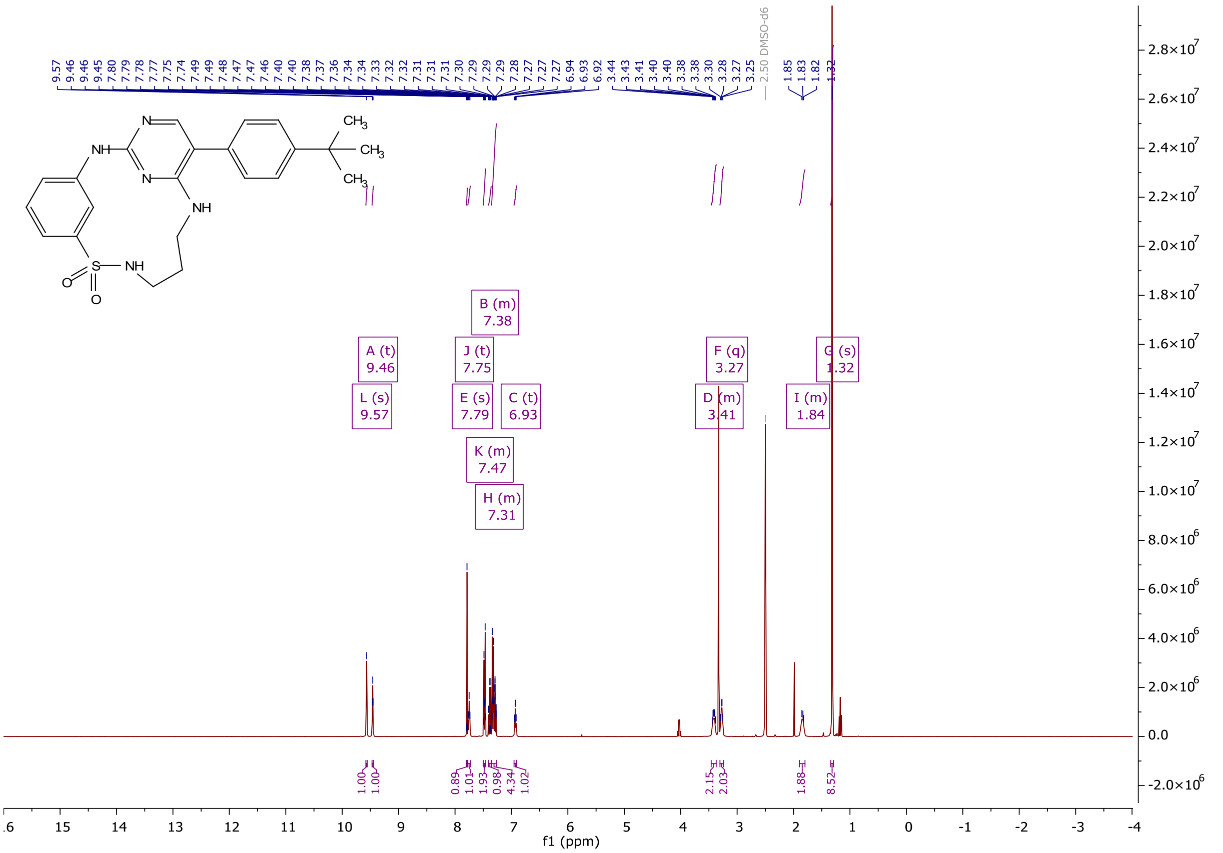

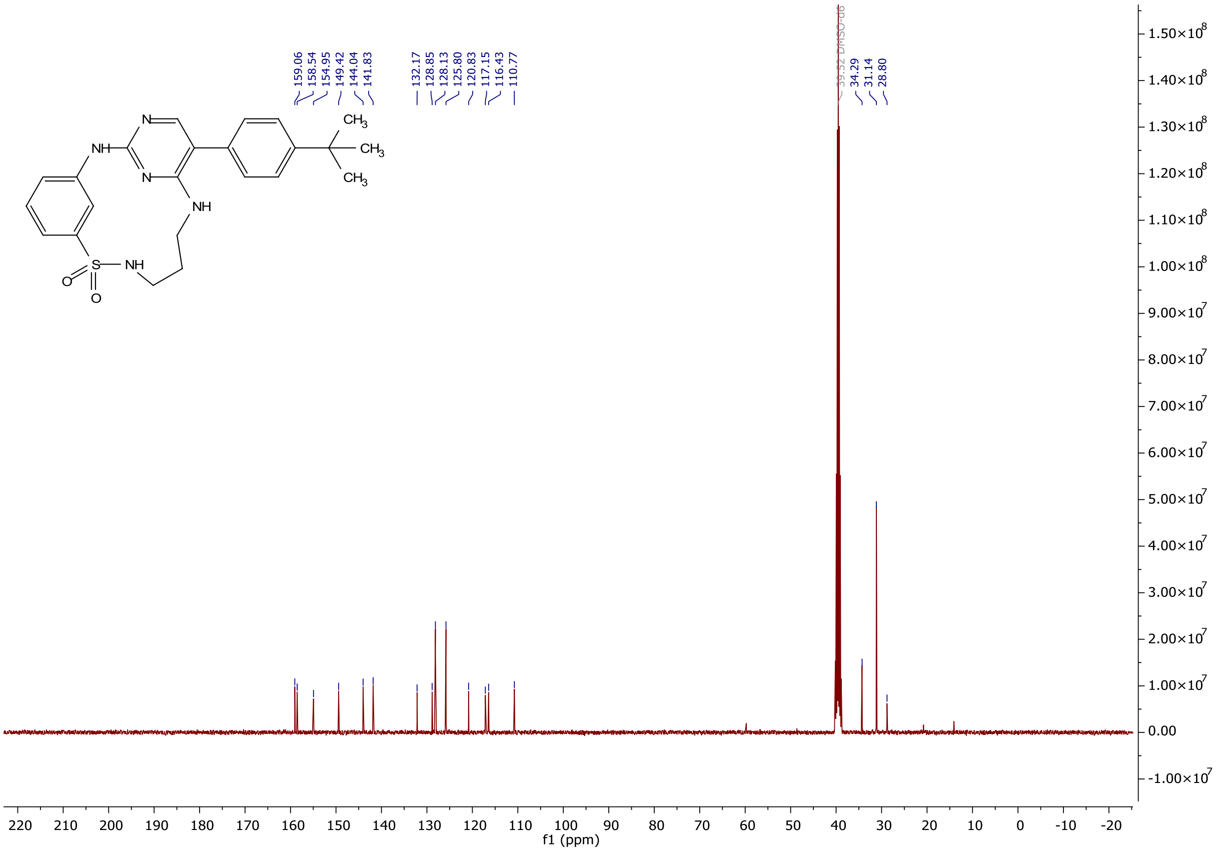

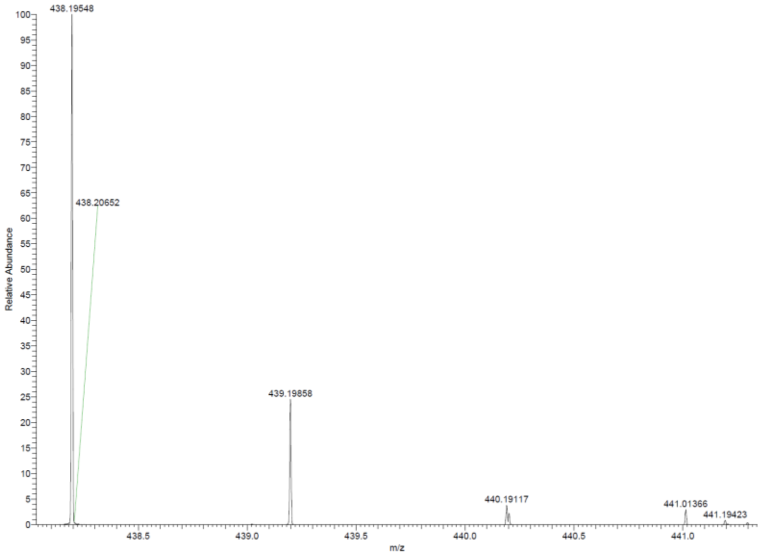

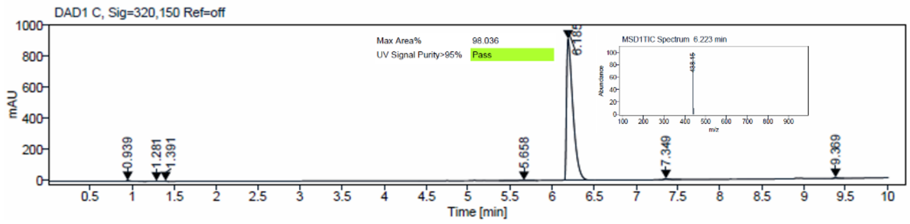

$^1$H and $^{13}$C {H} NMR, HRMS (Orbitrap) and HPLC/MS (ESI) data of compound **19**

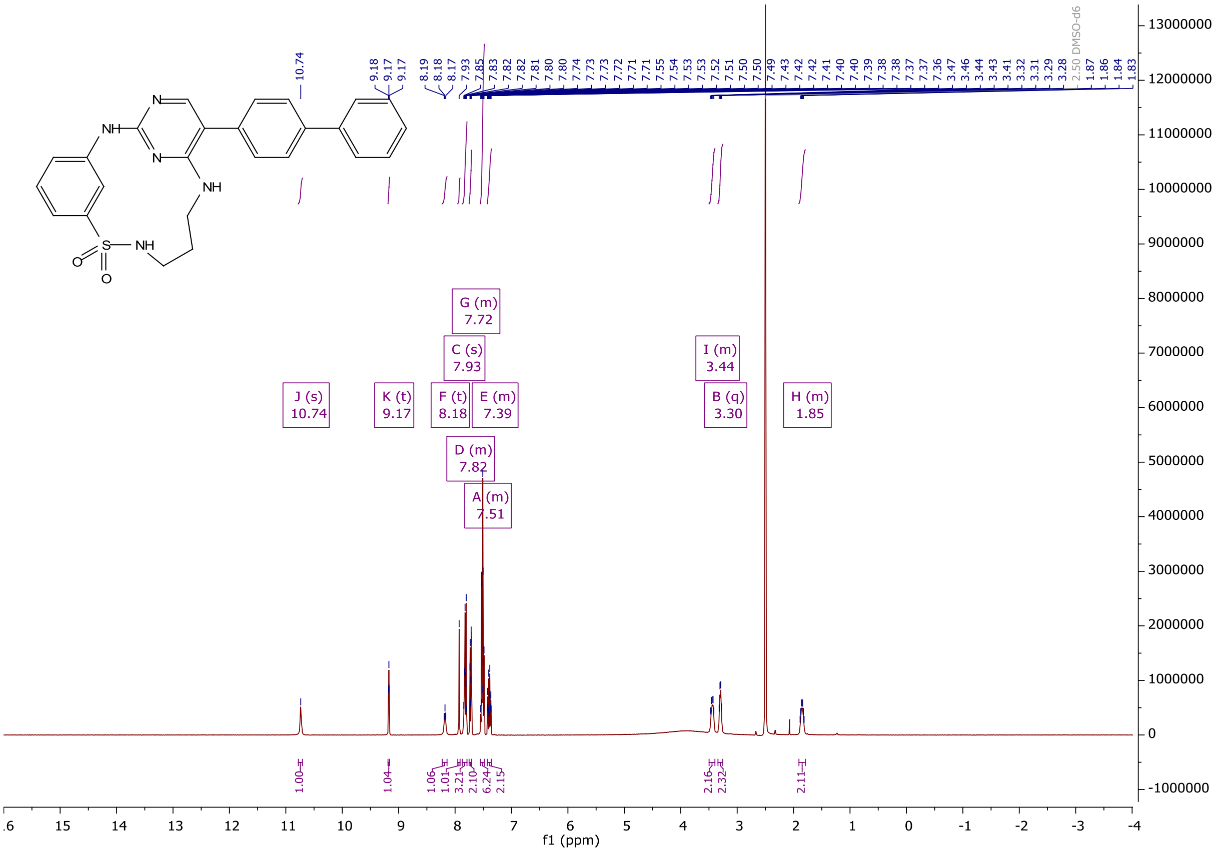

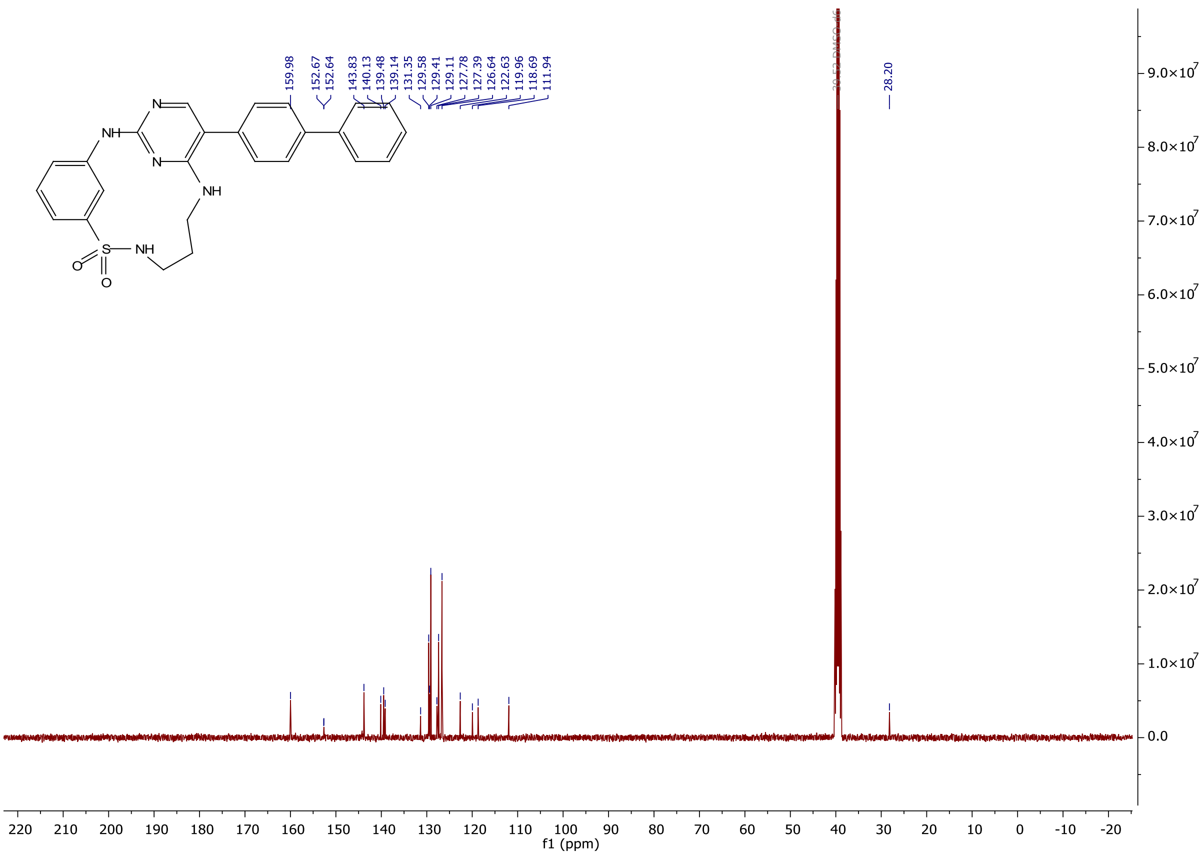

159.98
152.67
152.64
143.83
140.13
139.48
139.14
131.35
129.58
129.41
129.11
127.78
127.39
126.64
122.63
119.96
118.69
111.94
28.20
f1 (ppm)

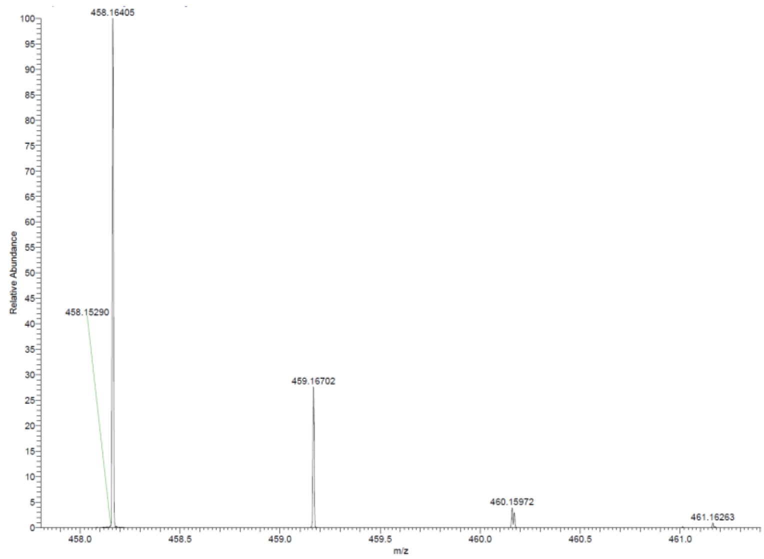

458.16405
458.15290
459.16702
460.15972
461.16263
Relative Abundance
m/z

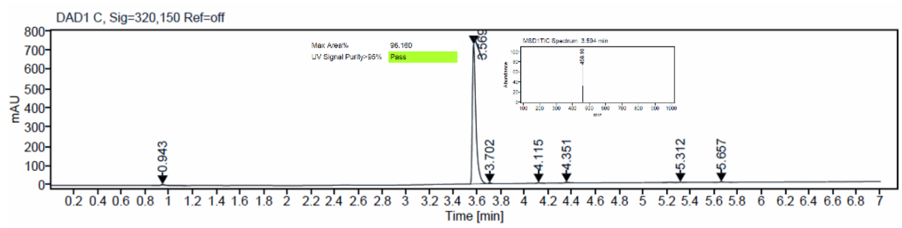

DAD1 C, Sig=320,150 Ref=off
Max Area% 96.160
UV Signal Purity>95% Pass
mAU
3.569
0.943
3.702
4.115
4.351
5.312
5.657
Time [min]

[1]H and [13]C {H} NMR, HRMS (Orbitrap) and HPLC/MS (ESI) data of compound **20**

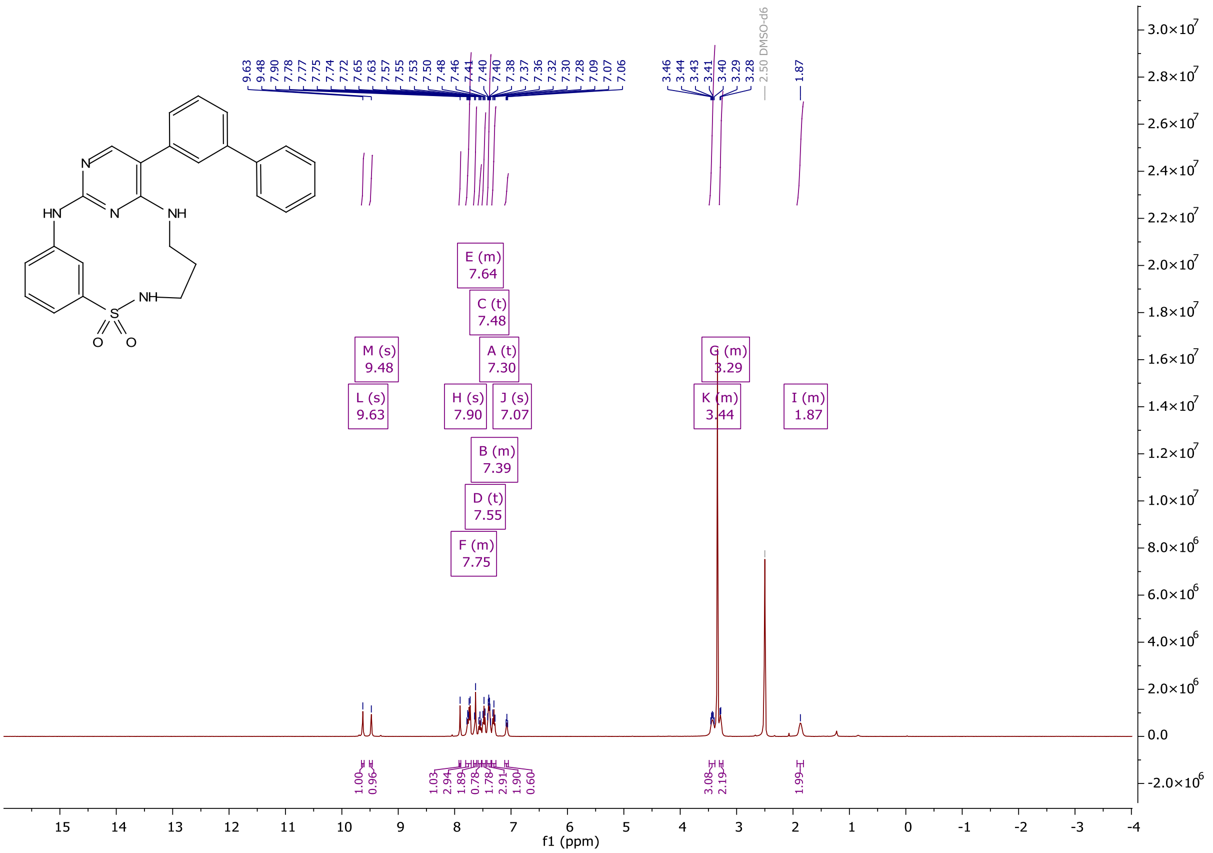

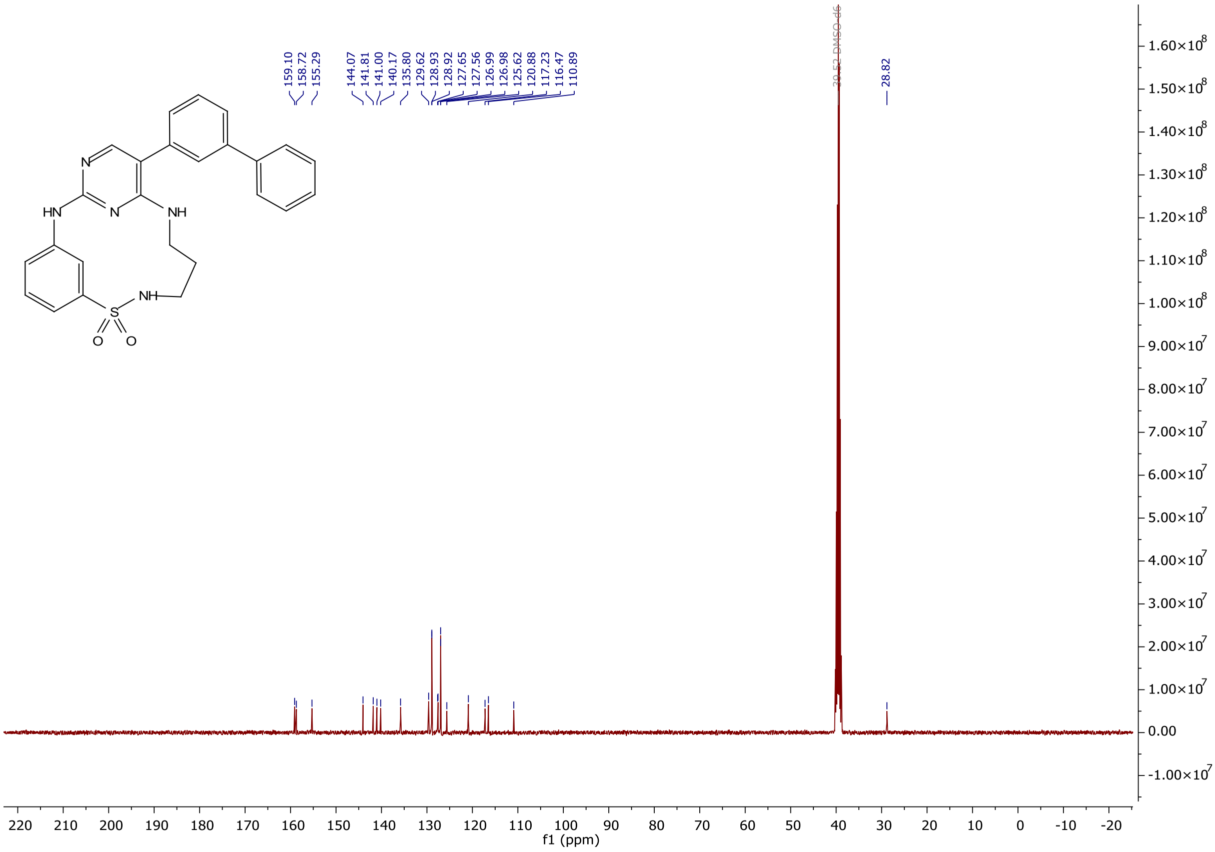

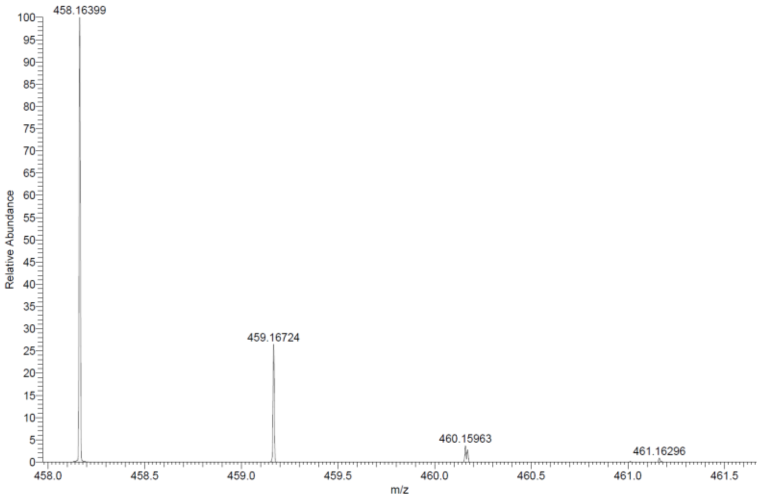

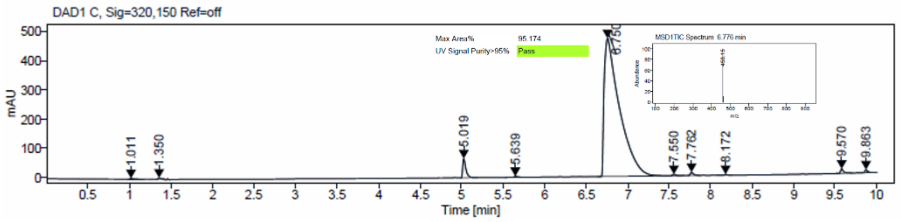

$^{1}H$ and $^{13}C$ {H} NMR, HRMS (Orbitrap) and HPLC/MS (ESI) data of compound **21**

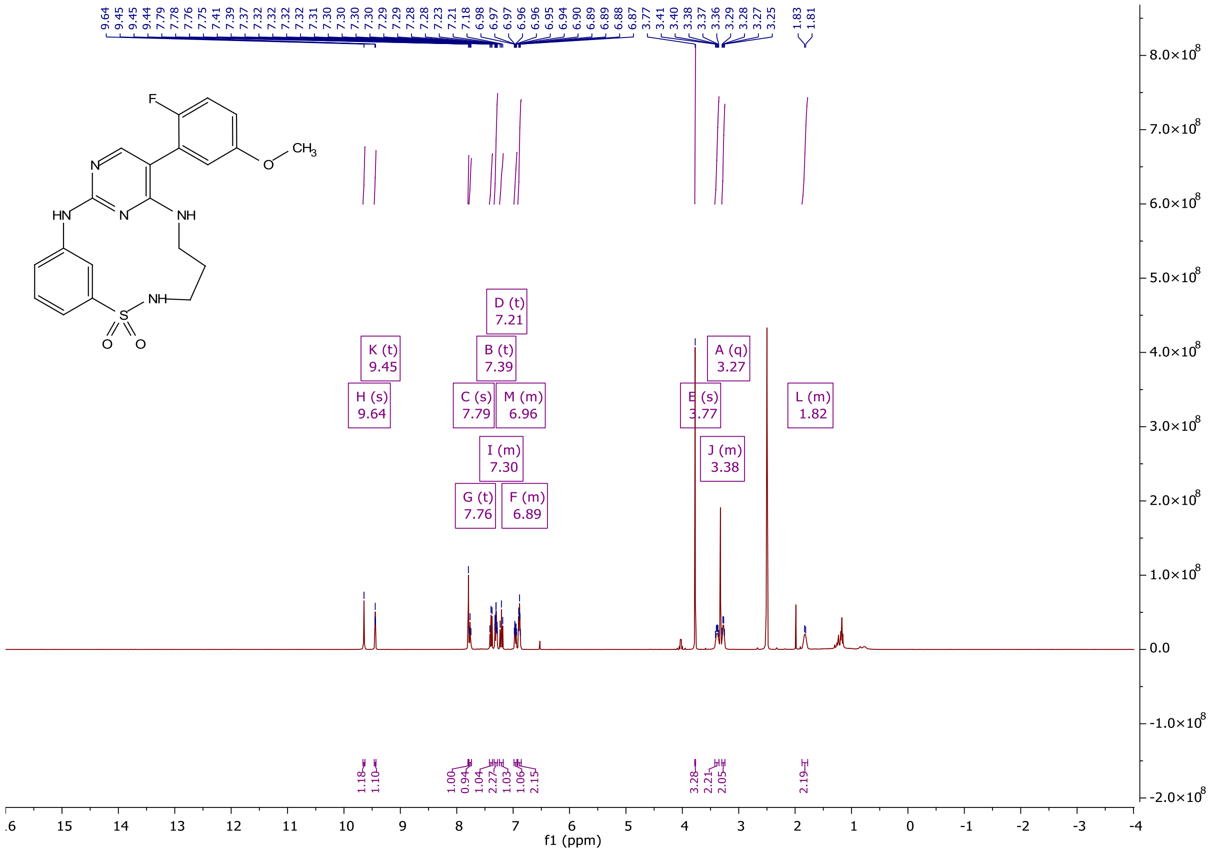

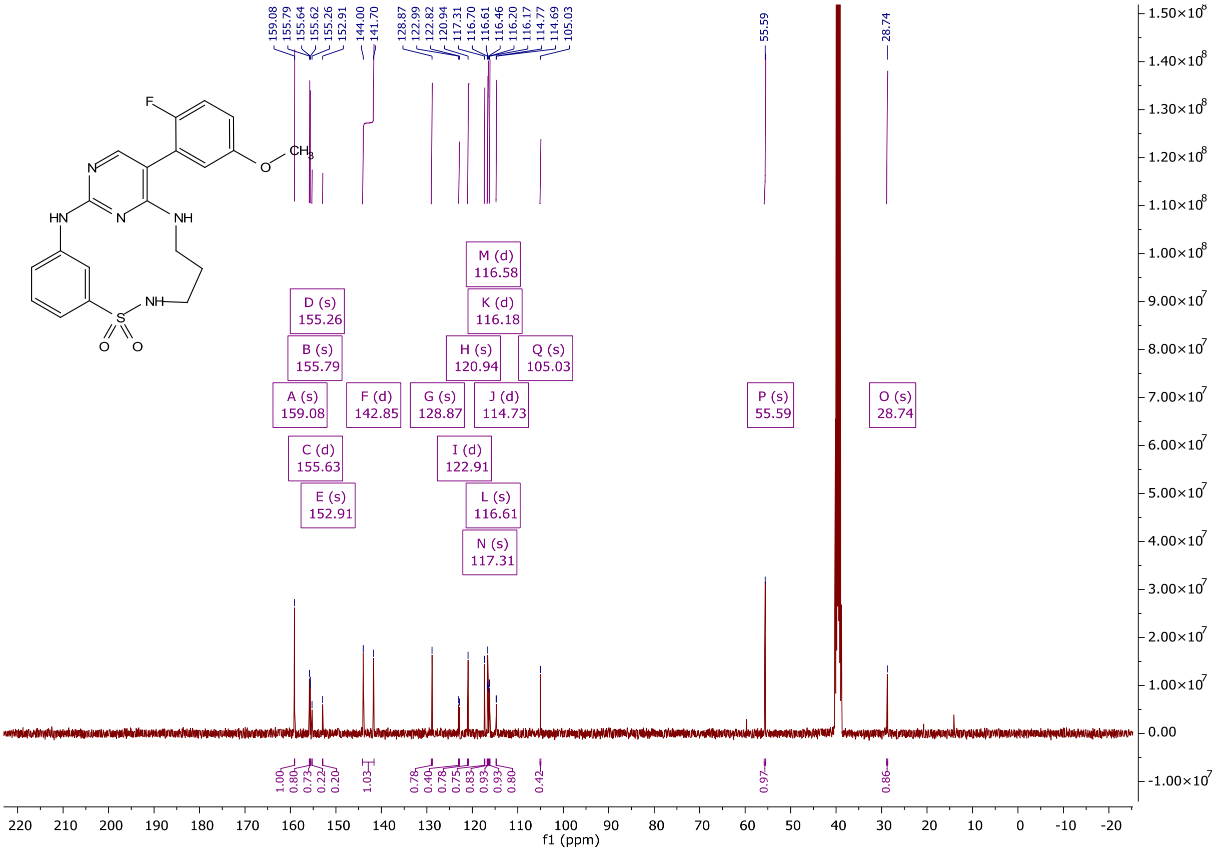

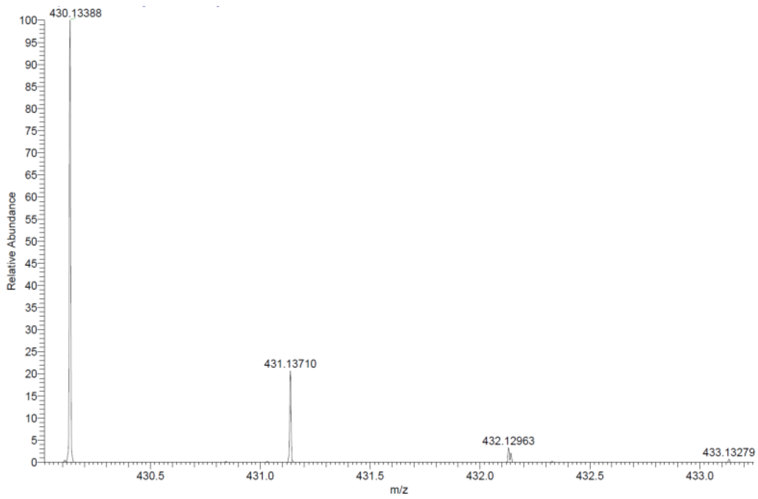

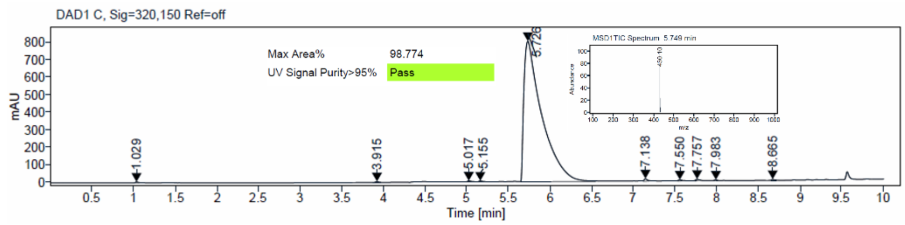

### [1]H and [13]C {H} NMR, HRMS (Orbitrap) and HPLC/MS (ESI) data of compound **22**

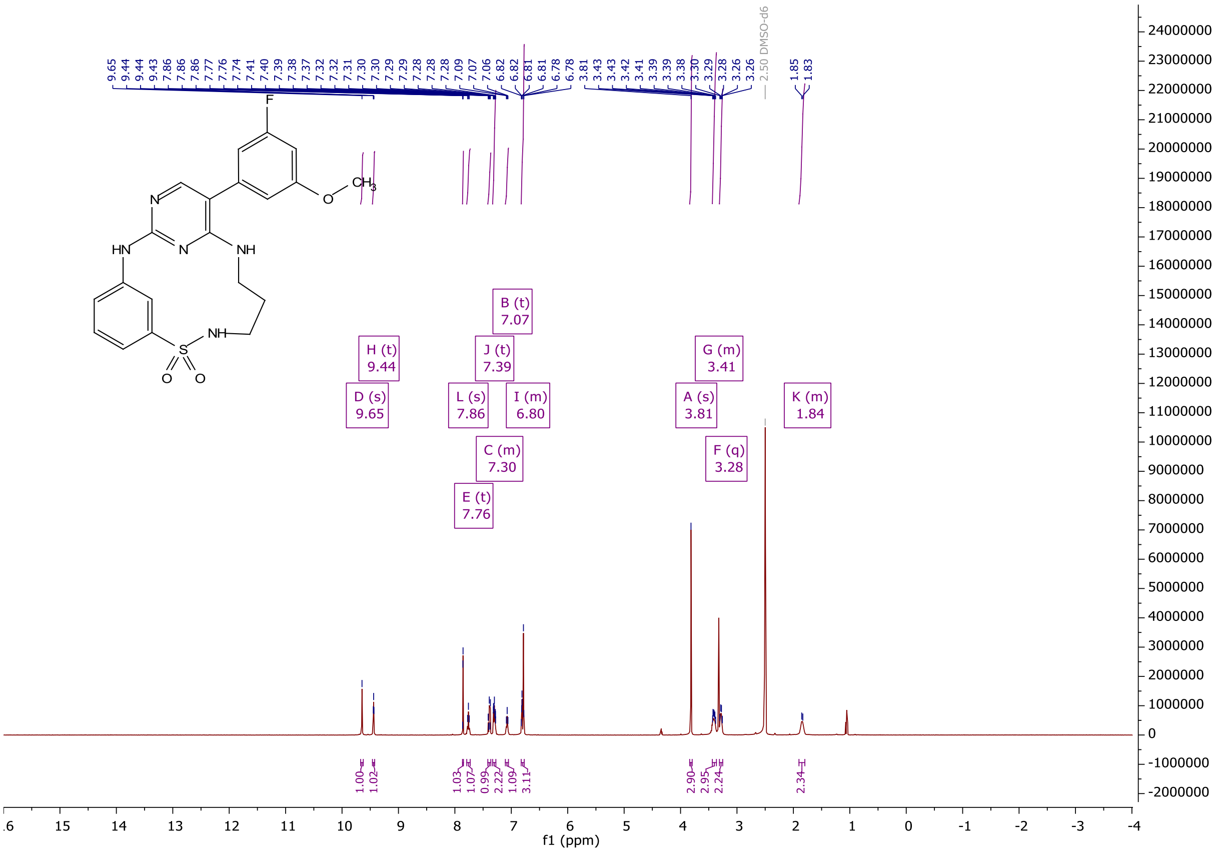

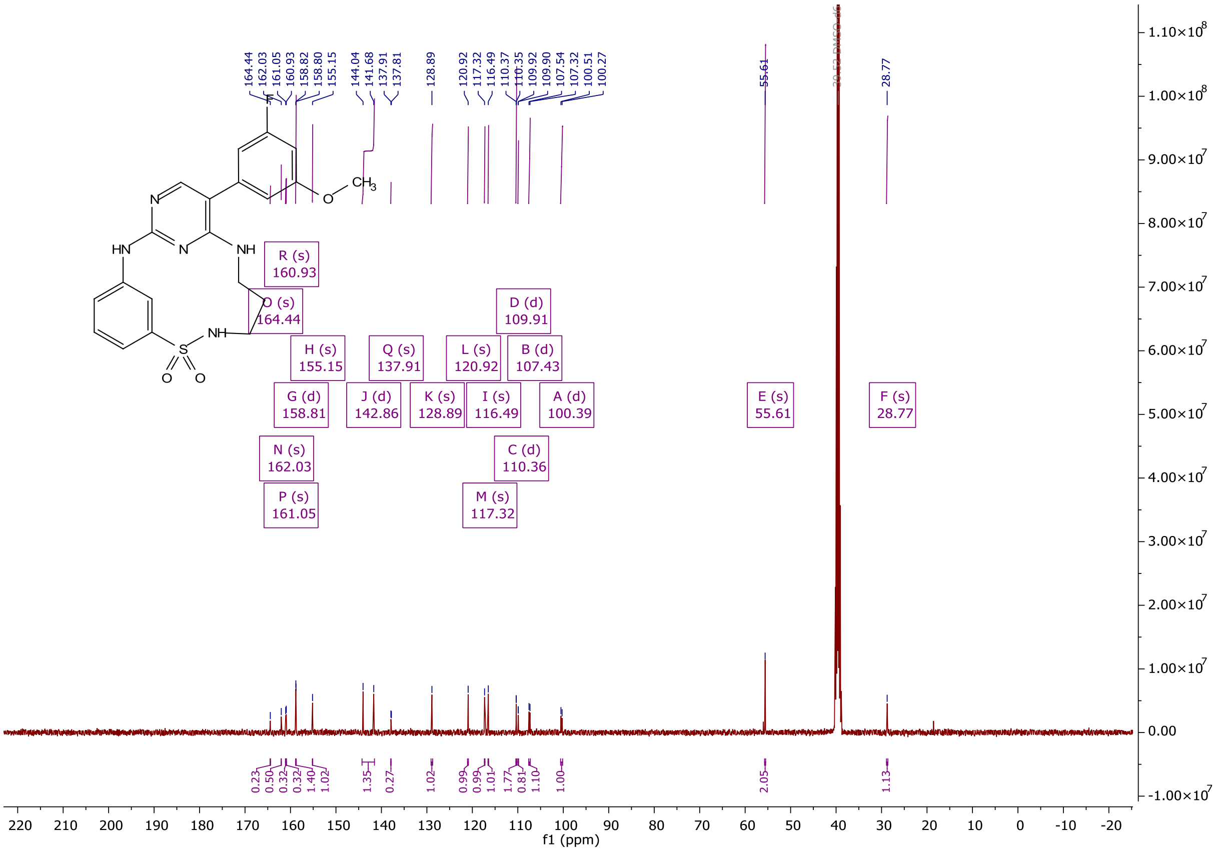

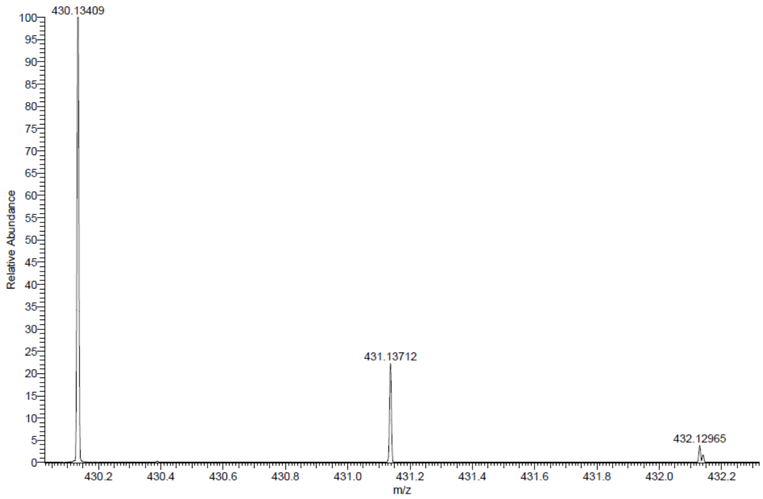

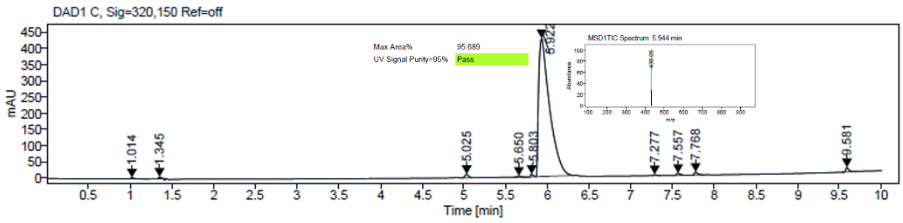

[1]H and [13]C {H} NMR, HRMS (Orbitrap) and HPLC/MS (ESI) data of compound **23**

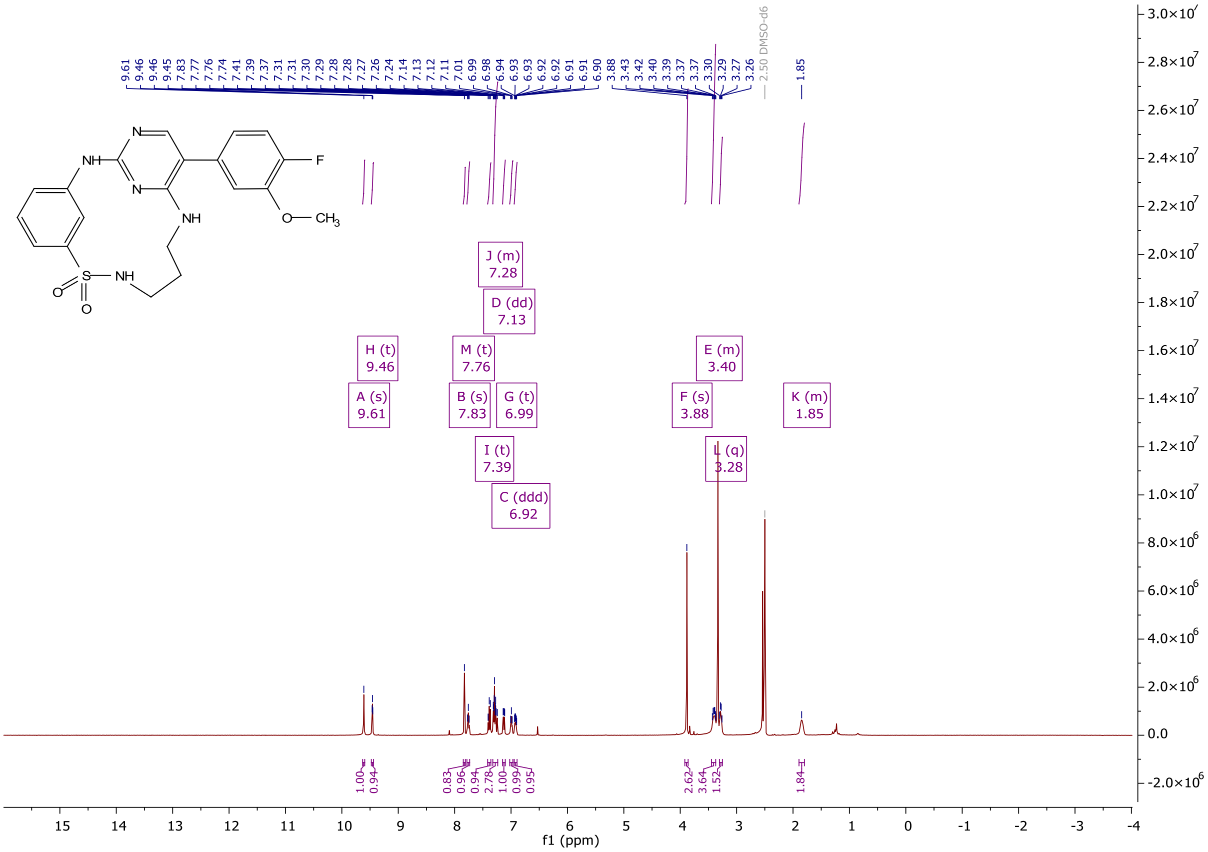

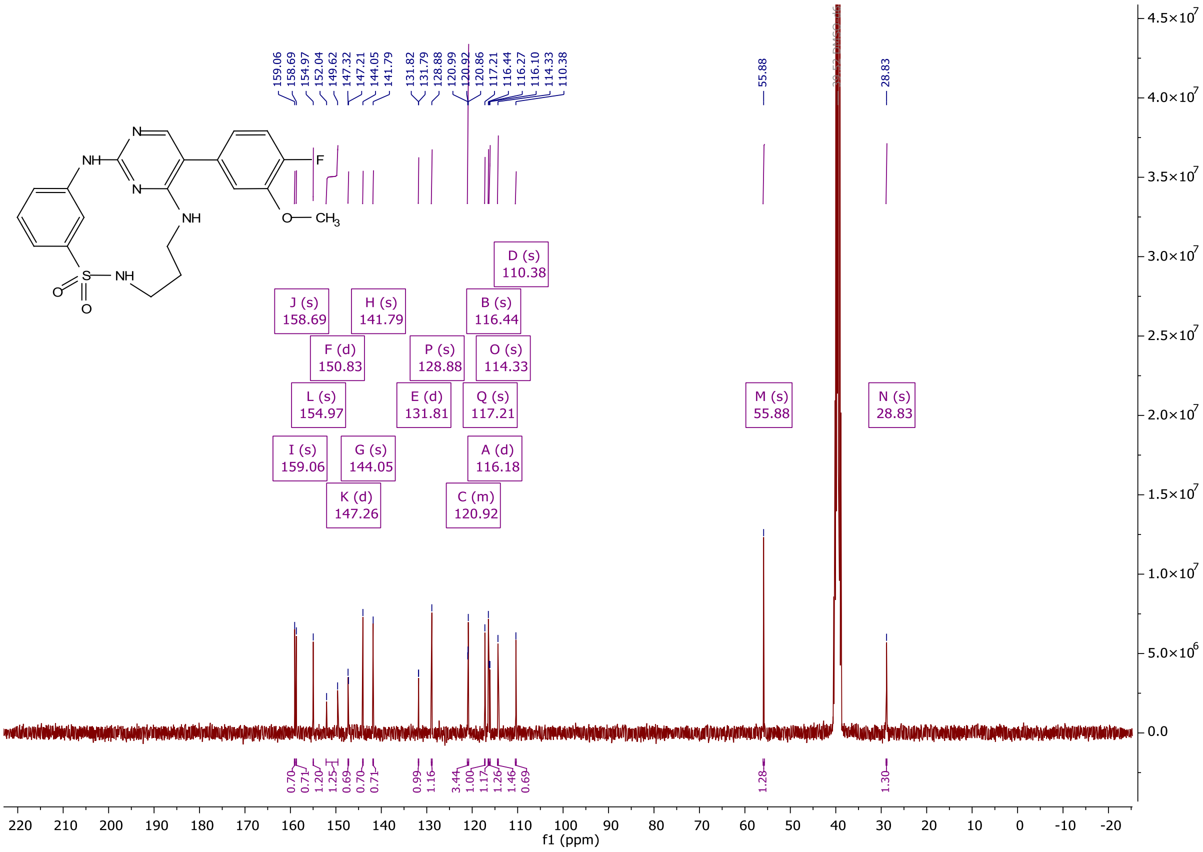

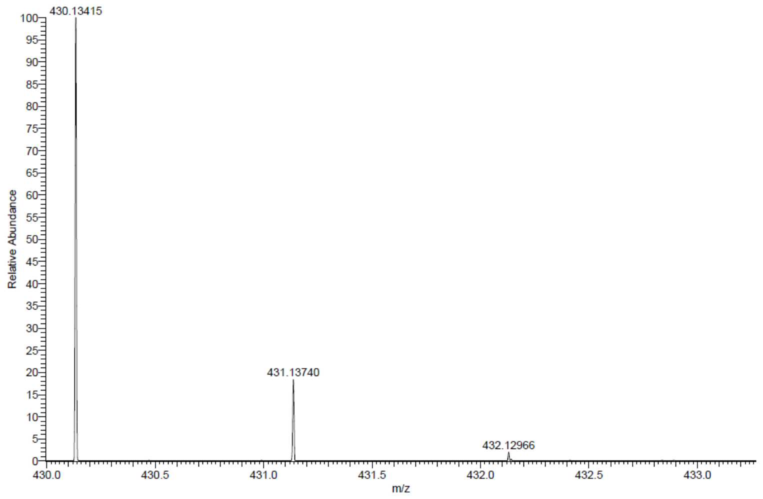

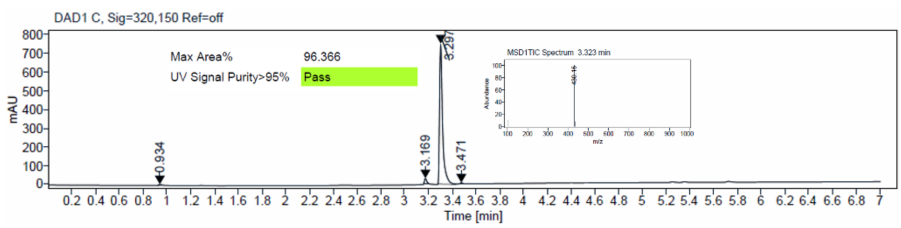
DAD1 C, Sig=320,150 Ref=off
Max Area% 96.366
UV Signal Purity>95% Pass
mAU
Time [min]

[1]H and [13]C {H} NMR, HRMS (Orbitrap) and HPLC/MS (ESI) data of compound **28**

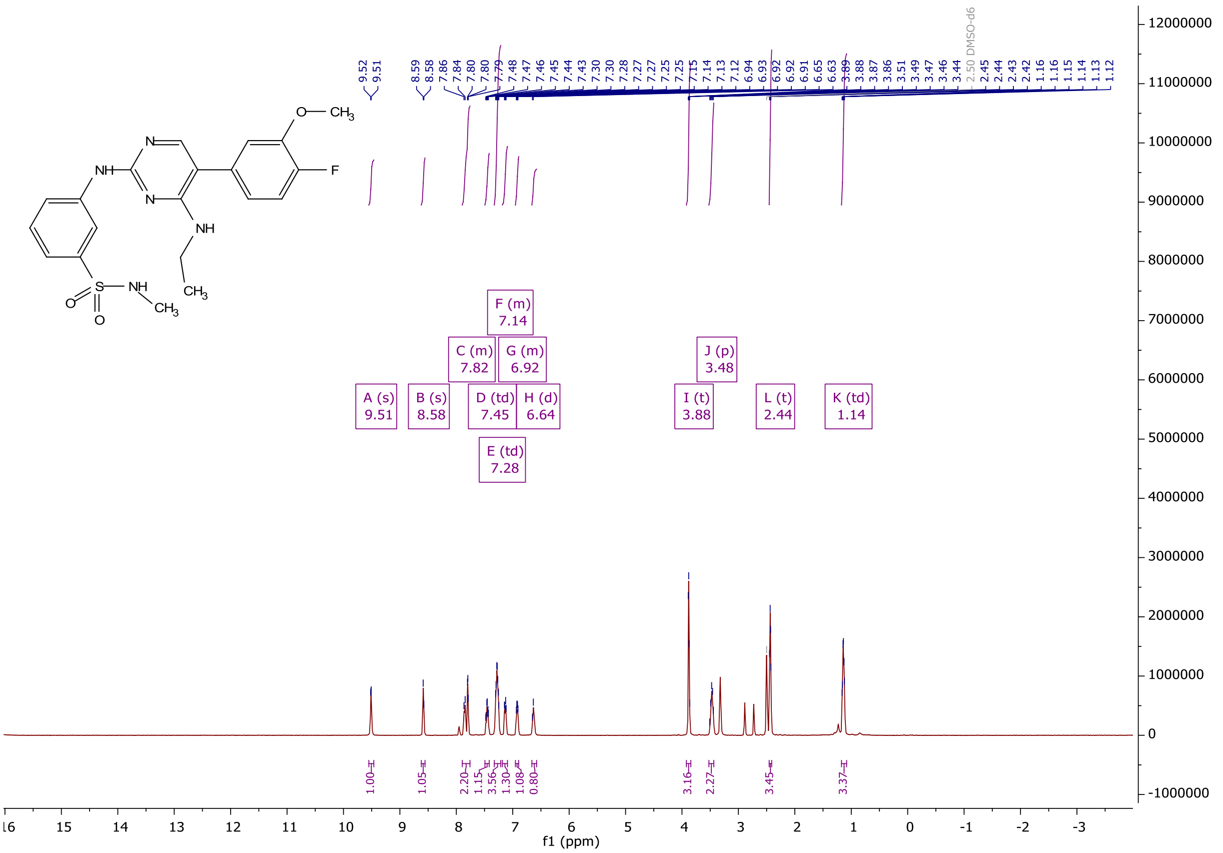

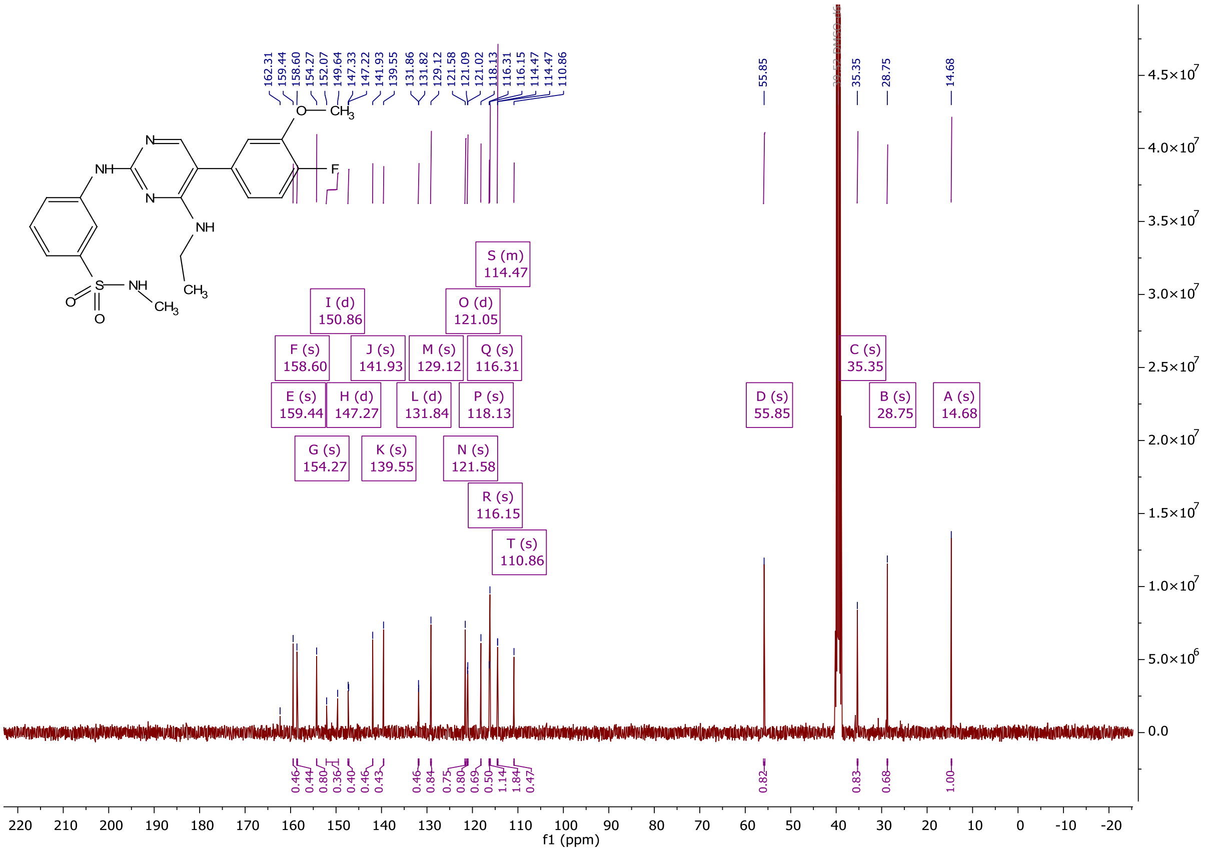

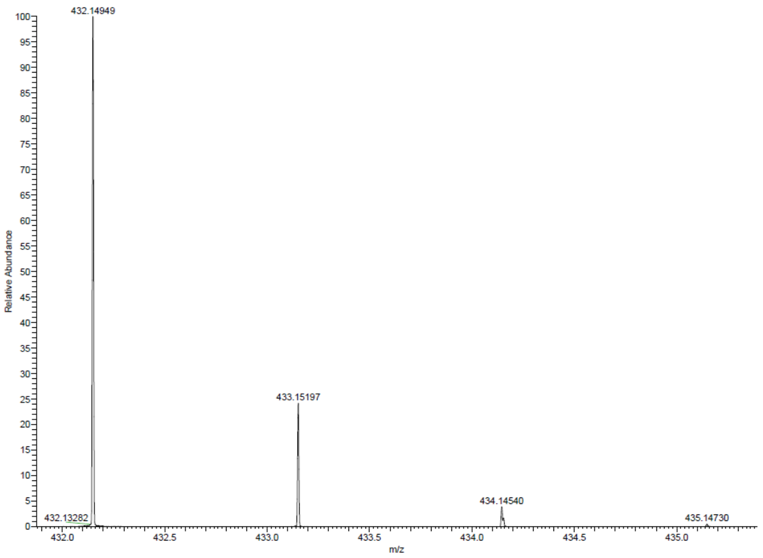

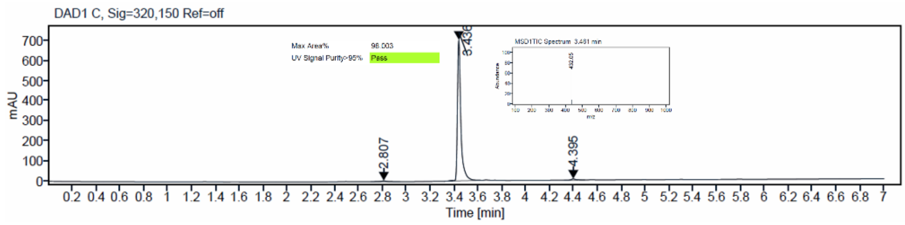

$^{1}$H and $^{13}$C {H} NMR, HRMS (Orbitrap) and HPLC/MS (ESI) data of compound **41**

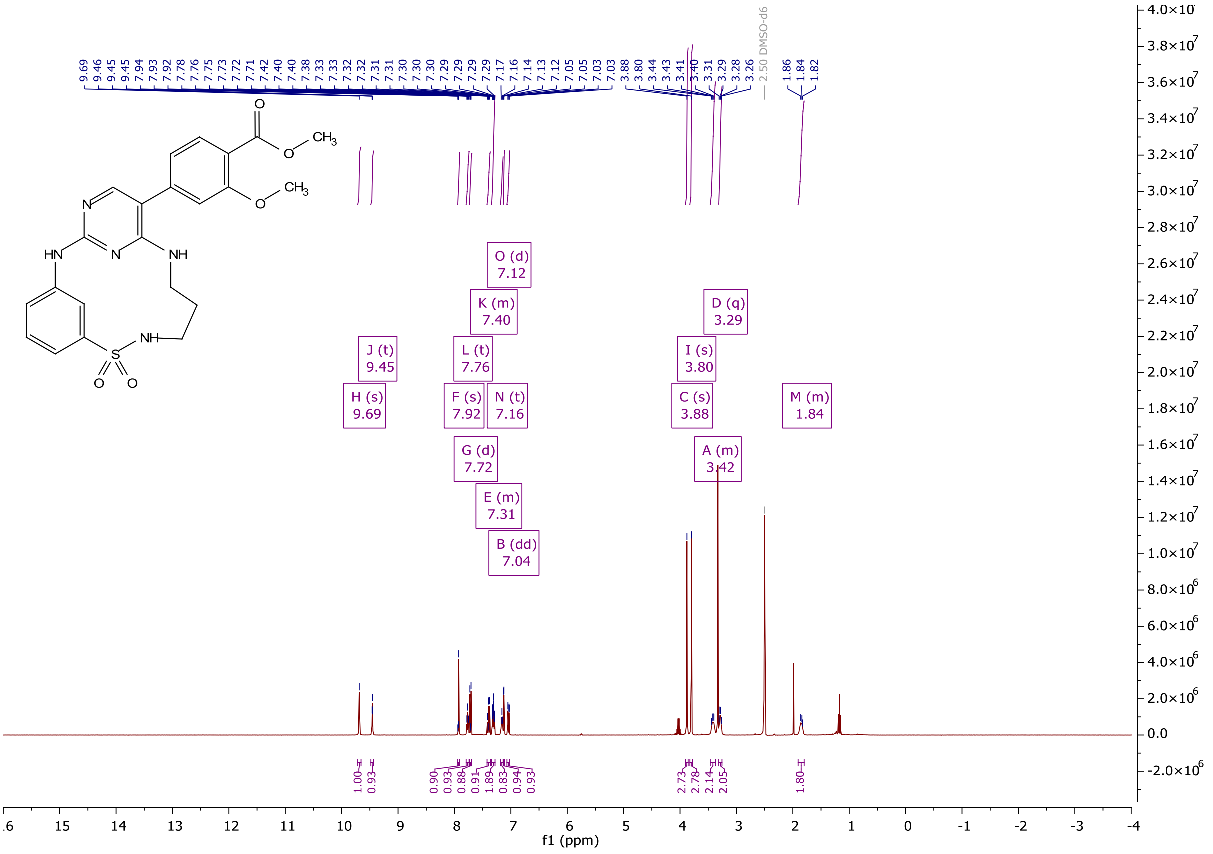

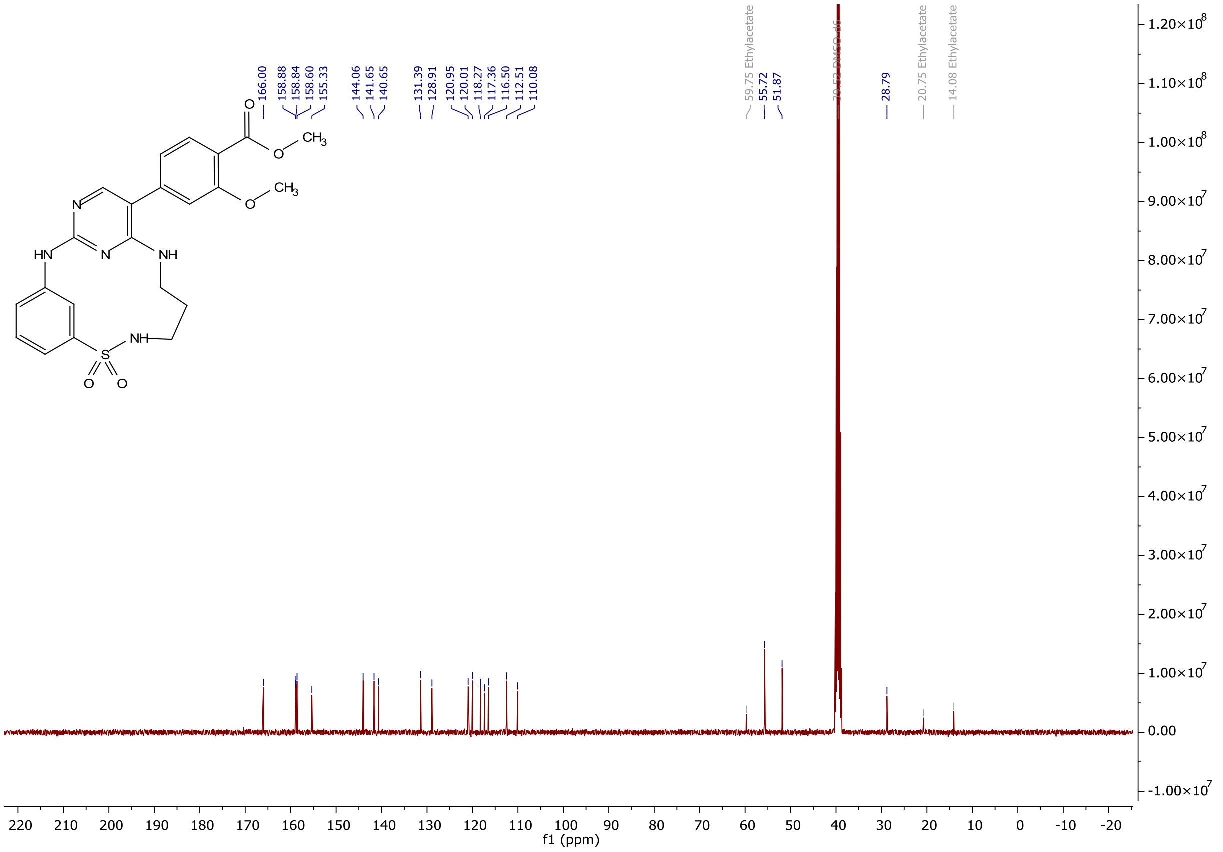
166.00
158.88
158.84
158.60
155.33
144.06
141.65
140.65
131.39
128.91
120.95
120.01
118.27
117.36
116.50
112.51
110.08
59.75 Ethylacetate
55.72
51.87
28.79
20.75 Ethylacetate
14.08 Ethylacetate
f1 (ppm)

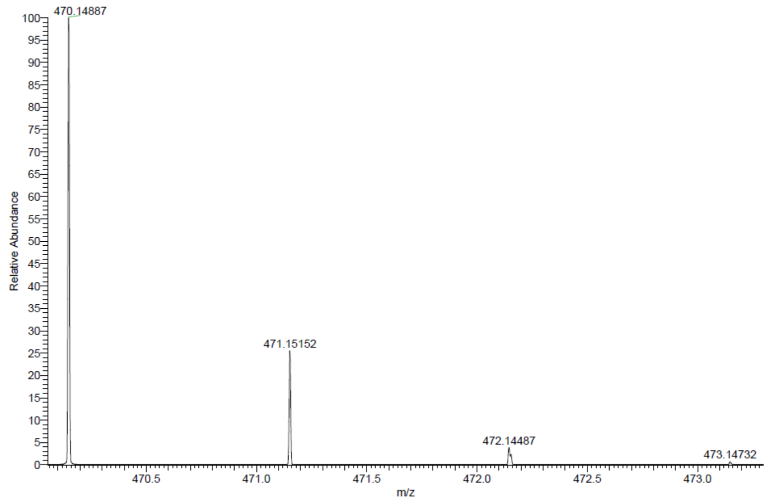
470.14887
471.15152
472.14487
473.14732
Relative Abundance
m/z

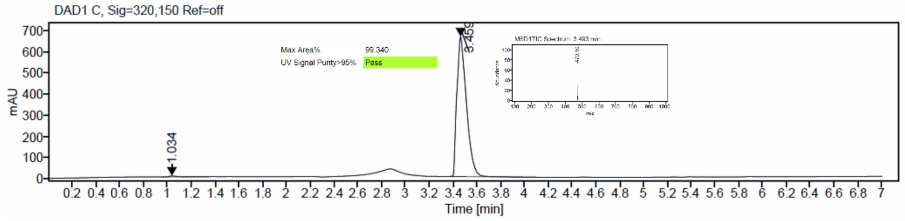
DAD1 C, Sig=320,150 Ref=off
mAU
Max Area%
99.340
UV Signal Purity>95%
Pass
3.459
1.034
Time [min]

$^{1}H$ and $^{13}C$ {H} NMR, HRMS (Orbitrap) and HPLC/MS (ESI) data of compound **42**

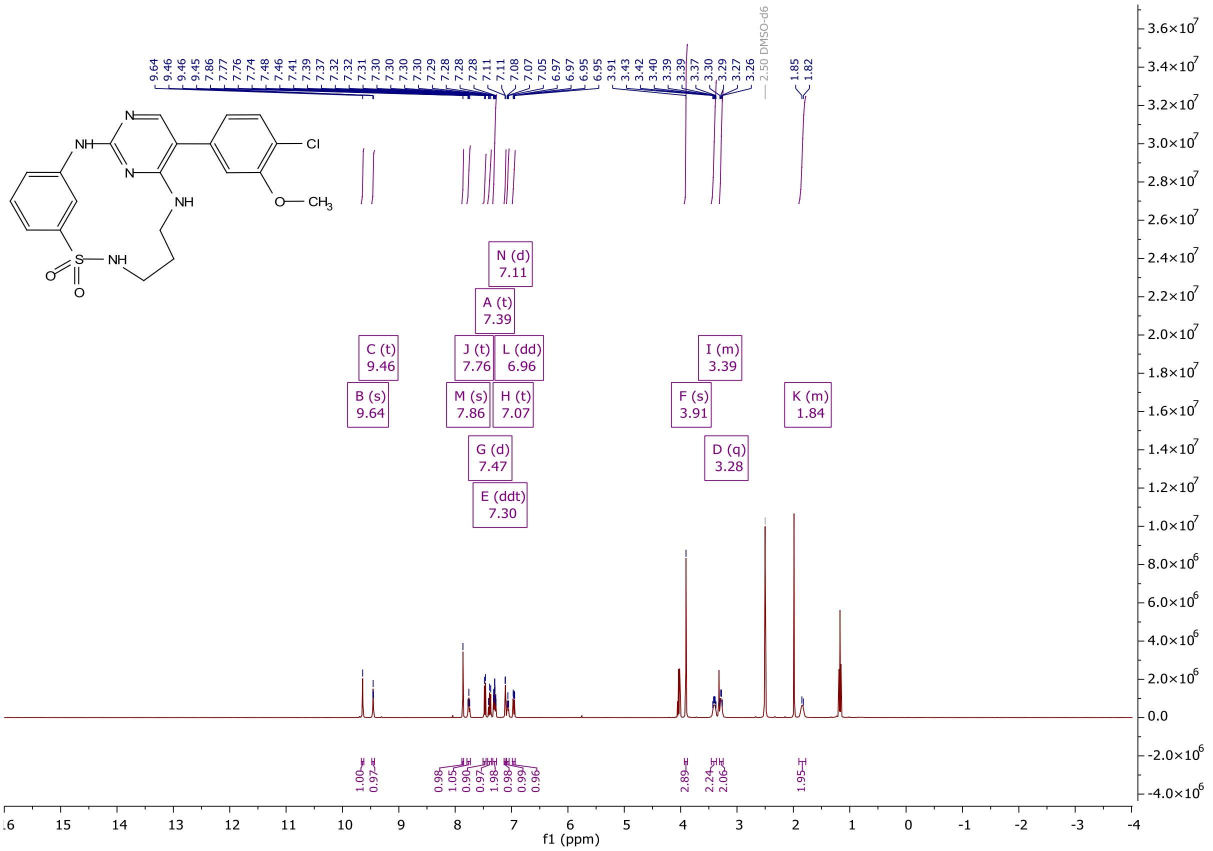

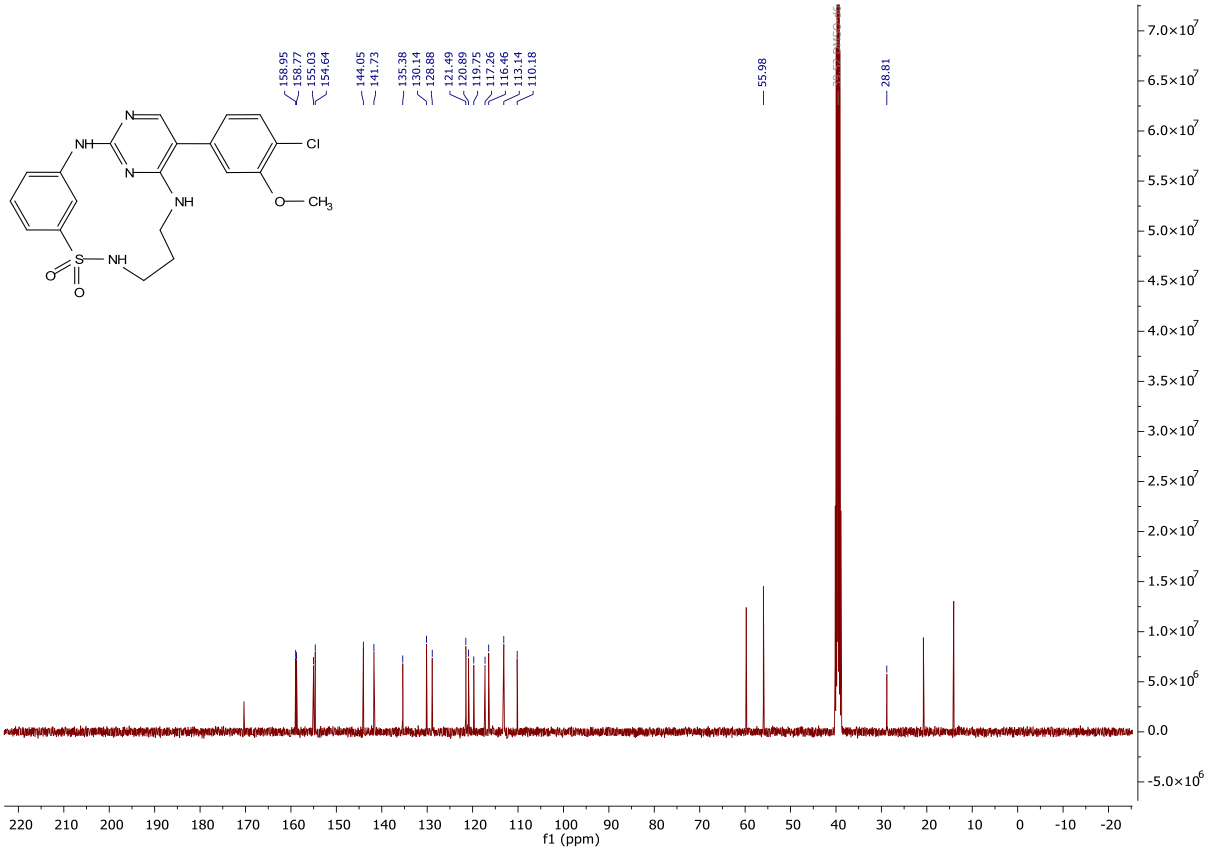

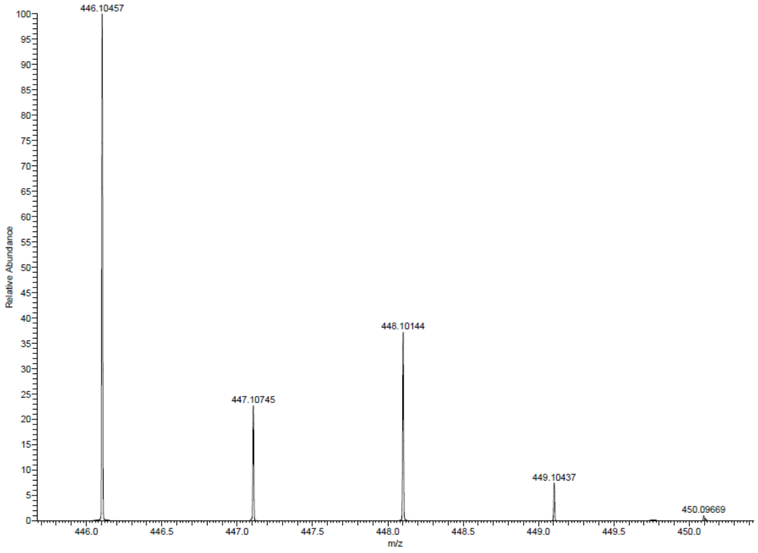

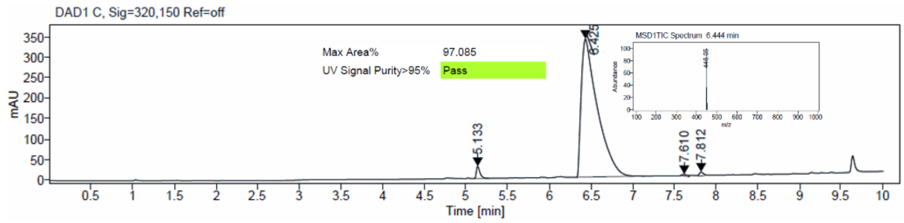

$^1H$ and $^{13}C$ {H} NMR, HRMS (Orbitrap) and HPLC/MS (ESI) data of compound **43**

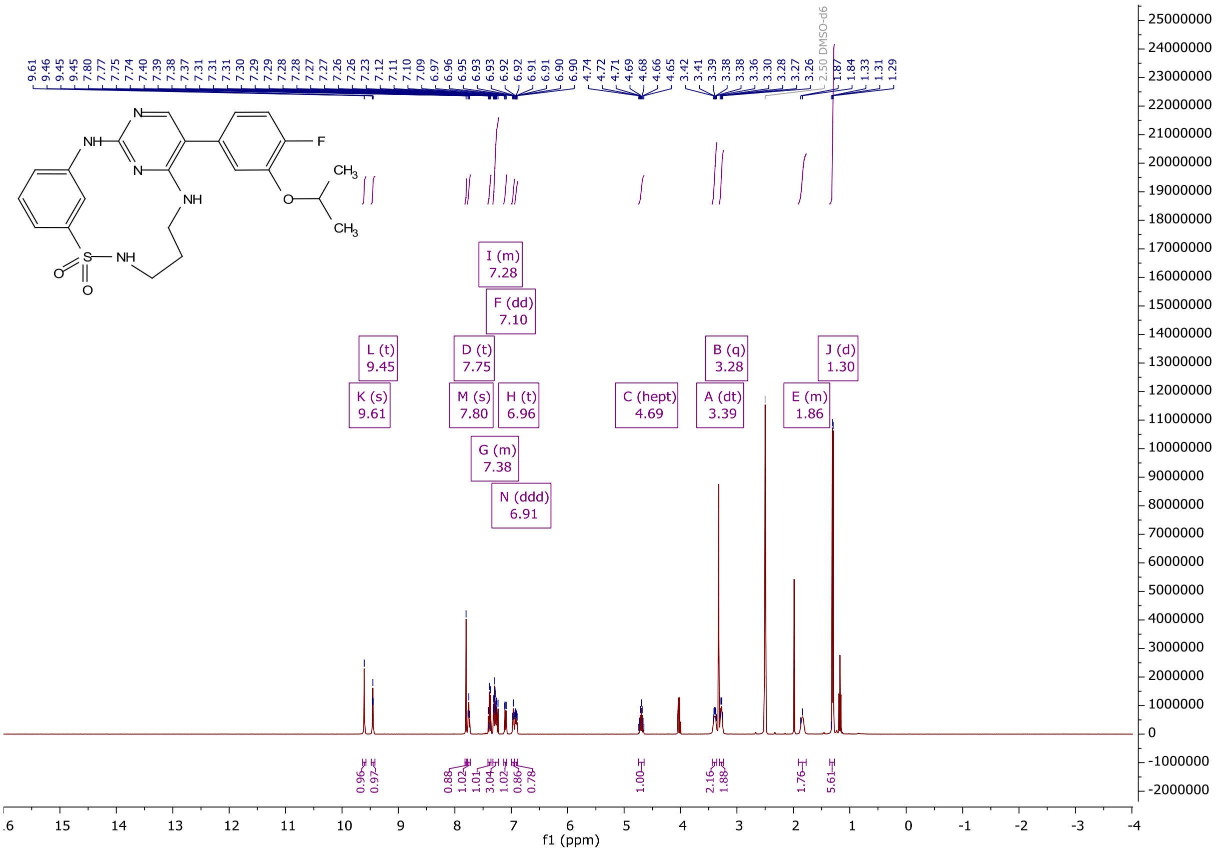

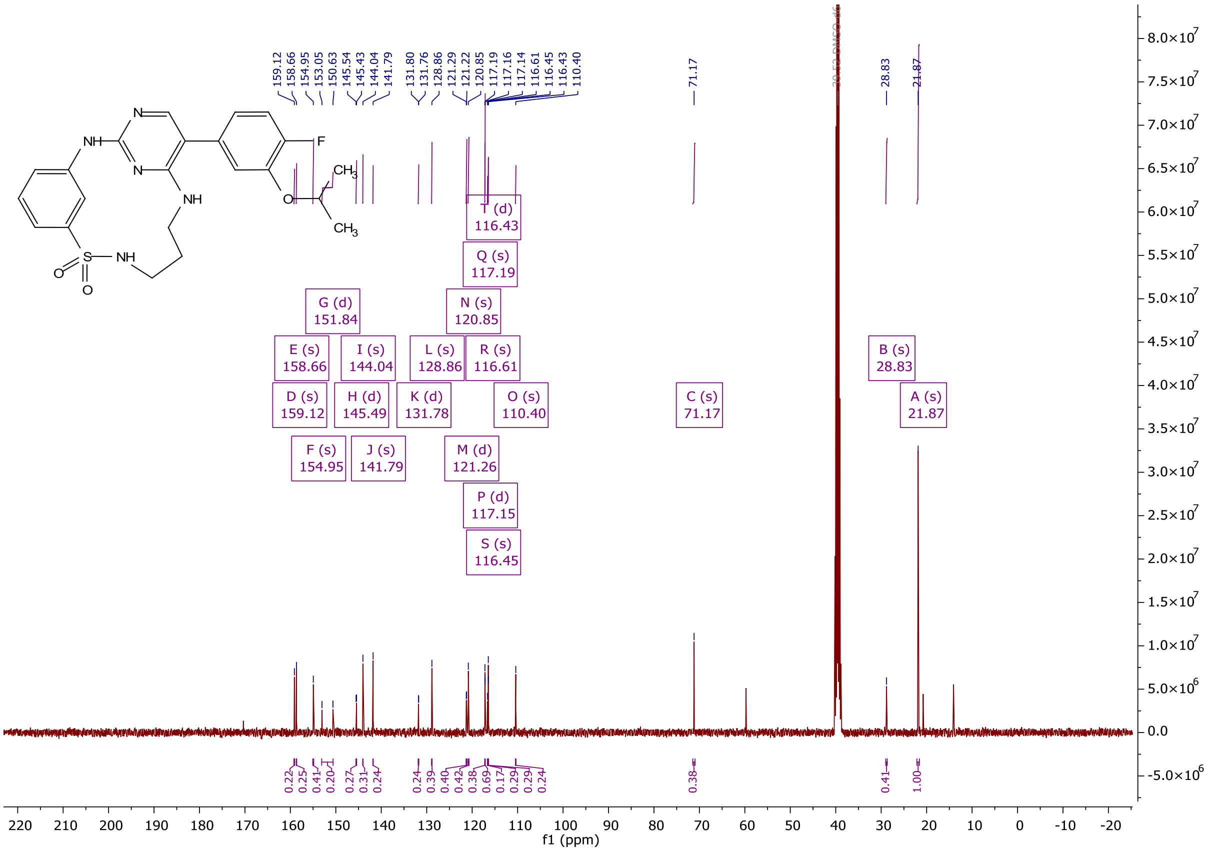

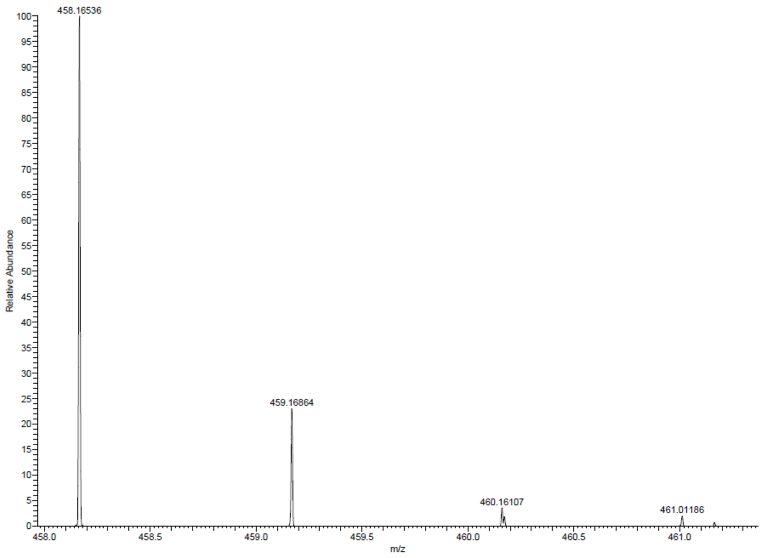

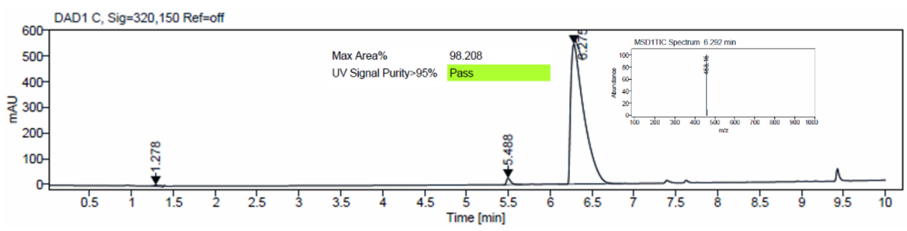

<sup></sup>[1]H and [13]C {H} NMR, HRMS (Orbitrap) and HPLC/MS (ESI) data of compound **44**

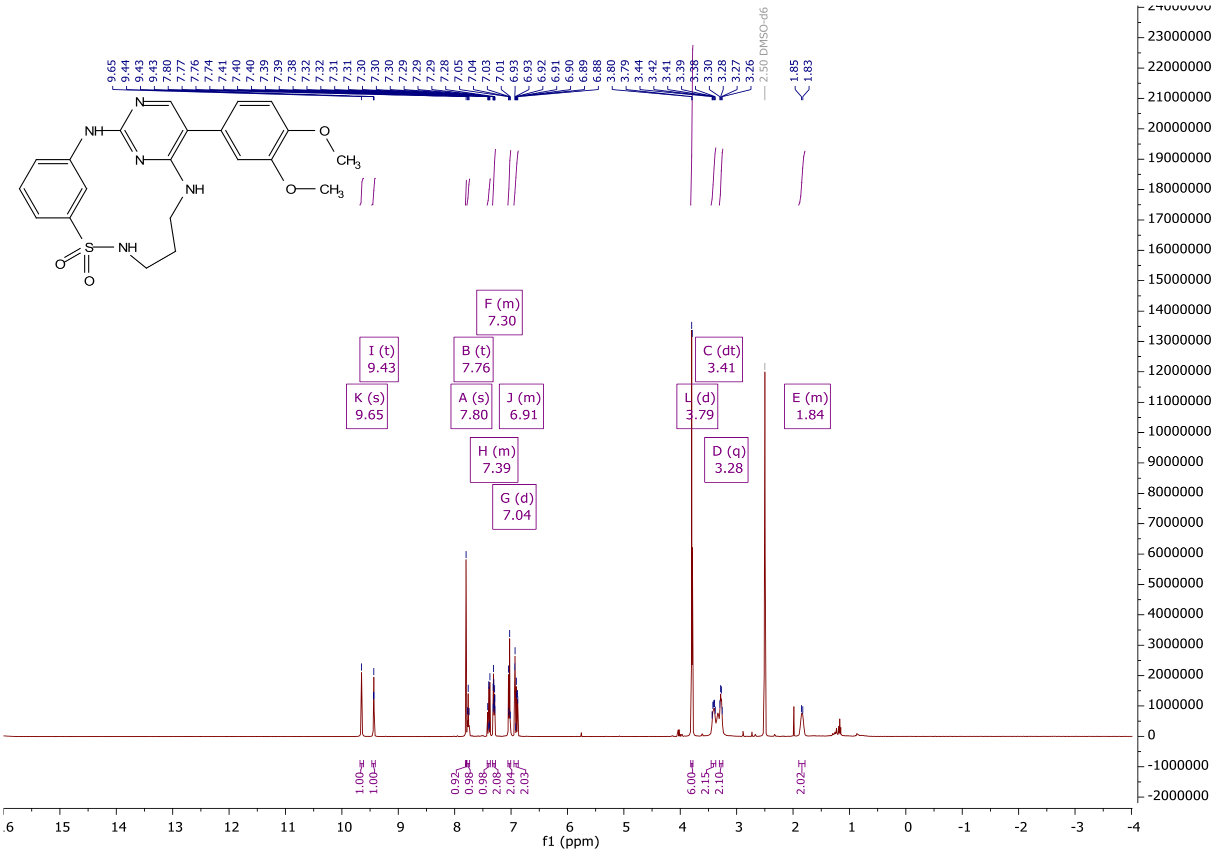

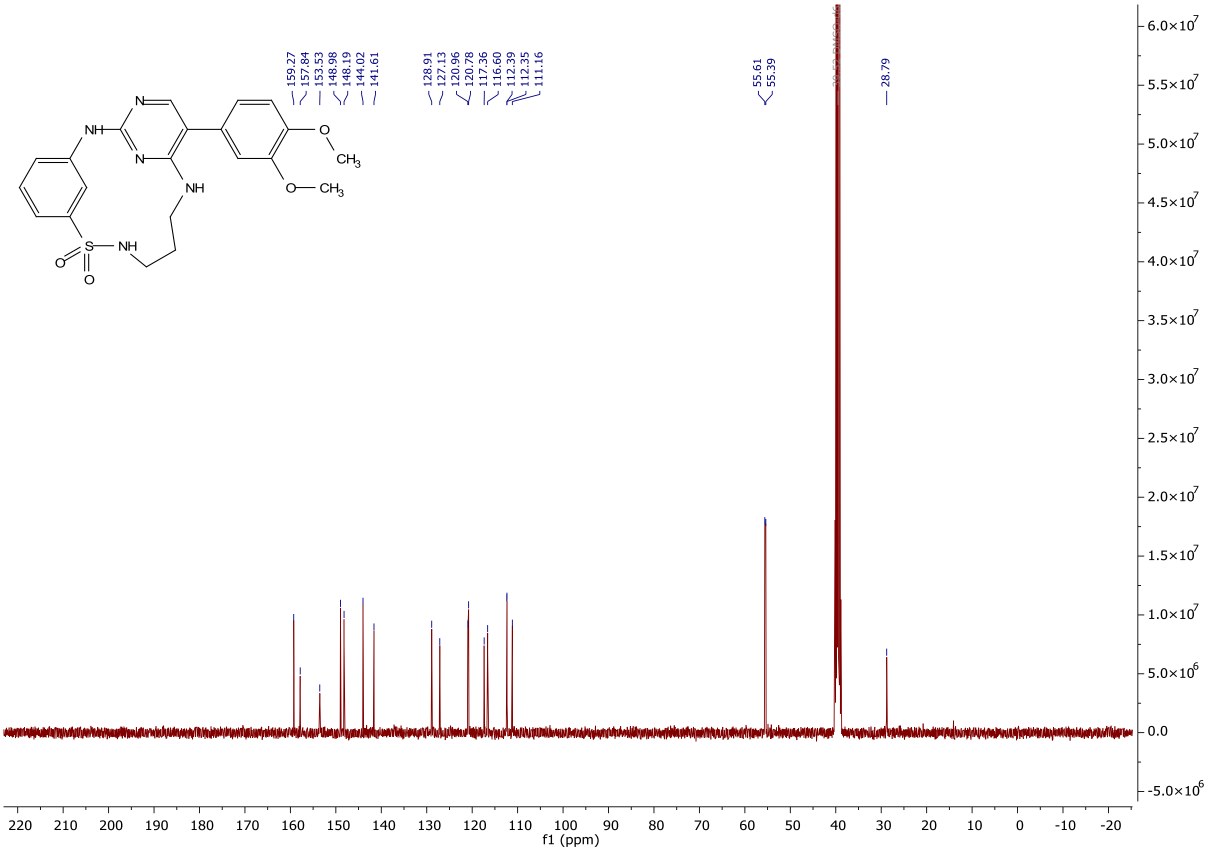

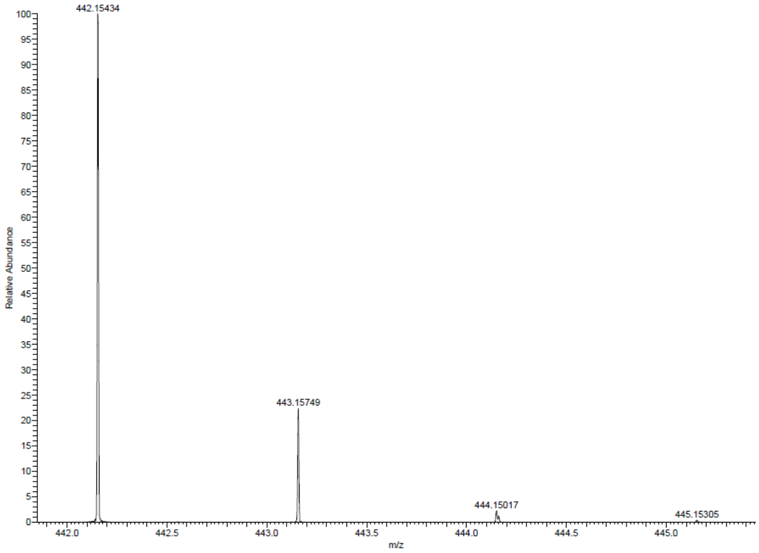

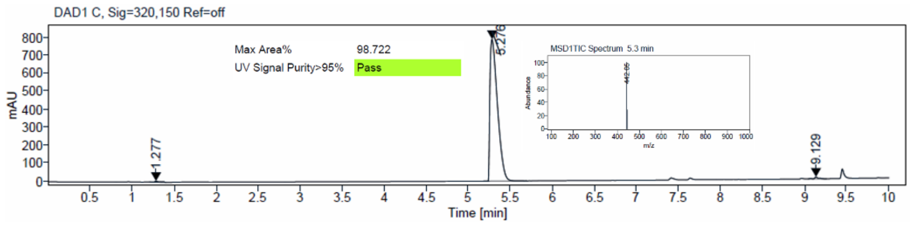

$^{1}H$ and $^{13}C$ {H} NMR, HRMS (Orbitrap) and HPLC/MS (ESI) data of compound **45**

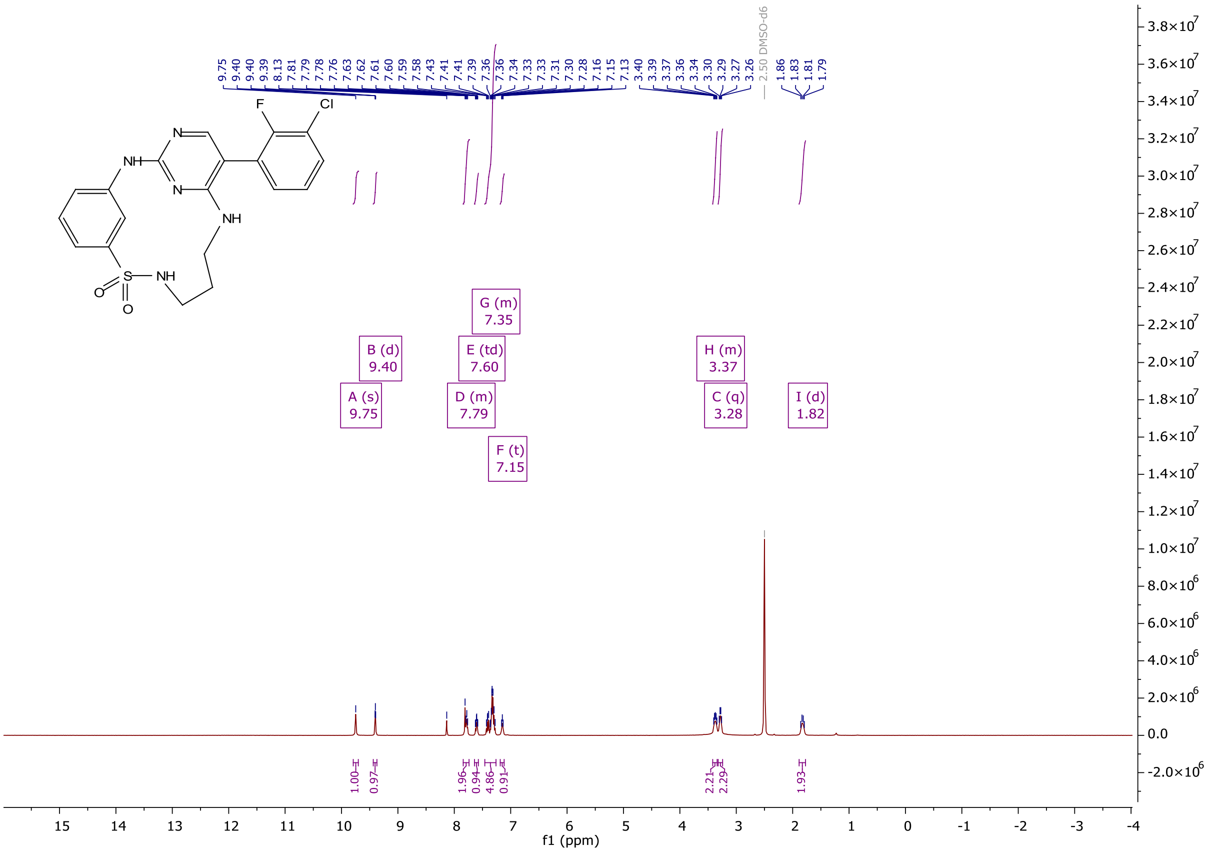

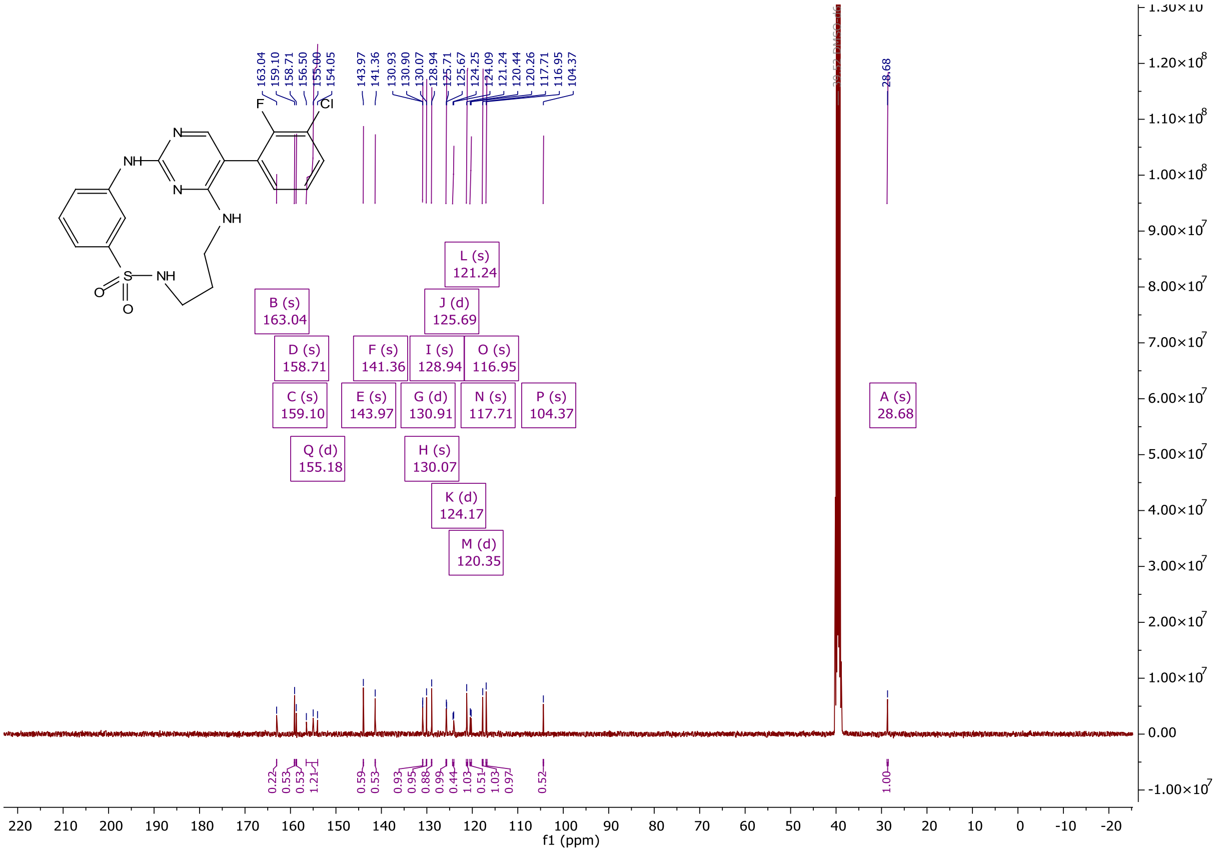

163.04
159.10
158.71
156.50
155.00
154.05
143.97
141.36
130.93
130.90
130.07
128.94
125.71
125.67
124.25
124.09
121.24
120.44
120.26
117.71
116.95
104.37
28.68
L (s) 121.24
B (s) 163.04
J (d) 125.69
D (s) 158.71
F (s) 141.36
I (s) 128.94
O (s) 116.95
C (s) 159.10
E (s) 143.97
G (d) 130.91
N (s) 117.71
P (s) 104.37
A (s) 28.68
Q (d) 155.18
H (s) 130.07
K (d) 124.17
M (d) 120.35
f1 (ppm)

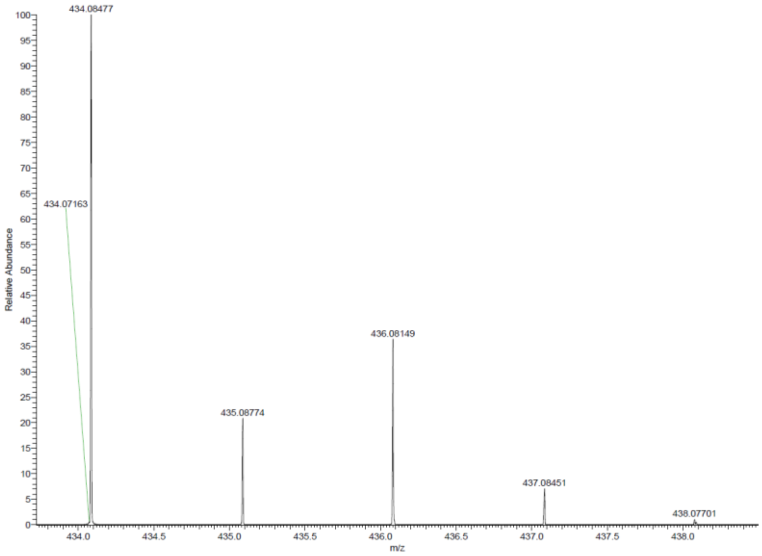

434.08477
434.07163
436.08149
435.08774
437.08451
438.07701
Relative Abundance
m/z

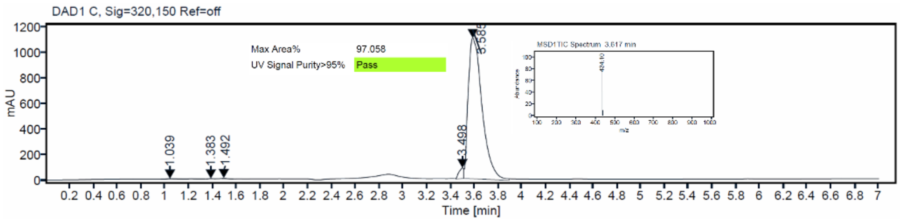

DAD1 C, Sig=320,150 Ref=off
Max Area% 97.058
UV Signal Purity>95% Pass
mAU
1.039
1.383
1.492
3.498
3.585
Time [min]

$^{1}H$ and $^{13}C$ {H} NMR, HRMS (Orbitrap) and HPLC/MS (ESI) data of compound **46**

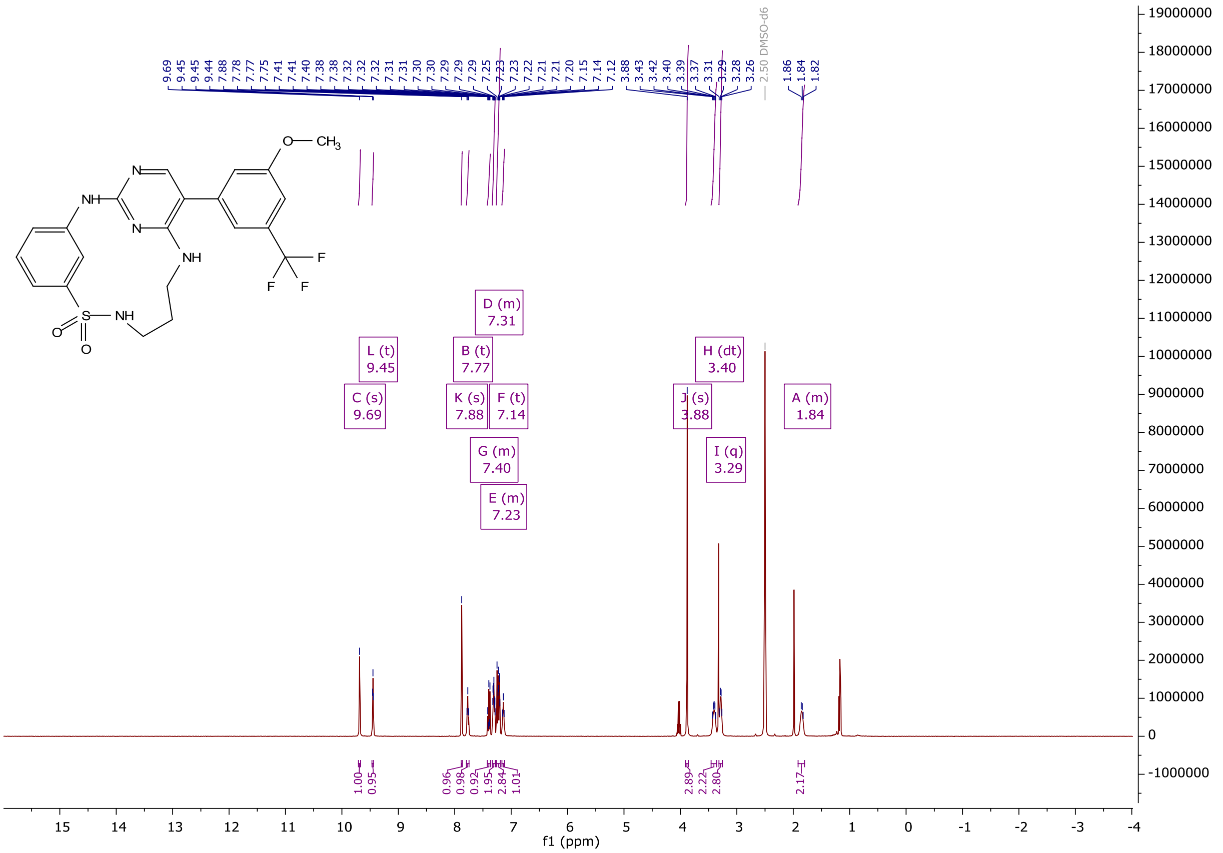

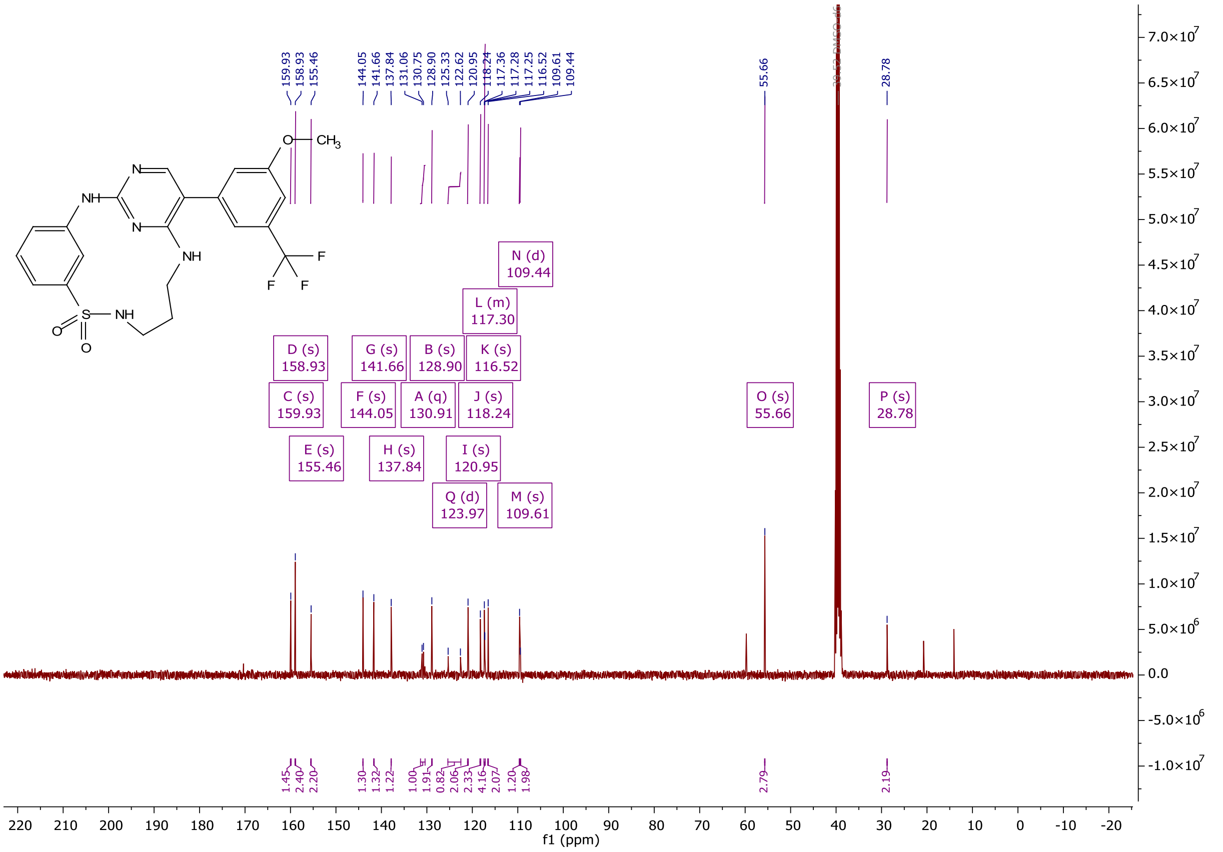

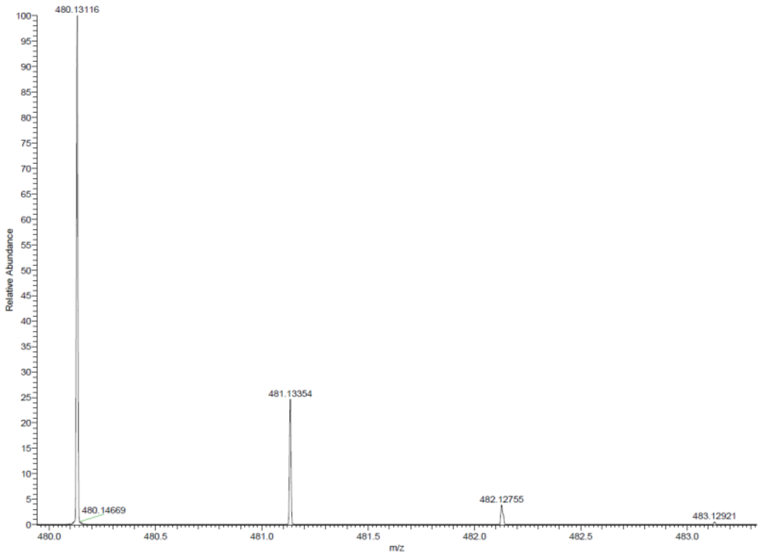

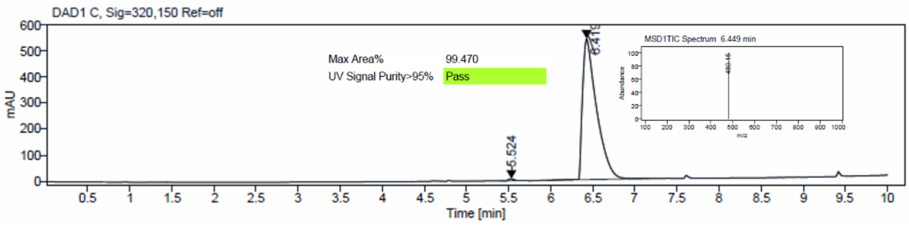

$^{1}$H and $^{13}$C {H} NMR, HRMS (Orbitrap) and HPLC/MS (ESI) data of compound **47**

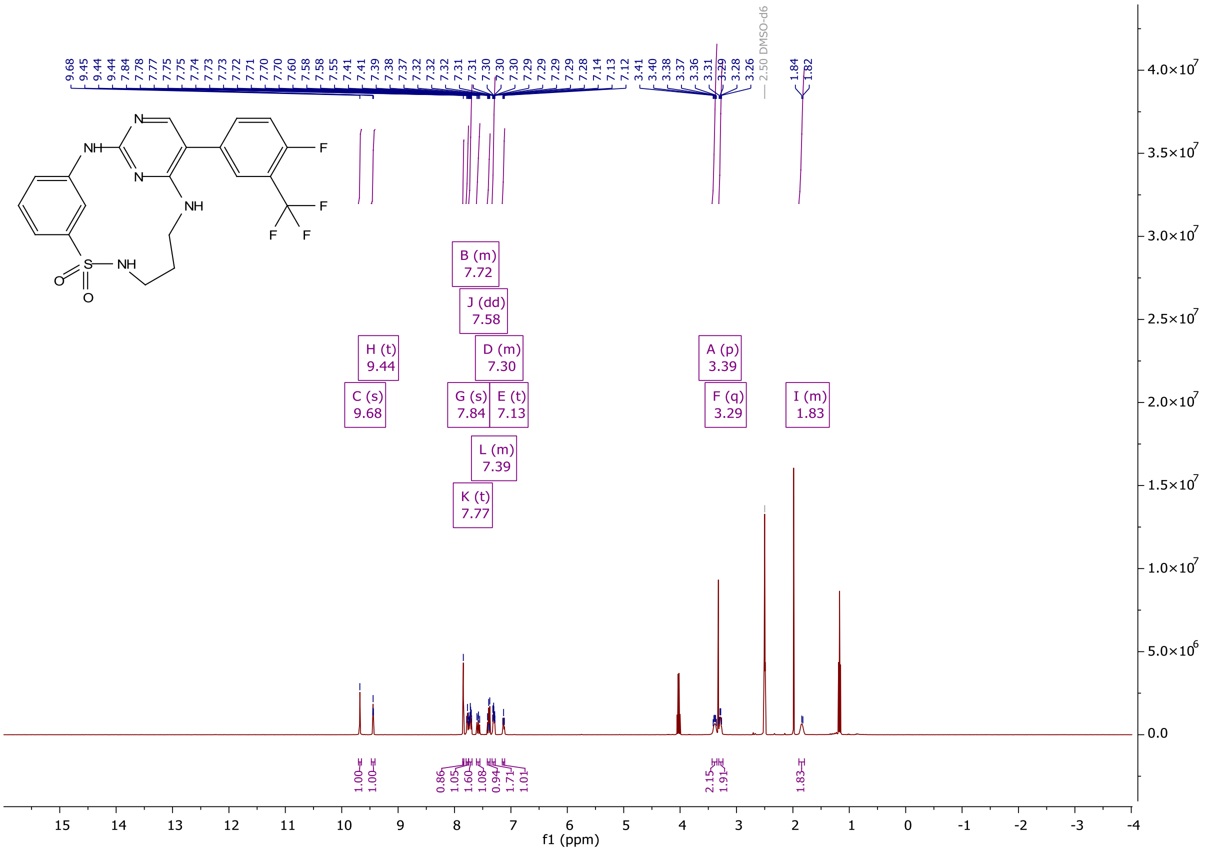

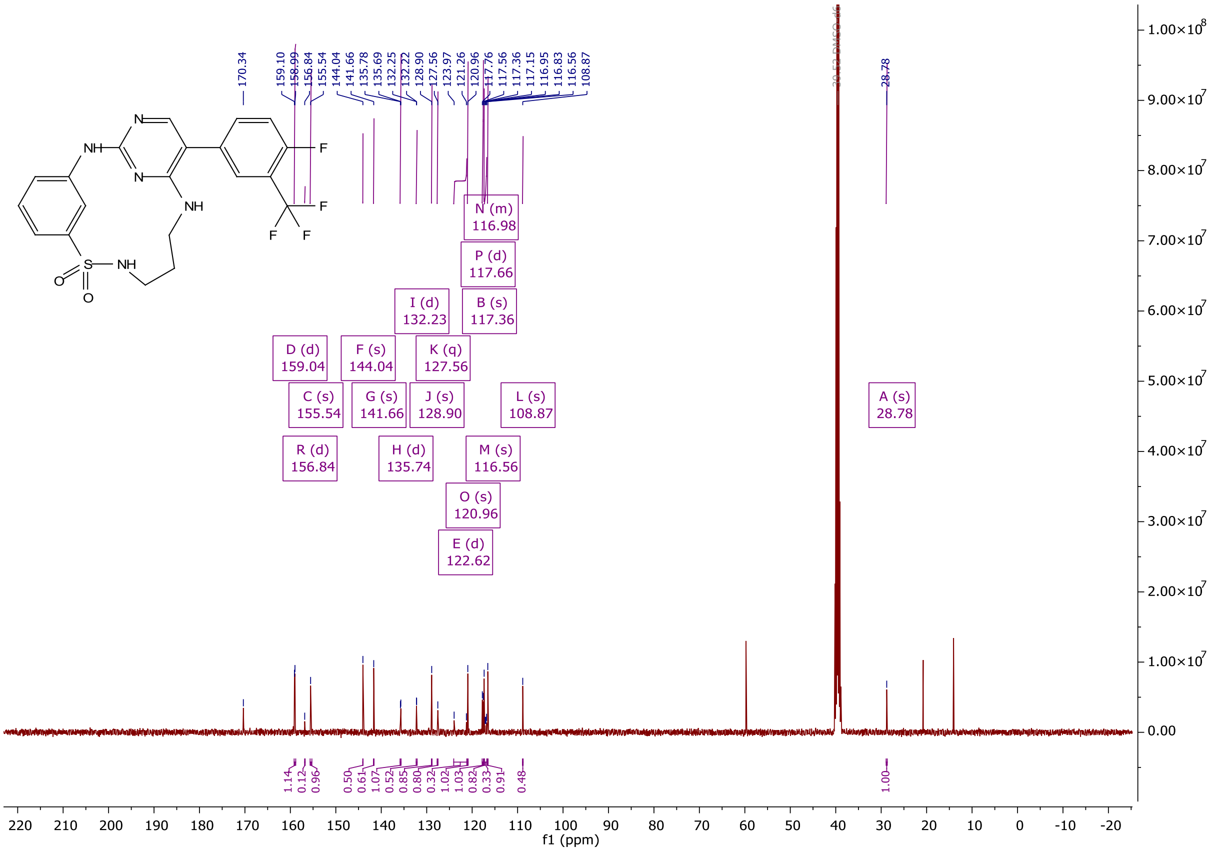

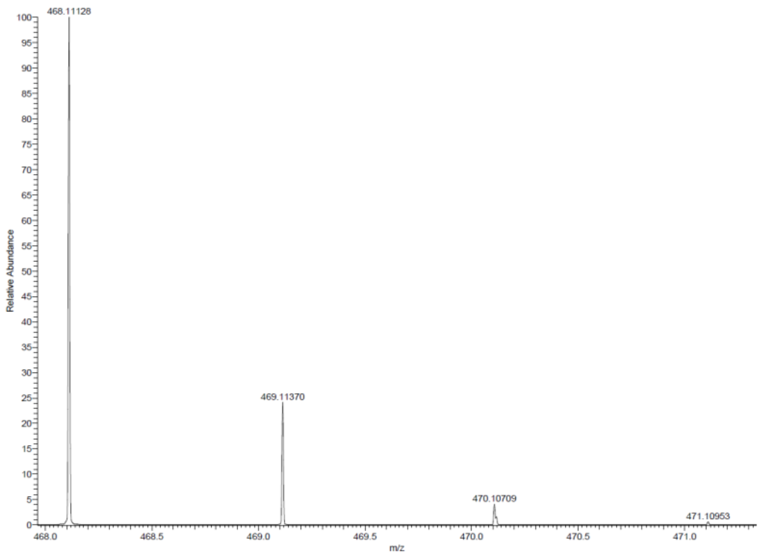

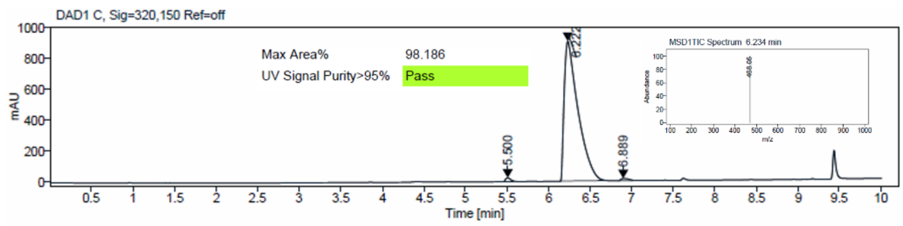
DAD1 C, Sig=320,150 Ref=off
Max Area% 98.186
UV Signal Purity>95% Pass
MSD1TIC Spectrum 6.234 min
Time [min]
mAU

[1]H and [13]C {H} NMR, HRMS (oTOF) and HPLC/MS (ESI) data of compound **48**

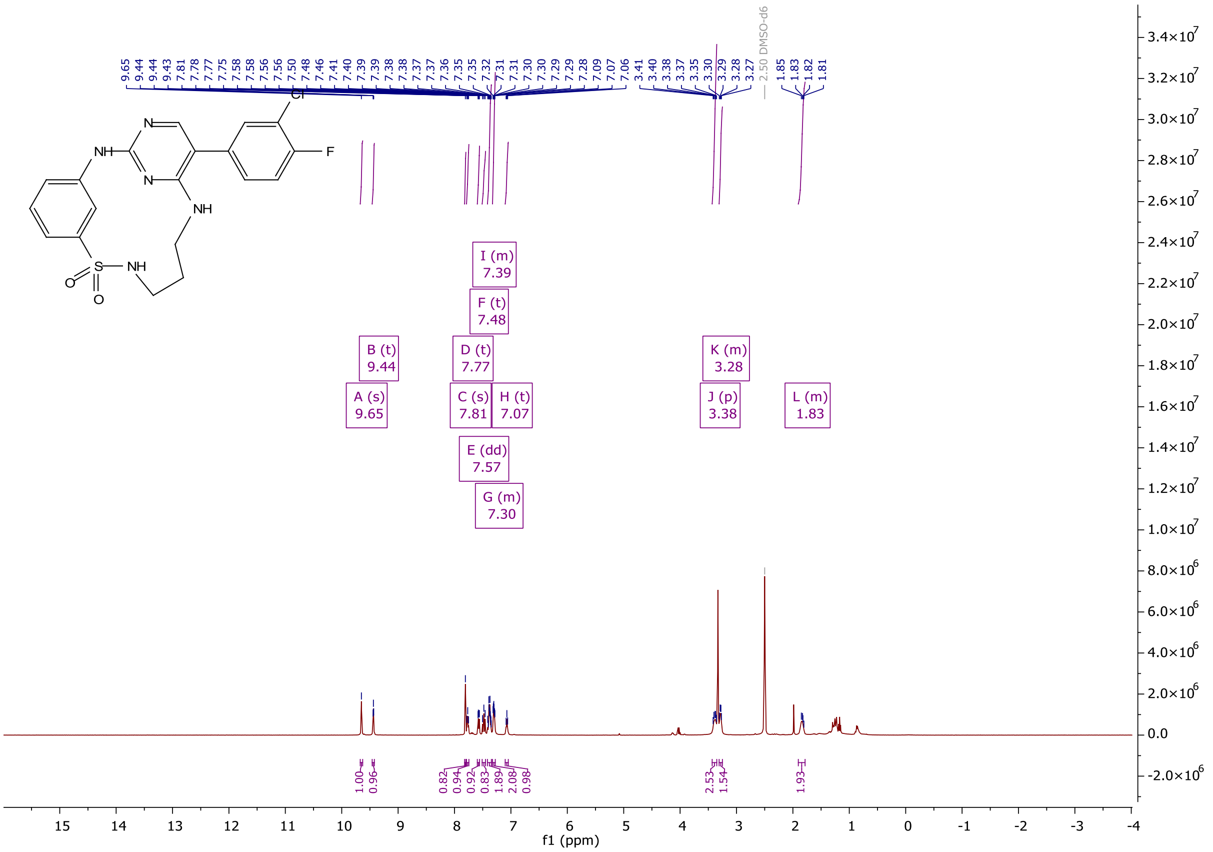

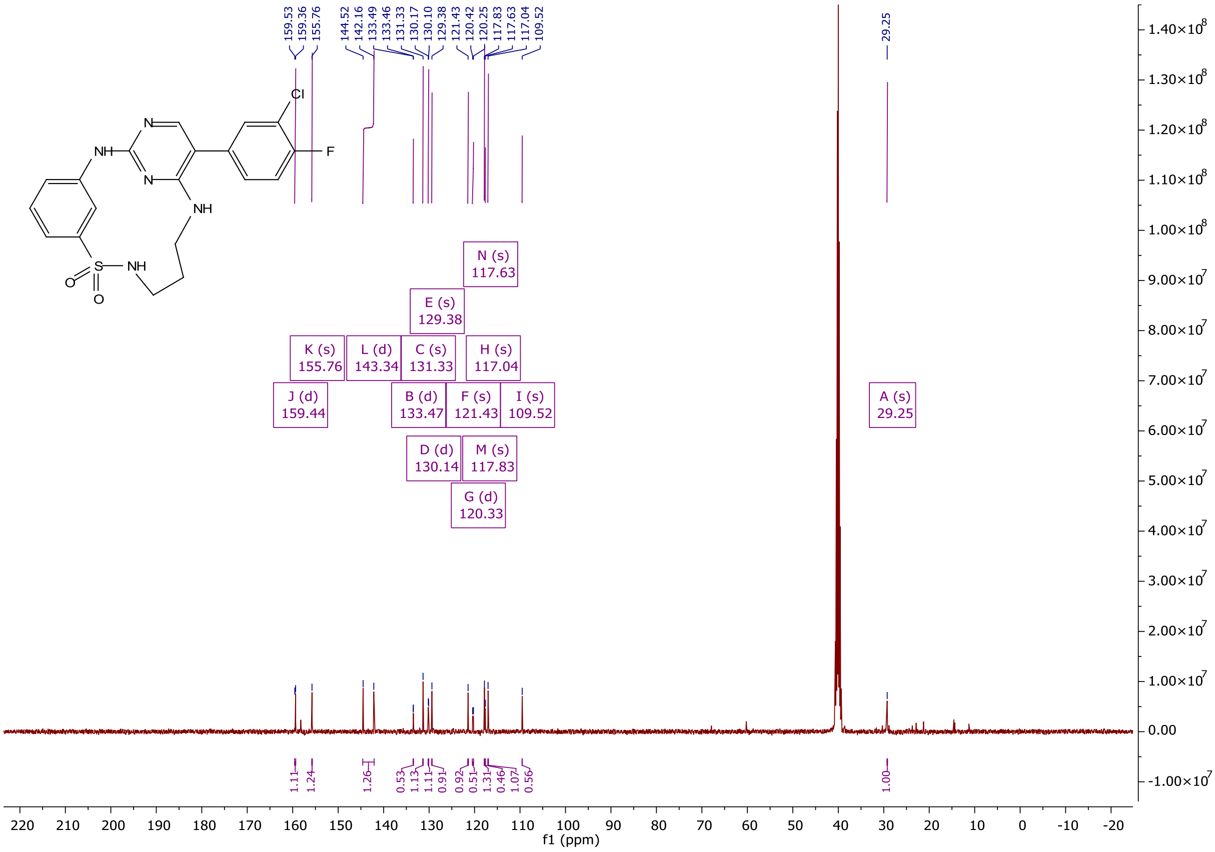

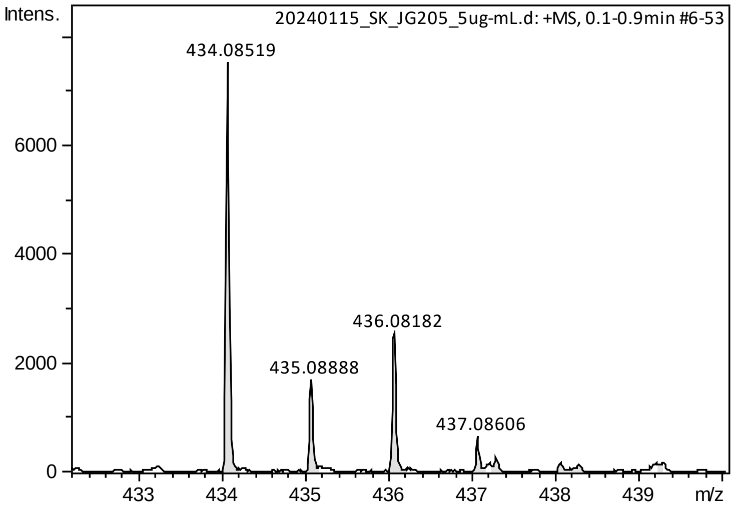

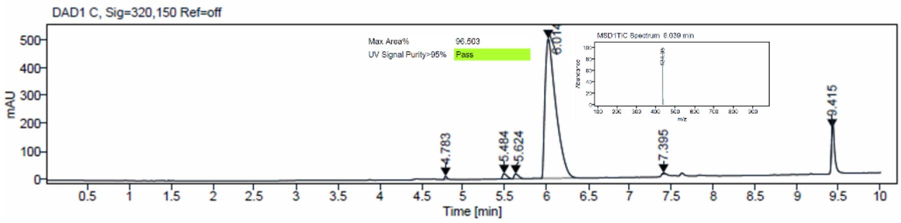

[1]H and [13]C {H} NMR, HRMS (oTOF) and HPLC/MS (ESI) data of compound **49**

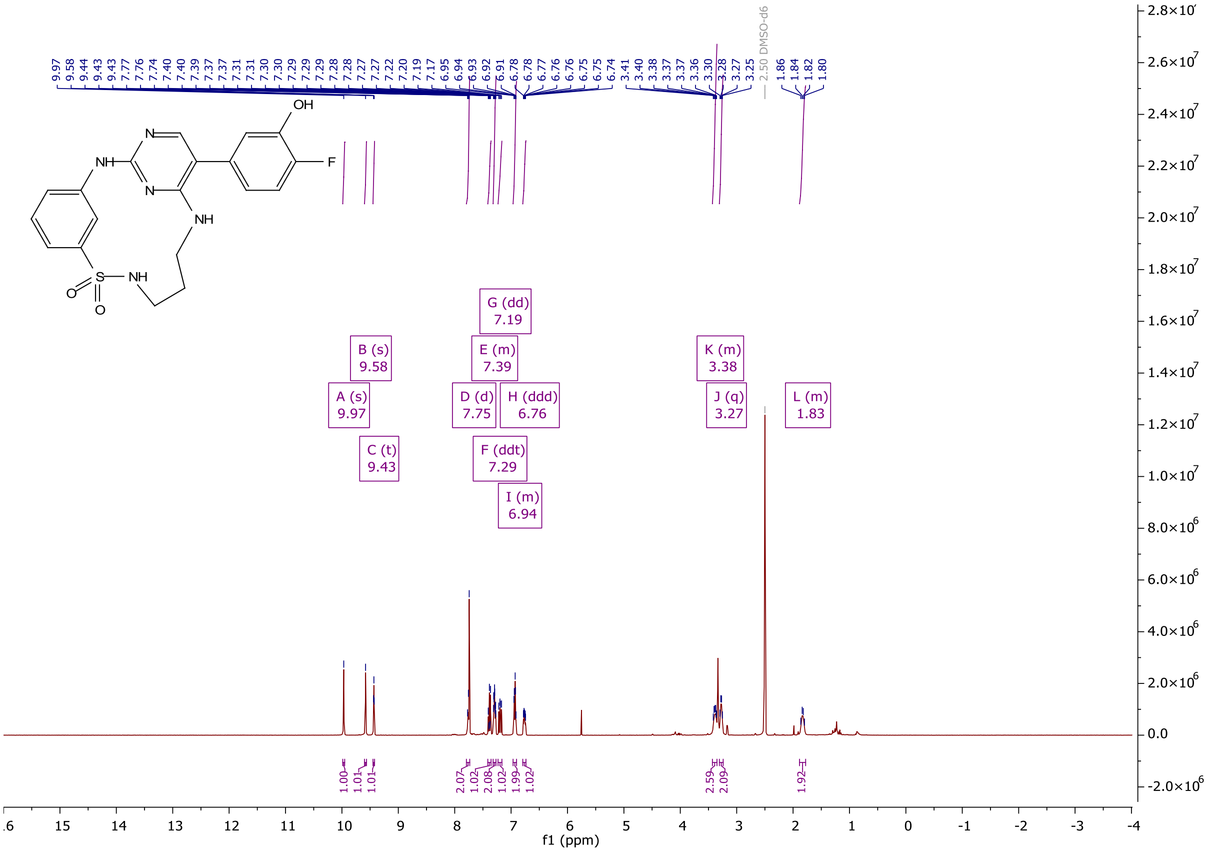

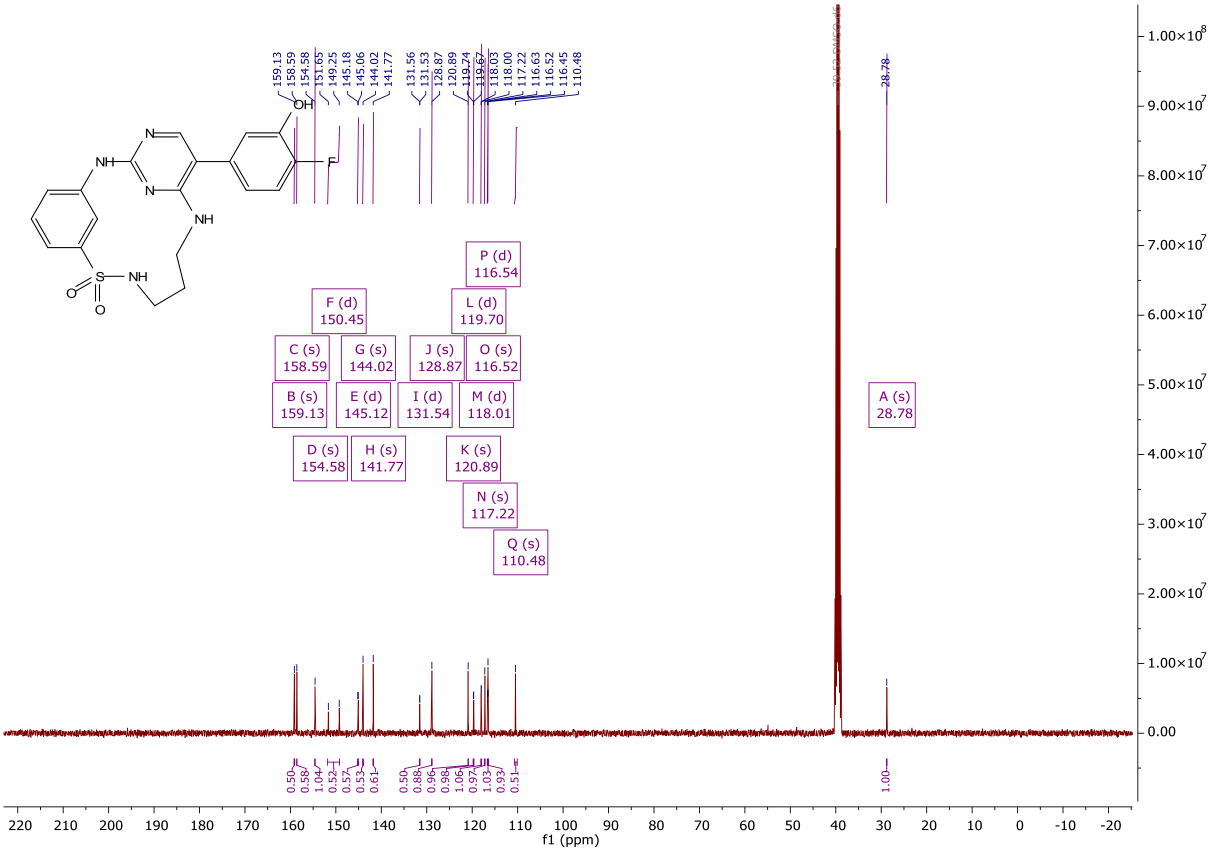

NH
N
N
NH
OH
F
S
NH
O
O
C (s) 158.59
B (s) 159.13
D (s) 154.58
F (d) 150.45
G (s) 144.02
E (d) 145.12
H (s) 141.77
J (s) 128.87
I (d) 131.54
K (s) 120.89
L (d) 119.70
M (d) 118.01
N (s) 117.22
O (s) 116.52
P (d) 116.54
Q (s) 110.48
A (s) 28.78
f1 (ppm)

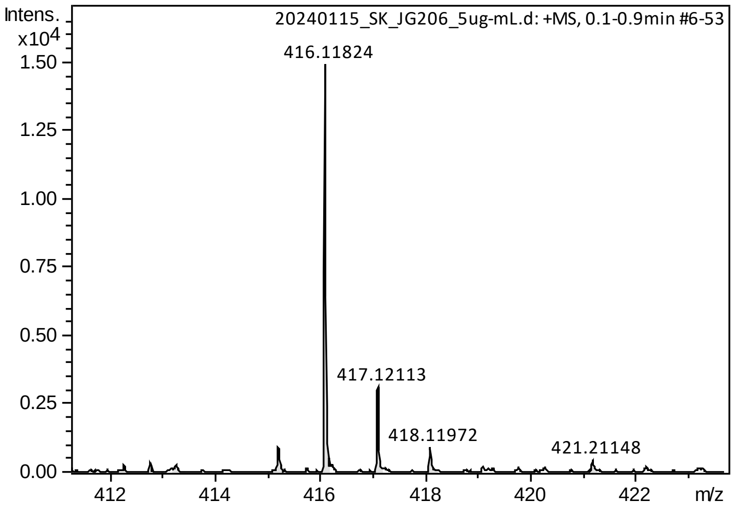

Intens.
x10^4
20240115_SK_JG206_5ug-mL.d: +MS, 0.1-0.9min #6-53
416.11824
417.12113
418.11972
421.21148
m/z

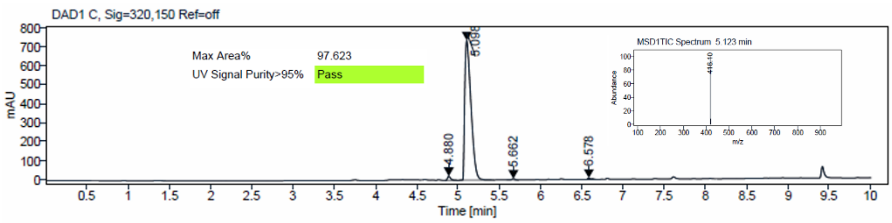

DAD1 C, Sig=320,150 Ref=off
Max Area% 97.623
UV Signal Purity>95% Pass
mAU
Time [min]
4.880
5.098
5.662
6.578
MSD1TIC Spectrum 5.123 min
Abundance
m/z

[1]H and [13]C {H} NMR, HRMS (Orbitrap) and HPLC/MS (ESI) data of compound **50**

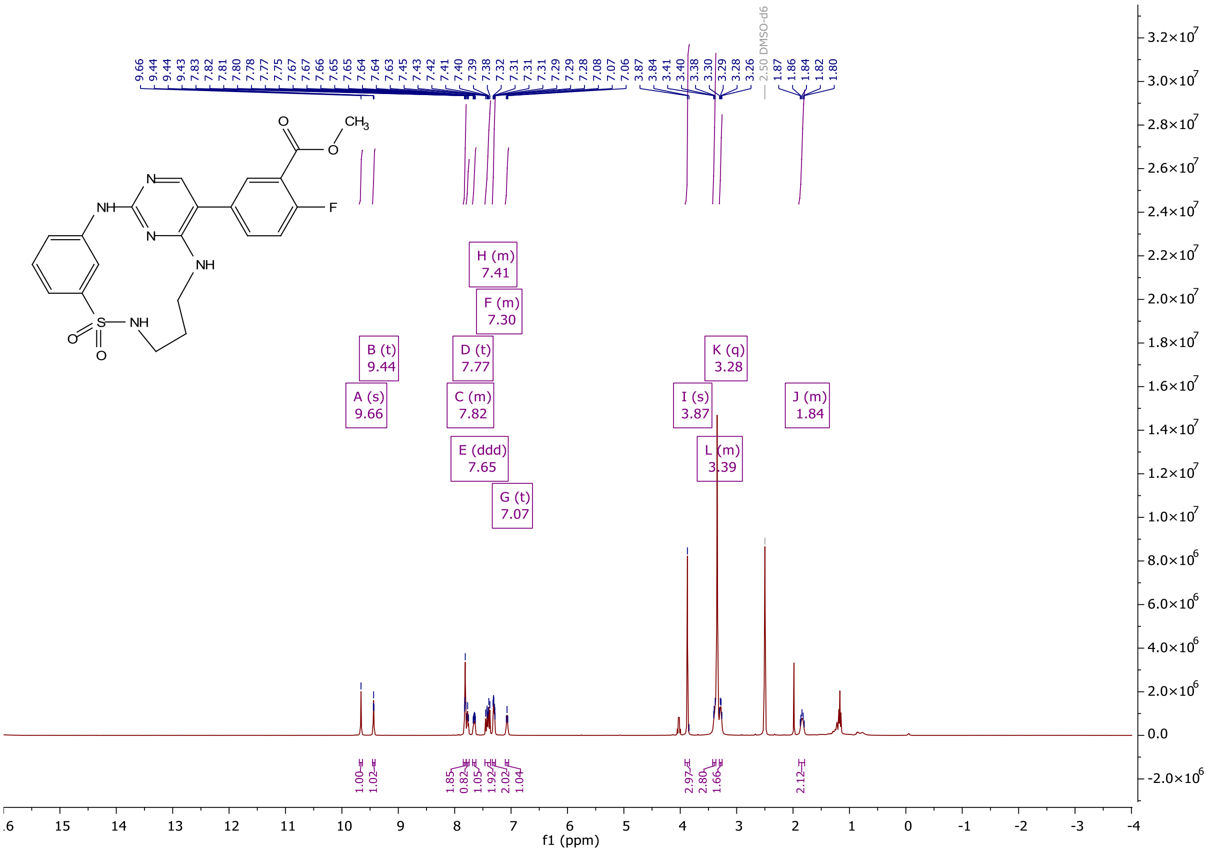

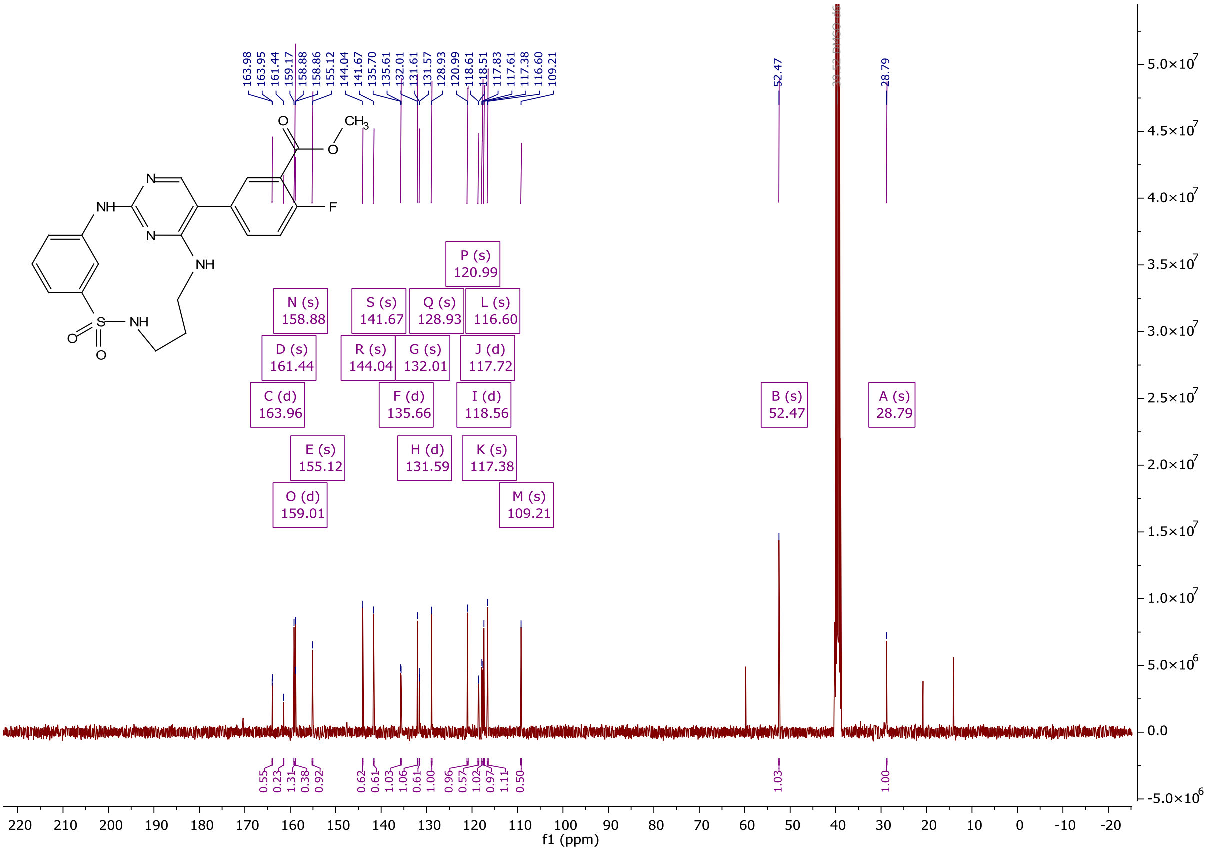

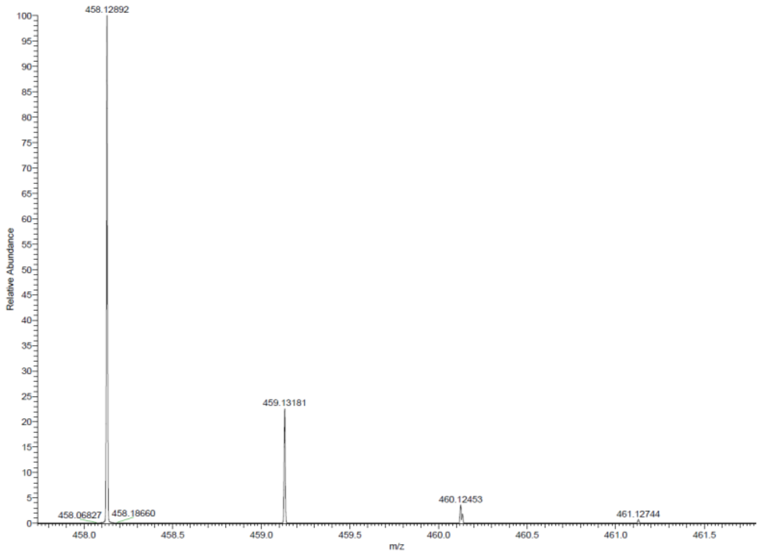

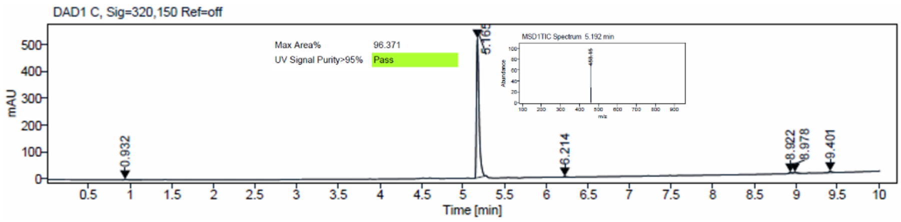

$^1$H and $^{13}$C {H} NMR, HRMS (Orbitrap) and HPLC/MS (ESI) data of compound **51**

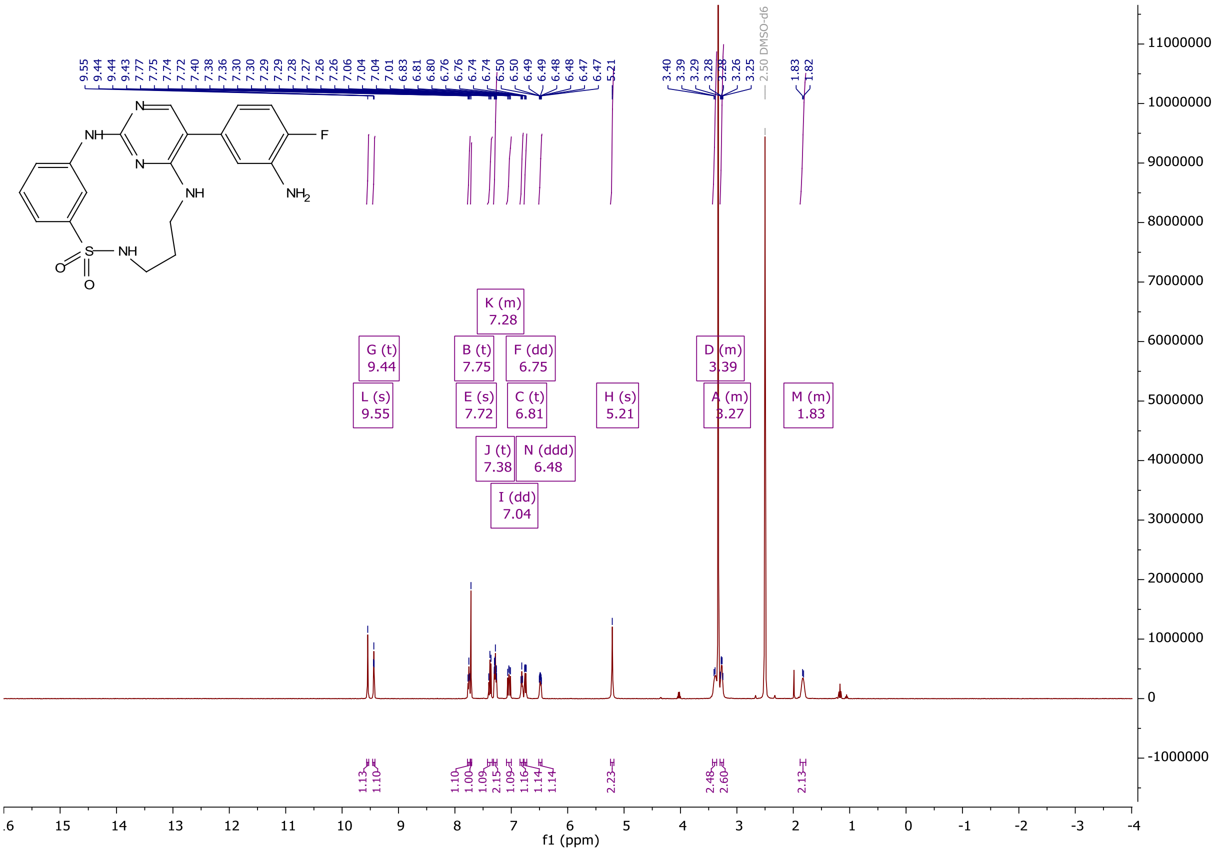

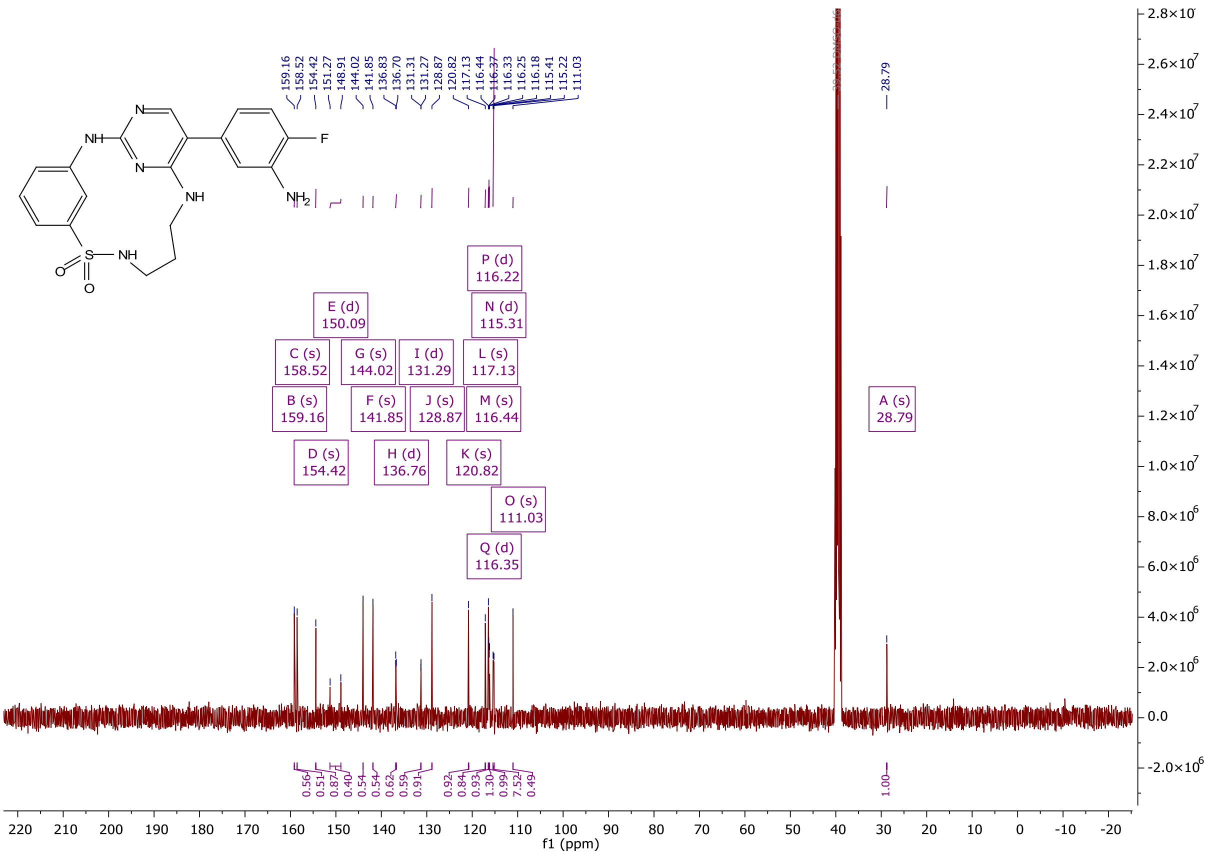
159.16
158.52
154.42
151.27
148.91
144.02
141.85
136.83
136.70
131.31
131.27
128.87
120.82
117.13
116.44
116.37
116.33
116.25
116.18
115.41
115.22
111.03
28.79
NH
N
N
NH
F
NH2
S
NH
O
O
P (d) 116.22
E (d) 150.09
N (d) 115.31
C (s) 158.52
G (s) 144.02
I (d) 131.29
L (s) 117.13
B (s) 159.16
F (s) 141.85
J (s) 128.87
M (s) 116.44
A (s) 28.79
D (s) 154.42
H (d) 136.76
K (s) 120.82
O (s) 111.03
Q (d) 116.35
f1 (ppm)

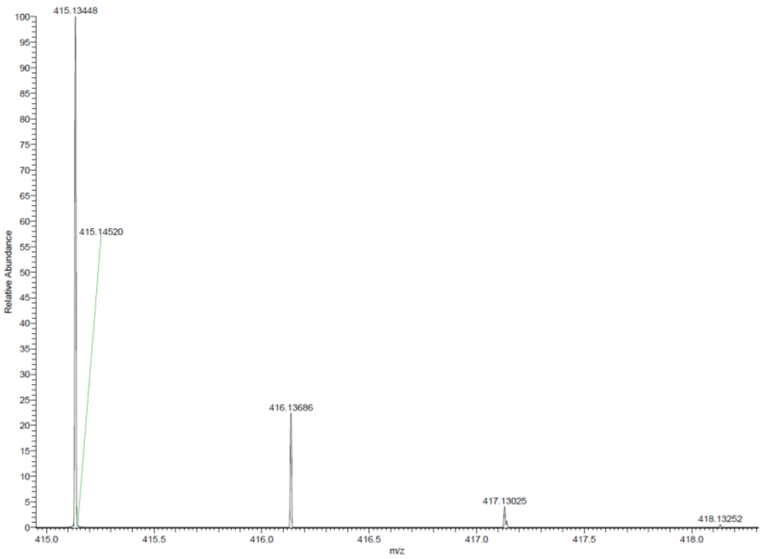
415.13448
415.14520
416.13686
417.13025
418.13252
Relative Abundance
m/z

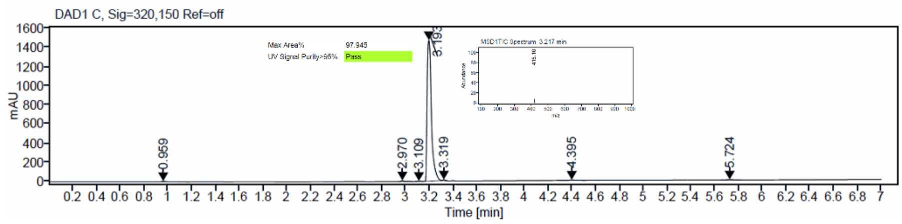
DAD1 C, Sig=320,150 Ref=off
Max Area% 97.945
UV Signal Purity>95% Pass
mAU
Time [min]
0.959
2.970
3.109
3.193
3.319
4.395
5.724

$^{1}H$ and $^{13}C$ {H} NMR, HRMS (Orbitrap) and HPLC/MS (ESI) data of compound **52**

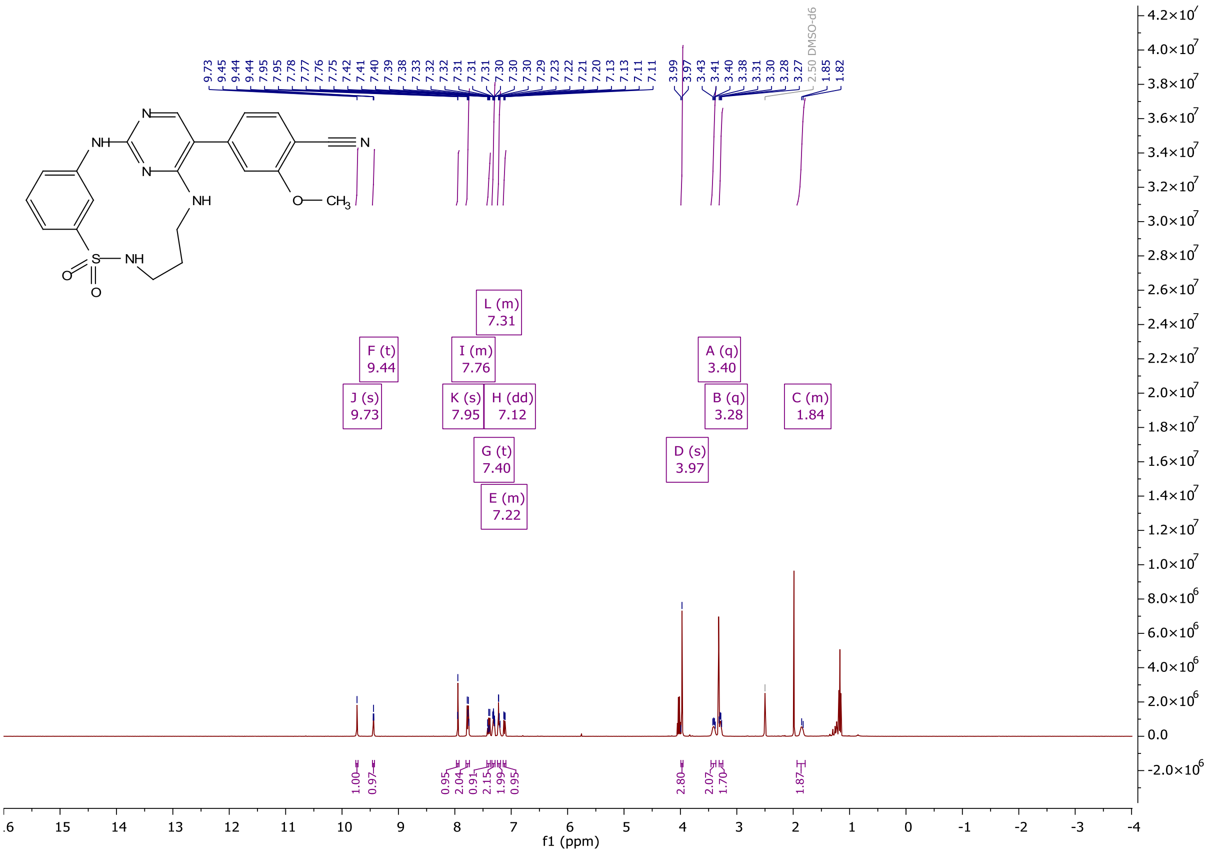

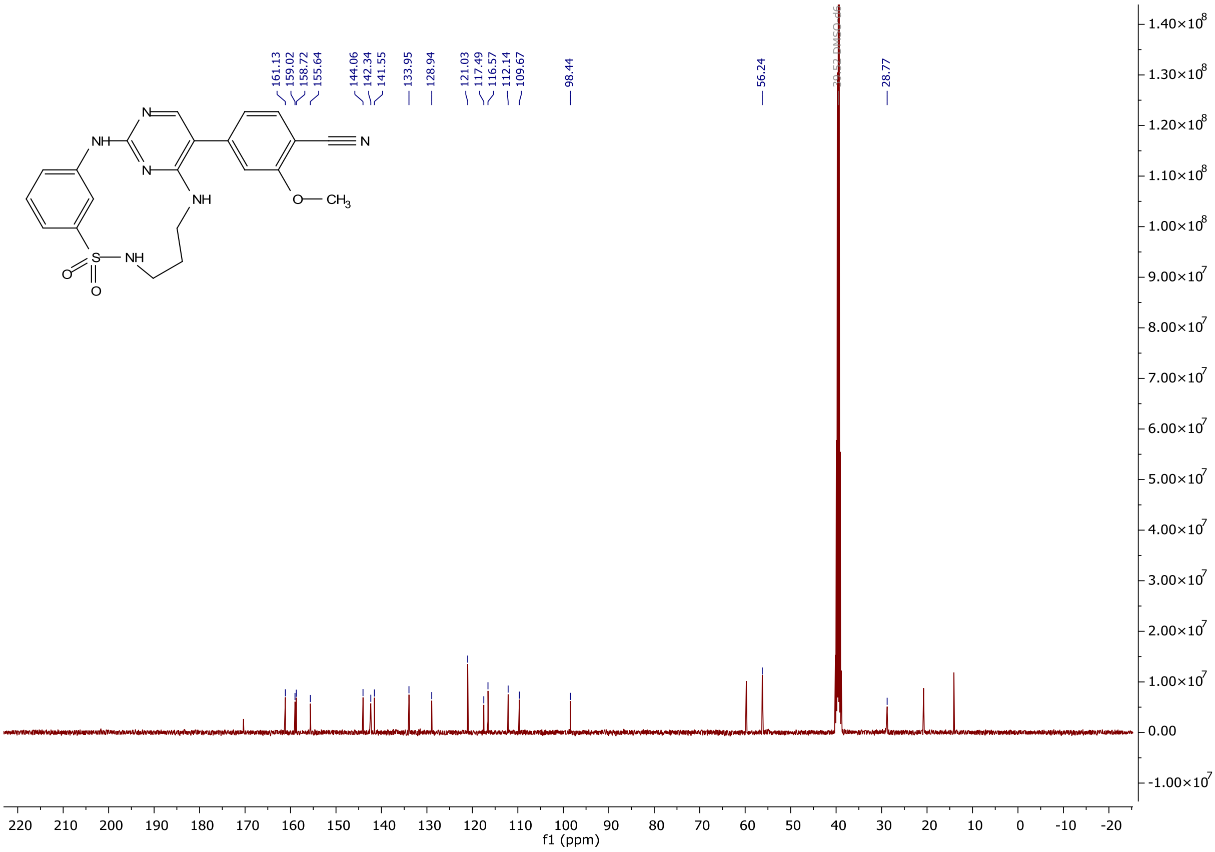

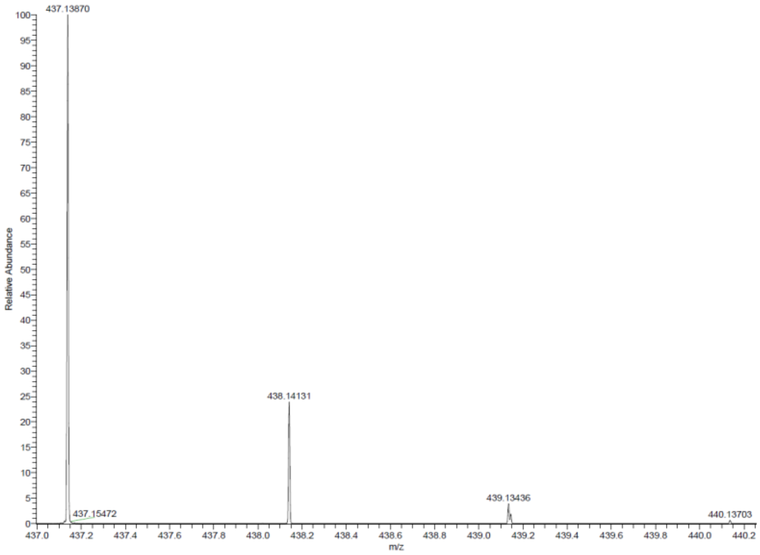

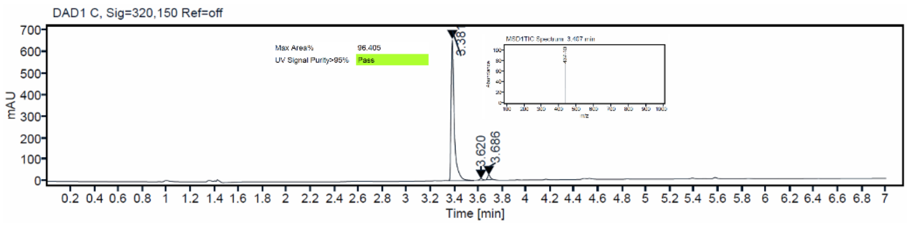

[1]H and [13]C {H} NMR, HRMS (oTOF) and HPLC/MS (ESI) data of compound **53**

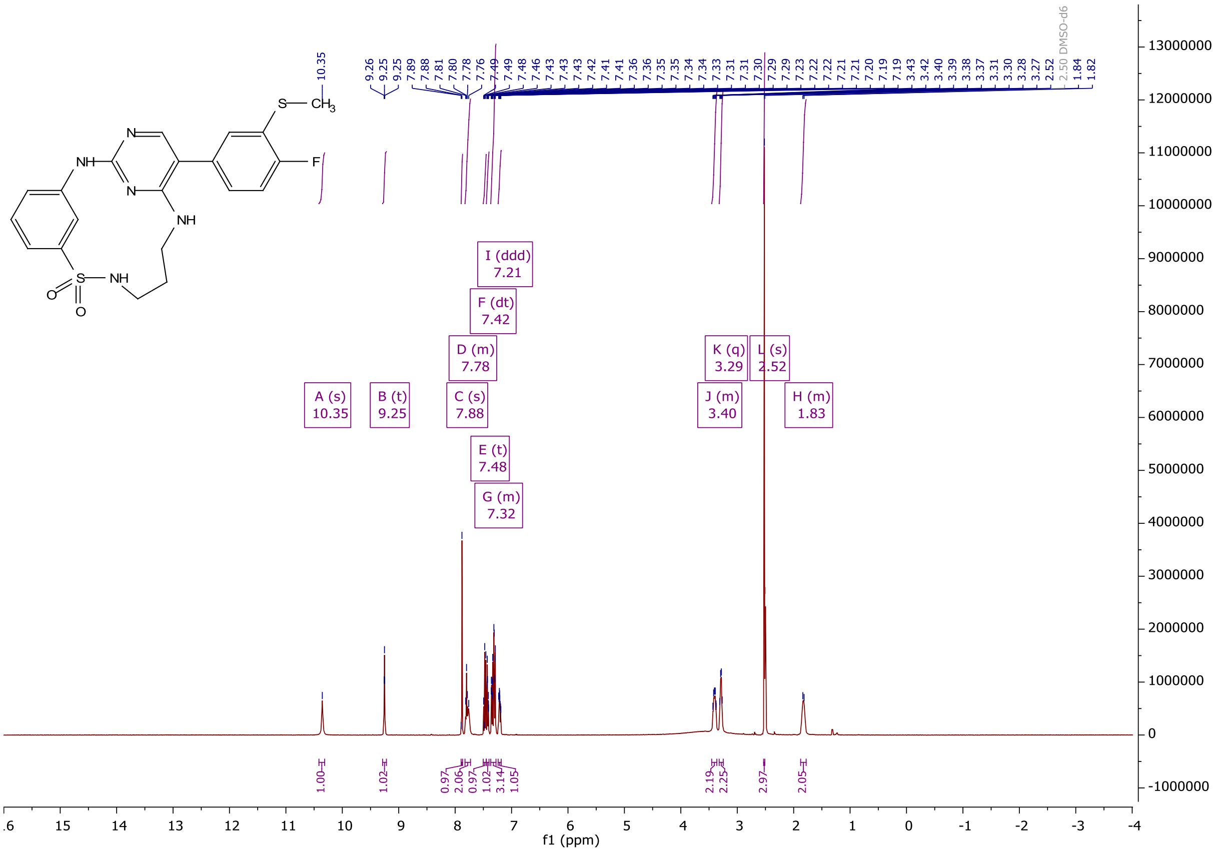

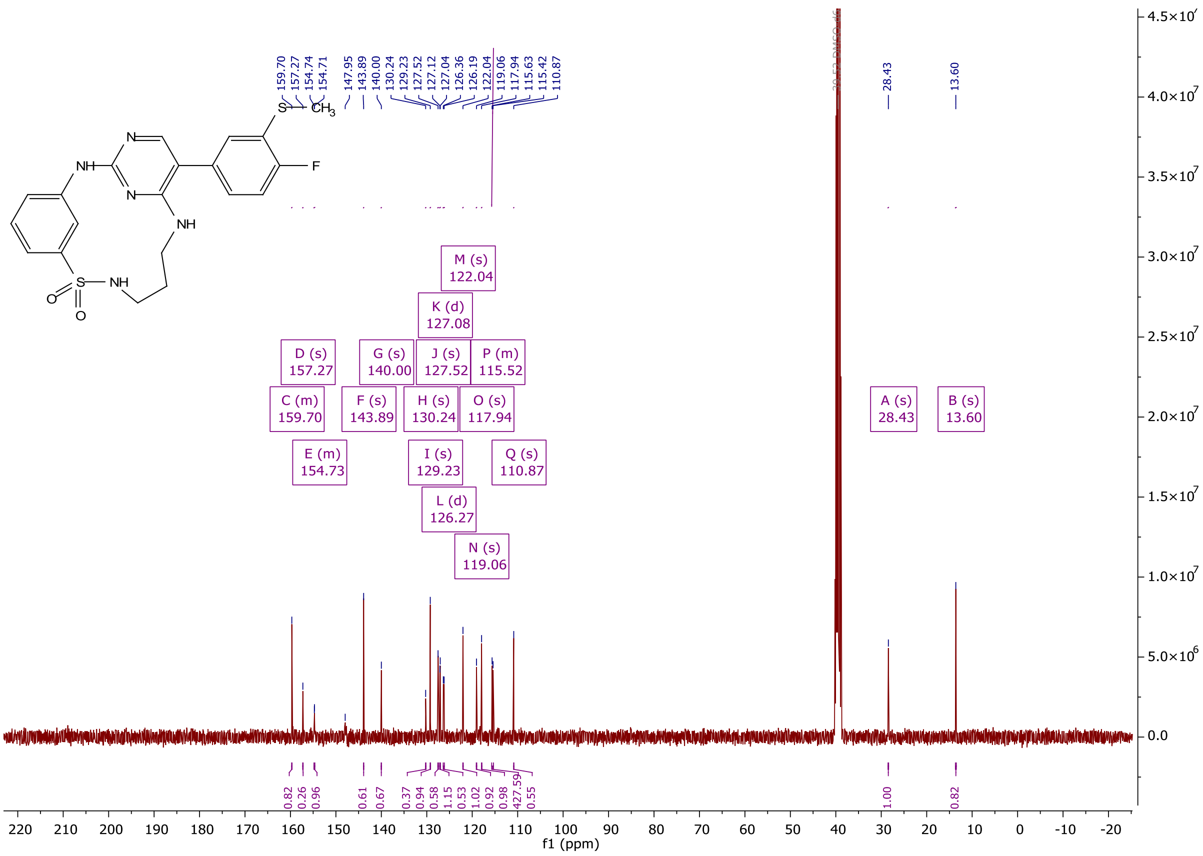

159.70
157.27
154.74
154.71
147.95
143.89
140.00
130.24
129.23
127.52
127.12
127.04
126.36
126.19
122.04
119.06
117.94
115.63
115.42
110.87
28.43
13.60
M (s) 122.04
K (d) 127.08
D (s) 157.27
G (s) 140.00
J (s) 127.52
P (m) 115.52
C (m) 159.70
F (s) 143.89
H (s) 130.24
O (s) 117.94
A (s) 28.43
B (s) 13.60
E (m) 154.73
I (s) 129.23
Q (s) 110.87
L (d) 126.27
N (s) 119.06
f1 (ppm)

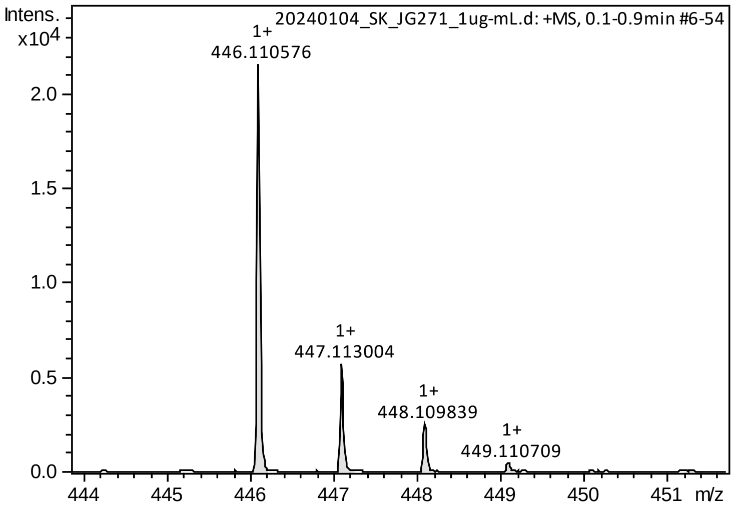

Intens.
x10^4
20240104_SK_JG271_1ug-mL.d: +MS, 0.1-0.9min #6-54
1+ 446.110576
1+ 447.113004
1+ 448.109839
1+ 449.110709
m/z

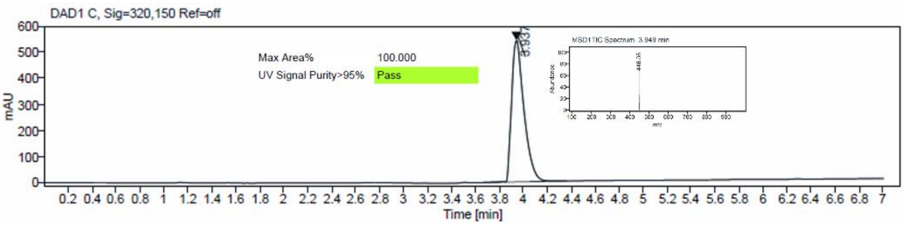

DAD1 C, Sig=320,150 Ref=off
Max Area%
100.000
UV Signal Purity>95%
Pass
mAU
Time [min]

<sup></sup>

[1]H and [13]C {H} NMR, HRMS (oTOF) and HPLC/MS (ESI) data of compound **54**

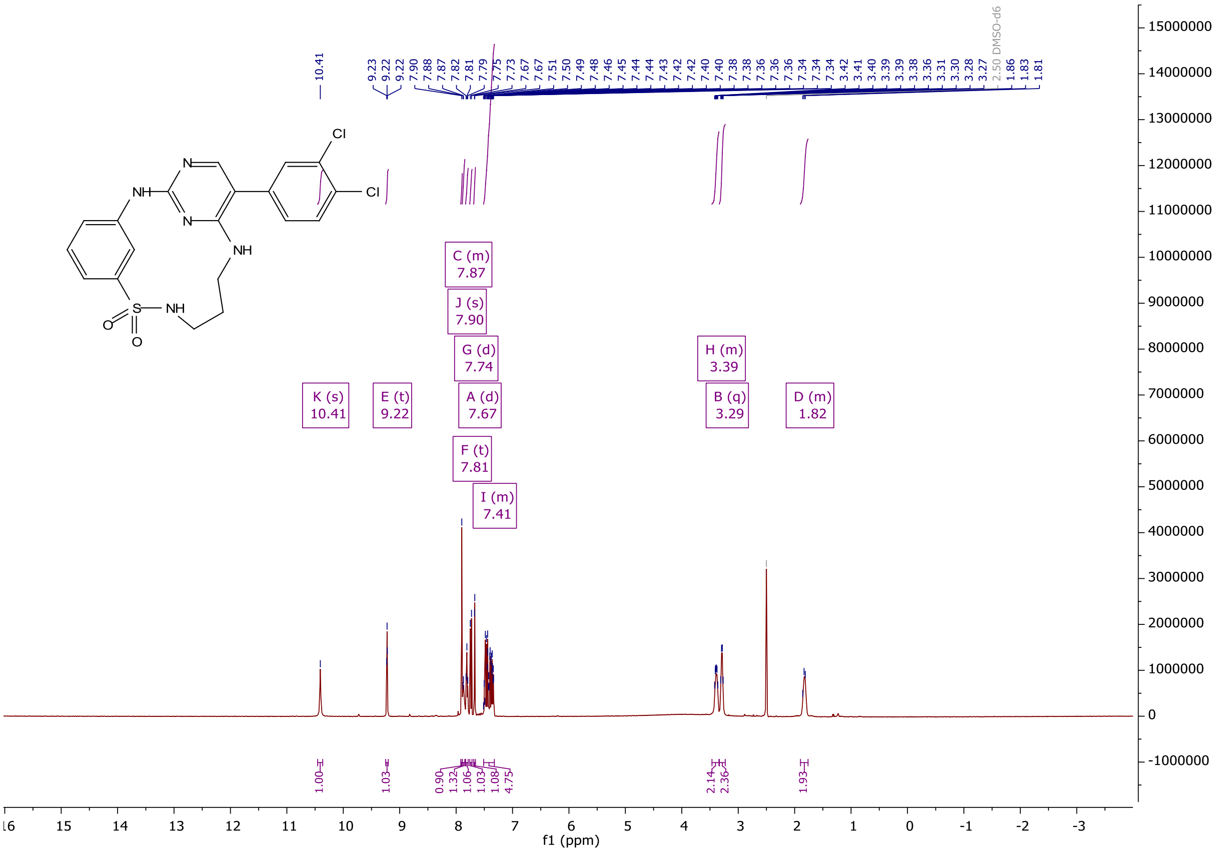

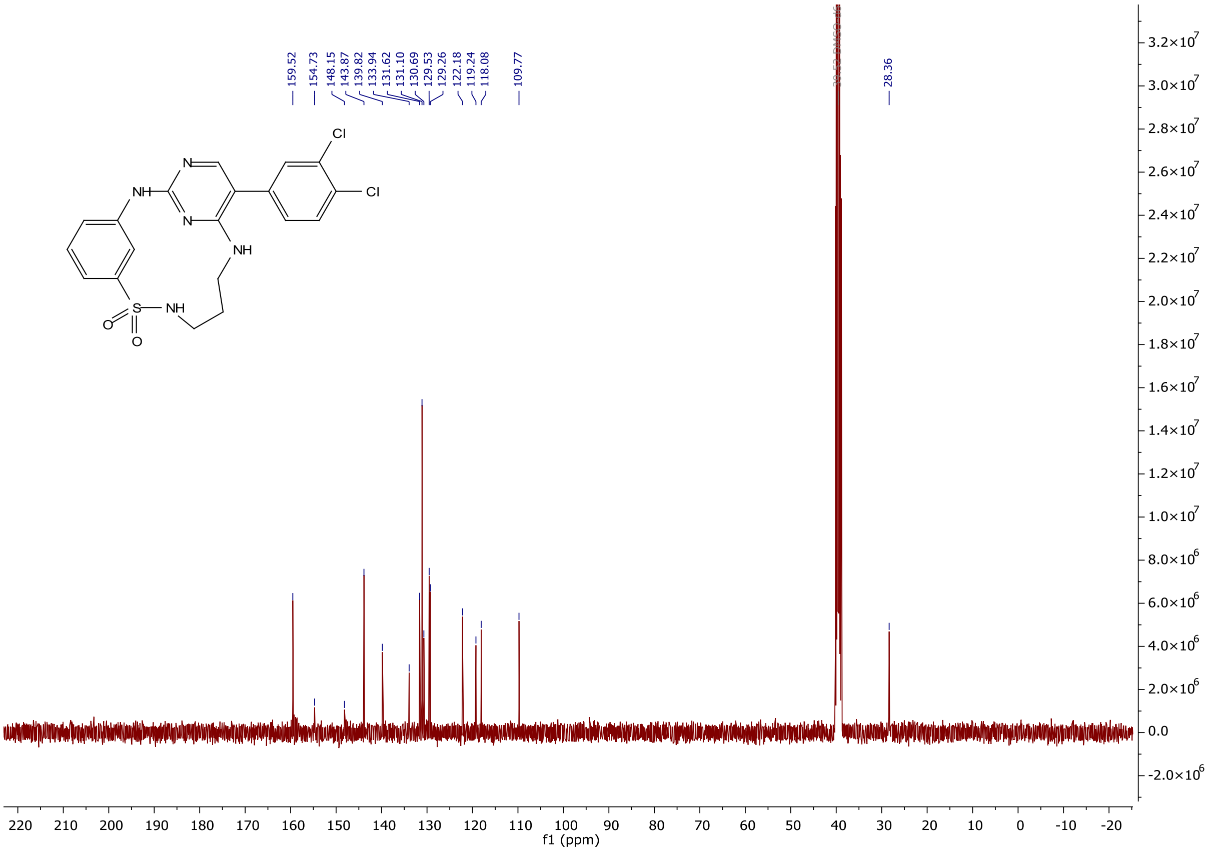

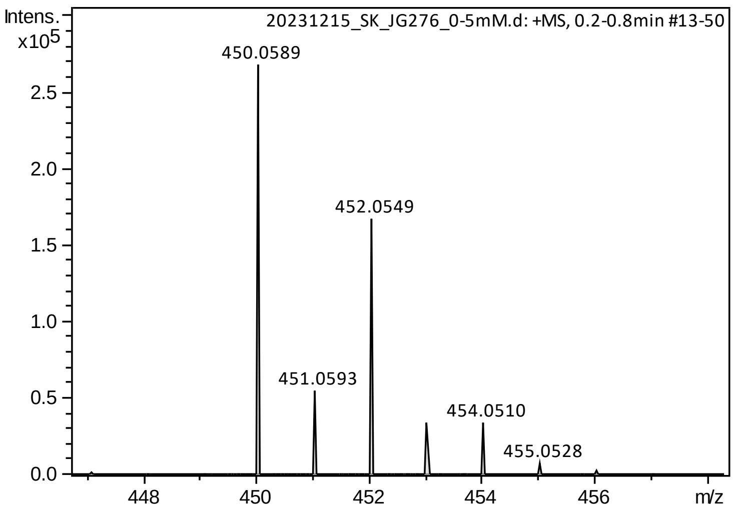

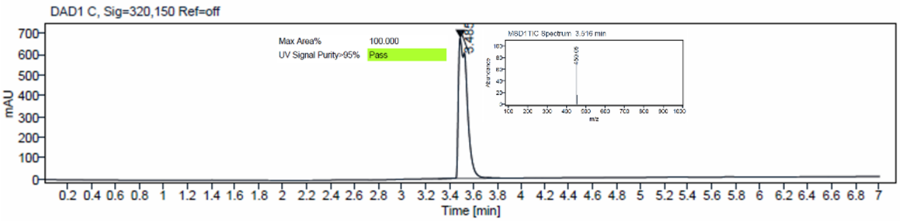

$^1$H and $^{13}$C {H} NMR, HRMS (oTOF) and HPLC/MS (ESI) data of compound **55**

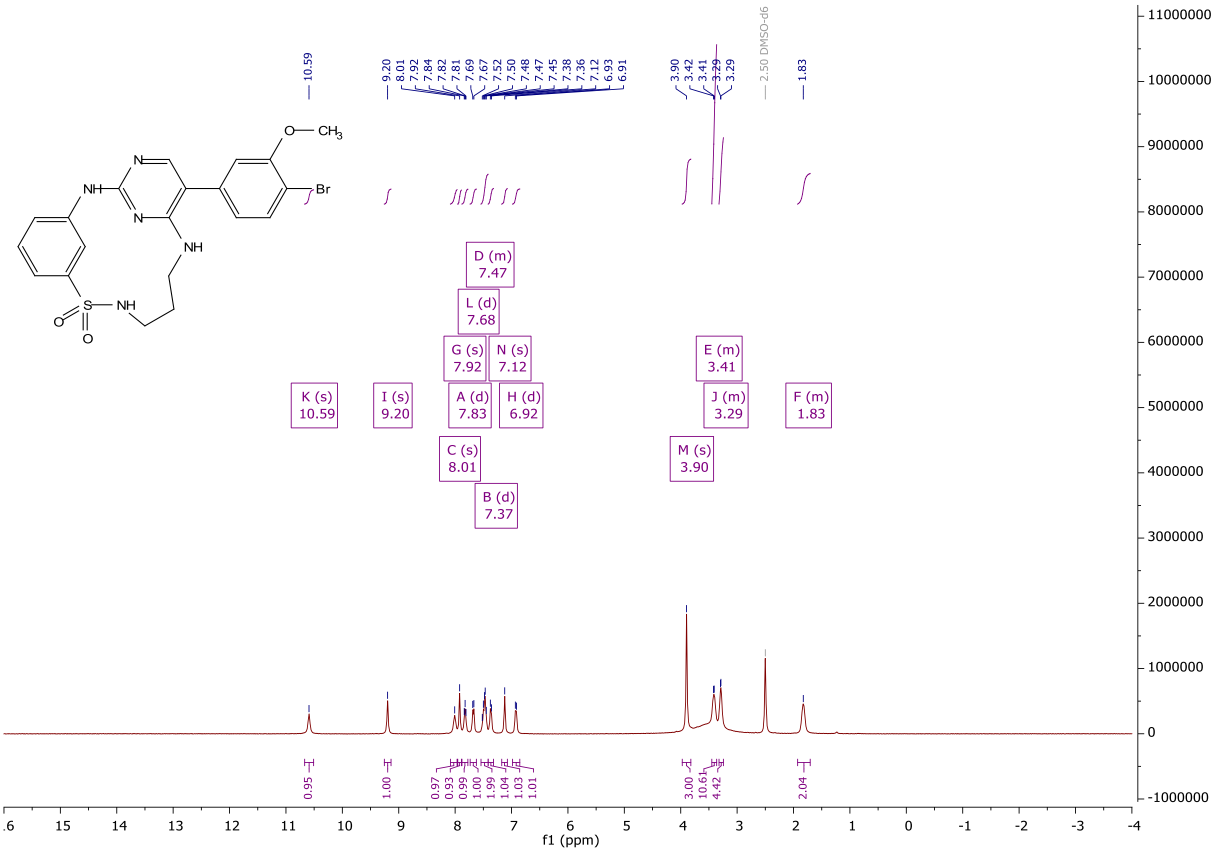

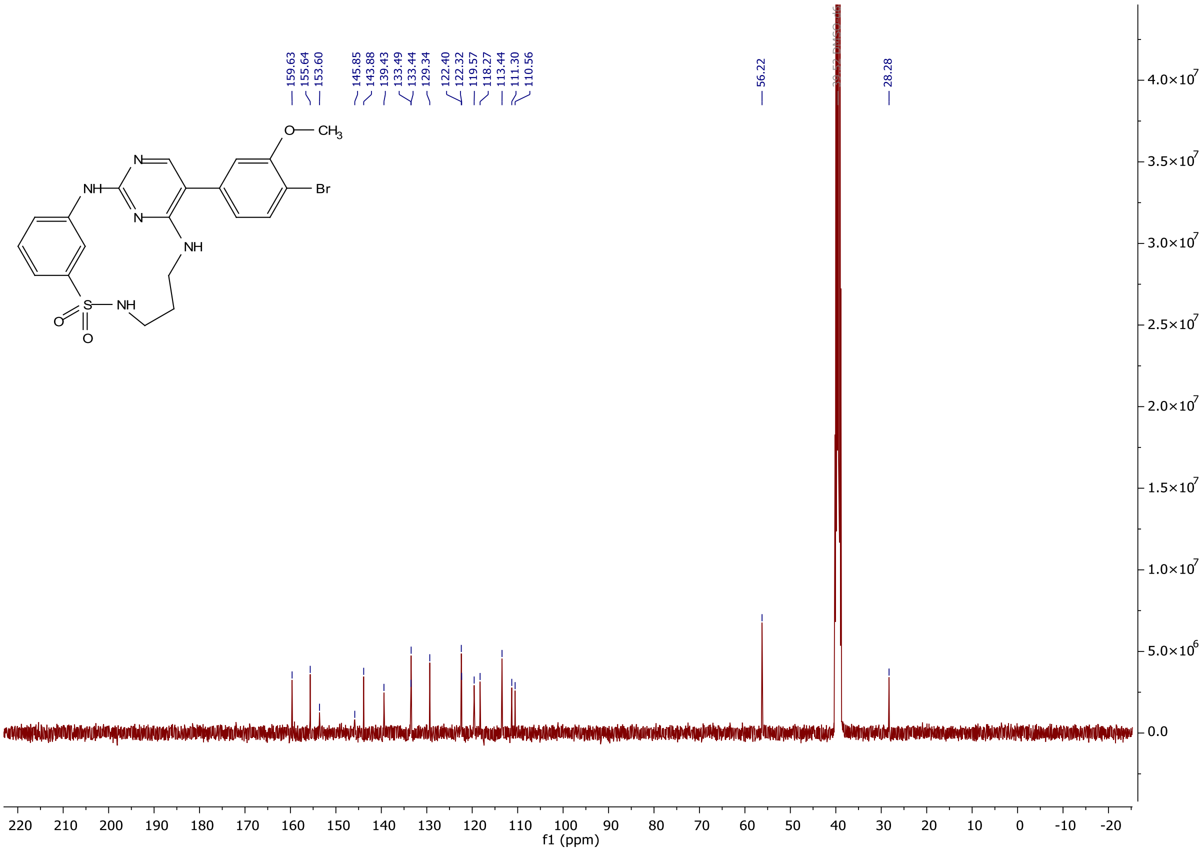
159.63
155.64
153.60
145.85
143.88
139.43
133.49
133.44
129.34
122.40
122.32
119.57
118.27
113.44
111.30
110.56
56.22
28.28
O— CH3
NH
N
N
NH
Br
S
NH
O
O
f1 (ppm)

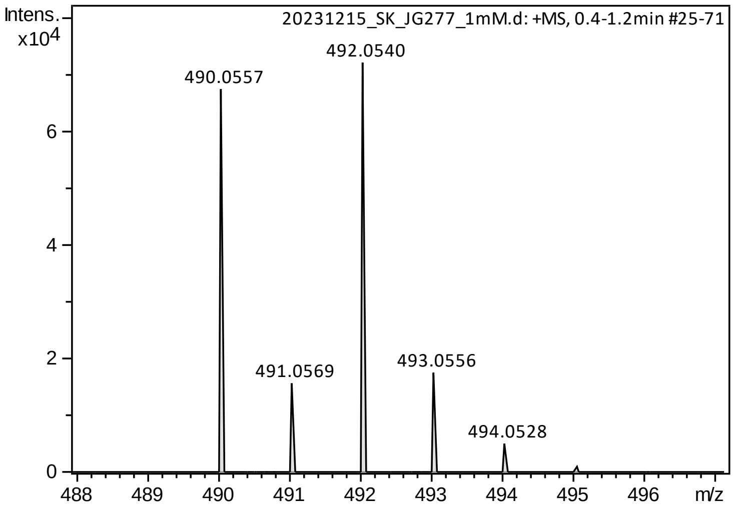
Intens.
x10^4
20231215_SK_JG277_1mM.d: +MS, 0.4-1.2min #25-71
490.0557
492.0540
491.0569
493.0556
494.0528
m/z

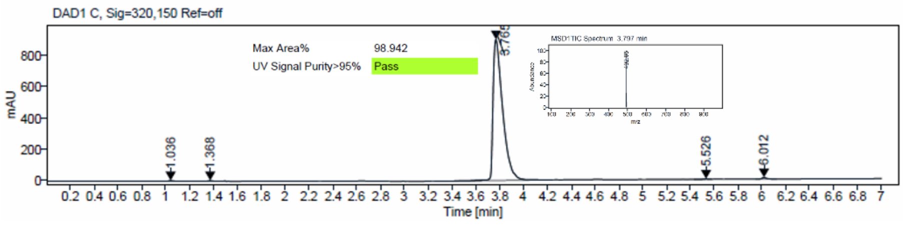
DAD1 C, Sig=320,150 Ref=off
Max Area%
98.942
UV Signal Purity>95%
Pass
mAU
Time [min]

$^{1}H$ and $^{13}C$ {H} NMR, HRMS (Orbitrap) and HPLC/MS (ESI) data of compound **56**

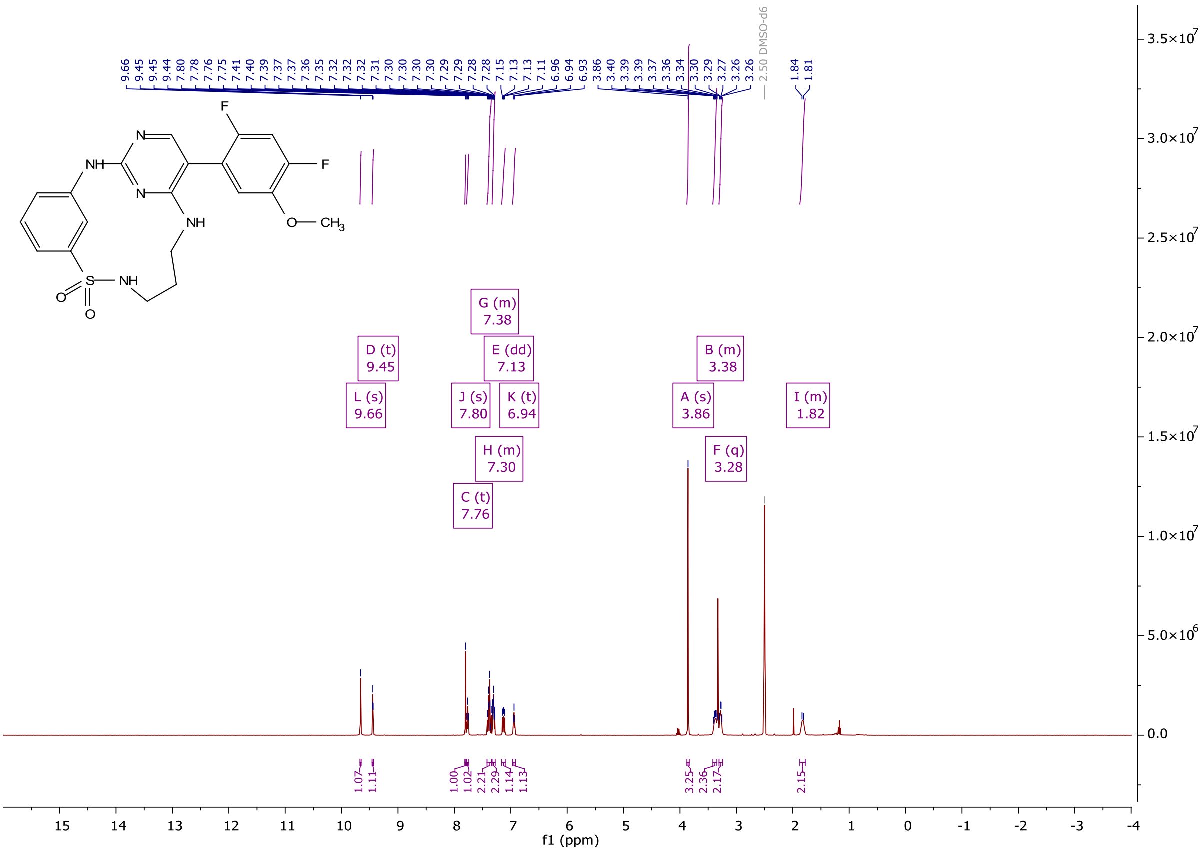

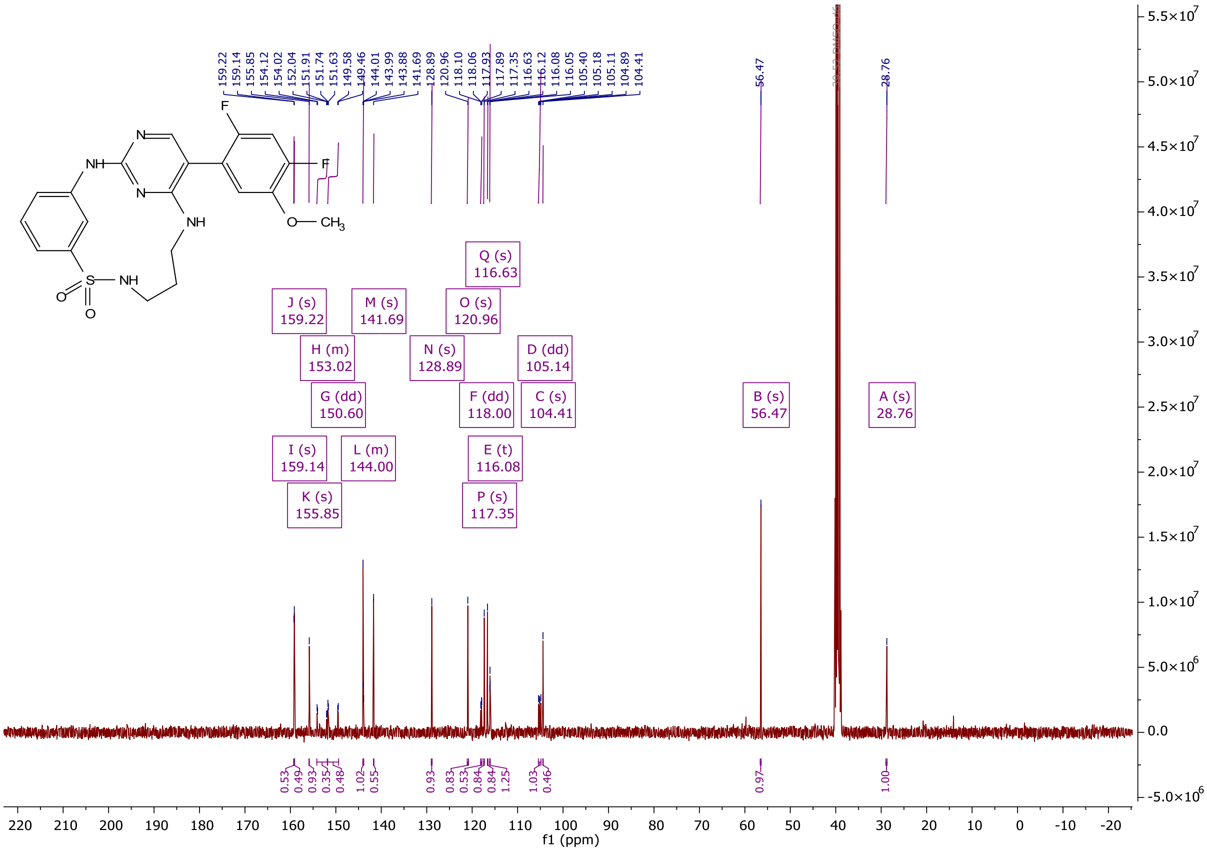

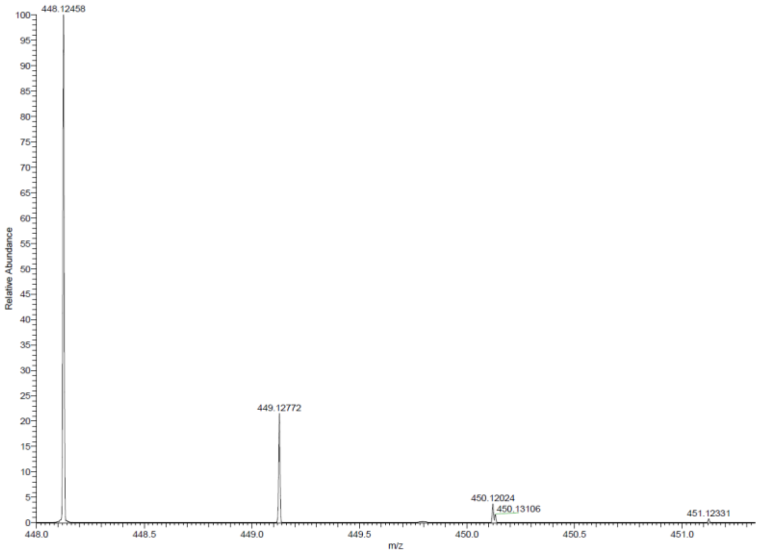

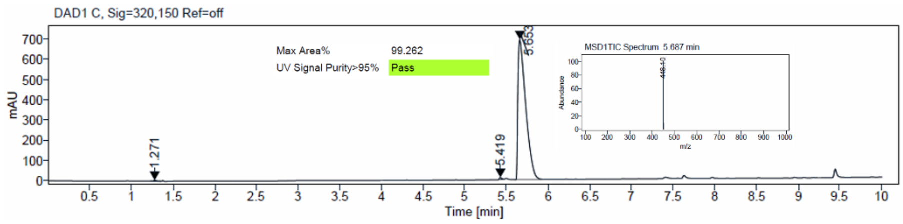

$^1$H and $^{13}$C {H} NMR, HRMS (Orbitrap) and HPLC/MS (ESI) data of compound **57**

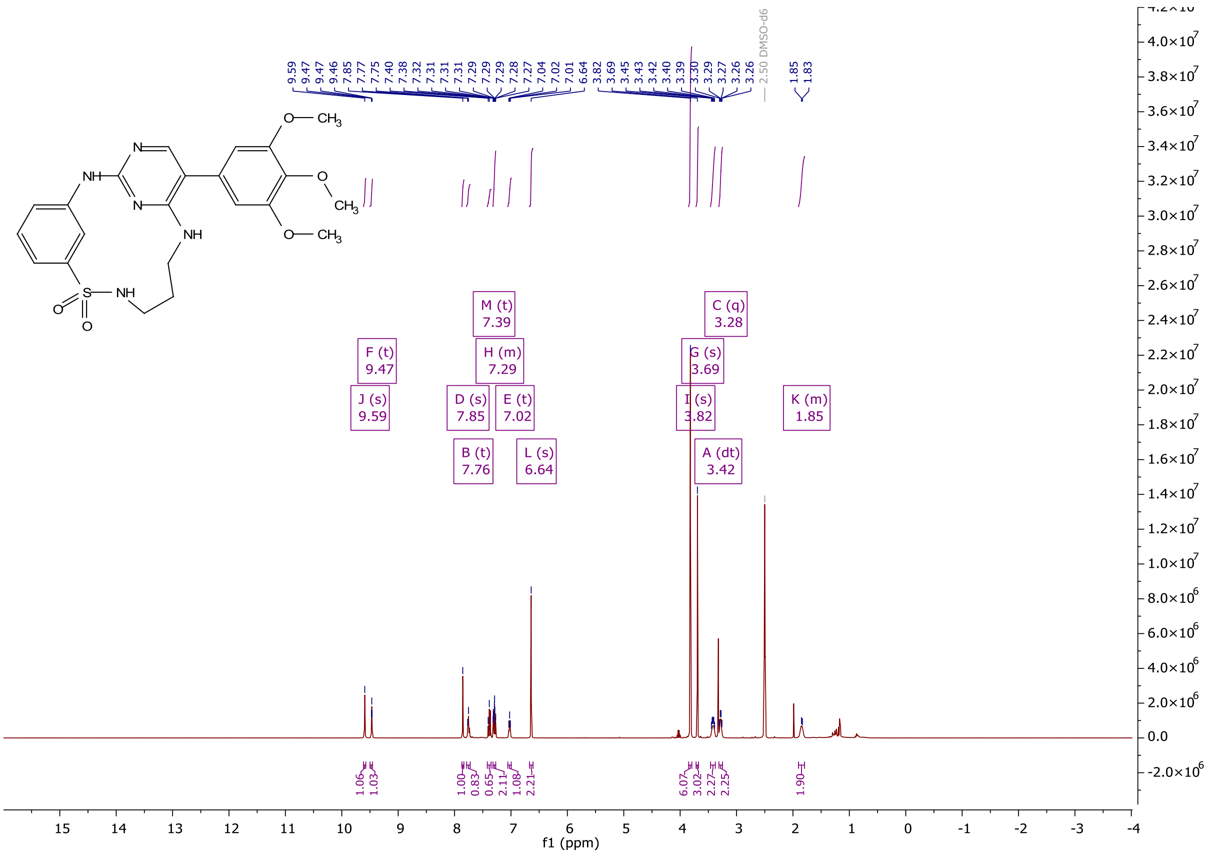

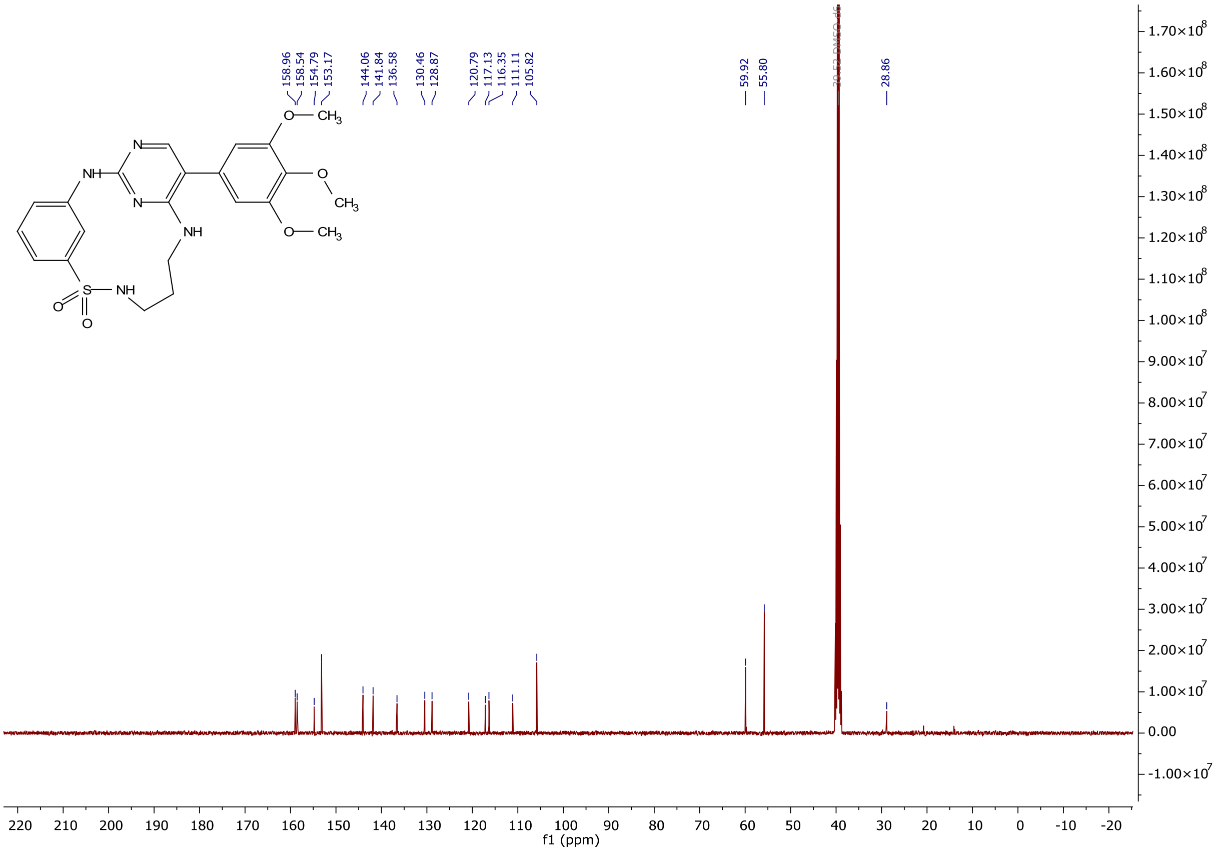
158.96
158.54
154.79
153.17
144.06
141.84
136.58
130.46
128.87
120.79
117.13
116.35
111.11
105.82
59.92
55.80
28.86
f1 (ppm)

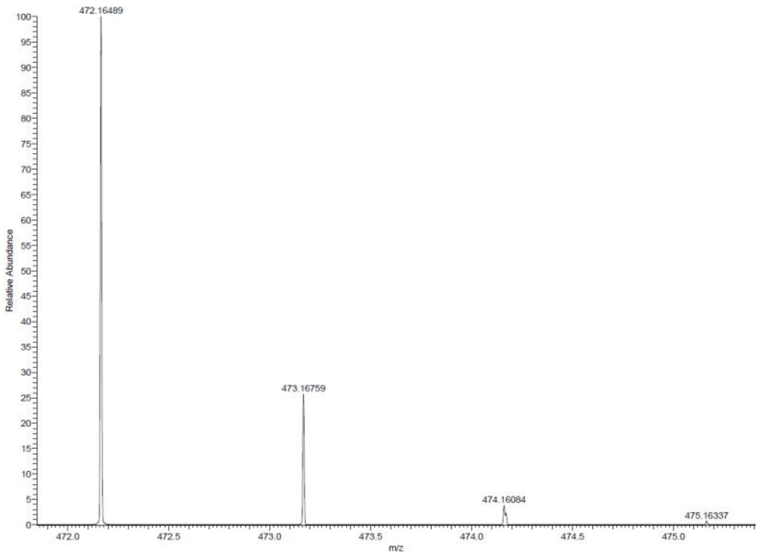
472.16489
473.16759
474.16084
475.16337
Relative Abundance
m/z

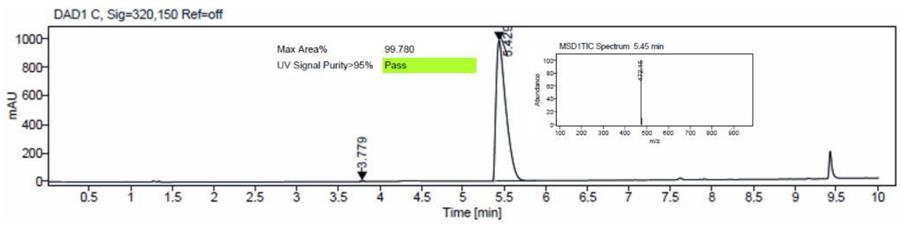
DAD1 C, Sig=320,150 Ref=off
Max Area%
99.780
UV Signal Purity>95%
Pass
MSD1TIC Spectrum 5.45 min
3.779
mAU
Time [min]

[1]H and [13]C {H} NMR, HRMS (Orbitrap) and HPLC/MS (ESI) data of compound **58**

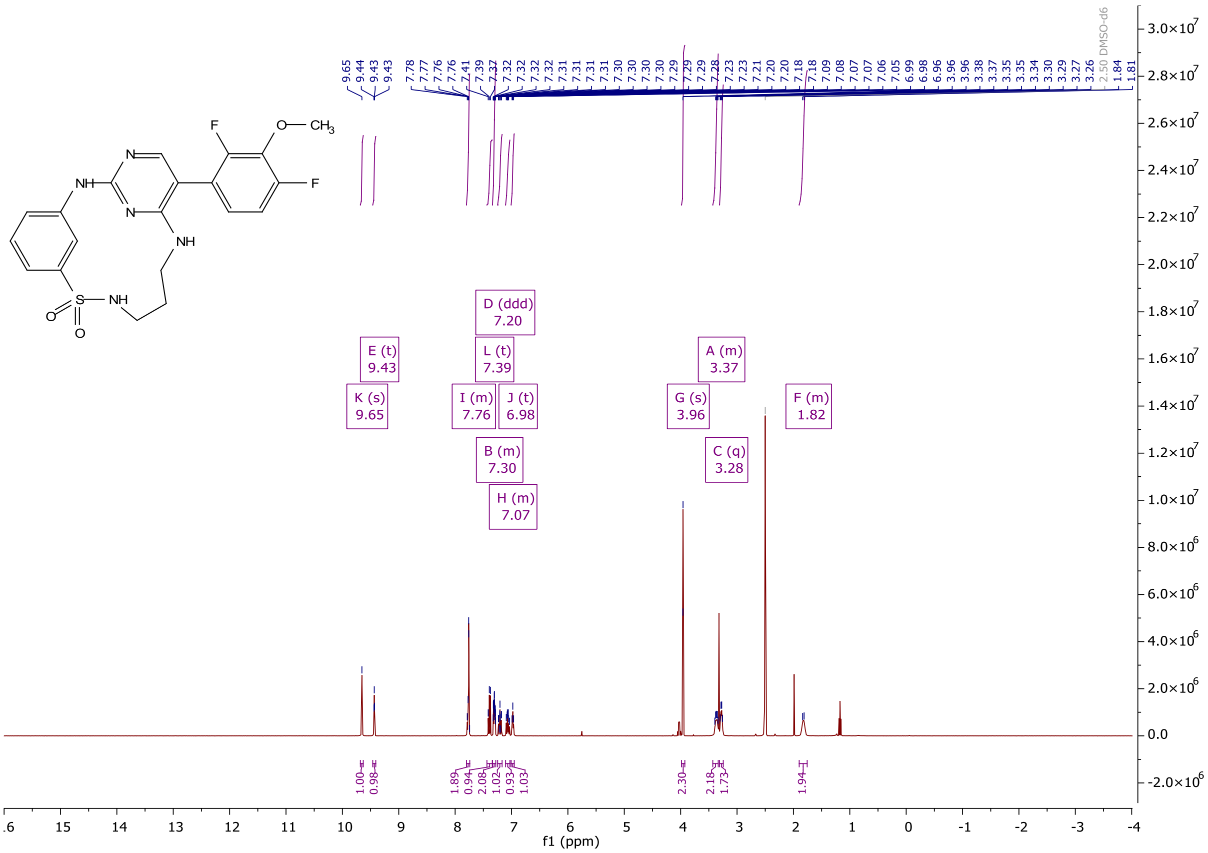

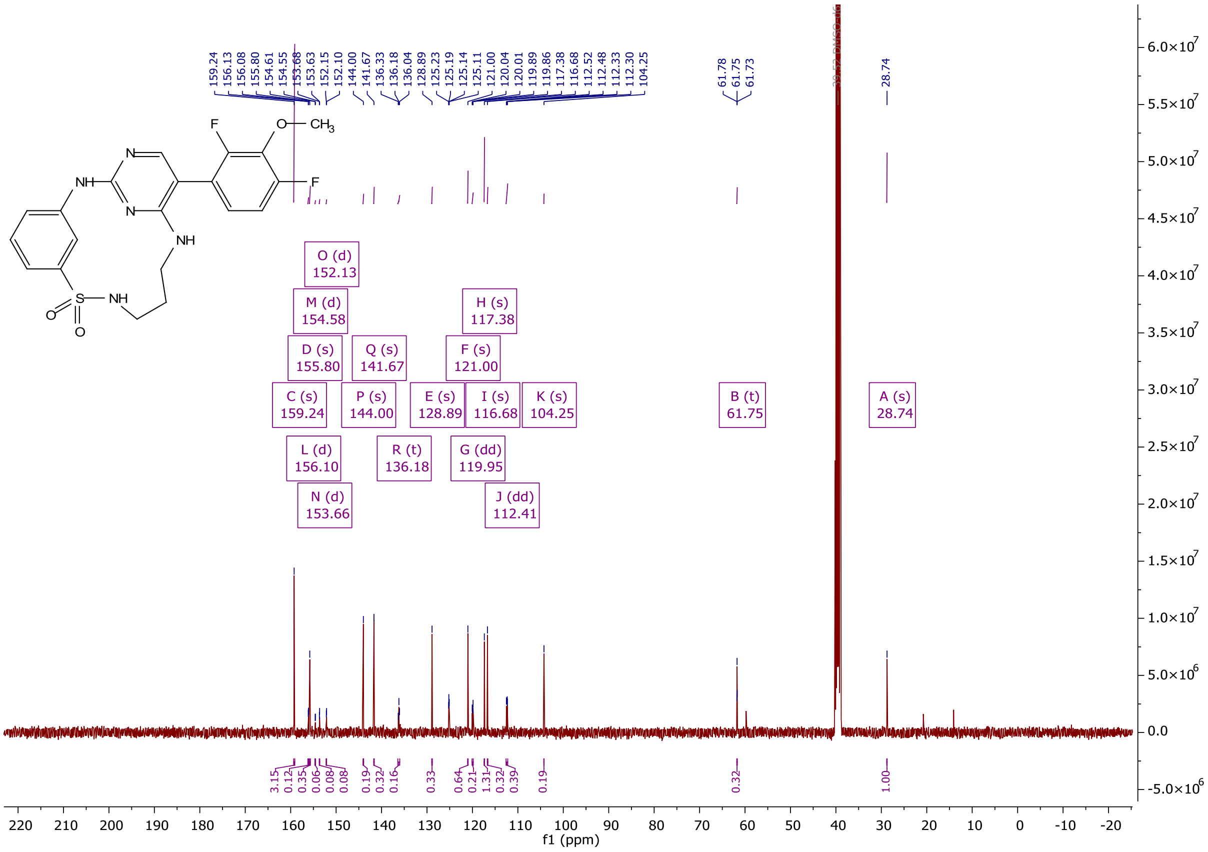

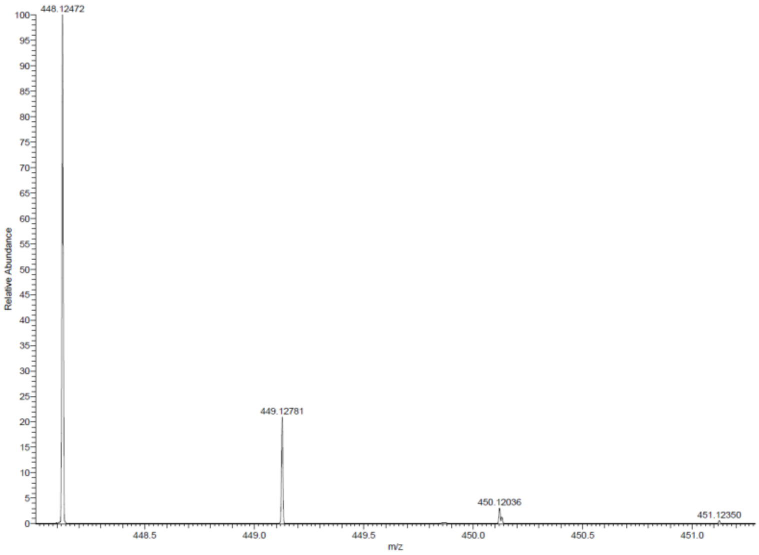

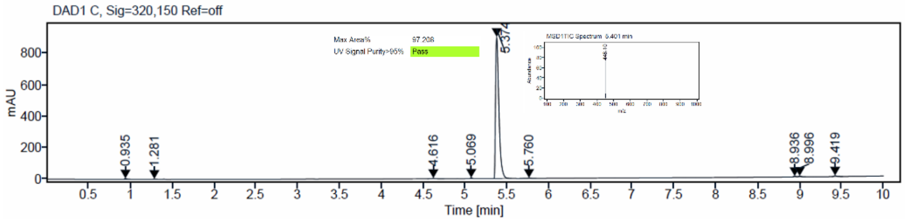

$^1$H and $^{13}$C {H} NMR, HRMS (Orbitrap) and HPLC/MS (ESI) data of compound **59**

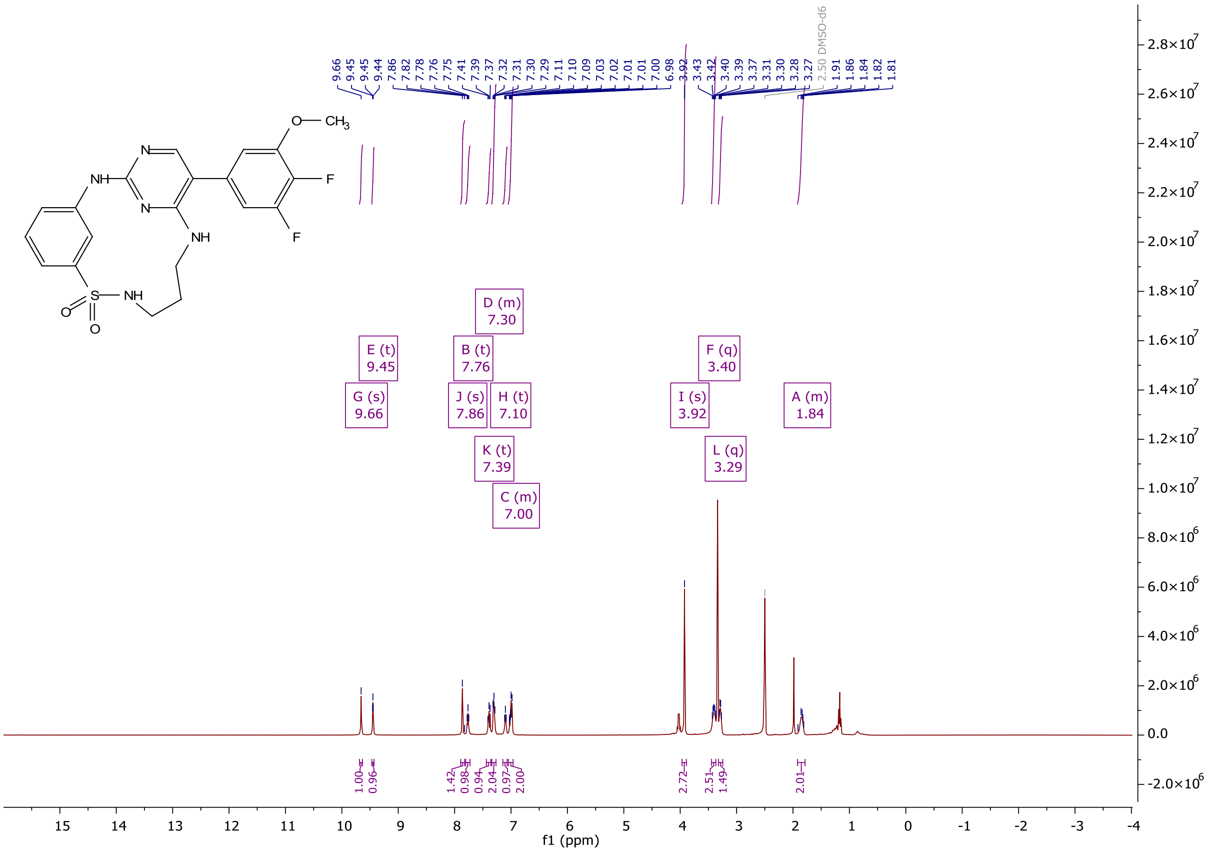

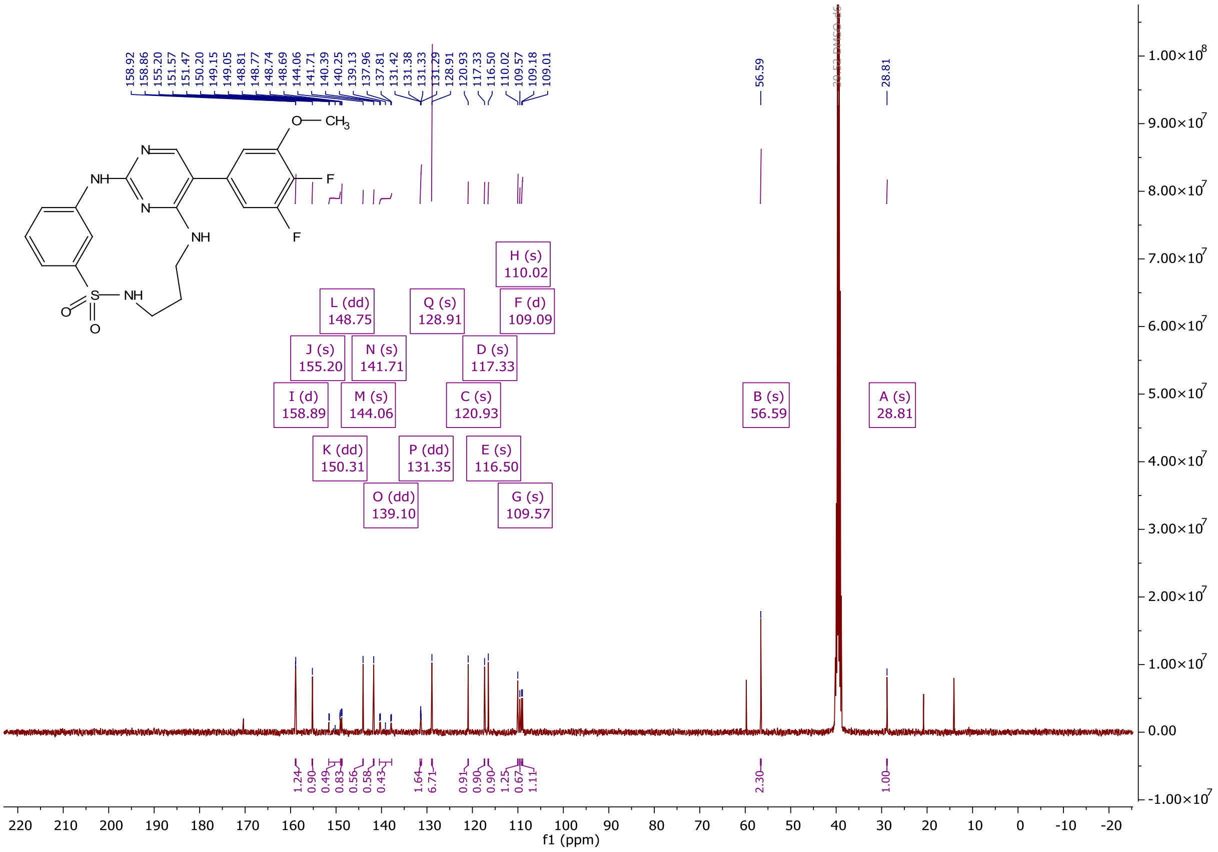

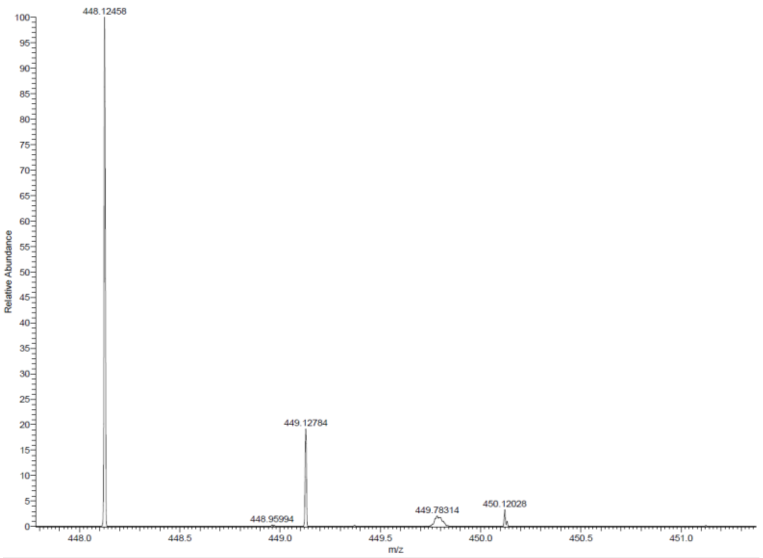

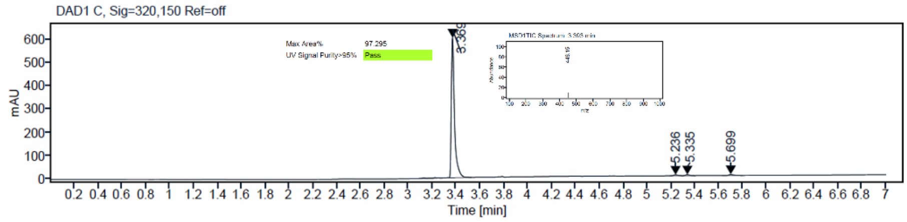

$^{1}H$ and $^{13}C$ {H} NMR, HRMS (Orbitrap) and HPLC/MS (ESI) data of compound **60**

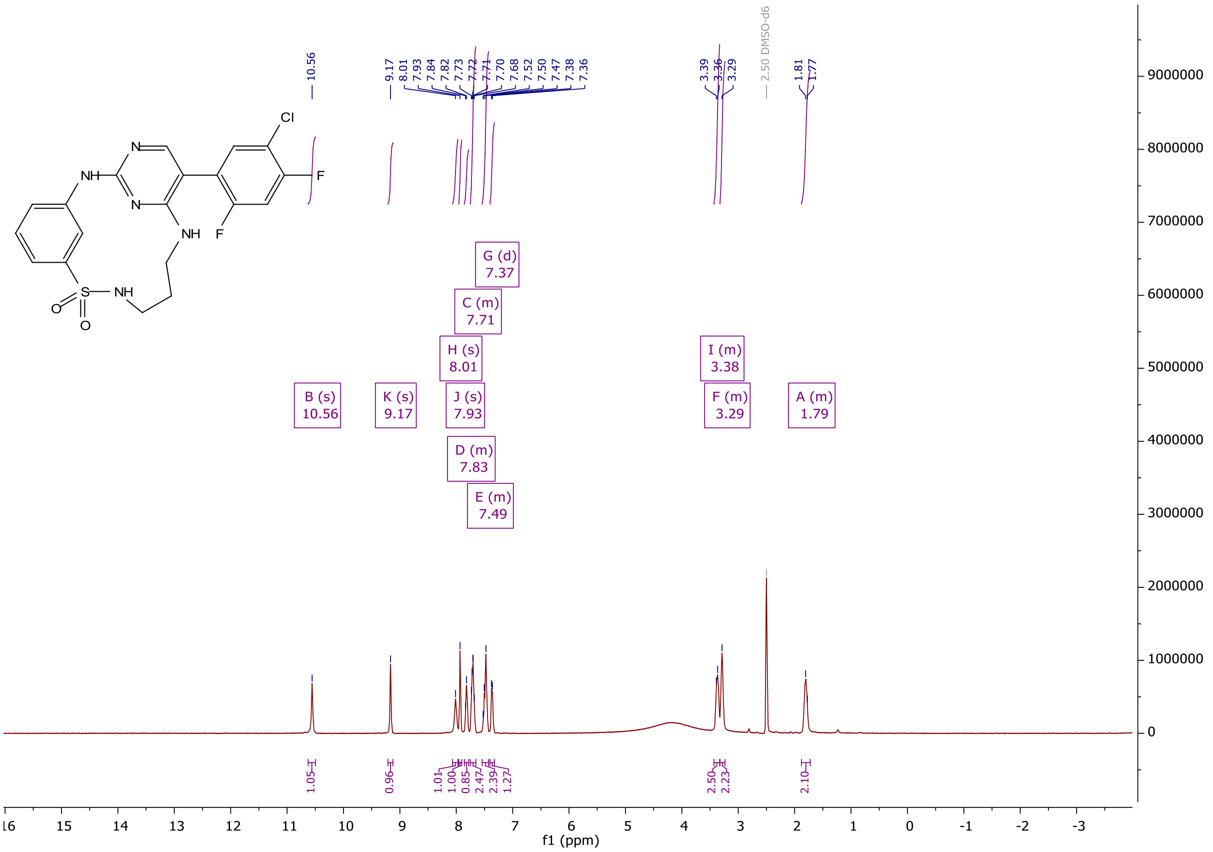

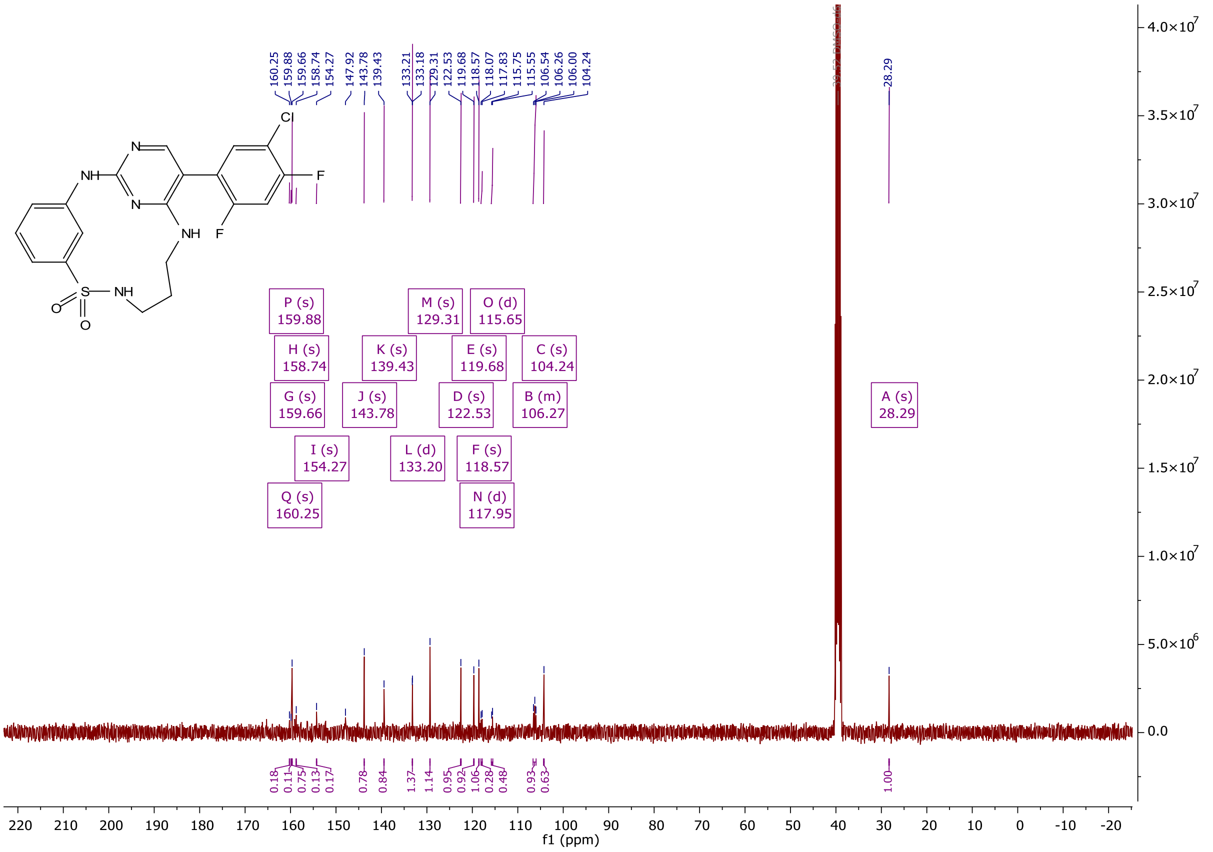

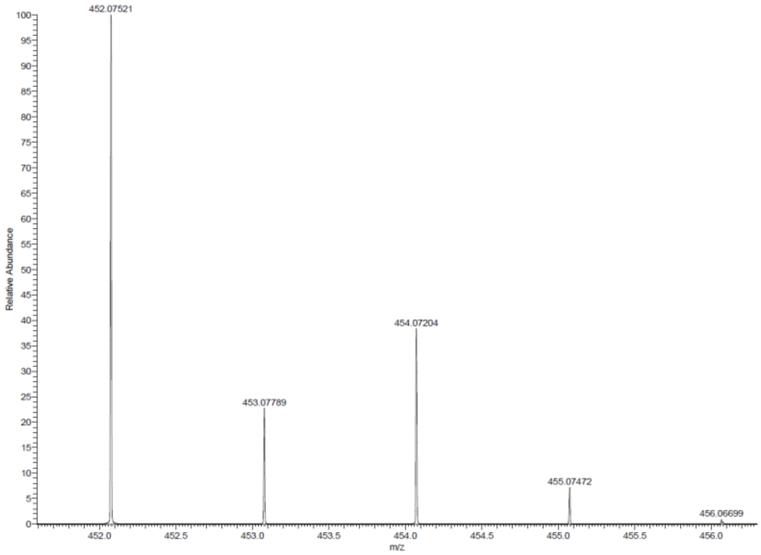

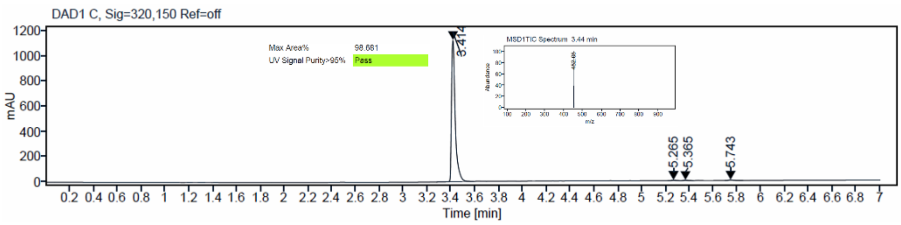

$^{1}$H and $^{13}$C {H} NMR, HRMS (oTOF) and HPLC/MS (ESI) data of compound **61**

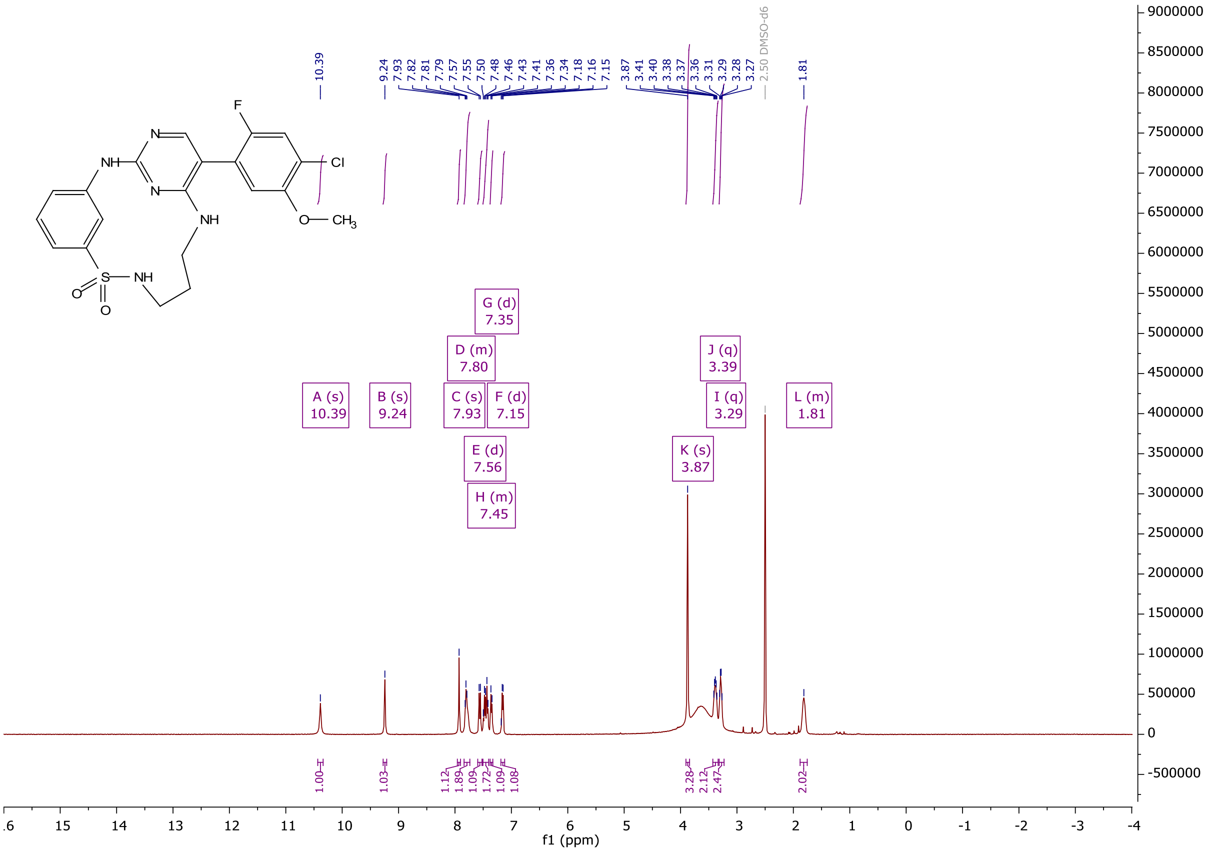

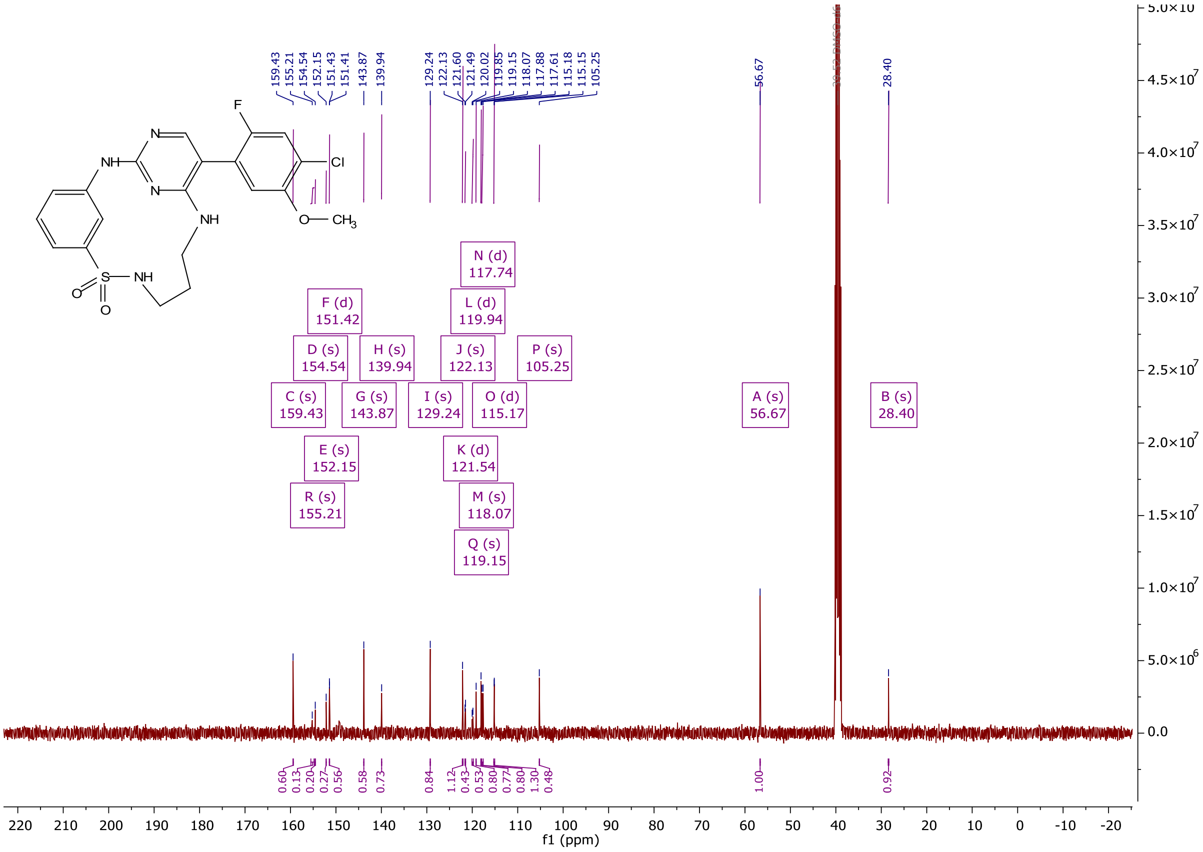
F
N
NH
N
NH
Cl
O— CH3
S— NH
O
O
C (s) 159.43
R (s) 155.21
D (s) 154.54
E (s) 152.15
F (d) 151.42
G (s) 143.87
H (s) 139.94
I (s) 129.24
J (s) 122.13
K (d) 121.54
L (d) 119.94
Q (s) 119.15
M (s) 118.07
N (d) 117.74
O (d) 115.17
P (s) 105.25
A (s) 56.67
B (s) 28.40
f1 (ppm)

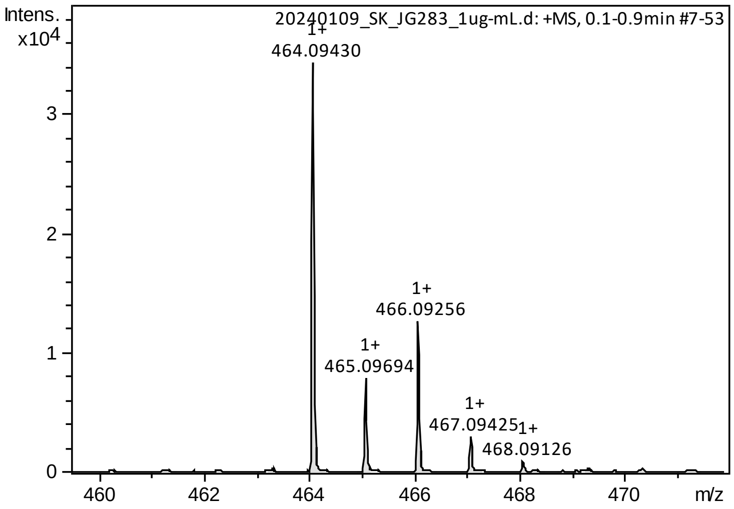
Intens.
x10^4
20240109_SK_JG283_1ug-mL.d: +MS, 0.1-0.9min #7-53
1+ 464.09430
1+ 465.09694
1+ 466.09256
1+ 467.09425
1+ 468.09126
m/z

#### HPLC/ESI-MS

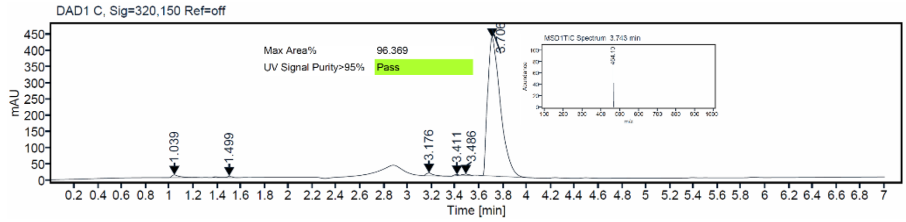
DAD1 C, Sig=320,150 Ref=off
Max Area% 96.369
UV Signal Purity>95% Pass
mAU
Time [min]

[1]H and [13]C {H} NMR, HRMS (oTOF) and HPLC/MS (ESI) data of compound **62**

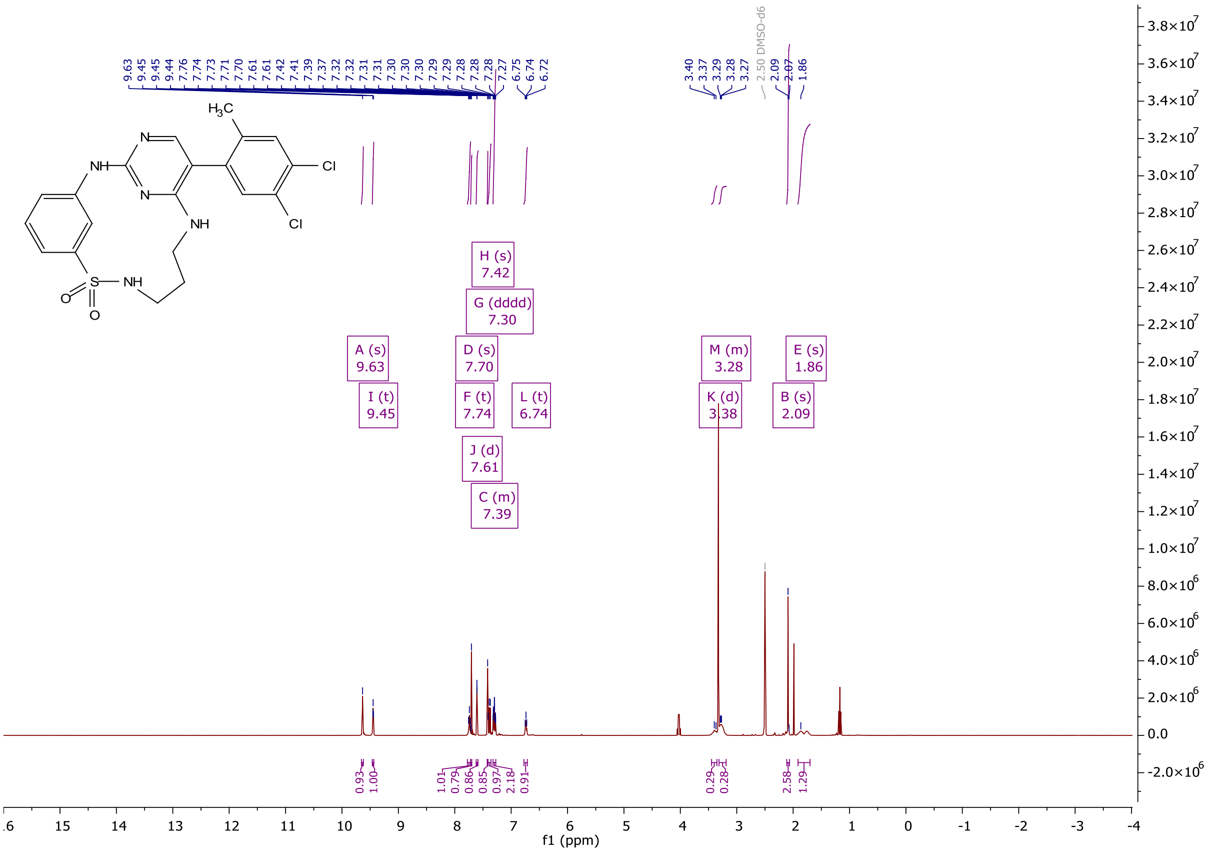

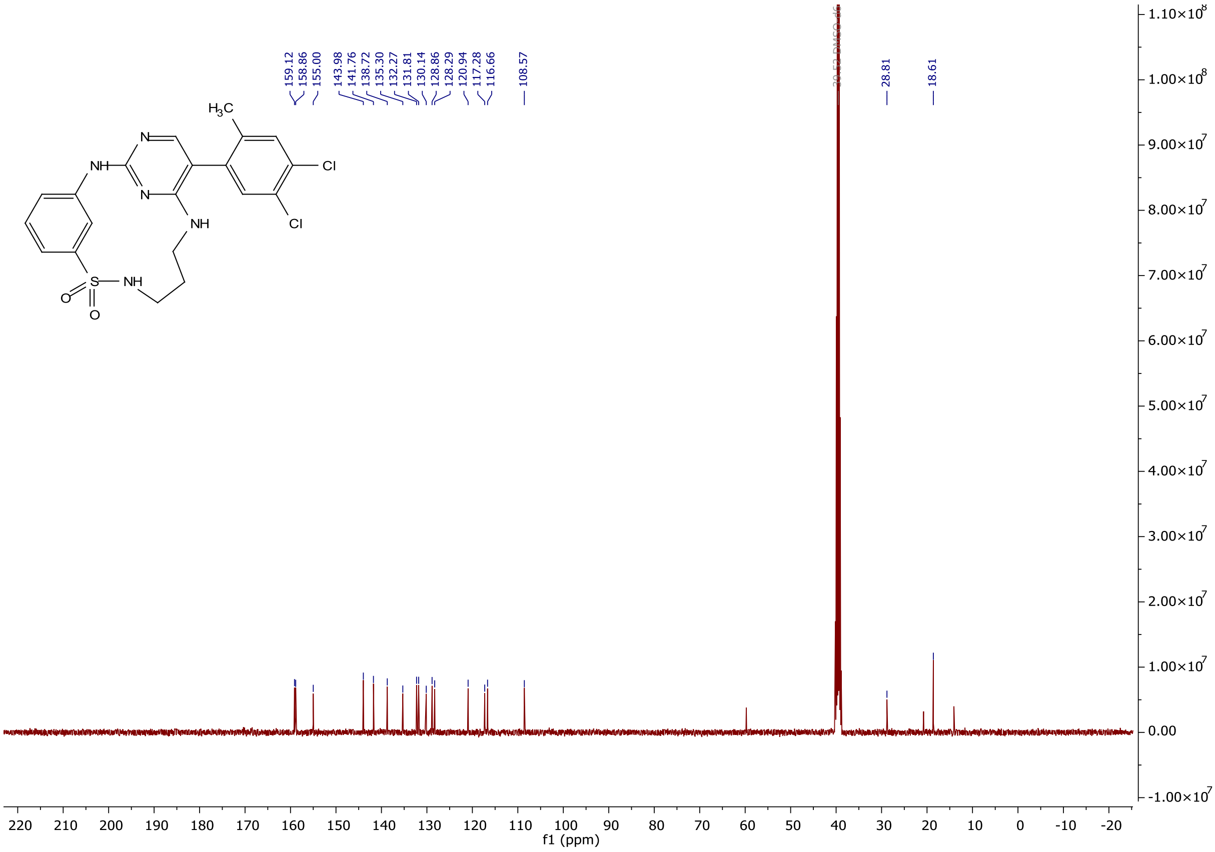

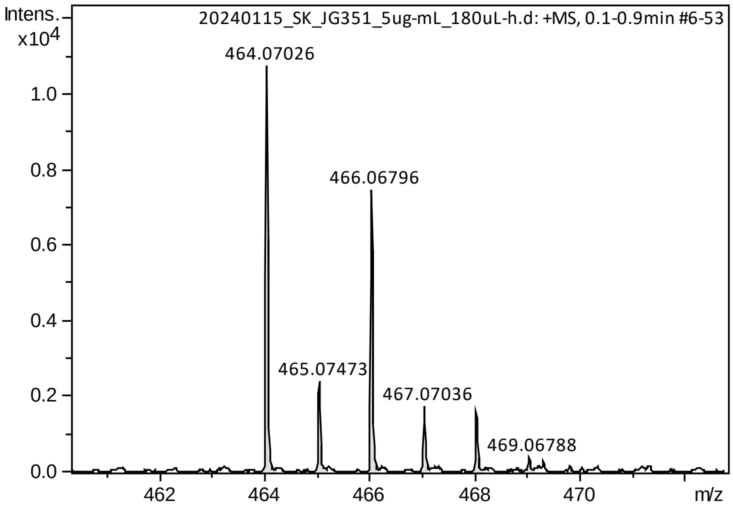

HPLC/MS (ESI)

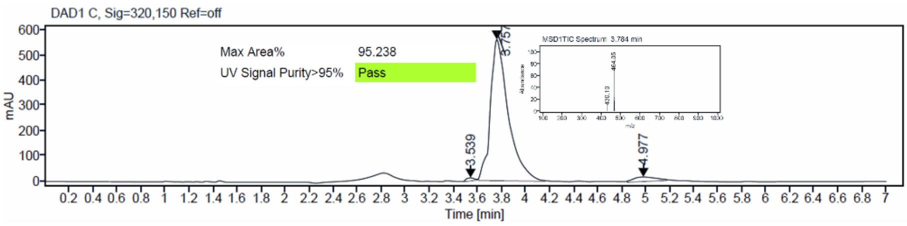

$^1$H and $^{13}$C {H} NMR, HRMS (oTOF) and HPLC/MS (ESI) data of compound **63**

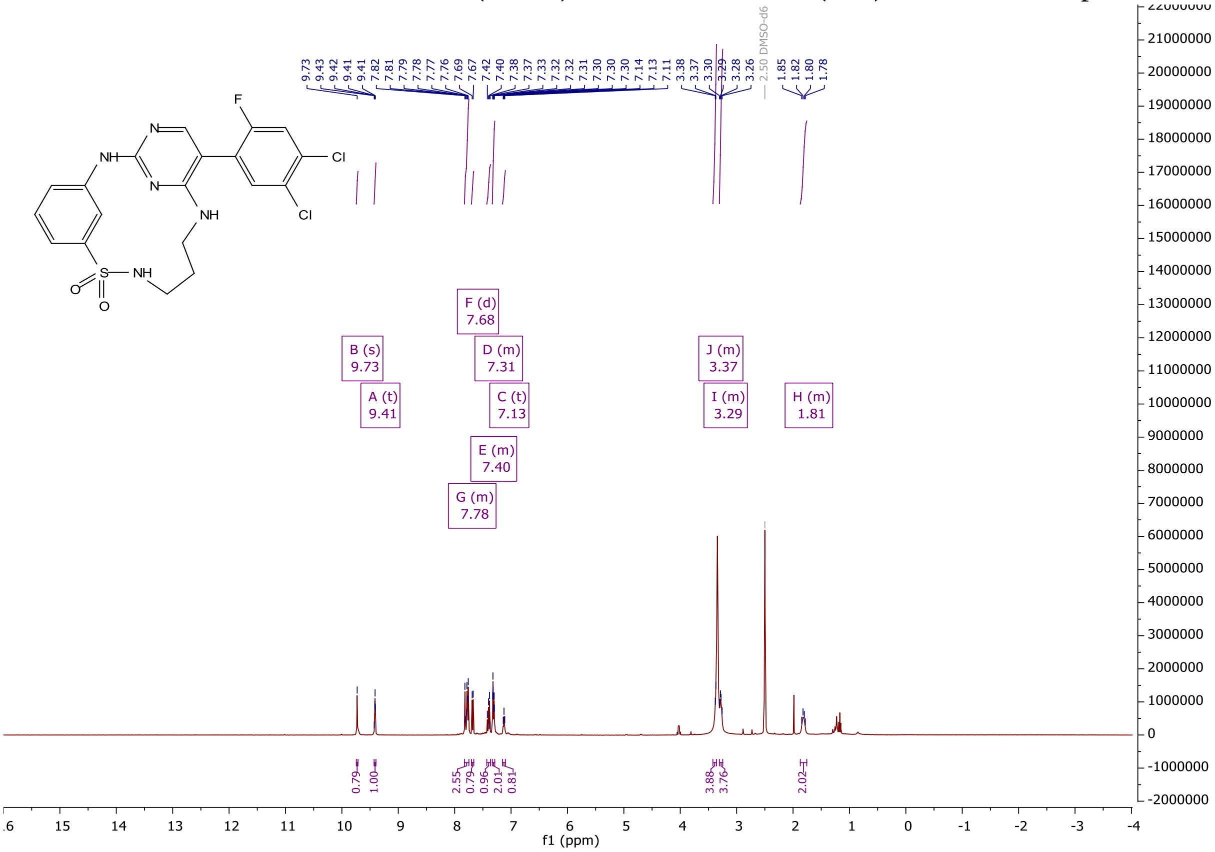

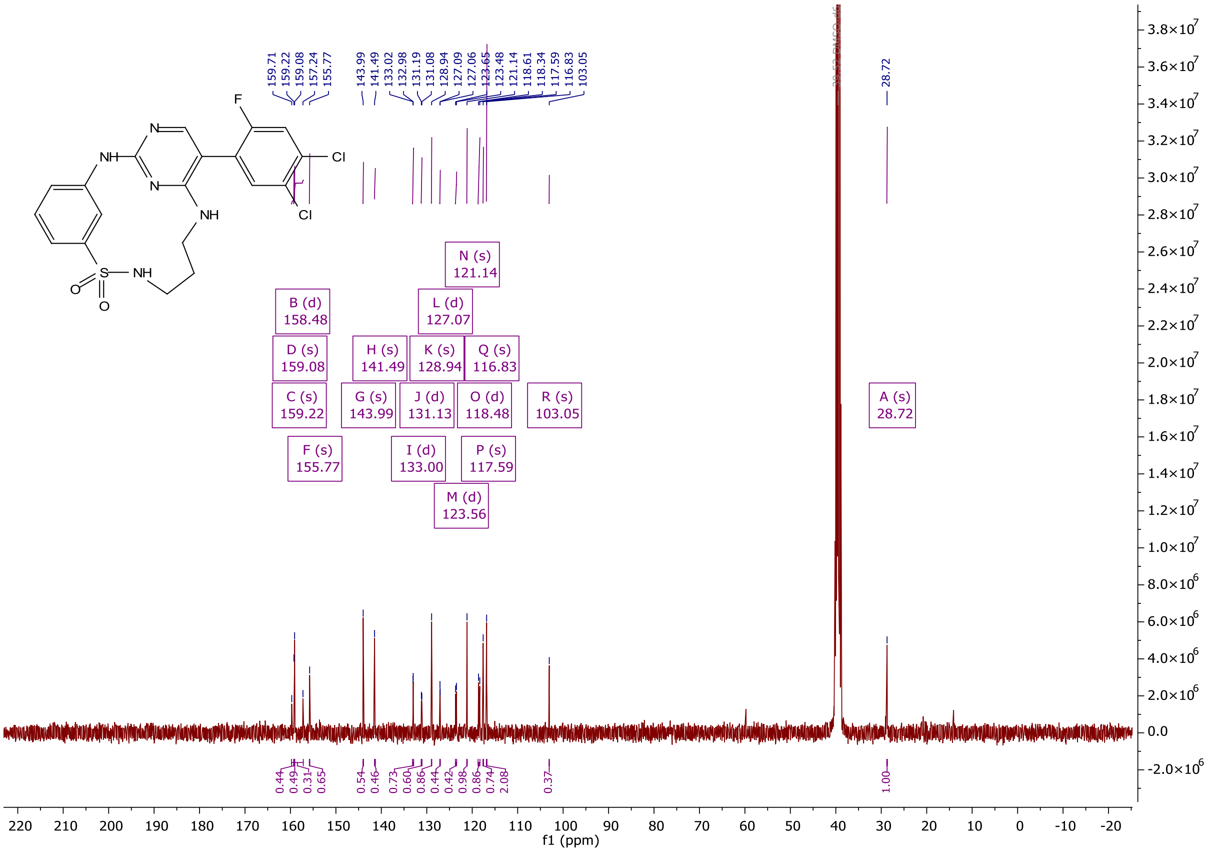
159.71
159.22
159.08
157.24
155.77
143.99
141.49
133.02
132.98
131.19
131.08
128.94
127.09
127.06
123.65
123.48
121.14
118.61
118.34
117.59
116.83
103.05
28.72
F
N
NH
N
Cl
Cl
NH
S
NH
O
O
N (s)
121.14
B (d)
158.48
L (d)
127.07
D (s)
159.08
H (s)
141.49
K (s)
128.94
Q (s)
116.83
C (s)
159.22
G (s)
143.99
J (d)
131.13
O (d)
118.48
R (s)
103.05
A (s)
28.72
F (s)
155.77
I (d)
133.00
P (s)
117.59
M (d)
123.56
f1 (ppm)

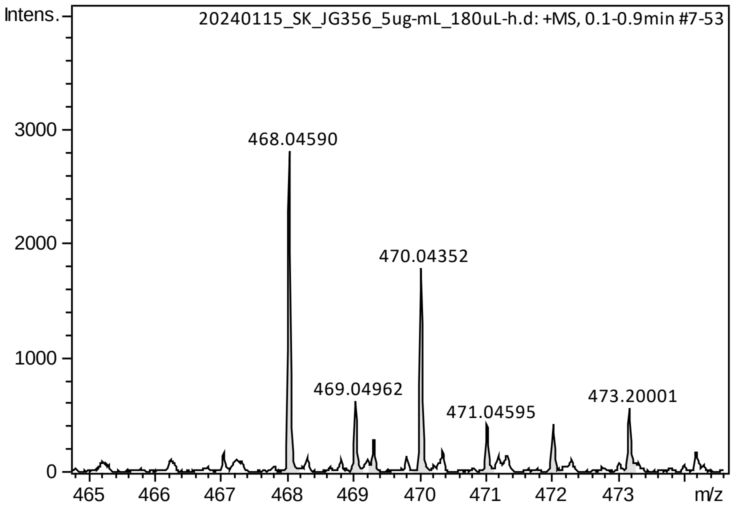
Intens.
20240115_SK_JG356_5ug-mL_180uL-h.d: +MS, 0.1-0.9min #7-53
468.04590
470.04352
469.04962
471.04595
473.20001
m/z

HPLC/MS (ESI)

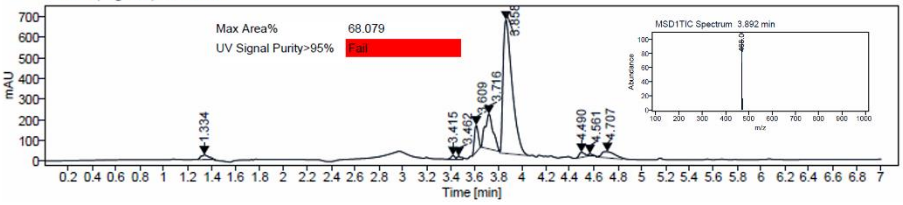
DAD1 C, Sig=320,150 Ref=off
Max Area%
68.079
UV Signal Purity>95%
Fail
MSD1TIC Spectrum 3.892 min
mAU
Time [min]

$^{1}H$ and $^{13}C$ {H} NMR, HRMS (Orbitrap) and HPLC/MS (ESI) data of compound **73**

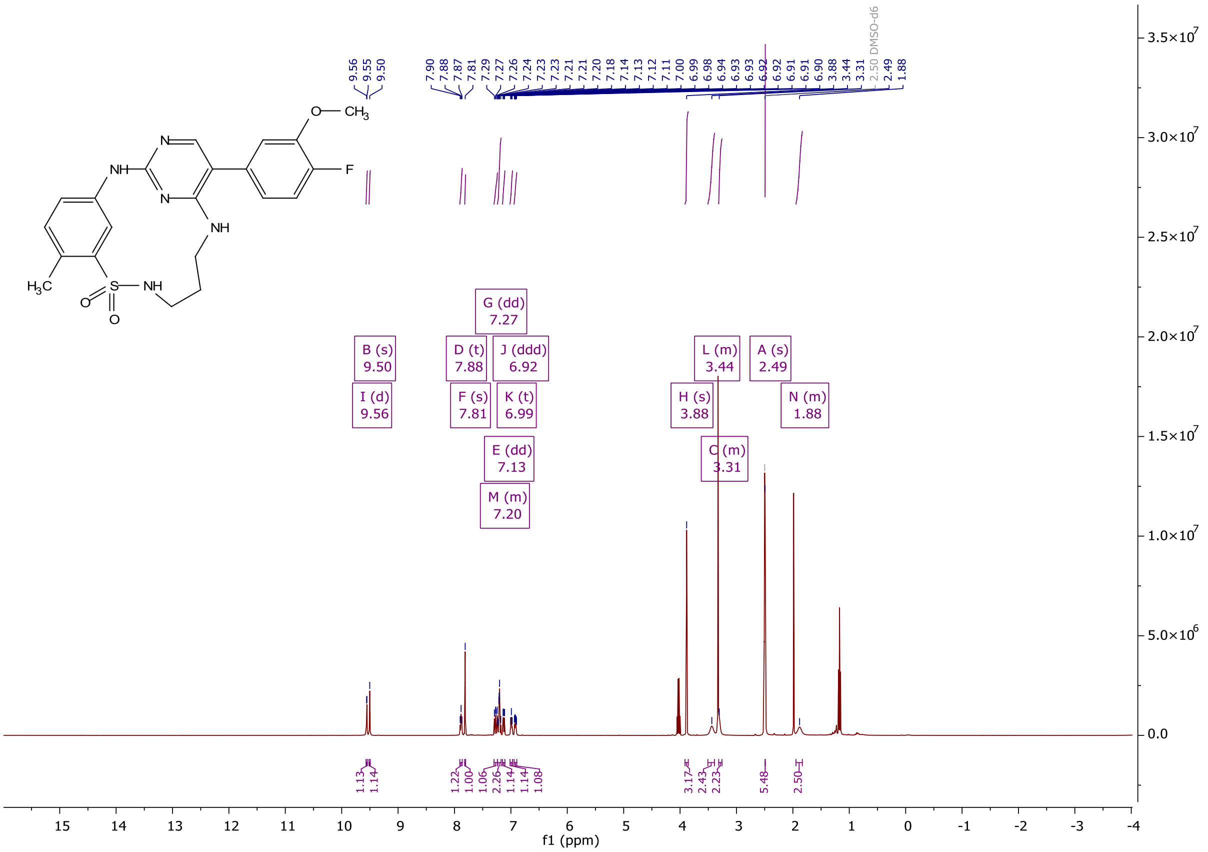

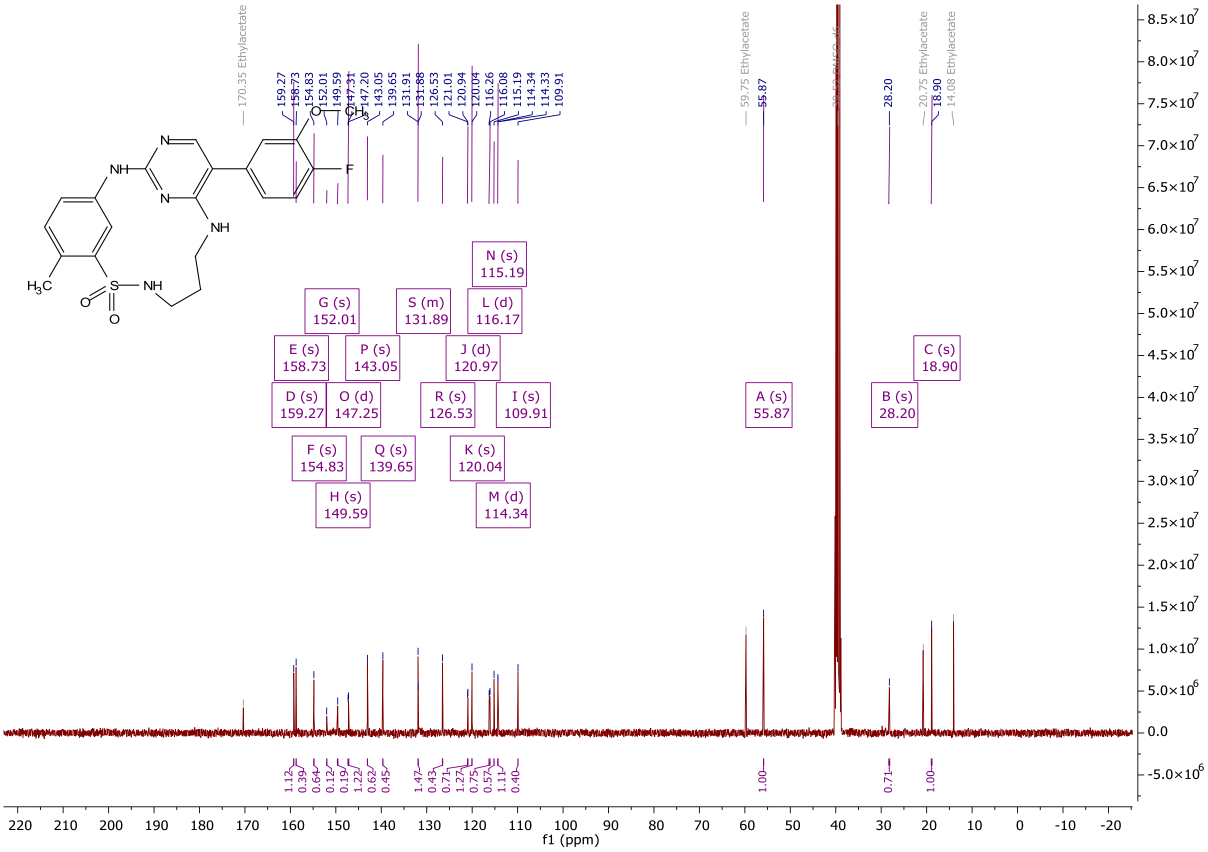

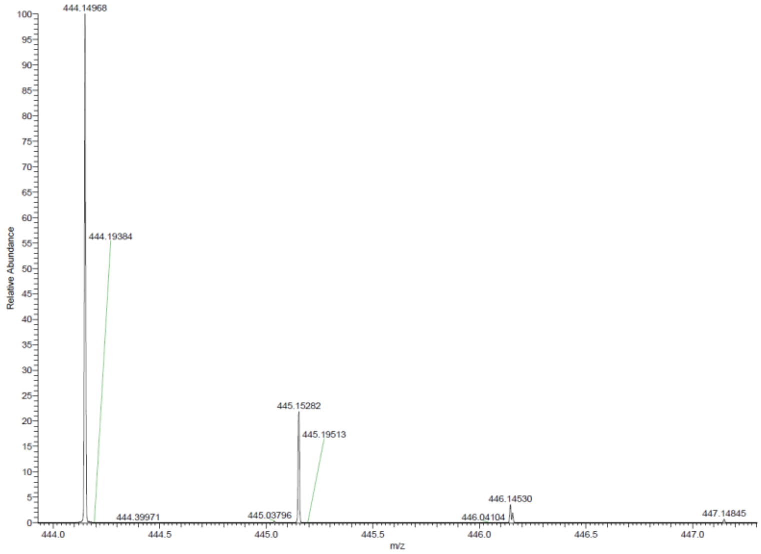

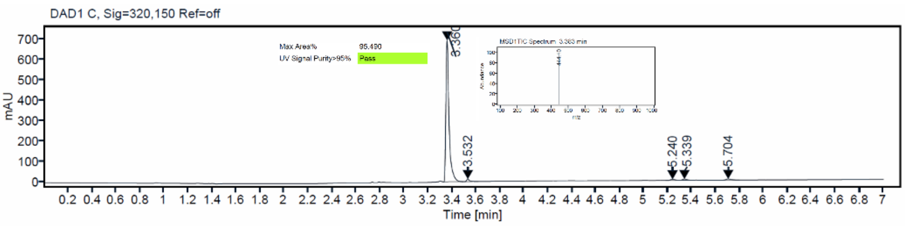

$^1$H and $^{13}$C {H} NMR, HRMS (Orbitrap) and HPLC/MS (ESI) data of compound **74**

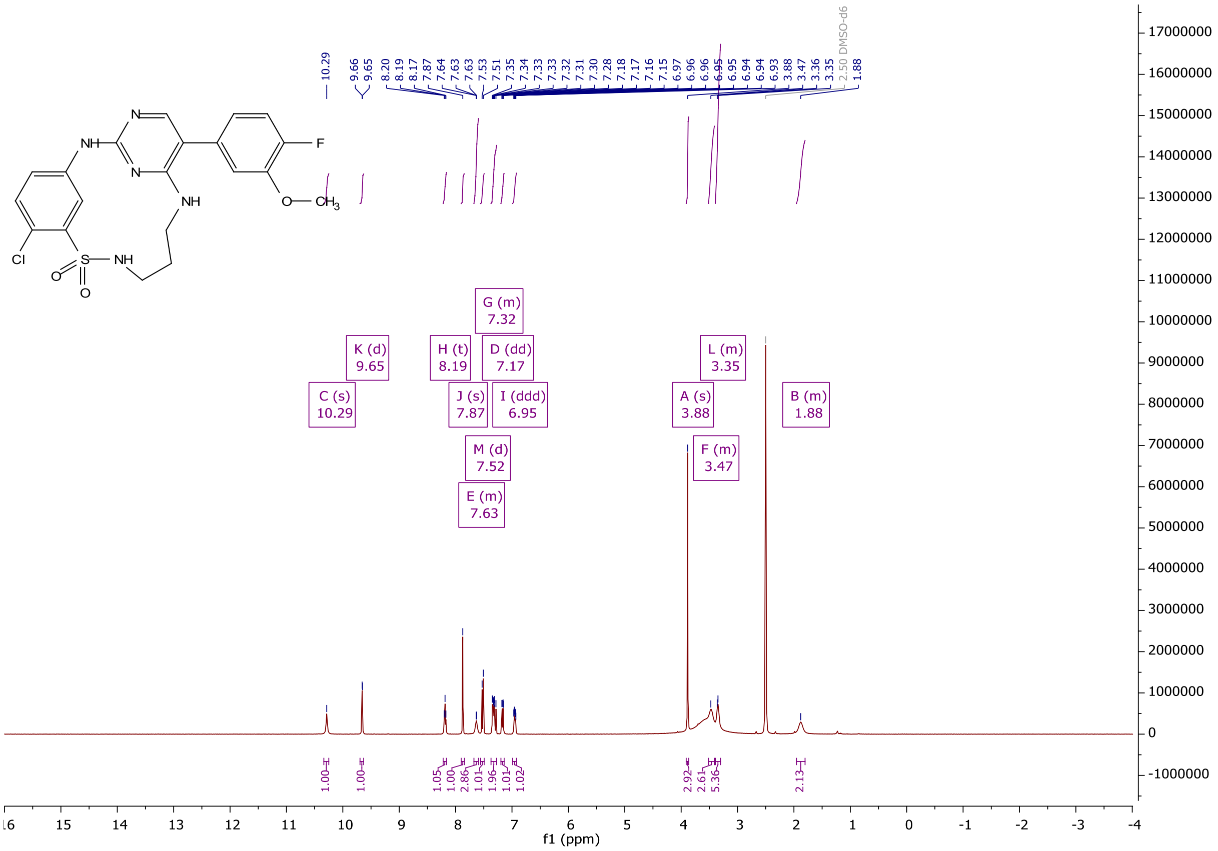

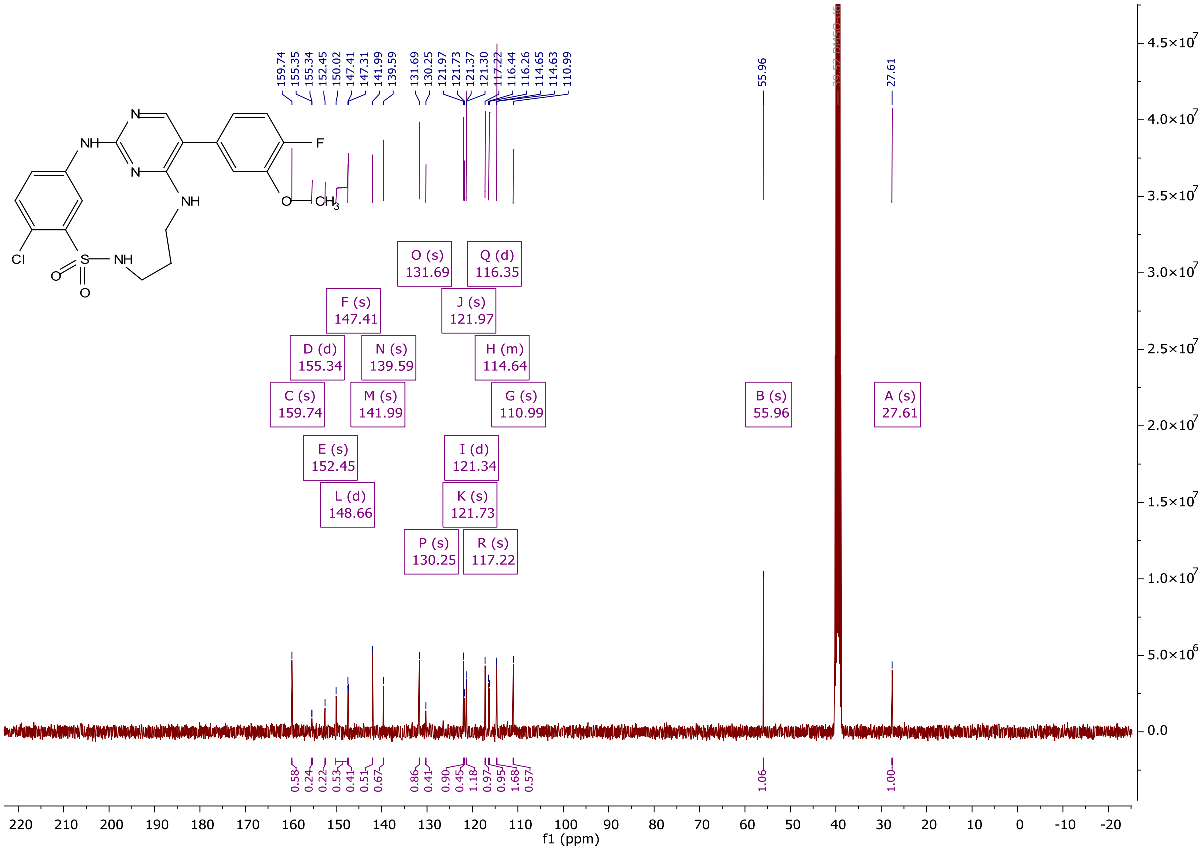

159.74
155.35
155.34
152.45
150.02
147.41
147.31
141.99
139.59
131.69
130.25
121.97
121.73
121.37
121.30
117.22
116.44
116.26
114.65
114.63
110.99
55.96
27.61
O (s) 131.69
Q (d) 116.35
F (s) 147.41
J (s) 121.97
D (d) 155.34
N (s) 139.59
H (m) 114.64
C (s) 159.74
M (s) 141.99
G (s) 110.99
B (s) 55.96
A (s) 27.61
E (s) 152.45
I (d) 121.34
L (d) 148.66
K (s) 121.73
P (s) 130.25
R (s) 117.22
0.58
0.24
0.22
0.53
0.41
0.51
0.67
0.86
0.41
0.90
0.45
1.18
0.97
0.95
1.68
0.57
1.06
1.00
f1 (ppm)

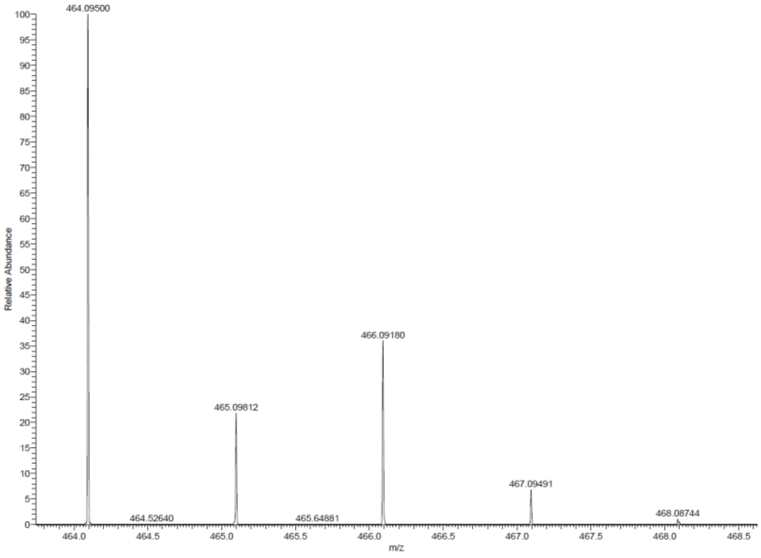

464.09500
465.09812
466.09180
467.09491
468.08744
464.52640
465.64881
Relative Abundance
m/z

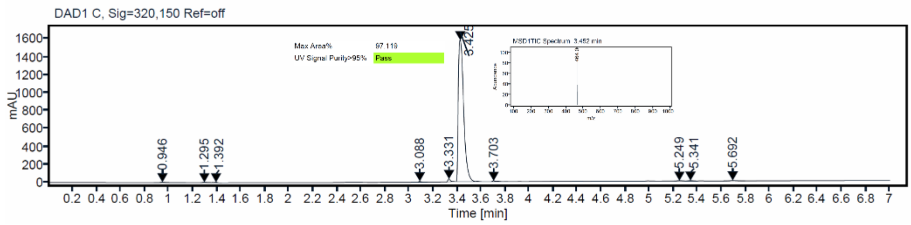

DAD1 C, Sig=320,150 Ref=off
Max Area%
97.119
UV Signal Purity>95%
Pass
mAU
0.946
1.295
1.392
3.088
3.331
3.425
3.703
5.249
5.341
5.692
Time [min]

[1]H and [13]C {H} NMR, HRMS (Orbitrap) and HPLC/MS (ESI) data of compound **75**

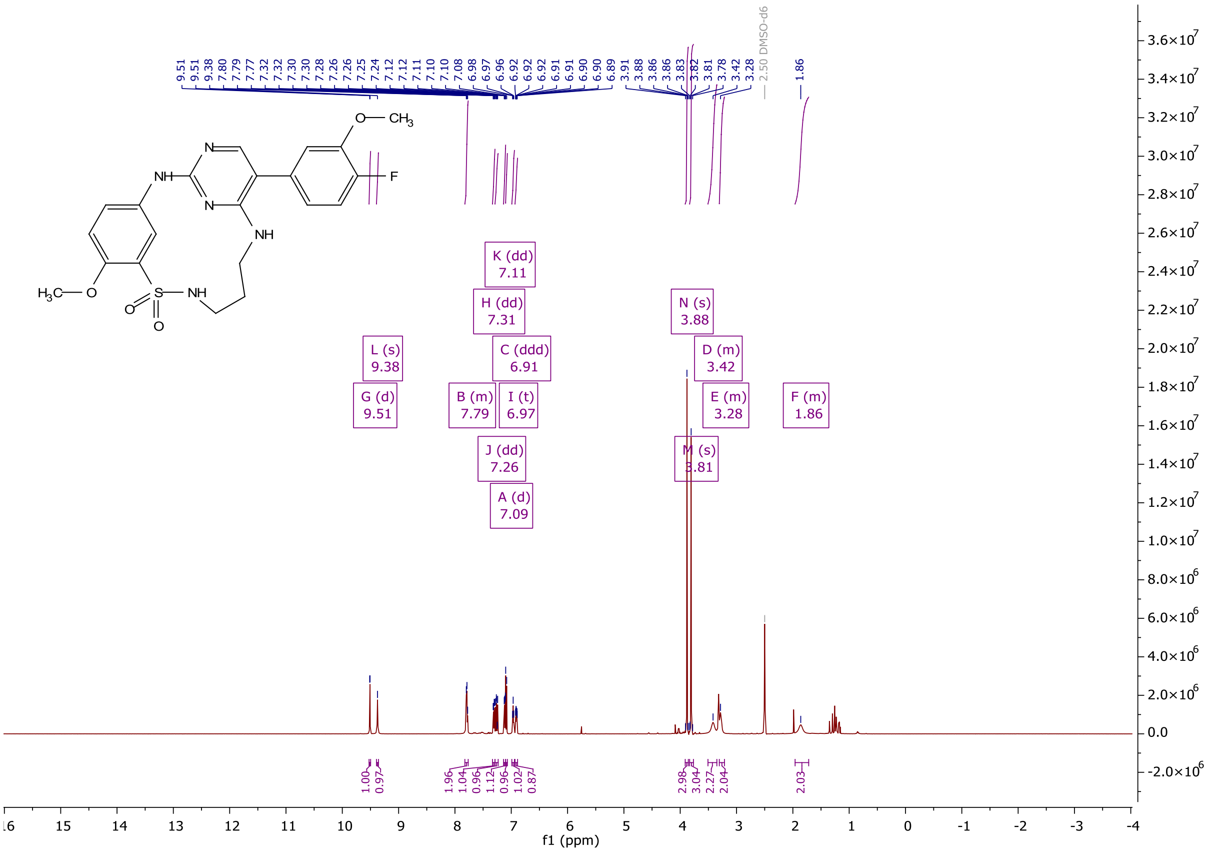

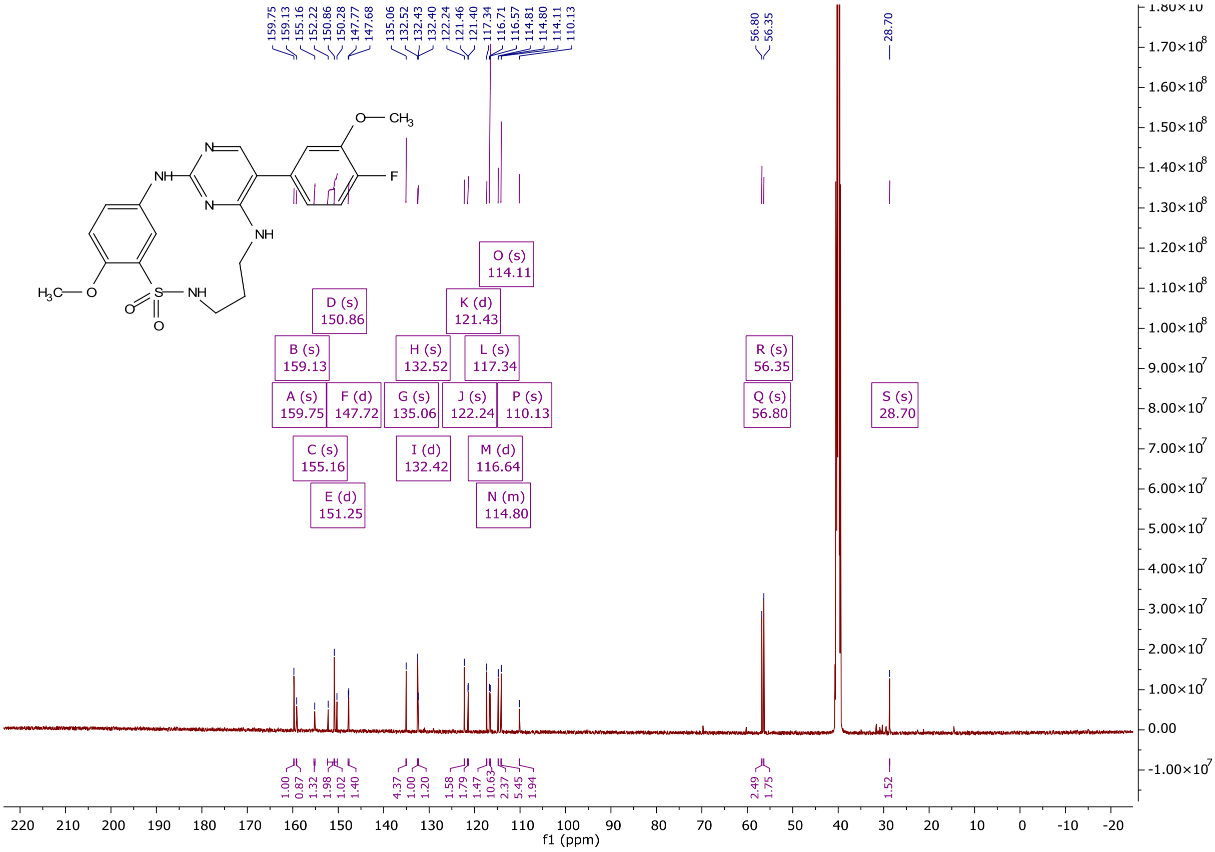

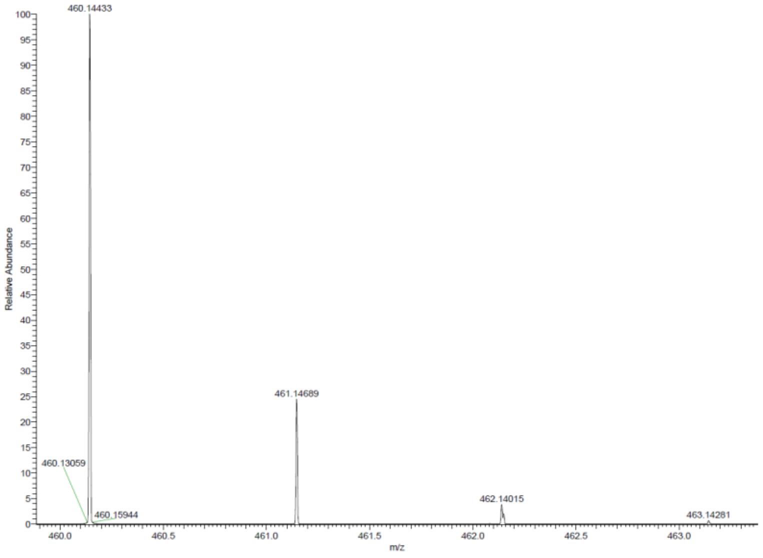

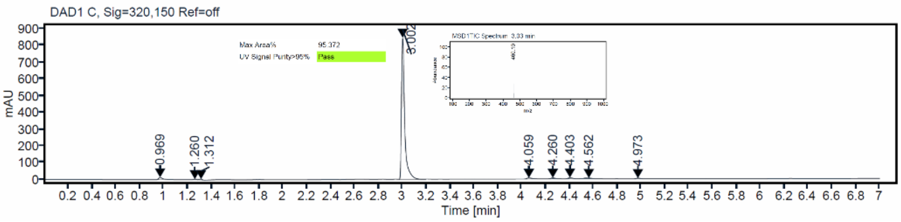

$^{1}H$ and $^{13}C$ {H} NMR, HRMS (oTOF) and HPLC/MS (ESI) data of compound **76**

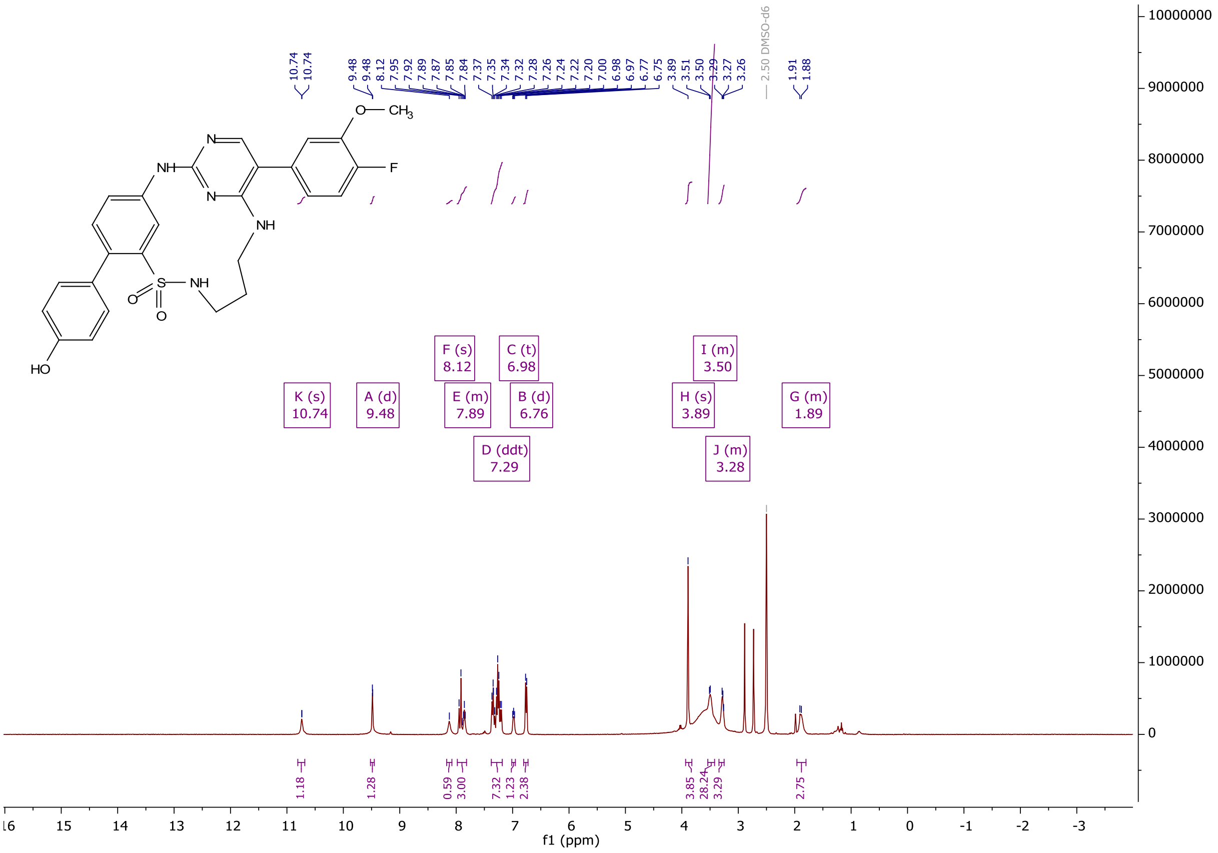

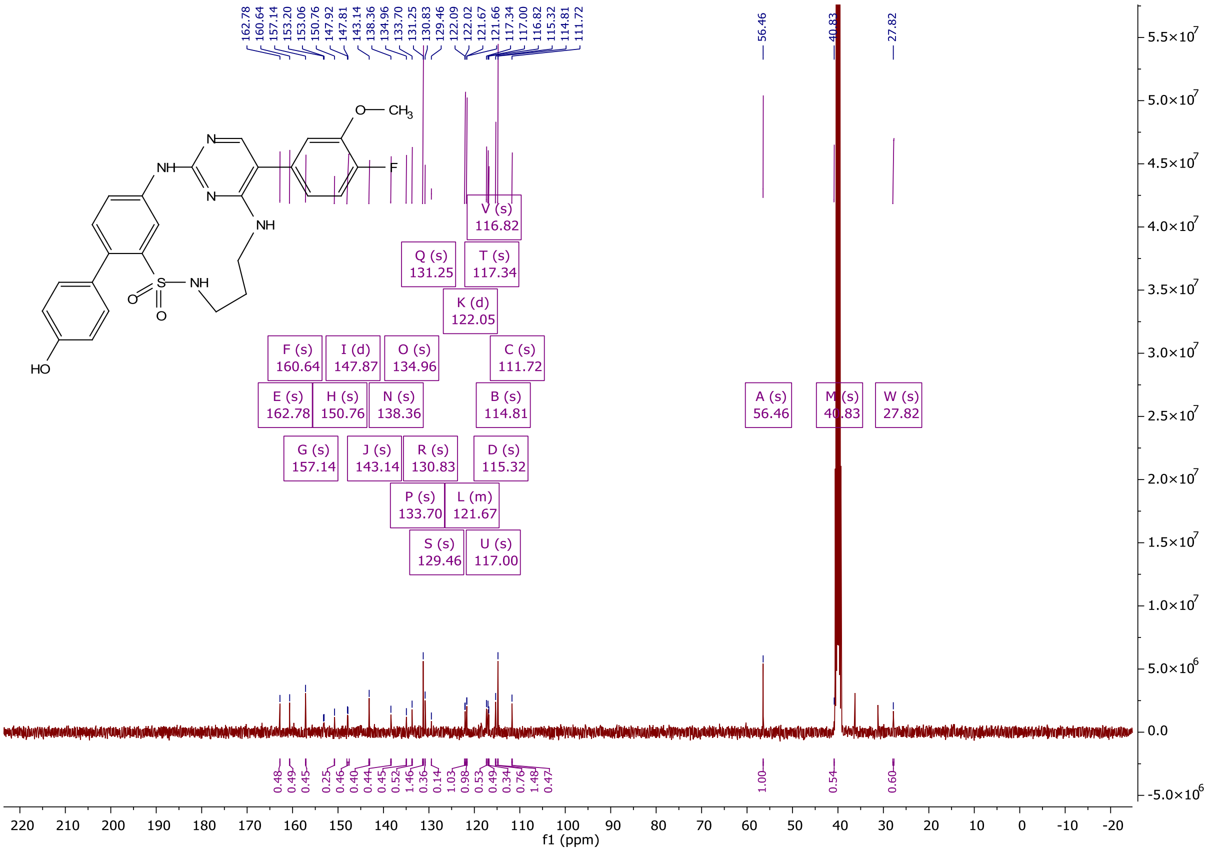

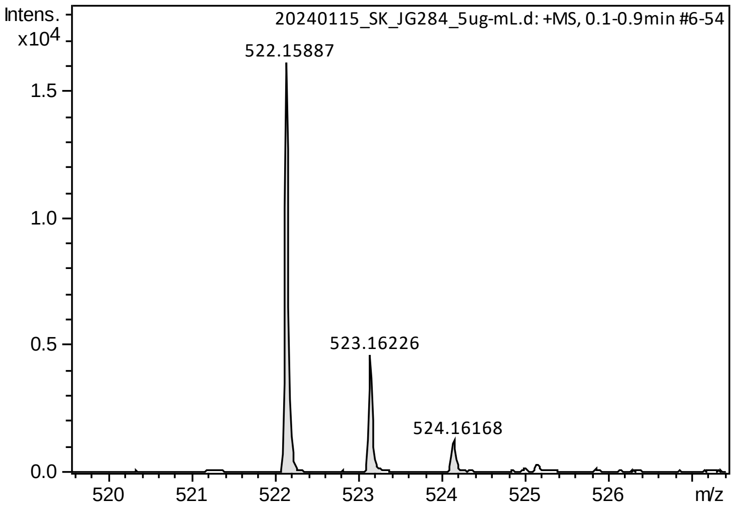
Intens.
x10^4
20240115_SK_JG284_5ug-mL.d: +MS, 0.1-0.9min #6-54
522.15887
523.16226
524.16168
m/z

HPLC/MS (ESI)

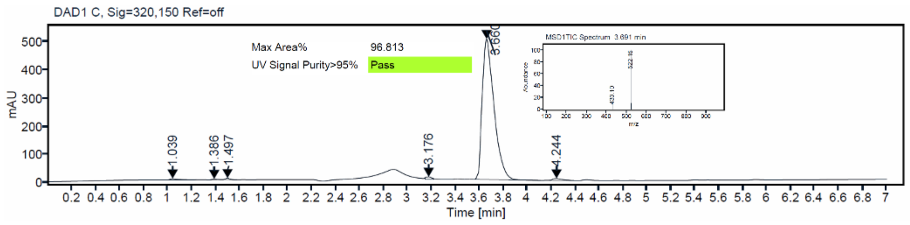
DAD1 C, Sig=320,150 Ref=off
Max Area% 96.813
UV Signal Purity>95% Pass
mAU
Time [min]

$^{1}H$ and $^{13}C$ {H} NMR, HRMS (oTOF) and HPLC/MS (ESI) data of compound **78**

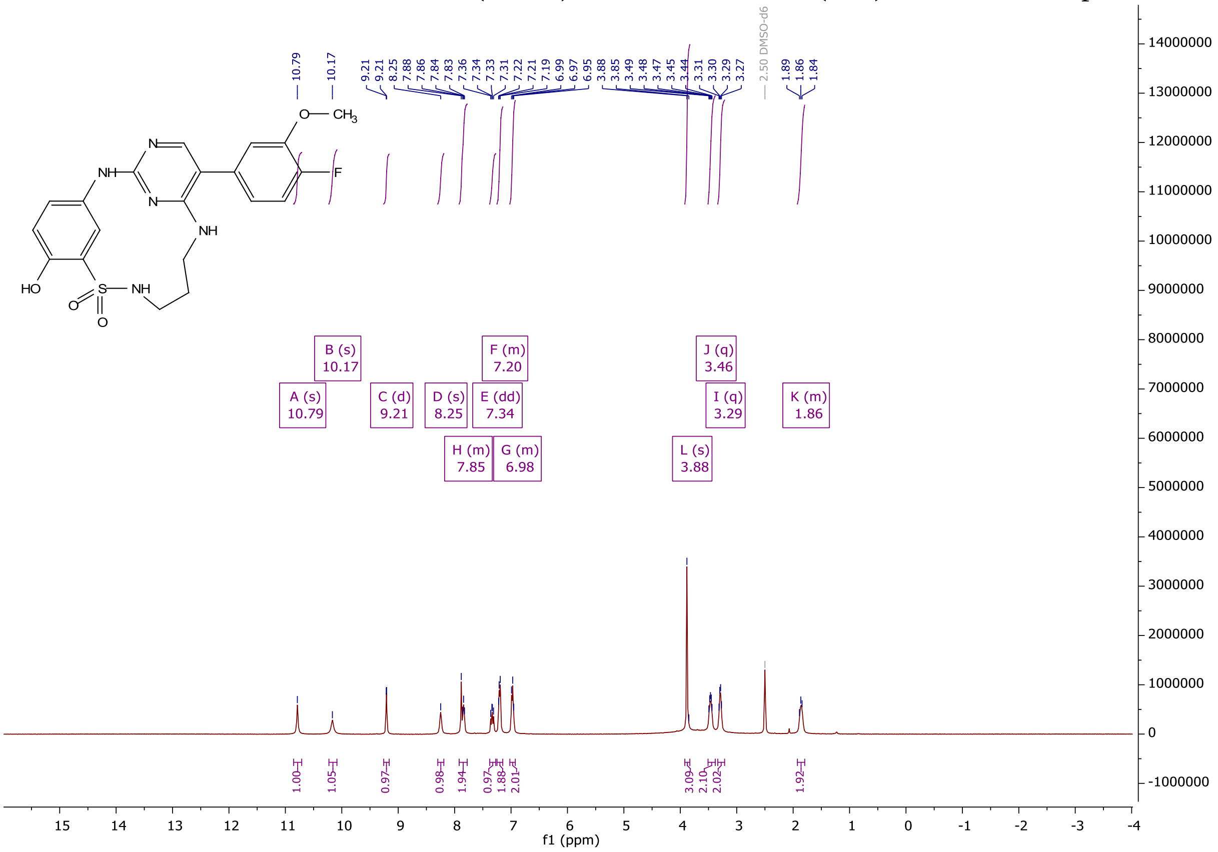

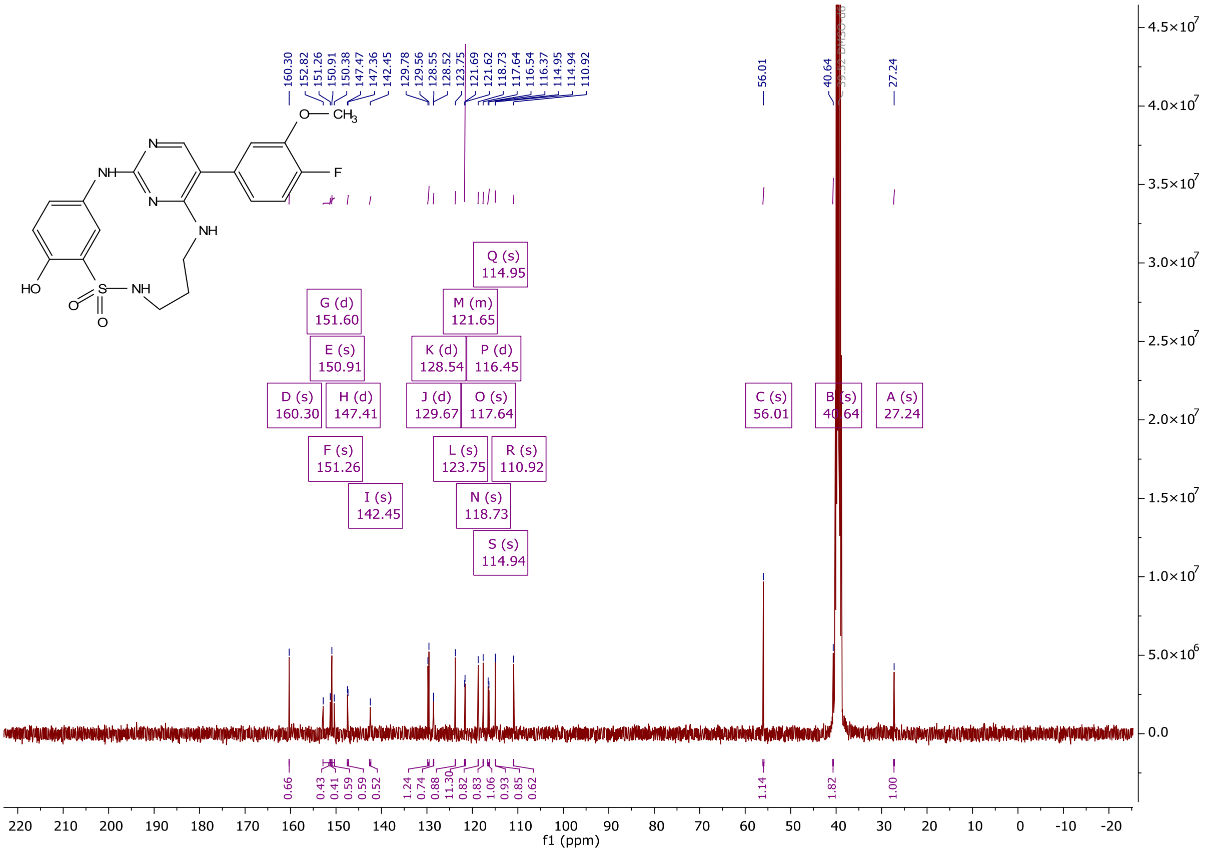

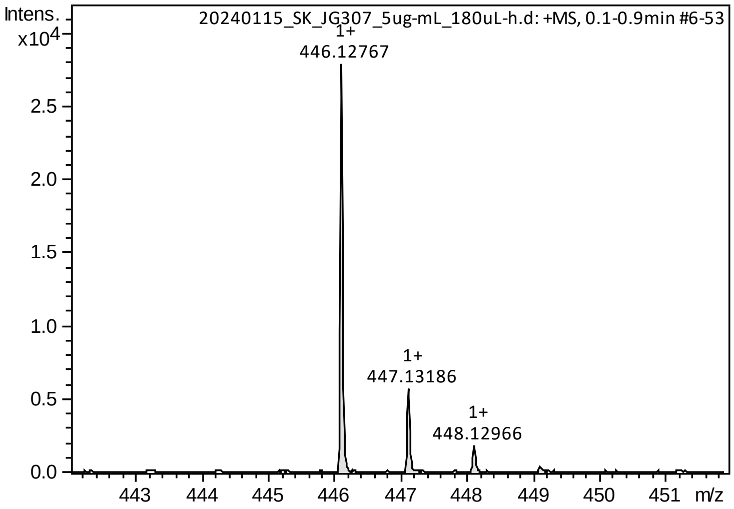

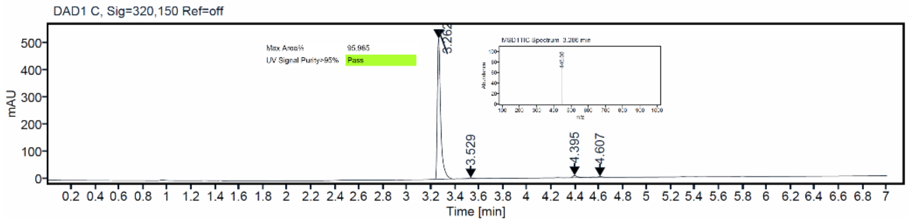

$^{1}$H and $^{13}$C {H} NMR, HRMS (Orbitrap) and HPLC/MS (ESI) data of compound **79**

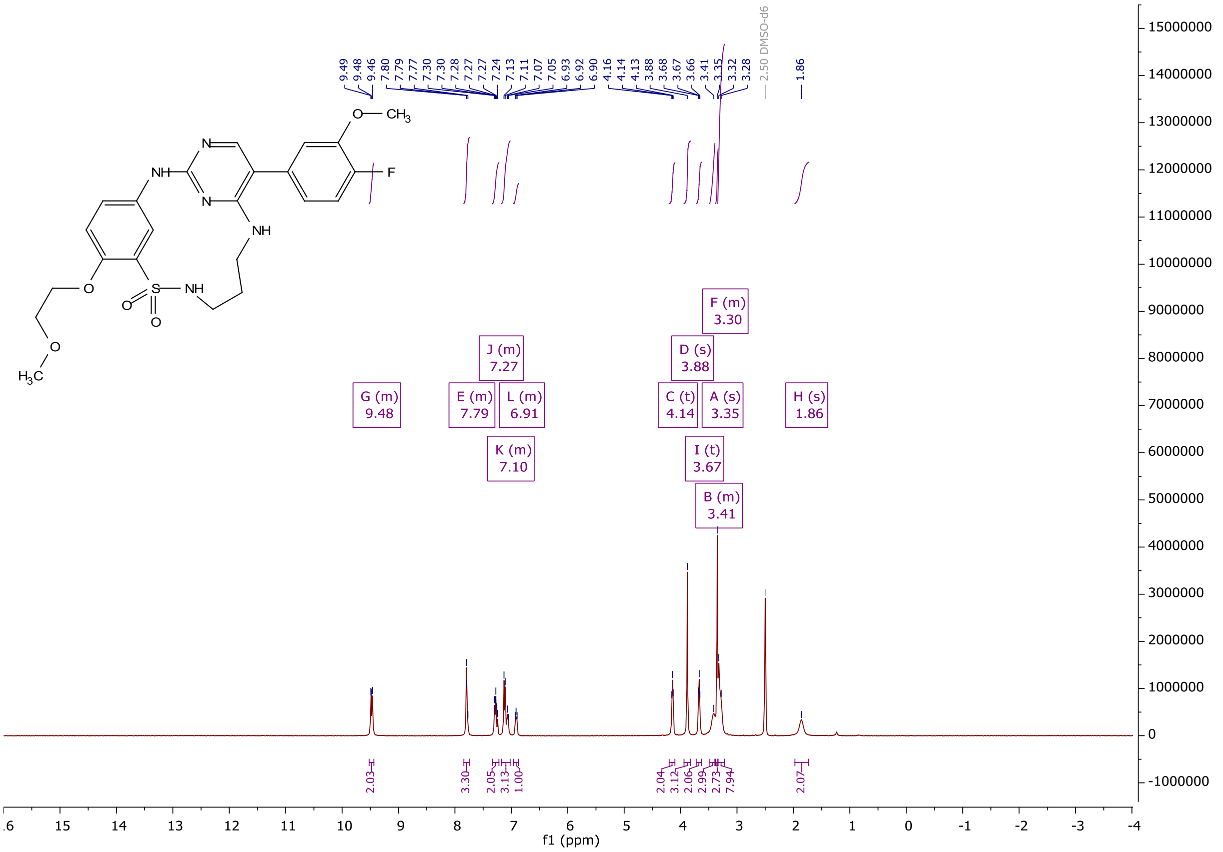

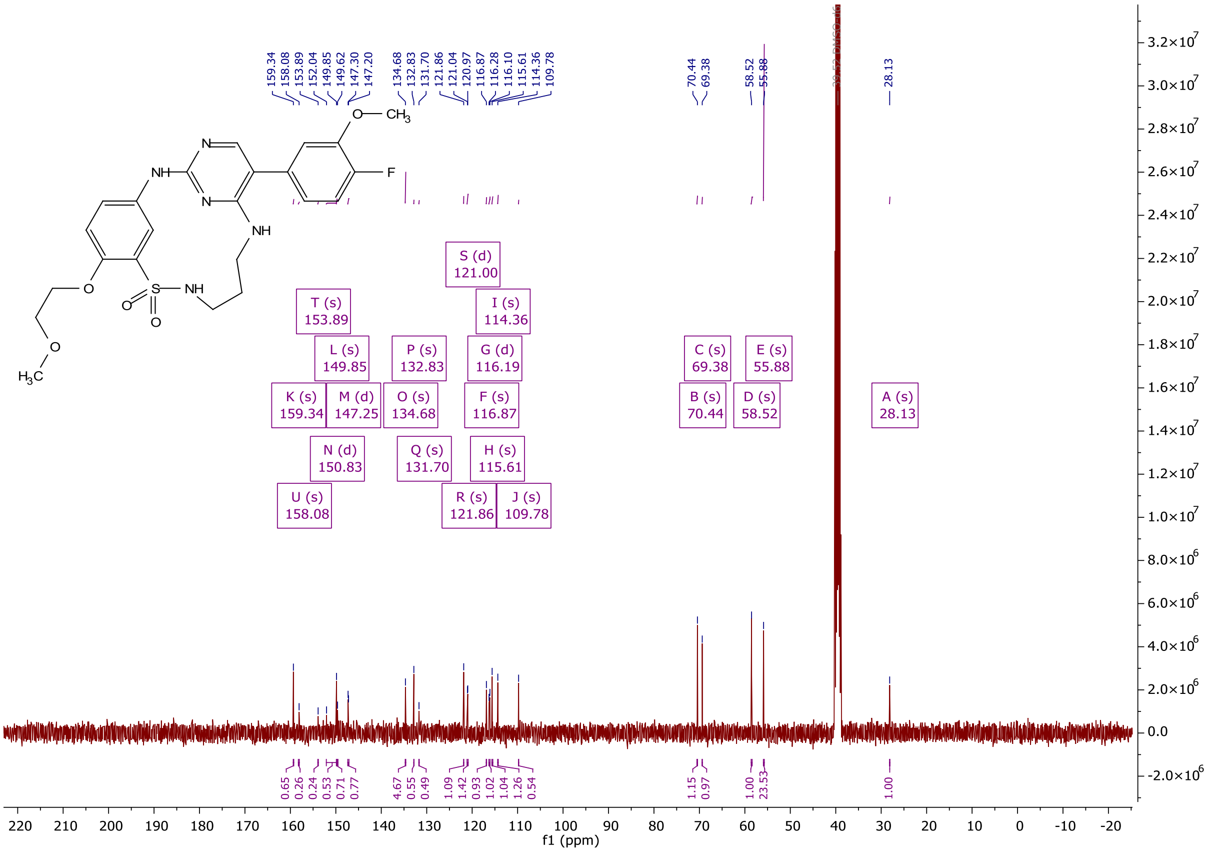
159.34
158.08
153.89
152.04
149.85
149.62
147.30
147.20
134.68
132.83
131.70
121.86
121.04
120.97
116.87
116.28
116.10
115.61
114.36
109.78
70.44
69.38
58.52
55.88
28.13
S (d) 121.00
T (s) 153.89
I (s) 114.36
L (s) 149.85
P (s) 132.83
G (d) 116.19
C (s) 69.38
E (s) 55.88
K (s) 159.34
M (d) 147.25
O (s) 134.68
F (s) 116.87
B (s) 70.44
D (s) 58.52
A (s) 28.13
N (d) 150.83
Q (s) 131.70
H (s) 115.61
U (s) 158.08
R (s) 121.86
J (s) 109.78
f1 (ppm)

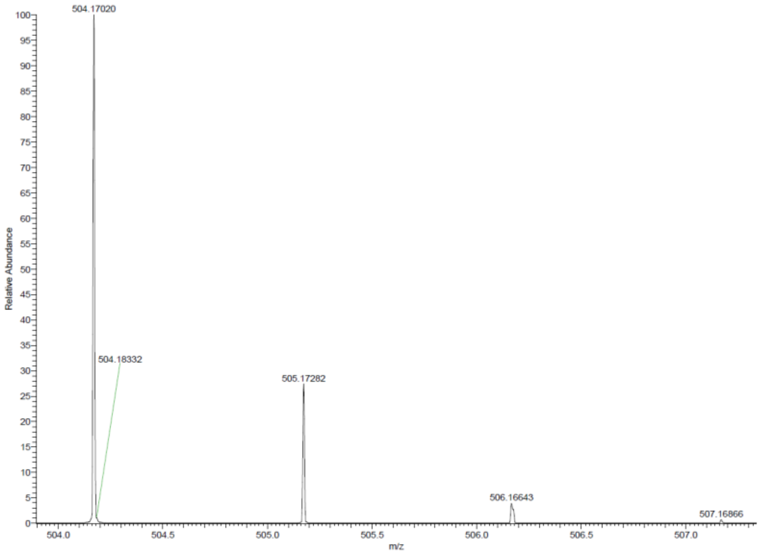
504.17020
504.18332
505.17282
506.16643
507.16866
Relative Abundance
m/z

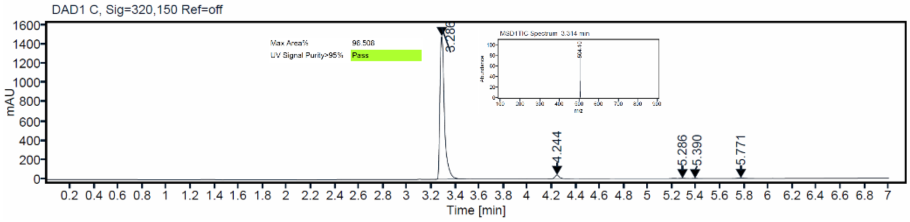
DAD1 C, Sig=320,150 Ref=off
Max Area% 98.508
UV Signal Purity>95% Pass
mAU
3.286
4.244
5.286
5.390
5.771
Time [min]

### $^{1}H$ and $^{13}C$ {H} NMR, HRMS (oTOF) and HPLC/MS (ESI) data of compound **80**

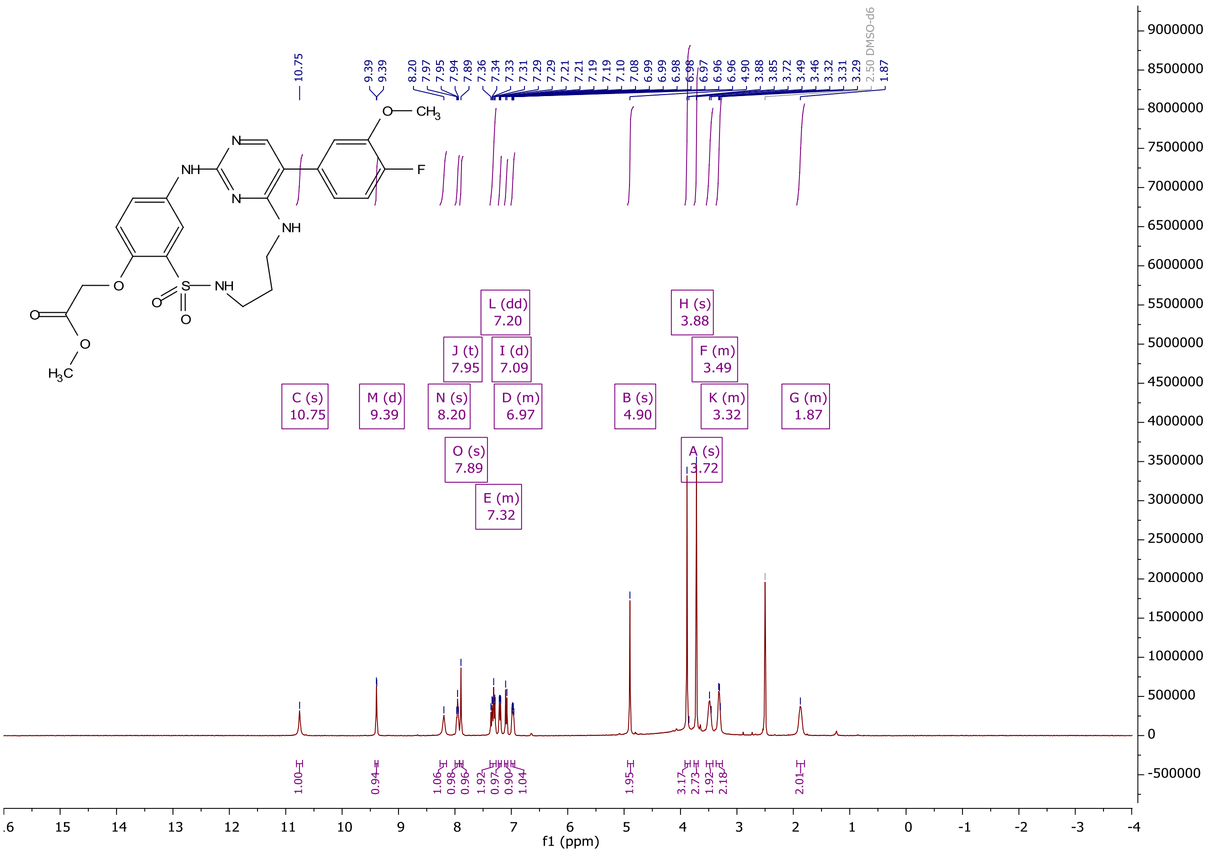

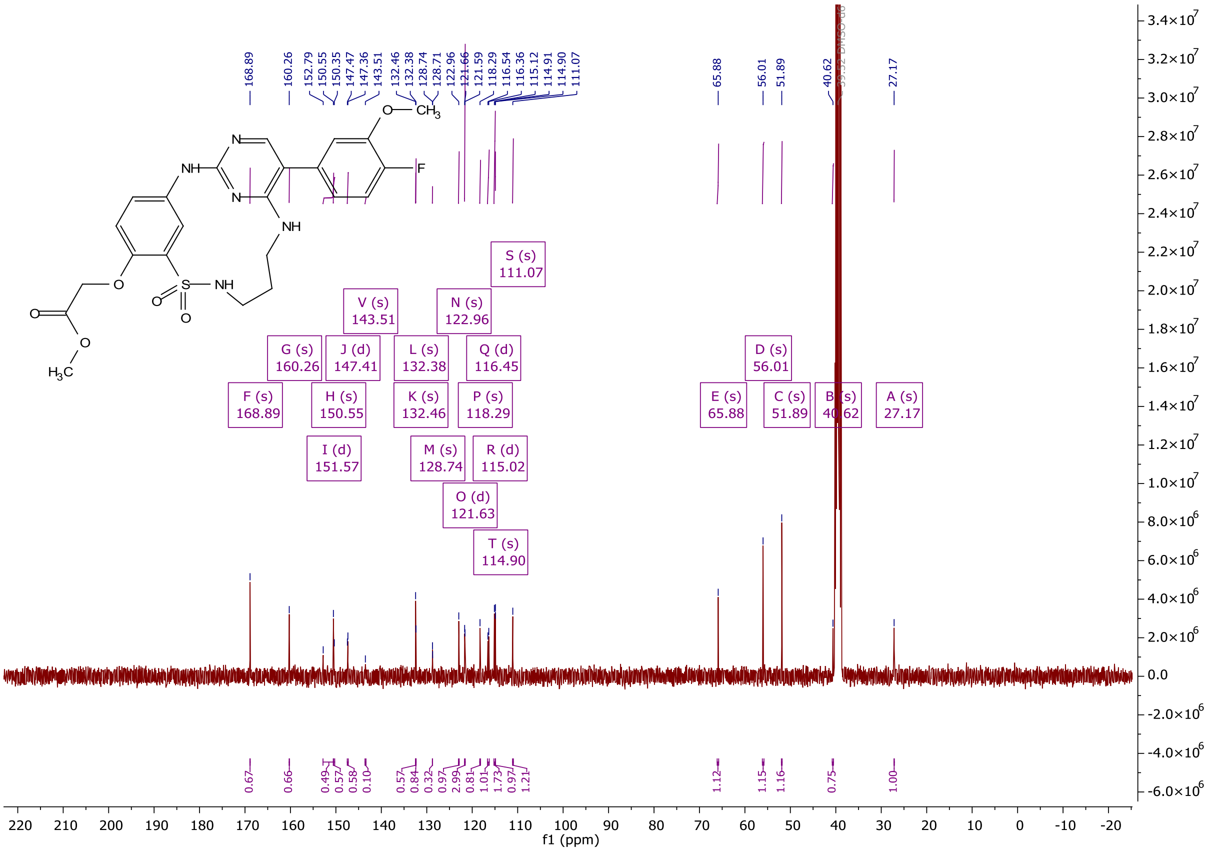

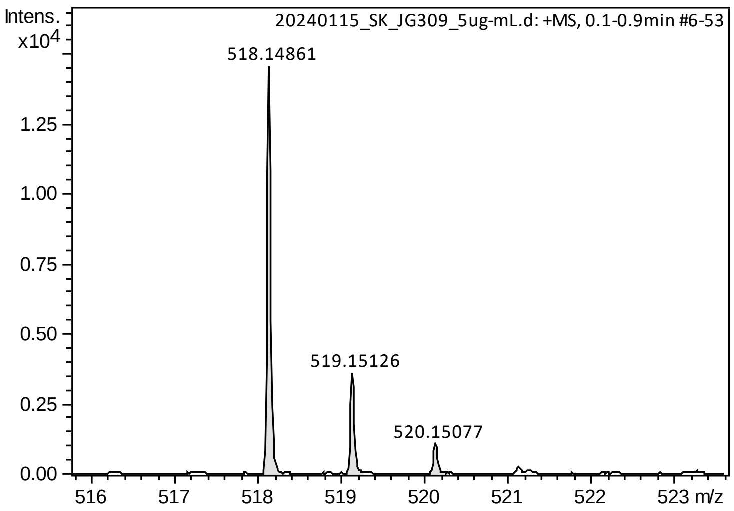

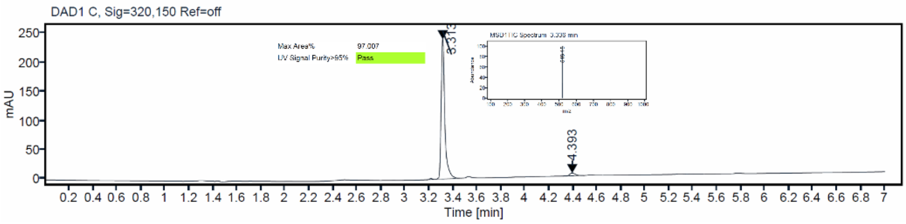

[1]H and [13]C {H} NMR, HRMS (oTOF) and HPLC/MS (ESI) data of compound **81**

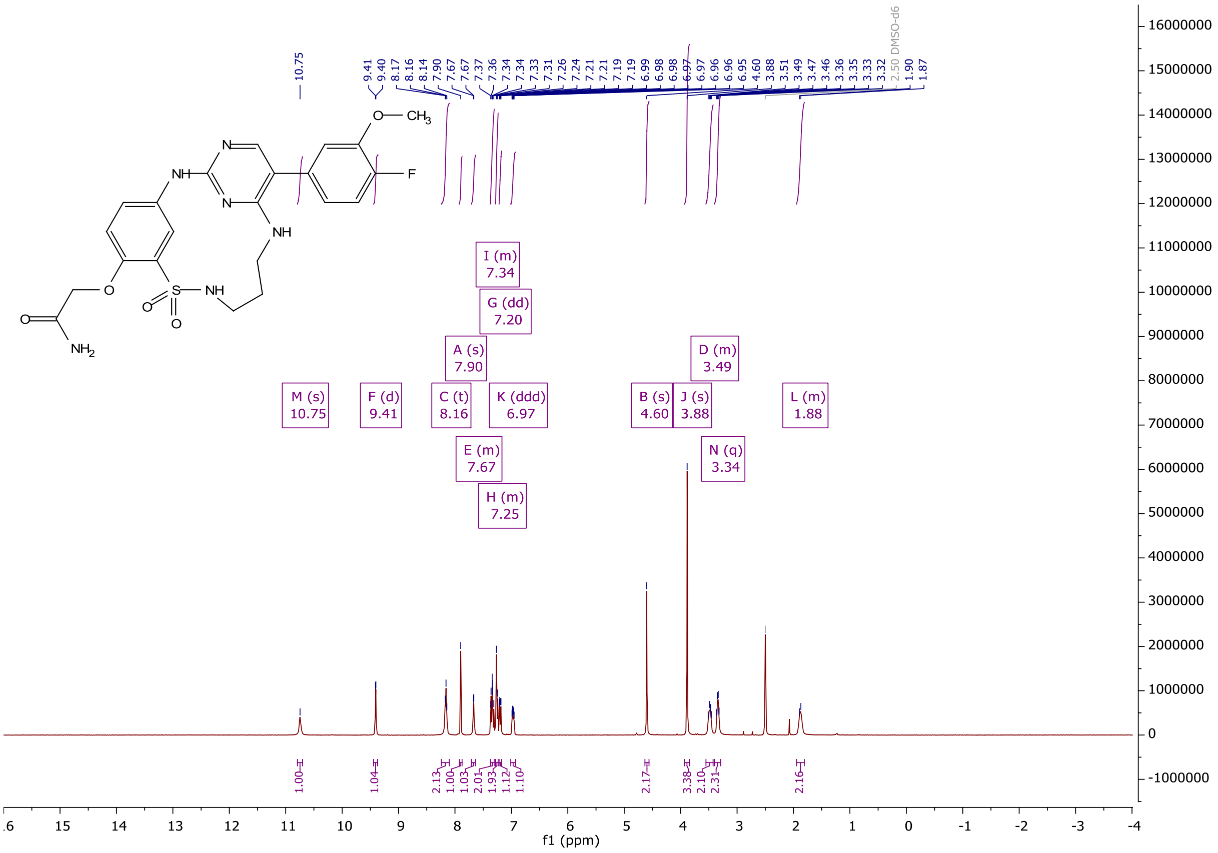

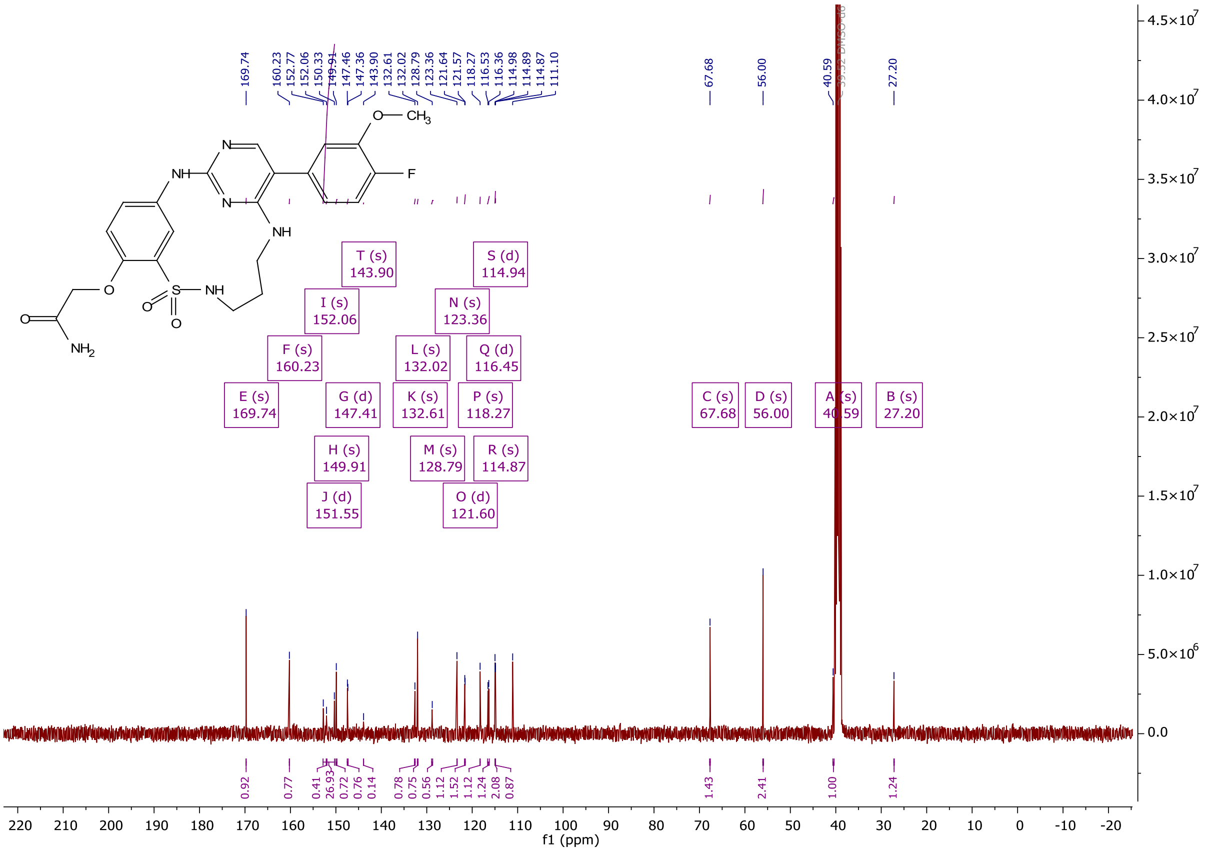

169.74
160.23
152.77
152.06
150.33
149.91
147.46
147.36
143.90
132.61
132.02
128.79
123.36
121.64
121.57
118.27
116.53
116.36
114.98
114.89
114.87
111.10
67.68
56.00
40.59
27.20
O— CH3
F
NH
N
NH
O
S— NH
O
NH2
T (s) 143.90
S (d) 114.94
I (s) 152.06
N (s) 123.36
F (s) 160.23
L (s) 132.02
Q (d) 116.45
E (s) 169.74
G (d) 147.41
K (s) 132.61
P (s) 118.27
C (s) 67.68
D (s) 56.00
B (s) 27.20
H (s) 149.91
M (s) 128.79
R (s) 114.87
J (d) 151.55
O (d) 121.60
0.92
0.77
0.41
26.93
0.72
0.76
0.14
0.78
0.75
0.56
1.12
1.52
1.12
1.24
2.08
0.87
1.43
2.41
1.00
1.24
f1 (ppm)

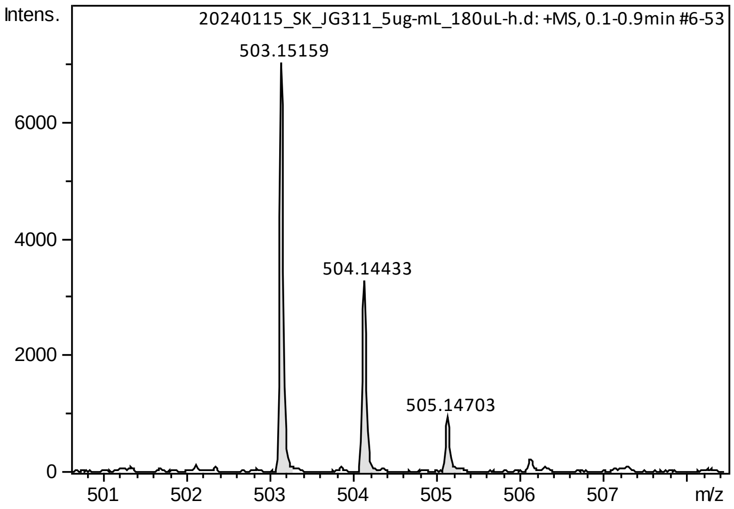

Intens.
20240115_SK_JG311_5ug-mL_180uL-h.d: +MS, 0.1-0.9min #6-53
503.15159
504.14433
505.14703
m/z

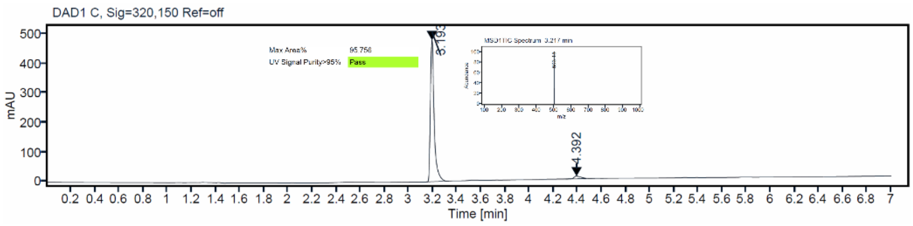

DAD1 C, Sig=320,150 Ref=off
mAU
3.193
4.392
Time [min]

[1]H and [13]C {H} NMR, HRMS (oTOF) and HPLC/MS (ESI) data of compound **82**

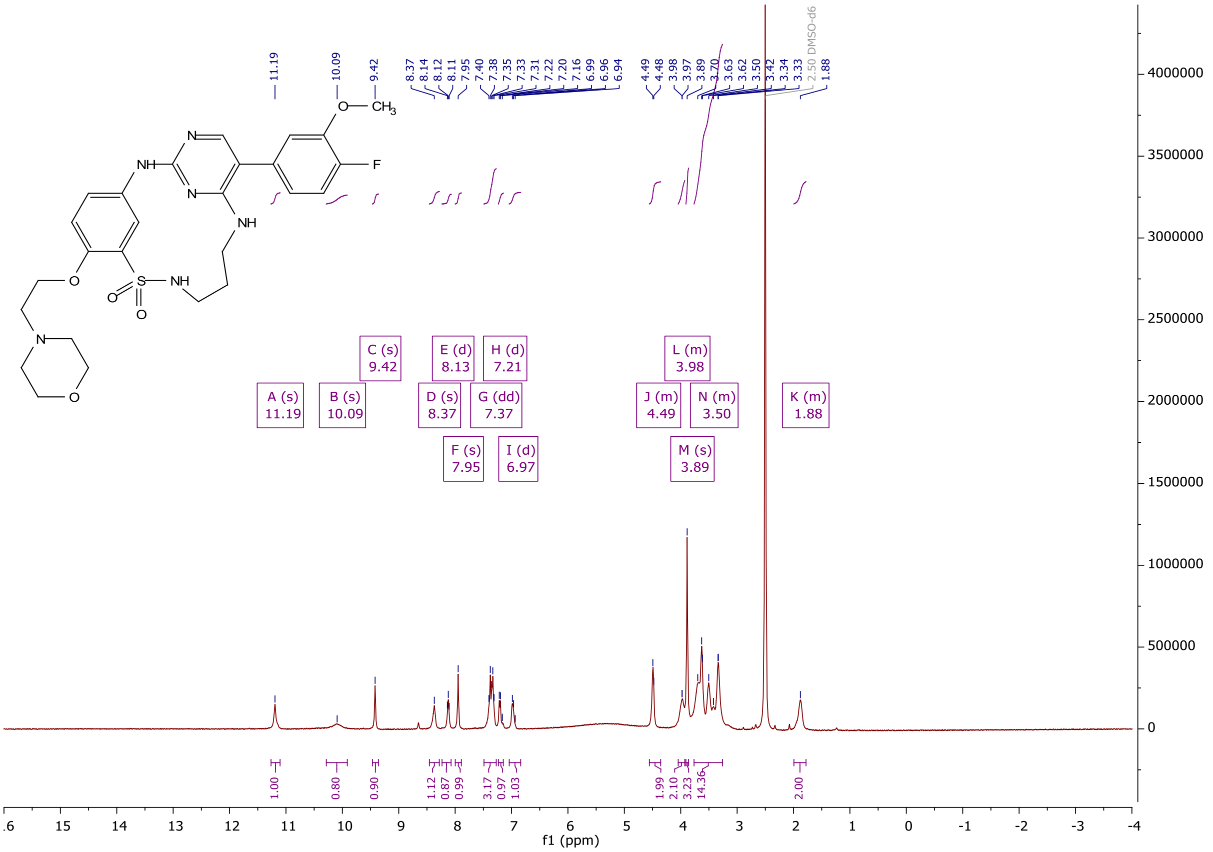

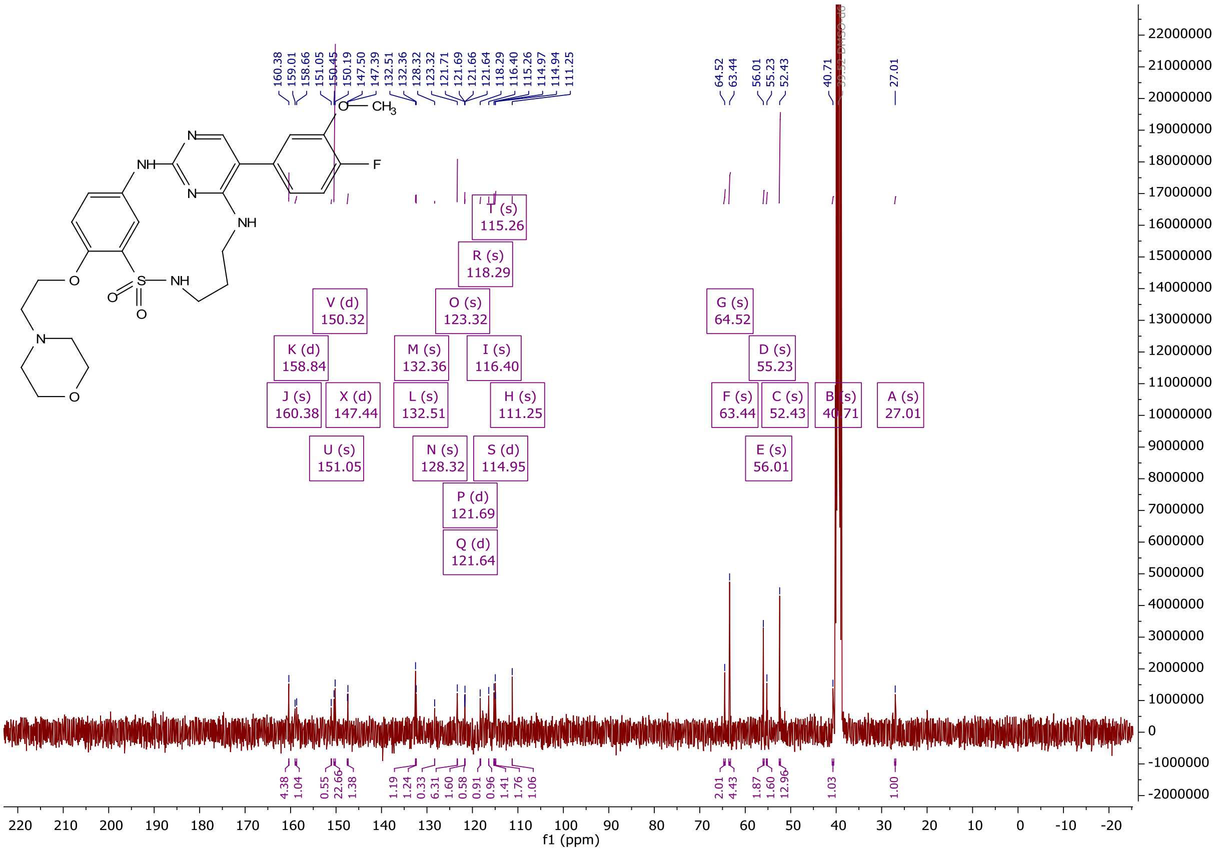

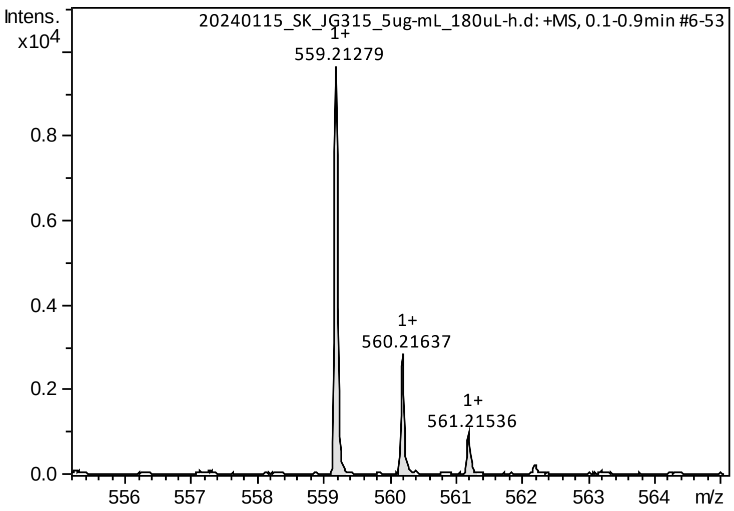

Intens.
x10^4
20240115_SK_JG315_5ug-mL_180uL-h.d: +MS, 0.1-0.9min #6-53
1+
559.21279
1+
560.21637
1+
561.21536
0.8
0.6
0.4
0.2
0.0
556
557
558
559
560
561
562
563
564
m/z

HPLC/MS (ESI)

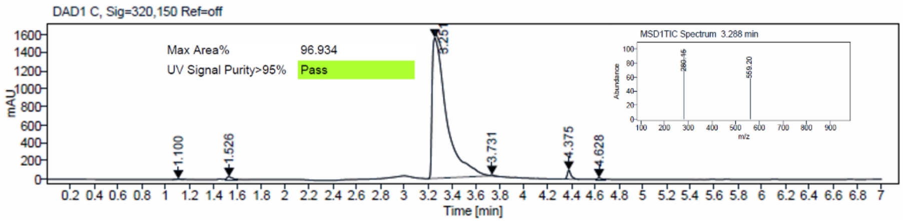

DAD1 C, Sig=320,150 Ref=off
Max Area%
96.934
UV Signal Purity>95%
Pass
mAU
Time [min]
1.100
1.526
3.251
3.731
4.375
4.628
MSD1TIC Spectrum 3.288 min
Abundance
m/z

#### Crystallization

**Table S3.** Data collection and refinement statistics

| **Data collection** | **EPHA2-23** |
|---|---|
| Beamline | X06SA/PXI SLS |
| Wavelength (Å) | 0.99999 |
| Space group | $P2_1$ |
| Cell dimensions | |
| *a, b, c (Å)* | 32.82, 107.68, 40.64 |
| α, β, γ (°) | 90.00, 109.04, 90.00 |
| Resolution (Å)* | 38.38-1.80 (1.84-1.80) |
| unique observations* | 24679 (1466) |
| $R_{pim}$* | 0.048 (0.356) |
| Completeness (%)* | 100 (100) |
| Multiplicity* | 6.9 (6.7) |
| Mean I/σI* | 12.6 (2.2) |
| $CC_{1/2}$* | 0.998 (0.750) |
| **Refinement** | |
| $R_{work}$ / $R_{free}$§ | 18.57 / 22.24 |
| No. of atoms | 2424 |
| Overall B-factors ($Å^2$) | 19.29 |
| Rms deviations | |
| Bond lengths (Å) | 0.007 |
| Bond angles (°) | 1.4 |
| Ramachandran (%) | |
| favored | 98 |
| allowed | 2 |
| outlier | 0 |
| **Protein Data Bank entry** | **8QQY** |

*Values for the highest-resolution shell are shown in parentheses.

§$R_{work}$ and $R_{free}$ = ∑||$F_{obs}$| - |$F_{calc}$||/∑|Fobs|, where $R_{free}$ was calculated with 5% of the reflections chosen at random and not used in the refinement

#### Surface Plasmon Resonance

**Table S4.** Affinity fit and kinetic fit data for SPR experiments with compounds 6, 23, 28, 55, 75 and 79 on EPHA2 and GAK.

| | *affinity fit* | | *kinetic fit* | | | | | | | |
|---|---|---|---|---|---|---|---|---|---|---|
| | **EPHA2** | **GAK** | **EPHA2** | | | | **GAK** | | | |
| **cpd ID** | **$K_D$ [nM]** | **$K_D$ [nM]** | **$K_D$ [nM]** | **$k_a$ [1/Ms]** | **$k_d$ [1/s]** | **τ [s]** | **$K_D$ [nM]** | **$k_a$ [1/Ms]** | **$k_d$ [1/s]** | **τ [s]** |
| **6**[*] | 307.0 ± 3.8 | 112.4 | 250 | 2,24E+06 | 5,67E-01 | 2 | 78 | 1,15E+06 | 8,89E-02 | 11 |
| **23** | 120.4 ± 4.6 | 7.7 | 103 | 8,71E+05 | 8,84E-02 | 11 | 8 | 1,52E+06 | 1,17E-02 | 85 |
| **28**[*] | N/A | 49.7 | N/A | 3,09E+04 | 3,59E-01 | 3 | 41 | 9,59E+05 | 3,97E-02 | 25 |
| **55** | 180.5 ± 3.5 | 19.2 | 166 | 4,22E+05 | 6,87E-02 | 15 | 15 | 7,04E+05 | 1,07E-02 | 93 |
| **75** | 95.3 ± 2.0 | 6.2 | 85 | 1,18E+06 | 9,94E-02 | 10 | 4 | 2,90E+06 | 1,24E-02 | 81 |
| **79** | 126.1 ± 9.9 | 5.0 | 118 | 1,06E+06 | 1,25E-01 | 8 | 4 | 2,24E+06 | 8,95E-03 | 112 |

[*]The kinetic measurement showed a very fast on set for 6 (EPHA2 and GAK) and 28 (EPHA2), which is why the kinetic fit results are on the edge of the device's limitations.

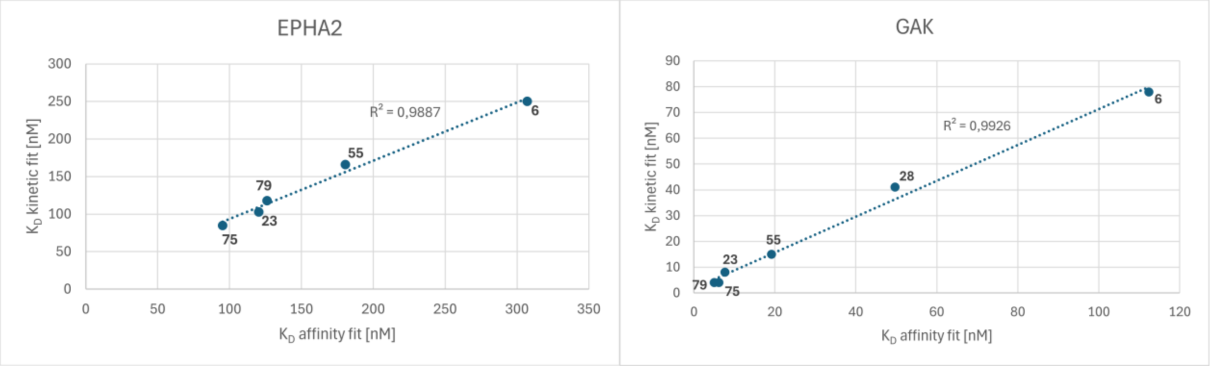

**Figure S1.** Affinity fit $K_D$ plotted against kinetic fit $K_D$ for EPHA2 and GAK. Each compound depicted as blue dot and labelled with compound ID. Linear regression trendline fitted into graph. $R^2$(EPHA2) ≈ 0.99; $R^2$(GAK) ≈ 0.99.

#### NanoBRET

**Table S5.** Specification of NanoBRET assays, which were conducted for this work.

| Protein | Plasmid catalog no. (Promega) | NanoLuc orientation | Tracer | Tracer catalog no. (Promega) | Tracer, concentration used [M] |
|---|---|---|---|---|---|
| EphA1 | NV1221 | C | K4 | N2540 | 1.30E-07 |
| EphA2 | NV1231 | C | K4 | N2540 | 3.10E-08 |
| EphA3 | NV3061 | C | K11 | N2850 | 6.30E-07 |
| EphA4 | NV1241 | C | K4 | N2540 | 1.60E-08 |
| EphA5 | NV1251 | C | K4 | N2540 | 1.60E-08 |
| EphA6 | NV1261 | C | K10 | N2840 | 2.50E-07 |
| EphA7 | NV1271 | C | K10 | N2840 | 5.00E-07 |
| EphA8 | NV1281 | C | K4 | N2540 | 6.30E-09 |
| EphA10 | kind gift of Promega | C | K4 | N2540 | 6.30E-08 |
| EphB1 | NV3071 | C | K9 | N2830 | 6.60E-07 |
| EphB2 | NV1291 | C | K4 | N2540 | 1.30E-08 |
| EphB3 | NV1301 | C | K4 | N2540 | 1.30E-07 |
| EphB4 | NV1311 | C | K4 | N2540 | 6.30E-09 |
| EphB6 | kind gift of Promega | C | K4 | N2540 | 1.00E-06 |
| ABL1 | NV1011 | N | K4 | N2540 | 1.30E-07 |
| FGFR1 | NV1341 | C | K10 | N2840 | 2.50E-07 |
| FLT1 | NV3111 | C | K9 | N2830 | 3.30E-07 |
| GAK | NV1421 | N | K10 | N2840 | 3.10E-08 |
